# AP2/ERF transcription factors enriched in the drought-response transcriptome of the Thar desert tree *Prosopis cineraria* show higher copy number and greater DNA-binding affinity than orthologs in drought-sensitive species

**DOI:** 10.1101/2024.06.17.599320

**Authors:** Vedikaa Dhiman, Debankona Marik, Amrita, Rajveer Singh Shekhawat, Asish Kumar Swain, Arpan Dey, Pankaj Yadav, Arumay Pal, Sucharita Dey, Ayan Sadhukhan

**Author notes:** Correspondence (PY), (AP), (SD), (AS). These authors contributed equally to this manuscript (Vedikaa Dhiman, Debankona Marik). Lead Contact (AS).

## Abstract

We sequenced the drought-response transcriptome of the keystone tree species *Prosopis cineraria* from the Indian Thar desert to understand the key factors in its drought tolerance mechanism. We identified a network of genes activated in *P. cineraria* involved in osmotic stress response, phytohormone, calcium, and phosphorelay signal transduction. Of these, up-regulation of 54 APETALA2/Ethylene-Responsive Factor (AP2/ERF) transcription factor genes, validated by real-time PCR, suggests their key role in the drought tolerance of *P. cineraria.* We conducted a genome-wide study of the AP2/ERF superfamily in *P. cineraria*, classifying its 232 proteins into 15 clades and analyzing their protein structures, gene structure, and promoter organization. The *P. cineraria* genome contains more copies of *AP2/ERF* genes than drought-sensitive plants. Further, we identified sequence polymorphisms in *AP2/ERF* genes between Arabian and Indian cultivars of *P. cineraria*. We modeled the DNA-protein complex structures of AP2/ERFs from drought-tolerant and sensitive species using AlphaFold to compare their DNA binding ability. Though the DNA binding domain (DBD) is relatively conserved across species, the unstructured region of these proteins possesses different charge distributions, which might contribute differently to their DNA search and binding. Using all-atom molecular dynamics simulations, we teased out a higher number of specific DBD-DNA hydrogen bonds in *P. cineraria,* leading to a stronger DNA-binding affinity compared to drought-sensitive *Arabidopsis thaliana*. These results directly support copy number expansion of AP2/ERF transcription factors and the evolution of their structures for more efficient DNA search and binding as drought adaptation mechanisms in *P. cineraria*.

## 1. INTRODUCTION

The Khejri tree (*Prosopis cineraria* (L.) Druce) is a perennial leguminous crop of the Thar desert in north-western Rajasthan, with its habitat stretching across Pakistan, Iran, and the Arabian Peninsula. Apart from its various uses as a vegetable, fodder, wood, and traditional medicine, it is also a keystone species crucial for sustaining the Thar desert ecosystem (Baibout et al. 2022). The leguminous nature of *P. cineraria* renders it highly tolerant to abiotic stresses viz., drought, heat, and nutrient deficiency. Due to its extensive root system, it can fix nitrogen through its nodulous roots and mine water from deep underground. The 691 Mb genome sequence of *P. cineraria* was recently available from an Arabian cultivar PC_DT_omini (GCF_029017545.1). The expansion of retrogenes in its genome gives critical insights into this plant’s extreme disease resistance and abiotic stress tolerance (Sudalaimuthuasari et al. 2022). However, the genome-wide organization, expression patterns, protein structure, and functions of its stress-tolerance genes remain elusive.

Plants have evolved stress-responsive transcription factors (TFs), master regulatory molecules induced in response to different biotic and abiotic stresses. These TFs interact with *cis*-acting elements in the promoters of numerous downstream stress-responsive genes. The APETALA2/Ethylene-Responsive Factor (AP2/ERF) superfamily consists of many TFs participating in all land plants’ growth, development, metabolism, and stress responses (Zhang et al. 2022). This superfamily consists of a highly conserved DNA-binding domain (DBD), a transcription regulation domain, an oligomerization site, and a nuclear localization signal (Liu et al. 2023). Following the convention of Nakano et al. (2006), the AP2/ERF superfamily is further classified into four families: AP2, ERF, Related to Abscisic Acid Insensitive 3/Viviparous 1 (RAV), and Soloist (Zhu et al. 2023). Among them, the *AP2* and *ERF* family genes are well known for regulating plant responses to abiotic stresses such as drought, salt, heat, and low temperature. The ERF family comprises two subfamilies: ERF and Dehydration-Responsive Element-Binding Protein (DREB). DREBs are induced in response to drought, salinity, heat, cold, and exogenous abscisic acid (ABA) but function through the ABA-independent signal transduction pathway. DREBs bind to the DRE/C**-**repeat element (CRT) having the core sequence motif A/GCCGAC (Liu et al. 1998; Nakano et al. 2006; Sakuma et al. 2006; Huang et al. 2020). While the *DREB1* genes are specific to cold, the DREB2s (of the A-2 subgroup) are specific to drought, salinity, and heat stress in Arabidopsis (Sakuma et al. 2006; Sadhukhan et al. 2014). On the other hand, ERF-subfamily proteins bind to the GCC box, which has the core sequence motif AGCCGCC (Fujimoto et al. 2000). *DREB* and *ERF* gene orthologs have been reported from drought-tolerant (DT) and drought-sensitive (DS) plants. *OsERF115* was recently identified as a novel positive regulator of heat and drought stresses and a negative regulator of ABA signaling in rice, poised as a candidate gene for developing heat and drought-resistant crops (Park et al. 2021). Furthermore, 224 AP2/ERF family members protect lettuce against cold, heat, drought, and salinity (Park et al. 2023). A recent genome-wide analysis identified 167 pearl millet *AP2/ERF* genes (Xu et al. 2024). Different abiotic stresses strongly induced the transcript levels of *PgERF* genes in different tissues. Recently, a study demonstrated the presence of many *DREB* genes in wheat, where overexpression of *TaDREB3* promoted tolerance to heat, dehydration, and salt stresses (Niu et al. 2020). In sugar beet, thirty *BvDREB* genes were identified at the genome-wide and transcriptome level, which regulated drought tolerance by elevating proline and ABA (Qiu et al. 2024). Overexpression of *DREB46* from *Populus trichocarpa* led to several drought-tolerance traits, including the scavenging capacity of Reactive oxygen species (ROS) and enhancing root growth (Geng et al. 2023). *AnDREB5* gene isolated from a desert plant *Ammopiptanthus nanus* conferred drought and cold tolerance in transgenic tobacco (Zhu et al. 2024). Despite these individual findings, the differences in DREB and ERF proteins between DT and DS plant species have yet to be explored in detail.

In this study, we conducted a transcriptome analysis of an Indian cultivar of *P. cineraria* treated with drought stress in the laboratory. An enrichment of TFs among genes highly induced by drought, particularly 54 genes of the AP2/ERF superfamily, prompted us to conduct a genome-wide analysis of this superfamily in this species. We explored the copy number variation (CNV) between the DBD of AP2/ERF proteins of DT and DS species to identify any possible correlation with drought tolerance. Structurally, DREB and ERF subfamily TFs have a single AP2 domain, differing at a few residue positions. For instance, ERF has alanine at the 14^th^ position, while the corresponding residue in DREB is valine (Sakuma et al. 2002). In binding affinity studies, DREB and ERF proteins exhibit strong interaction between the GCC box-binding domain of the proteins and the target DNA (Hassan et al. 2022). Structural studies have shown that the DBD, consisting of 59 amino acids, is tightly conserved in plants and helps bind the GCC box. While the N-terminal half of the DBD contributes to its high affinity binding with the GCC box, the divergent C-terminal half is important for modulating its specificity (Hao et al. 1998). Characterization of the DBD in terms of physico-chemical, geometrical, and structural features is crucial to understanding the differential binding strength of transcription factors as well as for accurate prediction of DNA-binding affinity (Dey et al. 2012; Pal et al. 2022; Harini et al. 2023). In this study, using molecular modeling and molecular dynamics (MD) simulations, we further dissected the structural differences between the DBD of AP2/ERF proteins of DT and DS species.

## 2. MATERIALS AND METHODS

### 2.1 Reference-guided transcriptome analysis of *Prosopis cineraria* under drought stress

Seeds of the Indian cultivar of *P*. *cineraria* were collected from the Indian Institute of Technology Jodhpur campus at the edge of the Indian Thar desert, removed from pods, and stored at room temperature. Surface sterilization of the seeds was done by rinsing with 70% ethanol and washing five times with deionized water. Sterilized seeds were germinated on moist cotton wool and grown on sterilized soil rite for a month. One-month-old seedlings were uprooted by carefully removing the soil rite without damaging the roots and washed in deionized water. The roots were dipped for six hours in either a 1/4^th^ Hoagland’s solution (control) or a 1/4^th^ Hoagland’s solution with 5% PEG-6000 (Himedia, Mumbai, India). The youngest compound leaf (the fifth from the cotyledon) was then picked and quickly frozen in liquid nitrogen. Total RNA was isolated by the CTAB-LiCl method. RNA quality and quantity were determined using the Agilent 4150 TapeStation system (Agilent Technologies, CA, USA) and Qubit 4 fluorometer (Thermo Fisher, Mumbai, India). The NebNext Ultra II RNA library prep kit (New England Biolabs, MA, USA) was used to synthesize RNA libraries. Paired end read (2×150 bp) sequencing of quality libraries was performed on the NovaSeq 6000 V1.5 platform (Illumina, CA, USA). The libraries’ raw sequence read quality, GC content, and complexities were determined using the FastQC software (Andrews, 2010). The raw transcriptome data were submitted to the NCBI Sequence Read Archive with a BioProject accession number PRJNA1023008. Low-quality sequences below a minimum size of 25 bp and a minimum quality score of 10 were removed by Bbduk software. The reads were aligned to the *P. cineraria* genome (GCF_029017545.1) by the Spliced Transcripts Alignment to a Reference (STAR) tool (Dobin et al. 2013). The *DEseq2*, *edgeR*, and *ggplot2* packages in R software (Chen et al. 2016; Love et al. 2014; Wickham, 2016) were used for differential gene expression analysis. Genes with a Benjamini-Hochberg false discovery rate *P* < 0.05 and -1 ≤ log_2_fold-change ≥ 1 were deemed differentially expressed genes (DEGs). The g:Profiler web server (Kolberg et al. 2023) was used for functional enrichment analysis of the up-and down-regulated genes.

### 2.2 *De novo* transcriptome assembly of the Indian cultivar of *Prosopis cineraria*

Genome-guided *de novo* transcriptome assembly was conducted using the Trinity pipeline (Grabherr et al. 2011). Compared to a genome-free *de-novo* assembly, a reference genome can be advantageous for paralogs or genes with shared sequences. Hence, coordinate-sorted binary alignment map (BAM) files were employed with the parameter *genome_guided_max_intron* set at 10,000 for the *de novo* assembly. The following methods assessed the assembly quality. First, assemblies were evaluated against a database of single-copy orthologous genes for plants, implemented in the Benchmarking Universal Single-Copy Orthologue (BUSCO) tool (Simão et al. 2015). Subsequently, the length of the contigs covering at least 50% of the assembly (N50 score) was measured for the assemblies.

### 2.3 Network analysis of *Prosopis cineraria* genes differentially expressed under drought

*P. cineraria* DEGs under drought stress were first converted into their nearest *Arabidopsis thaliana* orthologs using the following method. The protein sequences of *P. cineraria* were downloaded from NCBI (genome assembly GCF_029017545.1). *A. thaliana* protein sequences were downloaded from the Uniprot database and used for database creation using the “makeblastdb” module of the BLAST suit. The query sequences of *P. cineraria* proteins were searched against the *A. thaliana* protein database generated above. The homology search results were filtered, and the *A. thaliana* UniProt IDs were assigned to each of the *P. cineraria* protein IDs based on the topmost hit. The assigned *A. thaliana* IDs were mapped to AGI codes using the TAIR database (https://www.arabidopsis.org/). The Search Tool for the Retrieval of Interacting Genes (STRING) database and Cytoscape version 3.10.1 (Shannon et al. 2003) were used to construct a protein-protein interaction (PPI) network. To build the PPI network, the AGI codes of the *A. thaliana* orthologs of *P. cineraria.* The DEGs were fed into the software to construct two separate networks for up and down-regulated genes. Interactions with a score > 0.7 were deemed statistically significant. Functional annotation and enrichment analysis were conducted to understand the biological functions of the network genes using the STRING application within Cytoscape. The significance level was defined by a Benjamini–Hochberg false discovery rate (FDR) cut-off of 0.05. The important genes were identified by creating subnetworks with the MCODE application 2.0 available in Cytoscape. The following cut-offs were used: degree ≥ 2, node score ≥ 2, K-core ≥ 2, and max depth = 100. Further, hub genes were identified by observing and interpreting the MCODE and enrichment analysis results.

### 2.4 Phylogenetic classification of AP2/ERF proteins in *Prosopis cineraria*

The amino acid sequences of *AP2/ERF* genes differentially expressed (log_2_fold-change ≥ 1) in *P*. *cineraria* under PEG-induced drought stress were obtained from NCBI. A BLASTP search on de-duplicated 57 sequences against the whole proteome of *P. cineraria* identified the genome-wide paralogs of DREB and ERF proteins. A cut-off of *e*-value ≤ 0.05 and bit-score ≥ 150 were used for the BLAST search. After filtering unique BLAST hits, multiple sequence alignment (MSA) was conducted using Clustal Omega (Sievers et al. 2011). The maximum likelihood (ML) method with the Jones-Taylor-Thornton model and 1000 bootstrap replications were used to construct the phylogenetic tree. The phylogenetic tree was visualized, and the rooted tree was considered to further classify the proteins into different groups based on their evolutionary relationship by the Interactive Tree of Life (iTOL) tool (Letunic and Bork, 2021).

### 2.5 Real-time quantitative PCR analysis of *Prosopis cineraria AP2/ERF* genes under drought

A real-time quantitative PCR (qPCR) test confirmed the transcriptome result. One-month-old Indian cultivar *P. cineraria* seedlings were stressed with 5% PEG, and the fifth leaf from the cotyledon was harvested and snap-frozen in liquid nitrogen at different time points, 0 h, 1 h, 6 h, 12 h, and 24 h, for control as well as stressed samples. The tissues were crushed using a mortar and pestle. Total RNA was isolated using the CTAB-LiCl method, and cDNA synthesis was carried out using the PrimeScriptTM IV 1st strand cDNA Synthesis Mix (DSS Takara Bio India Pvt. Ltd., New Delhi, India). TB Green Premix Ex Taq II (Tli RNase H Plus) (Takara) was used for qPCR analysis in an Mx3000P qPCR machine (Agilent). The relative expression levels of five genes were estimated in control and stressed samples using the standard curve method, considering *PcActin* (gene ID: 129287134) as a housekeeping gene reference. Primers for qPCR were designed from the *de novo*-assembled Indian cultivar *P. cineraria* gene sequences using Primer3 web tool version 4.1.0 (https://primer3.ut.ee/).

### 2.6 Analysis of the conserved motifs and gene structures in *Prosopis cineraria* AP2/ERF proteins

The AP2/ERF proteins of *P. cineraria*, mined from NCBI, were subjected to motif discovery analysis using the MOTIF search tool (https://www.genome.jp/tools/motif/). The PEST sequences for protein degradation were identified using epestfind (https://emboss.bioinformatics.nl/cgi-bin/emboss/epestfind). TBtools v2.034 was used to illustrate the exon-intron organization within the coding sequence of the *AP2/ERF* genes (Chen et al. 2020). The coding sequences of 60 *P. cineraria AP2/ERF* genes, genome-wide paralogs, and the general feature format files were retrieved from NCBI and fed to TBtools for gene structure analysis.

### 2.7 Promoter and *cis*-acting element analysis of *AP2/ERF* superfamily genes in *Prosopis cineraria*

The Genome Data Viewer of NCBI was utilized to view the position of the highly expressed *AP2/ERF* genes from each group obtained from the phylogenetic classification of *P. cineraria*. The promoter sequences 2.0 kb of sequence upstream of the transcription start site of these *AP2/ERF* genes were retrieved from the NCBI. The promoter *cis*-acting elements of the highly expressed *AP2/ERF* genes were studied by the PlantCare database (Lescot 2002) and visualized by the TB tools software.

### 2.8 Polymorphism analysis in *AP2/ERF* genes between Indian and Arabian cultivars of *Prosopis cineraria*

To identify the nucleotide sequence variations between the Indian and Arabian cultivars of *P. cineraria*, we utilized the *de novo* assembled genome of the Indian cultivar and NCBI-retrieved nucleotide sequences of the Arabian cultivar. Burrows-Wheeler Alignment (bwa) tool, samtools, Picard, bcftools mpileup, haplotypecaller, and mutect2 of Genome Analysis Toolkit (GATK) were used for the DNA polymorphism analysis. The FASTA format sequences of the Indian cultivar were converted to FASTQ and aligned to the Arabian cultivar reference genome. Samtools was used to create a sequence alignment map (SAM), BAM, and sorted BAM files of the aligned FASTQ files of the *de novo* Indian cultivar. Further, the BAM files were checked for errors using the Picard tool. Finally, the genetic variants calling were performed through bcftools mpileup, haplotypecaller, and mutect2 of the GATK tool. A variant call format (VCF) file consisting of detailed information on the variants, including single nucleotide polymorphism (SNP), multiple nucleotide polymorphisms (MNP), insertions, deletions, and insertions and deletions (INDEL), was obtained after variant calling. The VCF file was analyzed in Integrative Genomics Viewer (Thorvaldsdóttir et al. 2013) to visualize the polymorphisms among 60 *AP2/ERF* genes induced under drought stress.

### 2.9 Copy-number variation analysis of the AP2/ERF superfamily between drought-tolerant and sensitive species

The CNVs in *AP2/ERF* superfamily genes of DT and DS species were identified by NCBI BLASTP search. For this purpose, 10 diploid DT species, cowpea (*Vigna unguiculata*), Khejri (*P. cineraria*), soybean (*Glycine max*), sorghum (*Sorghum bicolor*), poplar (*Populus trichocarpa*), foxtail millet (*Seteria italica*), cotton (*Gossypium raimondii*), cassava (*Manihot esculenta*), a desert legume (*Eremosparton songoricum*), and a desert moss (*Bryum argenteum*), and 10 diploid DS species, namely, rice (*Oryza sativa* subsp. *japonica)*, *A. thaliana*, pea (*Pisum sativum*), tomato (*Solanum lycopersicum*), potato (*Solanum tuberosum*), pigeonpea (*Cajanus cajan*), common bean (*Phaseolus vulgaris*), chickpea (*Cicer arietinum*), Medicago (*Medicago truncatula*), mungbean (*Vigna radiata*) were considered. The whole proteome of DT and DS species as databases and amino acid sequences of the AP2/ERF proteins as queries were used for an exhaustive BLASTP search. The query sequences were selected and extracted from NCBI based on the maximum number of amino acids present. The AP2/ERF sequences were filtered from the resultant BLAST hits, carefully deduplicated, and all partial sequences removed. Bar graphs were used to compare the CNVs between DT and DS species.

### 2.10 Modeling of *apo* and DNA-bound structures of the AP2/ERF and DREB proteins from drought-tolerant and sensitive species

The 3D structures of the *apo* form of individual proteins were modeled using AlphaFold2, downloaded from the GitHub repository and run locally (Jumper et al. 2021). We used AlphaFold2 pipeline and “full_dbs” to generate the MSA features for each protein sequence. A release date cut-off of February 1, 2024, was used for structure predictions using templates. Better prediction accuracy was ensured by increasing “num_recycles” value to 12. Next, the Alphafold2 structure of each protein was used to model their DNA-bound form, where the DNA-bound AP2/ERF structure from *A. thaliana* available in the Protein Data Bank (PDB id: 1GCC) was used as a template. Only the DBD of the proteins bound to the DNA from the complex structure was considered for the MD simulations.

### 2.11 Hydropathy and electrostatic profiles of AP2/ERF sequences

The hydropathy (hydrophilicity and hydrophobicity) of the AP2/ERF proteins along their amino acid sequence was evaluated using a moving-segment approach that continuously determines the average hydropathy within a window length as it advances through the sequence (Kyte and Doolittle 1982). Here, we took a window of five residues to calculate a running mean of hydropathy values for the reference sequences of *P.cineraria* and *A. thaliana* of groups VII and VIIIa for comparative analysis. Following the same Kyte-Woolridge scale, we generated a running mean of electrostatic values with a window of five amino acid residues to compare the electrostatic properties of *P. cineraria* and *A. thaliana* of groups VII and VIIIa. These comparisons were conducted region-wise for the N-terminus, DBD, and C-terminus.

### 2.12 All-atom molecular dynamics simulations

All-atom MD simulations for the protein-DNA model complexes were performed using GROMACS 2023 package with the AMBER03 force field (Ponder & Case, 2003). For better sampling, the production run of each of the four protein-DNA systems was repeated three times in explicit-solvent starting from different atomic velocity distributions, each lasting 200 ns. Each protein’s N- and C-termini regions were capped with NH_2_ and COOH groups. For each titratable residue, its default protonation state was considered. The complex structure was placed at the center of a cuboid box, the size of which was considered so that each protein or DNA atom was at least 1.2 nm away from the nearest edge of the box. Next, the box was solvated using water of SPC216 model type. The net charge of the system was neutralized by adding Na^+^ ions. The steepest descent algorithm avoided any unfavorable interaction by subjecting the solvated system to 1000 steps of energy minimization. Next, the system was gradually heated to 300K over a period of 500 ps under NVT conditions, following equilibration for 1ns using the NPT ensemble. Finally, at NPT conditions, a production run of 200 ns with a time step of 2 fs was performed. Electrostatic interactions were treated using the particle mesh Ewald (PME) method. Periodic boundary conditions were applied in x, y, and z directions. The simulation temperature of 300 K was set using a modified Berendsen thermostat. The pressure was maintained at 1 bar using Parrinello-Rahman pressure coupling. A cutoff distance of 1.2 nm was implemented for nonbonded interactions. All the proteins remained bound to the DNA, similar to the starting conformation throughout the simulations. Trajectory analysis was done using GROMACS tools and in-house codes.

### 2.13 Hydrogen bond analysis

For the hydrogen bonding analysis, conformations from the trajectories were saved at an interval of 500 picoseconds, resulting in 2000 structures from each simulation. Hydrogen bonds were calculated from each structure using the HBPLUS program (McDonald and Thornton 1994). Thus, hydrogen bond statistics for each system were obtained from 6000 structures from three independent simulations. Those bonds were consistently present in over 50% of cases and in at least two replicates were finally considered.

### 2.14 Binding energy calculation

Binding energy of the whole protein-DNA complex as well as contribution per residue was calculated using free energy decomposition method (Bashford & Case, 2000) on 2000 equally-spaced structures extracted from each trajectory as described above. MMPBSA.py (Miller et al. 2012) script in AMBERTools24 was used to calculate binding energy where Molecular Mechanics/ Generalized Born Surface Area (MM/GBSA) method was applied. Ions and waters were stripped from the trajectories, and the Generalized Born Solvation Model represented the solvent effect (Onufriev et al. 2004). A 100 mM salt concentration was considered. The ICOSA method estimated the nonpolar contribution to solvation-free energy from the solvent-accessible surface area (SASA). The average energy calculated from the three simulations was reported.

## 3. RESULTS

### 3.1 Stress-tolerance genes are highly induced in *Prosopis cineraria* under drought

Transcriptome analysis of the fifth leaf of *P. cineraria* seedlings treated with 5% PEG-6000 resulted in up-regulation of 2434 genes and down-regulation of 1377 genes with an absolute log_2_ fold-change ≥ 1 and *FDR* < 0.05 (Fig. 1A and Table S1). The functional enrichment analysis of the 2434 up-regulated genes in *P. cineraria* under drought stress revealed enriched biological processes like cell cycle regulation, olefinic compound metabolic process, microtubule-based movement, response to chemicals, and response to bacteria (Table 1). Several gene ontology (GO) molecular functions, including DNA-binding TF activity, protein binding, kinase regulator activity, galactinol-sucrose galactosyltransferase activity, and microtubule motor activity, were enriched among the induced genes. The enriched Kyoto Encyclopedia of Genes and Genomes (KEGG) pathways were plant-hormone signal transduction and MAPK signaling pathways. The enrichment analysis unraveled *P. cineraria*’s stress signaling pathways associated with drought tolerance.

**Fig. 1.**
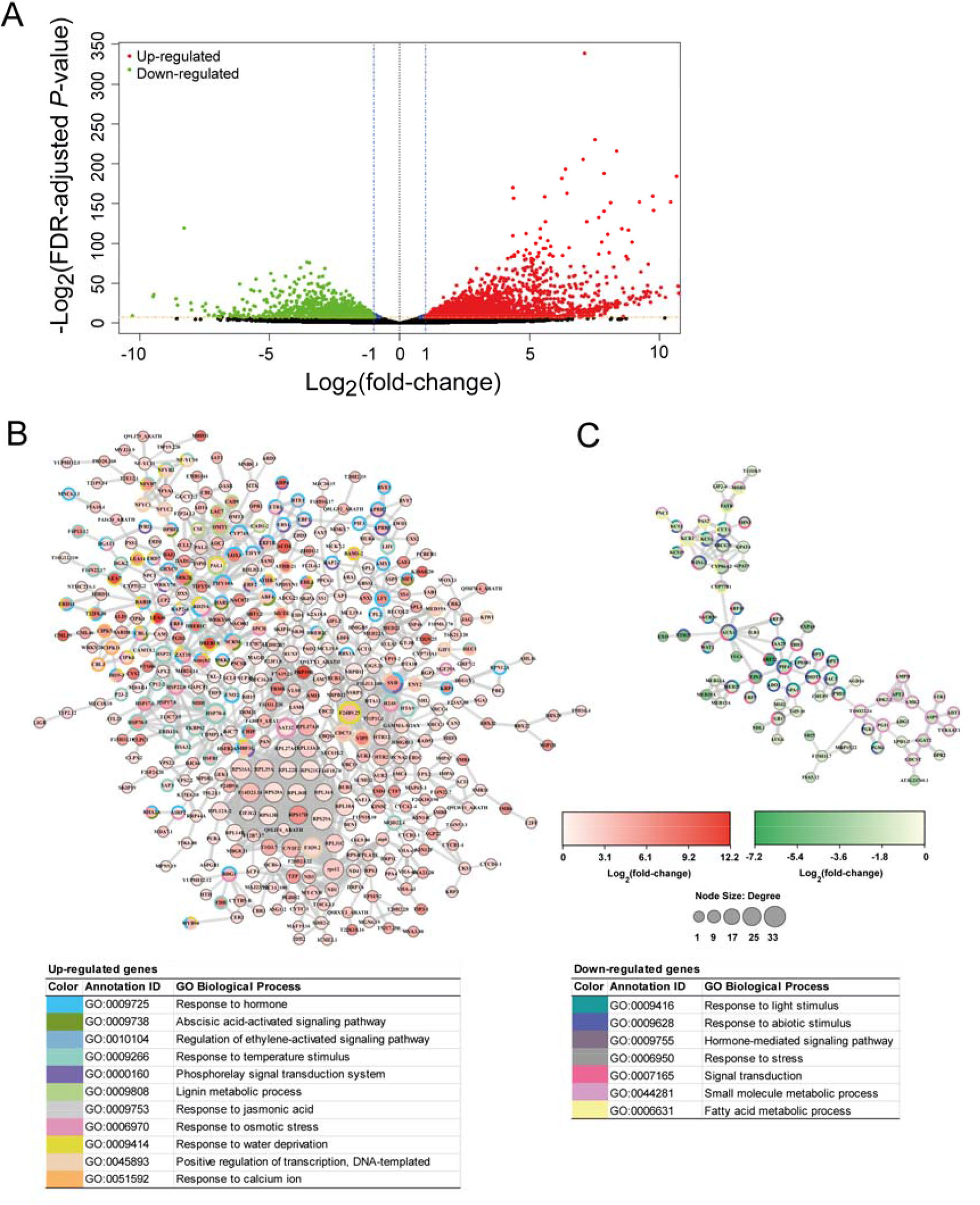
*Prosopis cineraria* genes differentially expressed under drought. Roots of one-month-old *P. cineraria* seedlings were immersed for 6 hours in ¼ Hoagland’s solution (control) and ¼ Hoagland’s solution with 5% PEG-6000 (drought stress). Transcriptome of the fifth compound leaf was analyzed from three control and three drought-stressed plants. **(A)** Volcano plot showing differentially expressed genes in *P. cineraria*. Red dots indicate 2434 up-regulated genes, and green dots 1377 down-regulated genes with false discovery rate (FDR)-adjusted *P*-value < 0.05 and three biological replicates. Protein-protein interaction (PPI) network of *Prosopis cineraria* **(B)** up-regulated genes with log_2_ fold-change > 1 and FDR < 0.05, **(C)** down-regulated genes with log_2_ fold-change < -1 and FDR < 0.05 under drought stress. The protein IDs were converted to their corresponding *Arabidopsis thaliana* orthologs (see Methods and Table S1). The pink node color indicates up-regulated genes, and the green color indicates down-regulated genes. The colored circles around the nodes represent different enriched pathways to which these nodes are linked. A key to the node size representing the degree of the nodes and a color key of the nodes representing the log_2_(fold-change) value of genes are shown at the bottom. The color chart represents enriched Gene Ontology biological process categories and annotation identities.

**Table 1.** Enriched functional categories of differentially expressed genes in *Prosopis cineraria* under drought stress.

| Enriched Terms | P value | Enriched Terms | P value |
| --- | --- | --- | --- |
| **Up-regulated genes** |  | **Down-regulated genes** |  |
| **GO molecular function** |  | **GO molecular function** |  |
| DNA-binding transcription factor activity | 1.35E-21 | protein binding | 3.70E-90 |
| sequence-specific DNA binding | 3.02E-17 | catalytic activity | 1.84E-76 |
| protein binding | 2.76E-07 | DNA binding | 3.79E-06 |
| microtubule motor activity | 8.50E-03 | metallopeptidase activity | 5.16E-05 |
| kinase regulator activity | 2.44E-02 | transporter activity | 6.60E-05 |
| galactinol-sucrose galactosyltransferase activity | 4.24E-02 | acetyltransferase activity | 1.08E-03 |
| kinase binding | 4.43E-02 | rRNA binding | 3.22E-03 |
| **GO Biological Process** |  | tRNA binding | 1.05E-02 |
| response to chemical | 5.84E-39 | exonuclease activity | 1.43E-02 |
| cell cycle process | 1.02E-09 | protein-macromolecule adaptor activity | 1.58E-02 |
| olefinic compound metabolic process | 7.49E-03 | oxidoreductase activity | 3.75E-02 |
| microtubule-based movement | 9.59E-03 | peptide receptor activity | 4.99E-02 |
| response to bacterium | 4.54E-02 | **GO Biological Process** |  |
| **GO Cellular Component** |  | cellular process | 4.94E-129 |
| cellular anatomical entity | 1.80E-09 | response to bacterium | 6.44E-09 |
| nucleus | 5.55E-09 | rhythmic process | 8.57E-08 |
| **KEGG Pathways** |  | protein maturation | 9.83E-06 |
| Plant hormone signal transduction | 2.21E-04 | mitochondrial gene expression | 1.34E-05 |
| MAPK signaling pathway - plant | 8.60E-04 | homeostatic process | 3.77E-04 |
|  |  | Group II intron splicing | 1.16E-03 |
|  |  | ta-siRNA processing | 3.43E-02 |
|  |  | entrainment of circadian clock | 4.51E-02 |
|  |  | sesquiterpenoid metabolic process | 4.52E-02 |
|  |  | **GO Cellular Component** |  |
|  |  | intracellular anatomical structure | 1.29E-218 |
|  |  | Cul4-RING E3 ubiquitin ligase complex | 7.53E-04 |
|  |  | preribosome | 9.40E-04 |
|  |  | **KEGG Pathways** |  |
|  |  | Metabolic pathways | 2.66E-03 |
|  |  | Nicotinate and nicotinamide metabolism | 6.18E-03 |
|  |  | DNA replication | 7.57E-03 |
|  |  | Biosynthesis of cofactors | 3.95E-02 |
| Enriched Gene Ontology and KEGG Pathways for 2434 up-regulated genes and 1377 down-regulated genes in *Prosopis cineraria* under drought stress identified using the g:Profiler software. The frequency of occurrence of a particular pathway in the set of 2434 genes (input number) versus the frequency in the *P. cineraria* genome (background number) determined the enrichment. Enrichment categories with P-value < 0.05 are presented (Benjamini-Hochberg FDR). |  |  |  |

A PPI network analysis of the identified DEGs was performed to understand the molecular interactions in *P. cineraria* under drought stress (Fig. 1B, 1C). We separately constructed the up-regulated and down-regulated PPI networks. A bird’s eye view of the PPI networks indicated that the number of DEGs was higher in the up-regulated network and fewer in the down-regulated network. In the PPI network, the degree of the nodes is denoted by the relative size, and a color scale shows the expression pattern. We performed an enrichment analysis of GO biological processes to unravel the important hub gene functions in the interaction network. A colored ring around each node represents enriched biological pathways. Our analysis showed significant enrichment of the specific processes such as “response to hormone”, “ABA-activated signaling pathway”, “regulation of ethylene-activated signaling pathway”, “response to temperature stimulus”, “phosphorelay signal transduction system”, “lignin metabolic process”, “response to jasmonic acid”, “response to osmotic stress”, “response to water deprivation”, “positive regulation of transcription”, “response to calcium ion” for the up-regulated genes network (Fig. 1B, Table S2). The MCODE analysis revealed 138 central up-regulated network genes, which were further narrowed down to 12 up-regulated hub genes based on their involvement in more than two functionally enriched pathways or biological processes and having a threshold value for degree > 6 and log_2_(fold-change) > 4 (Table S3). Similarly, for the down-regulated network, the significantly enriched processes were “response to light stimulus,” “response to abiotic stimulus,” “hormone-mediated signaling pathway,” “response to stress,” “signal transduction,” “small molecule metabolic process,” “fatty acid metabolic process” (Fig. 1C, Table S2). Likewise, MCODE analysis identified 35 down-regulated genes narrowed down to 11 hub genes based on their involvement in more than two functionally enriched pathways or biological processes and having a threshold value for degree > 5 and log_2_(fold-change) < -1.5 (Table S3). In a nutshell, most of the developmental and defense-related genes and a subset of stress-related genes were found to be down-regulated in this study, and the induced genes chiefly belonged to the family of stress response-related TFs and signaling proteins that potentially provide tolerance to *P. cineraria* against drought.

### 3.2 Classification of *Prosopis cineraria AP2/ERF* genes

The transcriptome data analysis indicated up-regulation of 54 and down-regulation of six *AP2/ERF* superfamily genes in *P. cineraria* under drought. We conducted a genome-wide study on these proteins to gain insights into the evolution of AP2/ERF superfamily proteins of the DT tree *P*. *cineraria*. Firstly, a BLASTP search was performed on the 60 AP2/ERF proteins differentially expressed (54 up-regulated and 6 down-regulated) under drought against the entire protein set of *P*. *cineraria.* This led to 232 unique genome-wide paralogs of AP2/ERF superfamily proteins. Next, a combined phylogenetic analysis of the AP2/ERF superfamily proteins of *P*. *cineraria* and *A. thaliana* showed their clustering into 12 major groups I to X, AP2, and Soloist (Fig. 2, S1). Moreover, groups II, VIII, and X were further classified into two sub-groups according to their evolutionary relationship with *A. thaliana* proteins of corresponding sub-groups. The proteins of groups I to Xb were classified into subgroups A1 A6 and B1 B6 for DREBs and ERFs, respectively, according to Sakuma et al. 2002. Groups I, IIa, IIb, III, IV, VI, VII, VIIIa, VIIIb, X, Xb, AP2, and Soloist comprised 8, 4, 10, 34, 9, 20, 8, 14, 10, 5, 55, 8, two, 40, and 5 *P*. *cineraria* protein members, respectively. Of these, groups III, IX, and AP2 comprised the maximum number of proteins. *A. thaliana* RAV proteins did not cluster with any protein of *P. cineraria*, confirming no RAV members in this desert tree.

**Fig. 2.**
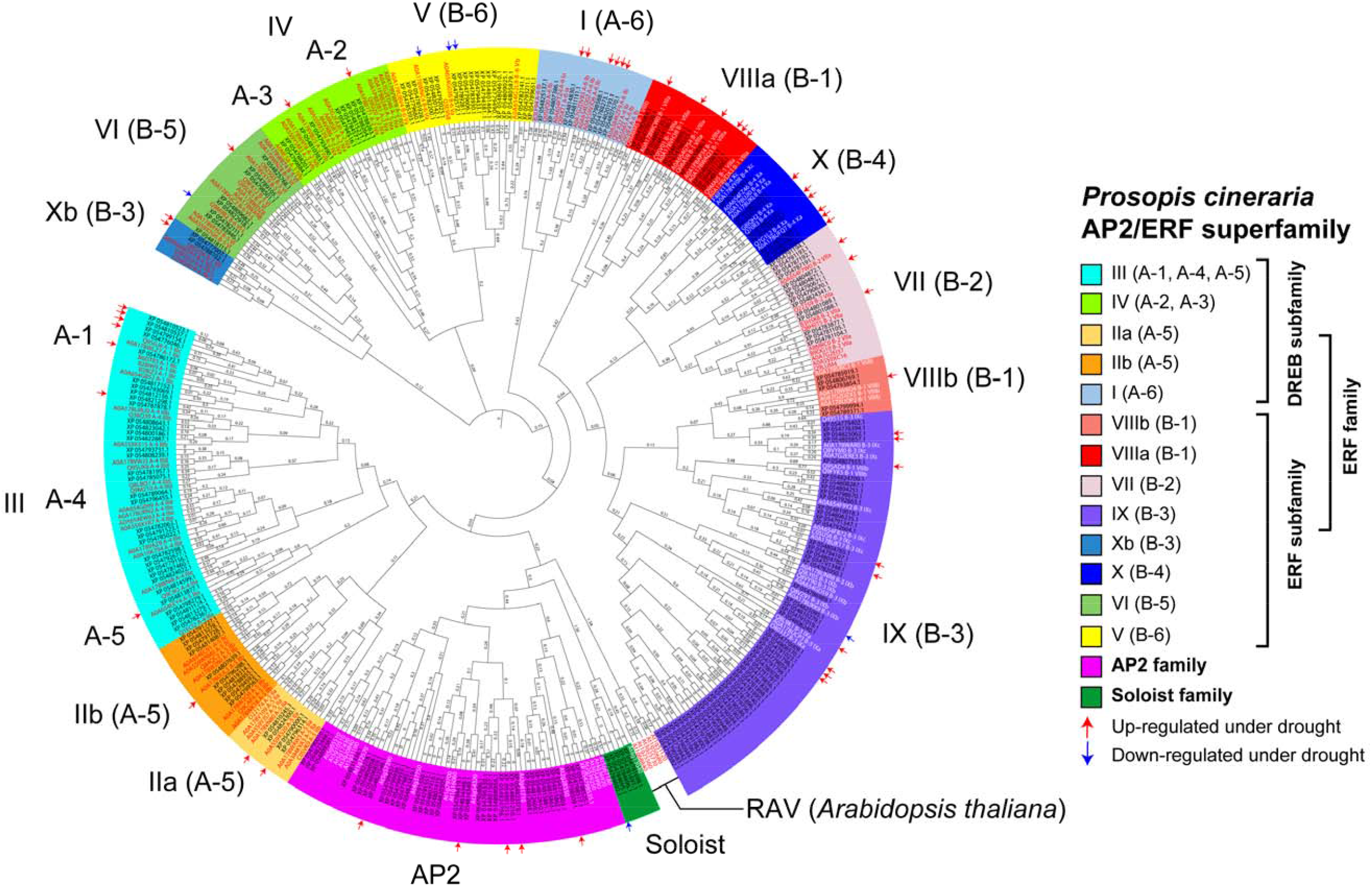
Classification of *Prosopis cineraria AP2/ERF* superfamily genes. The amino acid sequences of 60 differentially expressed genes of the AP2/ERF superfamily of *P. cineraria* under drought stress (see Table S1) were mined from the NCBI database. An NCBI BLASTP search with an e-value of ≤ 0.05 and a bit-score of ≥ 150 was performed exhaustively on these 60 proteins. Multiple sequence alignment (MSA) was performed using Clustal Omega on the 232 unique hits obtained from the NCBI BLASTP search, together with 154 *A. thaliana* AP2/ERF superfamily proteins. Phylogenetic tree was constructed using the maximum likelihood method with the Jones-Taylor-Thornton model with 1000 bootstrap replicates in the MEGA11 application. The groups I-X were divided based on the classification of Nakano et al. (2006), whereas groups A1 A6 and B1 6 followed the classification of Sakuma et al. (2002). Red and blue arrows indicate *P. cineraria genes* up-regulated and down-regulated under drought, respectively. *A. thaliana* proteins in each group are shown in red or white text.

### 3.3 *Prosopis cineraria AP2/ERF* genes display different expression patterns under drought stress

Total 54 *P. cineraria* genes from groups I, IIa, IIb, III, IV, VI, VII, VIIIa, VIIIb, IX, X, Xb, and AP2 of the phylogenetic tree (Fig. 2) were overexpressed (log_2_FC > 1) under drought stress in our transcriptome analysis (Table S1). In contrast, three genes from group V and one each from IX, VI, and Soloist were down-regulated with log_2_FC < -1 (Table S1). We validated the up-regulation of five randomly chosen genes, *PcDREB1A-like* (LOC129312011) from group III (A–1), *PcDREB1E-like* (LOC129311911) from group IV (A-2), *PcDREB2F* (LOC129320117) from group III (A–4), *PcERF109* (LOC129293970) from group Xb (B-3), and *PcAIL5* (LOC129320898) from group AP2, by qPCR using primers designed against the corresponding *de novo*-assembled sequences from the Indian cultivar (Table S4). The relative expression levels of the selected genes under drought compared to non-stressed control are shown in Fig 3A. The expression levels of *PcDREB2F* and *PcERF109* increased gradually in a time-dependent manner up to 24 h under drought. In contrast, the expression levels of *PcAIL5* and *PcDREB1A-like* sharply increased up to 12 h but dropped after long-term exposure of 24 h.

**Fig. 3.**
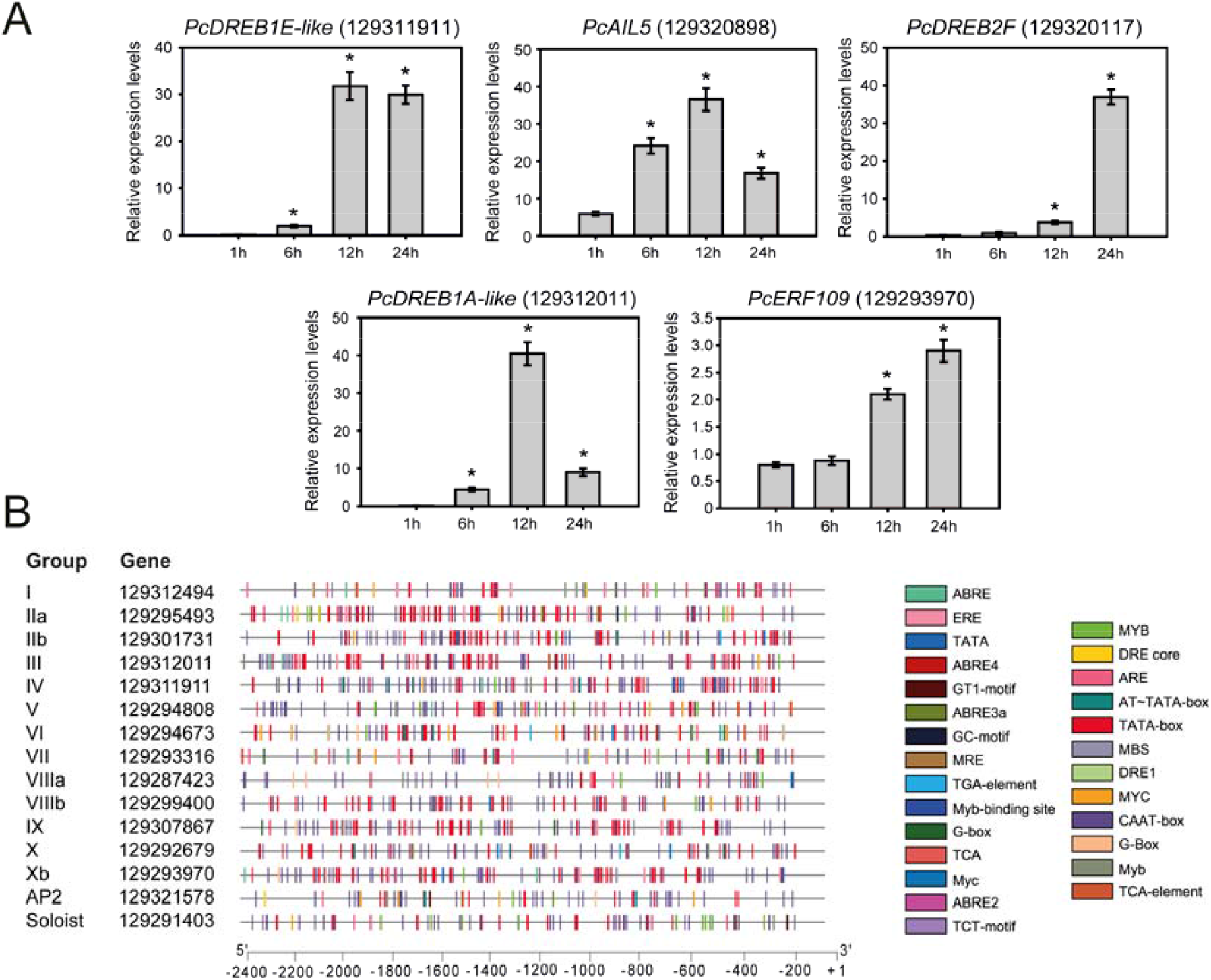
Expression and promoter *cis*-element analysis of *Prosopis cineraria AP2/ERF* genes. **(A)** Time-dependent expression analysis of *P. cineraria AP2/ERF* genes under drought stress is shown. *P. cineraria* seedlings, grown for one month in soil, were uprooted carefully, washed with distilled water, and submerged in ¼ strength Hoagland’s media for control samples and ¼ Hoagland’s media supplemented with 5% PEG 6000 for stressed samples. The fifth leaf from the cotyledon for both control and stressed samples was harvested at 0 h, 1 h, 6 h, 12 h, and 24 h and snap-frozen in liquid nitrogen. Total RNA was isolated and converted to cDNA for real-time PCR. The expression levels of five upregulated genes, *PcDREB1E-like* (gene ID: 129311911), *PcAIL5* (129320898), *PcDREB2F* 129320117), *PcDREB1A-like* (129312011), and *PcERF109* (129293970) in the transcriptome analysis (see Table S1) were validated by quantitative real-time PCR, by the standard curve method. *PcActin* (129287134) was the housekeeping gene to normalize sample expression levels. The relative fold changes in expression levels under PEG versus control at different time points are shown in panel A. The bars indicate an average of three biological replicates with standard errors. Asterisks indicate significant differences between relative expression levels at different time points (*P* < 0.05; Student’s *t*-test). **(B)** Promoter *cis-*elements of *P. cineraria* 15 *AP2/ERF* genes. The promoter sequence up to 2.4 kb of a representative member from each group of the phylogenetic tree (see Fig. 2) having the highest fold change of expression (Table S1) is shown. The *cis*-elements were identified by the PlantCare database and visualized by the TBtools. The scale at the bottom shows the promoter positions in base pairs. Colored boxes on the right indicate different *cis*-elements in the promoters.

### 3.4 *Prosopis cineraria AP2/ERF* promoters contain *cis*-elements for abiotic stress response

We studied the promoter architectures of *AP2/ERF* genes of *P. cineraria* to understand their roles in gene expression regulation under drought. We selected 15 genes with the highest log_2_(fold-change) values from all the phylogenetically classified groups. From groups I, IIa, IIb, III, IV, V, VI, VII, VIIIa, VIIIb, IX, X, Xb, AP2 and Soloist, the selected genes had log_2_(fold-change) values of 9.6, 3.2, 6.7, 10.7, 7.3, -2.4, 3.5, 3.2, 5.4, 5.0, 7.0, 8.7, 8.9, 1.9 and -1.2, respectively. Interestingly, our analysis of promoter *cis*-acting elements showed the presence of *cis*-elements responsible for abiotic stress tolerance (Fig. 3B). These included anaerobic response element (ARE), ethylene response element (ERE), G-box, MYB-binding site, MYC-binding site, TCA elements, TGA elements, abscisic acid-responsive element (ABRE), ethylene-responsive element (ERE), and the GT-1 motif. Drought stress response-related MYB and MYC-binding sites were present in all the groups. ABRE was found in 12 out of 15 groups, except for IIb, VIIIb, and Soloist. DRE was present only in groups I, IIa, VII, and AP2. ARE was present in groups I, III, IV, V, VI, VII, X, and AP2. ERE was present in groups I, IIa, III, IV, VII, VIIIa, IX, and Xb. The salt stress response-related GT-1 motif was found in groups IIa, IIb, AP2, V, and Soloist. The salicylic acid-responsive TCA element was present in groups I, III, IV, V, VI, VII, X, and AP2. The auxin-responsive TGA element was found in groups IIa, VI, and X.

### 3.5 Gene structure and conserved motifs of *Prosopis cineraria* AP2/ERF proteins

To gain insights into the structures of *P. cineraria AP2*/*ERF* genes, a gene map depicting exons and introns in the coding sequences and the 5’-untranslated regions (UTRs) was constructed. The genes were interrupted by introns varying in position and count for each group. In groups IIb, III, IV, V, VI, VII, VIIIa, IX, X, Xb, AP2, and Soloist, a varying number of introns were observed, i.e., 2, 8, 5, 11, 4, 17, 1, 7, 9, 1, 300 and 30, respectively. Noteworthy, groups I and IIa have zero introns, and these groups have highly conserved coding sequence regions. The *AP2* family genes contain the highest number of introns within the genes. Fig. 4 depicts the gene structures and conserved protein motifs. The amino acid sequences of all *P. cineraria* AP2/ERF proteins contain the conserved AP2 domain. A total of 31 *P. cineraria* AP2 family TFs contain two AP2 domains. The AP2 domain was present at the N-terminal end of the proteins of groups IIb, III, IV, V, VIIIa, and VIIIb, while in other groups, it was observed at the center (Fig. 4, S1). PEST sequences for protein degradation, rich in proline, glutamic acid, serine, and threonine, were detected in all groups except group VII. Another conserved domain, the dehydrin motif, was found only in four members of group IX (Fig. 4).

**Fig. 4.**
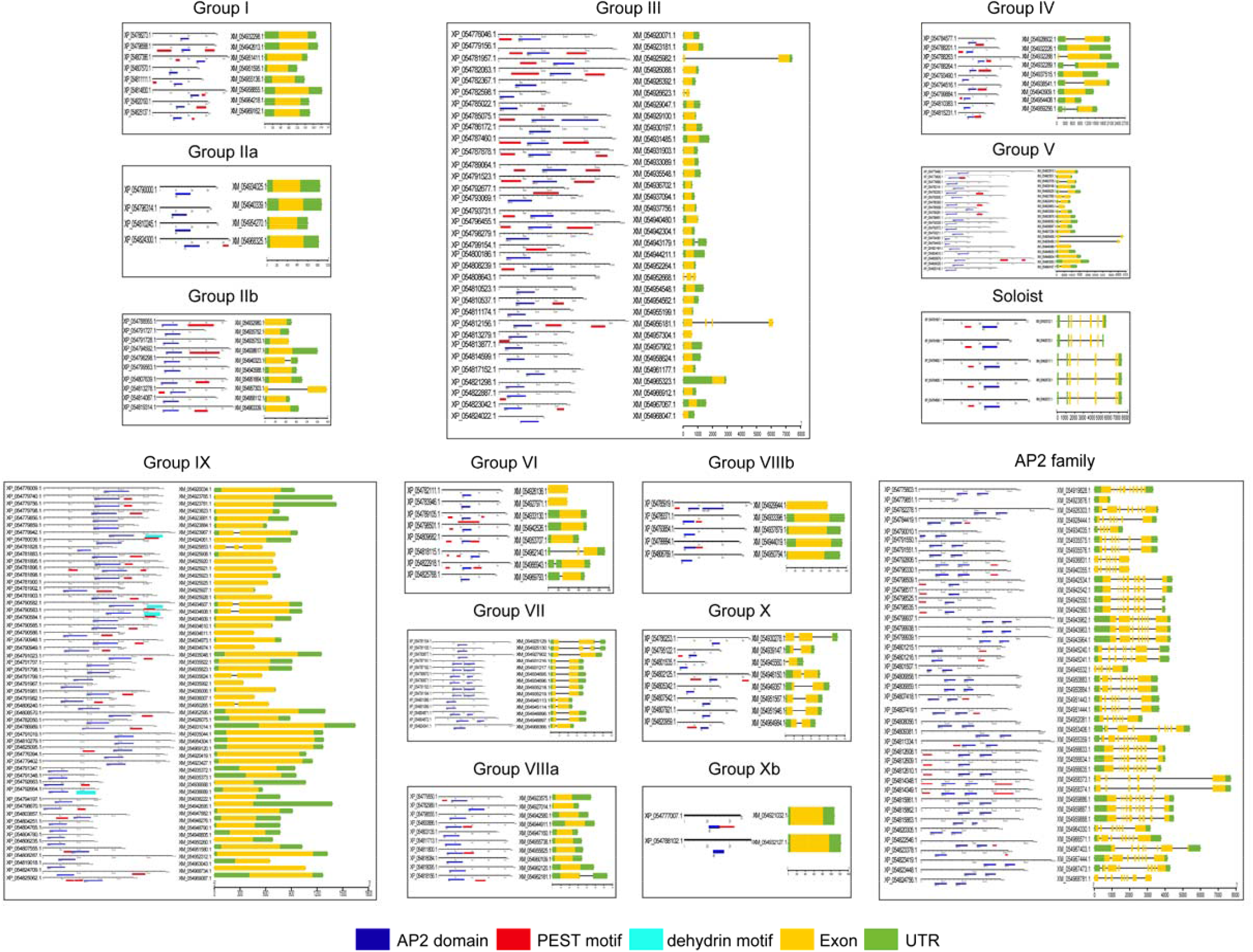
Conserved motifs and gene structures of *Prosopis cineraria* AP2/ERF superfamily proteins. The boxes show an arrangement of protein motifs and the gene structures according to the phylogenetically classified I– Xb, AP2, and Soloist of *P. cineraria* (see Fig. 2). *P. cineraria* AP2/ERF mRNA and amino acid sequences were mined from NCBI. A motif search tool identified conserved AP2 and dehydrin motifs (https://www.genome.jp/tools/motif/). PEST motifs were identified using epestfind (https://emboss.bioinformatics.nl/cgi-bin/emboss/epestfind). Blue, red, and cyan boxes represent the AP2 domain, PEST, and dehydrin motifs. The intron/exon arrangement-based gene structure of the AP2/ERF superfamily, which was analyzed in TBtools software, is next to the conserved motifs. The yellow and green boxes show exons and untranslated regions (UTR) in each mRNA sequence.

### 3.6 *AP2/ERF* gene polymorphisms between Indian and Arabian cultivars of *Prosopis cineraria*

*P. cineraria* has adapted to the deserts of Arabia and the Indian Thar desert, which have different rainfall patterns and ecosystems. To gain insights into the nucleotide sequence level variation between the Indian and Arabian cultivars, viz., SNP, MNP, insertions, deletions, and INDELS, we prepared a *de novo* assembly of the Indian cultivar from the RNA sequencing data. Two assemblies were generated: one containing the control leaf transcriptome (assembly-1) and the other incorporating both control and drought-stressed leaf transcriptomes (assembly-2). Assembly-1 consisted of 121,770 contigs, while assembly-2 comprised 156,767 contigs. The BUSCO assessment revealed that both assemblies exhibited high gene representation, with 97.6% for assembly-1 and 100% for assembly-2. Similarly, N50 scores of 2,227 bp for assembly-1 and 2,415 bp for assembly-2 were obtained. These assessments confirmed the high quality of the generated assemblies. We focused on polymorphism in the 60 *AP2/ERF* genes of *P. cineraria*, differentially expressed in our transcriptome analysis conducted with the Indian cultivar. We found that *mutect2* of the GATK tool outperformed bcftools, mpileup, and haplotypecaller in identifying the variations. Our analysis showed that out of 60 *AP2/ERF* genes, 49 genes contained SNPs, MNPs, and INDELs. Moreover, among 49 genes, the *ERF060-like* gene (LOC129309848) harbored the highest number (total 46) of SNPs, MNPs, and INDELs. Further, 19 SNPs were detected in the *ethylene-responsive transcription factor SHINE 2-like* gene (LOC129298195). Table 2 summarizes all identified variations in *AP2/ERF* genes between the Indian and Arabian cultivars of *P. cineraria*.

**Table 2.**
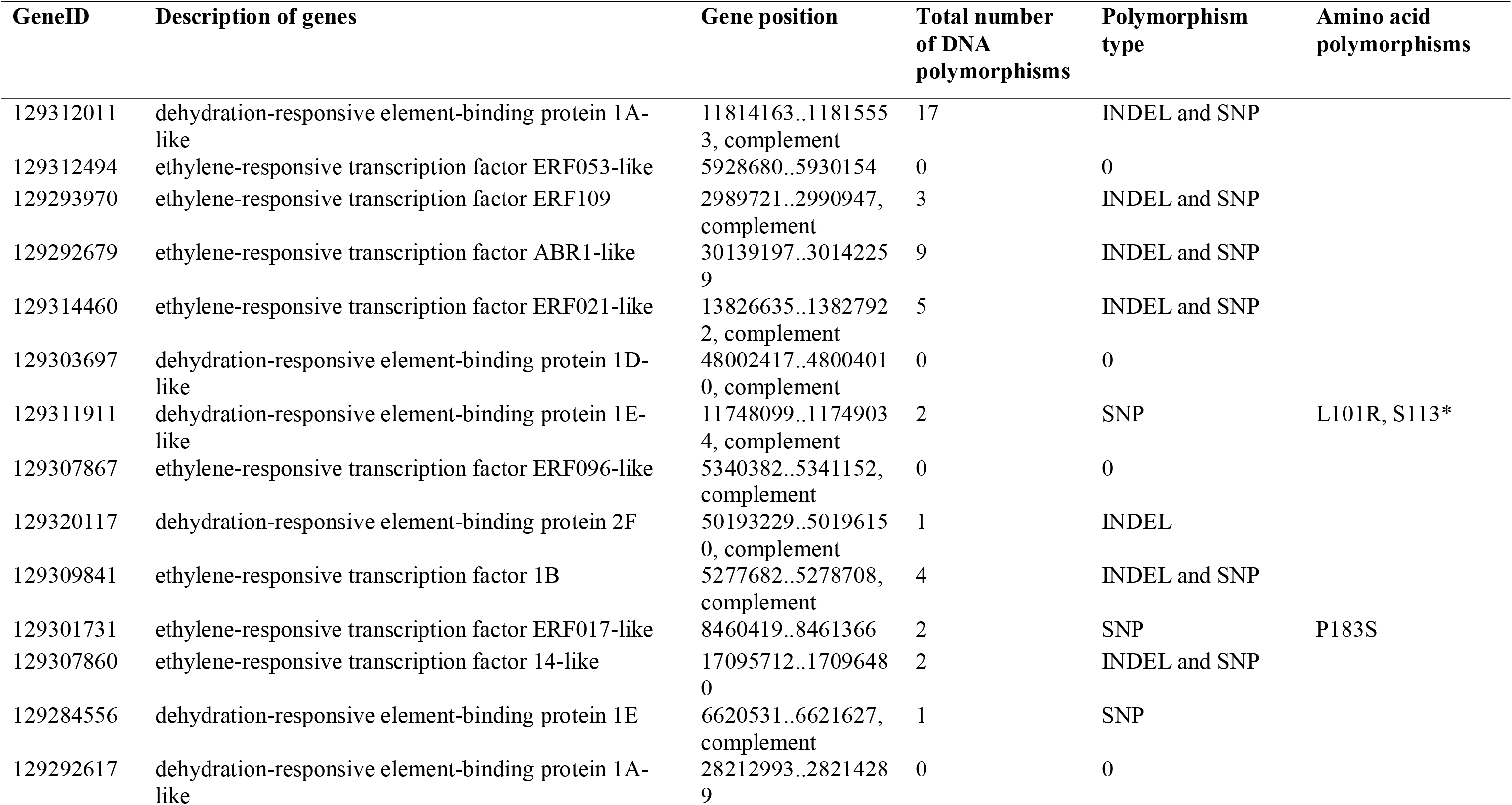
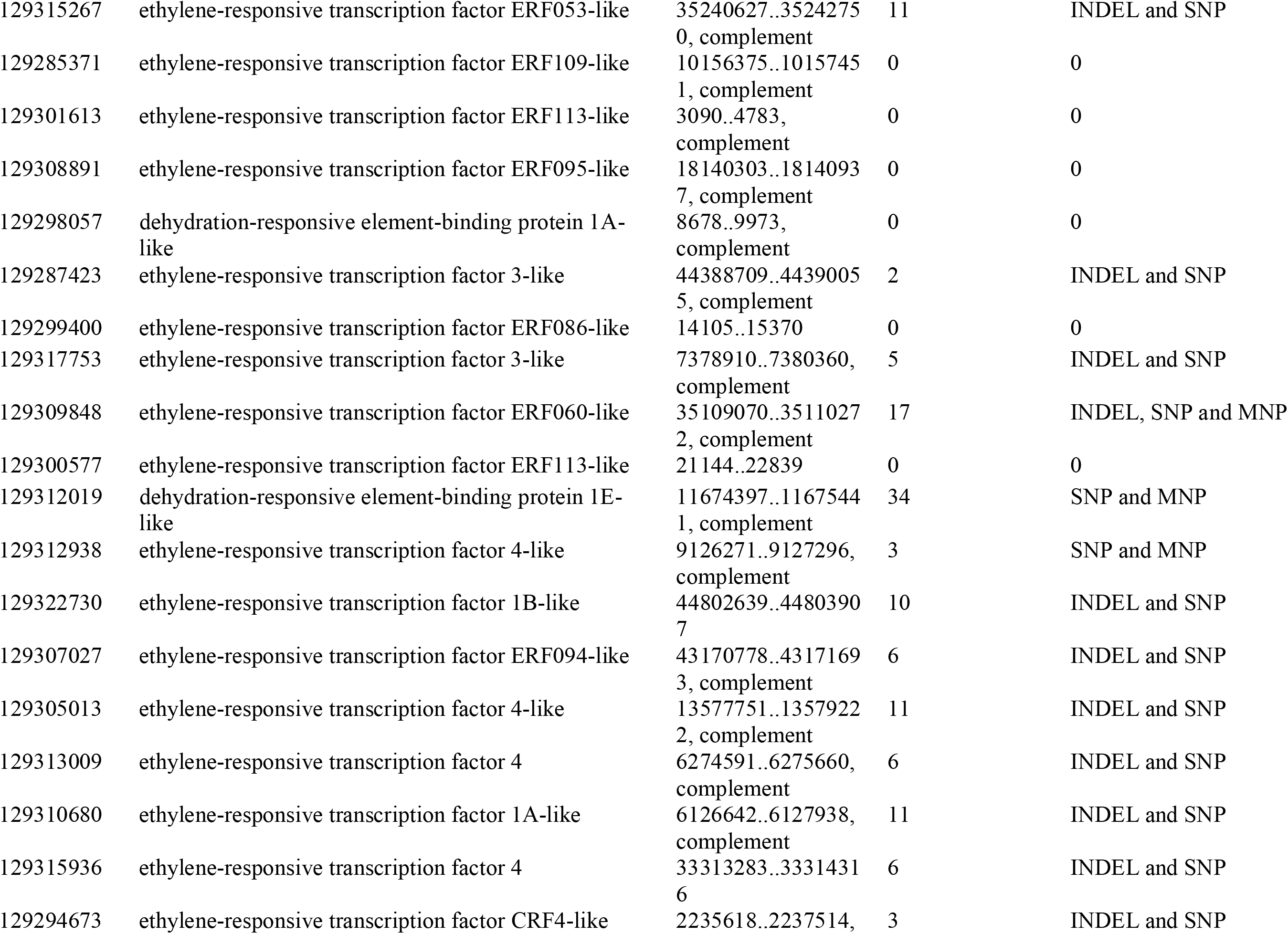
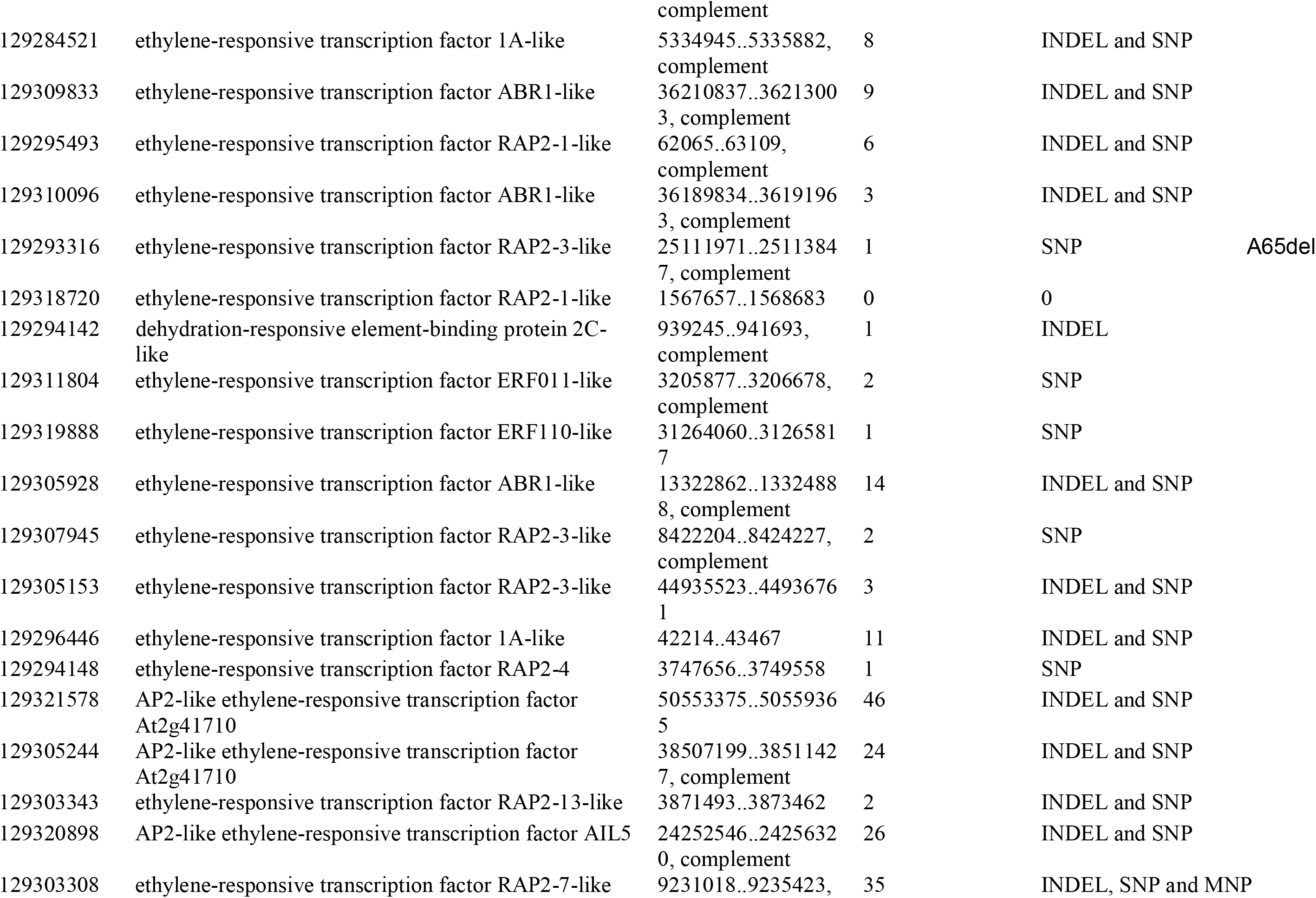
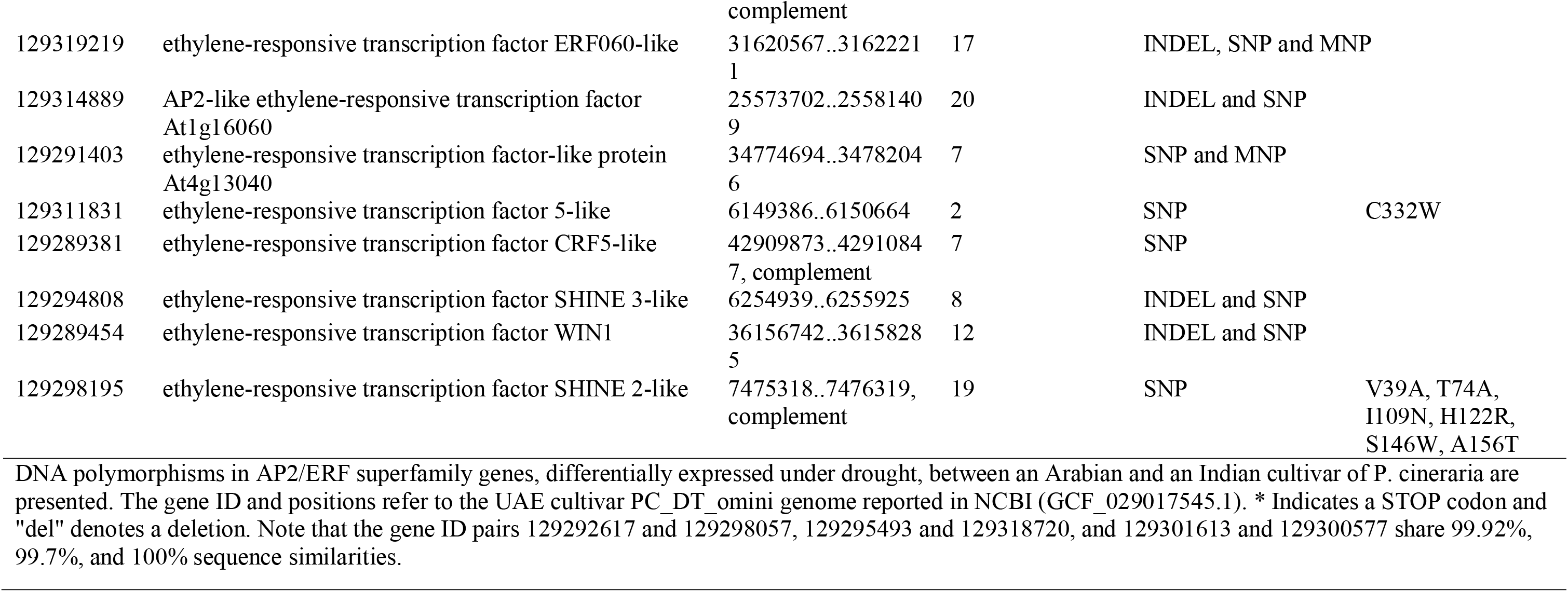
Polymorphism analysis in *AP2/ERF* genes between Indian and Arabian cultivars of *Prosopis cineraria*.

### 3.7 Drought-tolerant legumes contain more copies of *AP2*/*ERF* genes than drought-sensitive plants

We explored the CNVs of *AP2/ERF* genes between 10 DS and 10 DT species of equal ploidy (diploid) to understand their functional impact on drought tolerance (Fig. 5). We noticed a higher copy number of *AP2/ERF* genes in DT species than in DS species. This suggests an evolutionary strategy of expansion of *AP2/ERF* genes for drought adaptation (Agarwal et al. 2016). The DT legumes like cowpea, *P. cineraria*, and *Eremosparton songoricum* contain 309 (Agarwal et al. 2016), 232 (mined from NCBI for this study), and 153 copies (Zhao et al. 2022) of *AP2/ERF* genes, respectively. Non-legume DT trees like poplar contain 200 copies (Zhuang et al. 2008), and the diploid cotton *Gossypium raimondii* and *Gossypium arboreum*, 120 and 118 gene copies, respectively (Zafar et al. 2022). DT cassava contains 147 copies of *AP2/ERF* genes (Wei et al. 2016). The DT millets, foxtail millet, and sorghum contain 190 and 187 *AP2/ERF*s (Agarwal et al. 2016), and only 83 *AP2/ERF*s are found in the desert moss *Bryum* (Li et al. 2018). Again, among DS species, legumes possess the highest number of *AP2/ERFs*. Mungbean contains 186 (Chen et al. 2022), pea 178 (Agarwal et al. 2016), and common bean contains 179 *AP2/ERF*s (Agarwal et al. 2016). Other DS legumes, such as chickpea and Medicago, contain 147 and 131 *AP2/ERF*s, respectively (Agarwal et al. 2016). In Solanaceae, tomato has 184 and potato 155 *AP2/ERFs* (Mariam et al. 2015). Other DS species, Arabidopsis and rice, contain 159 and 139 *AP2/ERF*s (Nakano et al. 2006). Our results indicate that two DT legumes, cowpea and *P. cineraria*, out of diploid plants, contain more copies of *AP2/ERF* TFs than other DT or DS plant families. In addition, legumes of higher ploidy, groundnut (*Arachis hypogaea*), and soybean (*Glycine max*) contain a much higher number of 400 and 440 *AP2/ERFs*, respectively (Zhao et al. 2022).

**Fig. 5.**
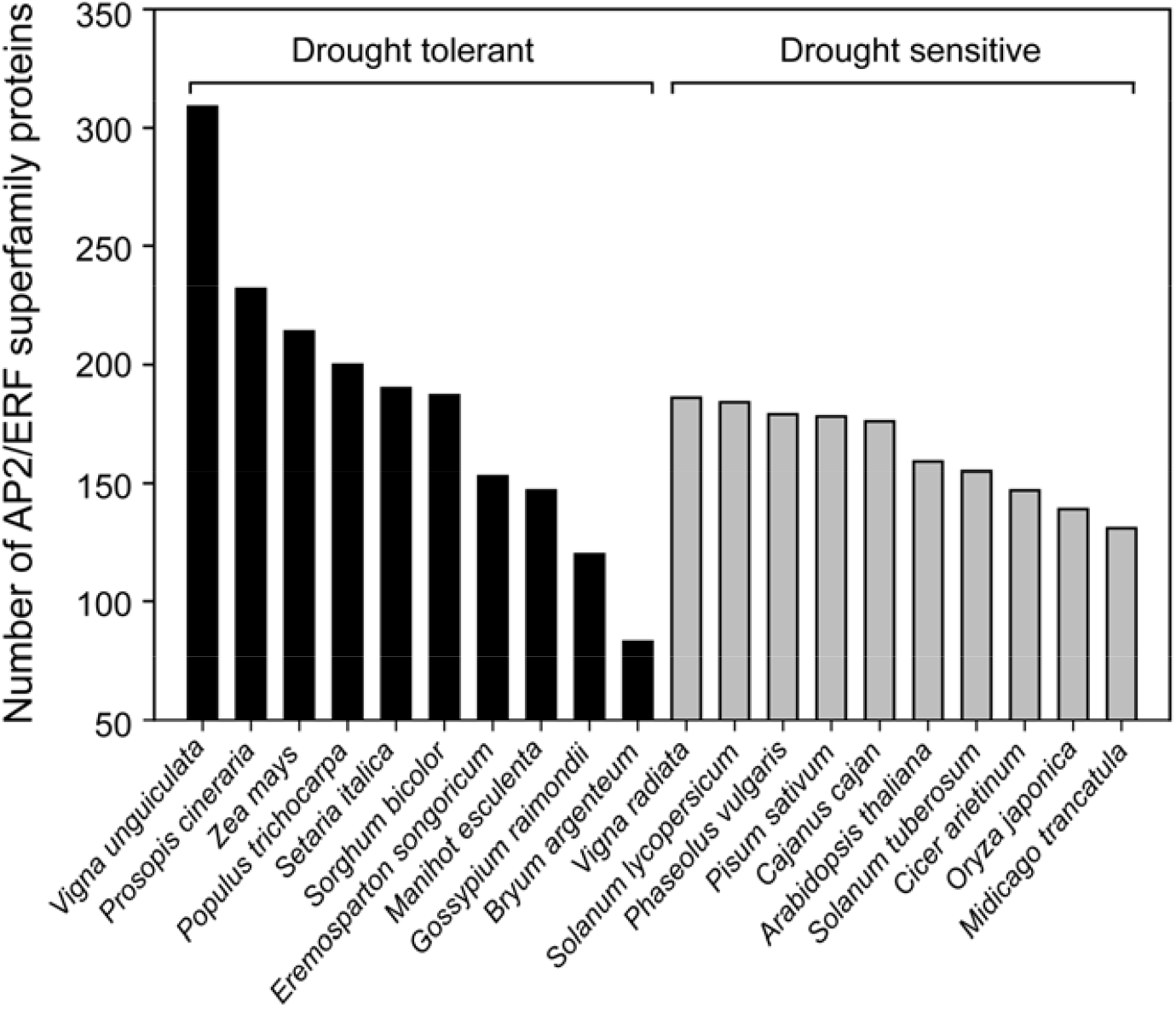
Copy number variation of AP2/ERF proteins between drought-tolerant (DT) and drought-sensitive (DS) species. Ten DT and 10 DS diploid plant species were selected to analyze copy number variations in the AP2/ERF superfamily proteins. The study was conducted through an exhaustive NCBI BLASTP search and copy numbers reported in the literature (see Methods). The vertical axis indicates the number of unique AP2/ERF proteins in each species. Black and grey bars indicate DT and species, respectively. Among the DT species, *P. cineraria* has the second highest copy number of AP2/ERF proteins.

### 3.8 The DNA-binding domains of *Prosopis cineraria* and *Arabidopsis thaliana* ERFs adopt the same fold but have chemical dissimilarity at a few positions

The protein data bank (PDB) has one solution structure of ERF1 GCC box-binding domain and one crystal structure of ERF/AP2 domain from *A. thaliana* (PDB id 1GCC and 5WX9, respectively, Fig. 6A, Fig. S1). Both the structures have the same fold (C□ RMSD = 1.31Å) consisting of ∼60 residues with three-stranded antiparallel β-sheet comprising strand 1, strand 2, and strand 3 connected by two turns and one □-helical region (Allen et al. 1998). Of these, four positively charged residues and two aromatic residues from the β-sheet (R150, R152, R162, and R170 in 1GCC; R16, R19, R21, and R39 in 5WX9; Fig. S1) are involved in specific hydrogen bonding with DNA-bases. We compared the amino acid sequences of AP2/ERF orthologs from *A. thaliana* and *P. cineraria* separately for each group of the phylogenetic tree (Fig. 2) by MSAs (Fig. S2). We found that most residues involved in DNA binding were fully conserved between the two species except for a few positions. We found clear segregation in the chemical nature (polar vs. non-polar) of a single position within the DBD between DT *P. cineraria* and DS *A. thaliana* in only two groups of the phylogenetic tree, VII and VIIIa. Hence, we modeled the full-length structures of four ERF or ERF-like proteins from groups VII and VIIIa of *A. thaliana* and *P. cineraria* AP2/ERF superfamily using AlphaFold2. These are (i) *A. thaliana* protein RAP2.2 (AT3G14230; UniProt ID: A0A5S9XC16) and (ii) *P. cineraria* protein ethylene-responsive transcription factor RAP2-3-like isoform X1 (XP_054790670.1) from group VII (Fig. 6B, C); and (iii) *A. thaliana* protein ERF9 (AT5G44210; UniProt ID: A0A654G7S4) and (iv) *P. cineraria* protein ethylene-responsive transcription factor 3-like (XP_054818156.1) from group VIIIa (Fig. 6D, E). All these four models show a similar fold in their DNA binding domain. When superimposed on the known solution structure 1GCC, the RMSD values considering the C□ atoms were within 0.7 Å, showing their high structural similarity. Interestingly, the DNA-bound complexes of these four proteins modeled from the latest online version of AlphaFold (AlphaFold3, Abramson et al. 2024) came out to be the same, except for a slight variation in the DNA conformation.

**Fig. 6:**
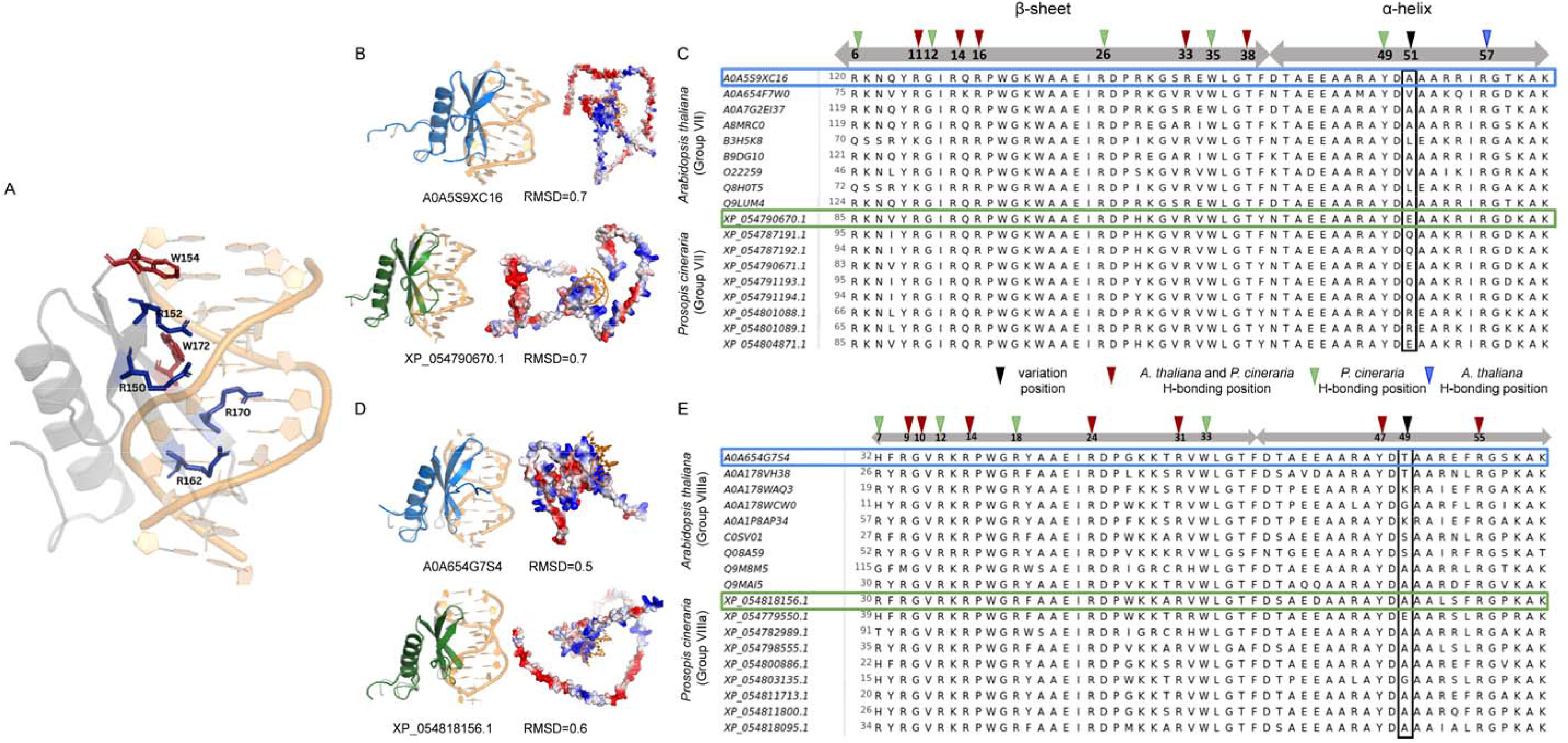
Structure and multiple sequence alignment of DNA binding domains of ERF proteins from *Prosopis cineraria* and *Arabidopsis thaliana*. **(A)** Crystal structure of DNA binding domain (DBD) of ERF protein from PDB (1GCC.pdb). **(B)** Modelled structures of DBD of ERF proteins from *A. thaliana* group VII (upper panel in blue cartoon) and *P. cineraria* group VII (lower panel in green cartoon). **(C)** MSA of homologous ERF sequences from group VII of *A. thaliana* and *P. cineraria*. **(D)** Modelled structures of DBD of ERF proteins from *A. thaliana* group VIIIa (upper panel in blue cartoon) and *P. cineraria* group VII (lower panel in green cartoon). In both **(B)** and **(D)**, the figures show the superimposition of ERF cartoon structure (shades of blue for *A. thaliana* and shades of green for *P. cineraria*) with 1GCC along with their root mean square deviation (RMSD). The electrostatic surface of the proteins is shown on the right, where blue and red represent positively and negatively charged surfaces, respectively. **(E)** MSA of ERF homologous sequences from group VIIIa of *A. thaliana* and *P. cineraria.* In both **(C)** and **(E)**, only a representative section from the DBD corresponding to the α-helix and the β-sheet region is shown. A difference in hydrogen bond donor position between the two species is highlighted with a black box and triangle. Similar triangles also show other specific hydrogen bonding positions (blue for *A. thaliana*, green for *P. cineraria*, and maroon when bonding was observed in both species).

While comparing the amino acid sequences from group VII ERFs, the critical residues involved in DNA binding and hydrogen bonding obtained from MD simulations in *P. cineraria* and *A. thaliana* are R11, R14, R16, R33, and T38 (Fig. 6C, right panel). The hydrogen bonds for *P. cineraria* are at G12, R26, T35, and Y49, while R57 was involved in hydrogen bonding only for *A. thaliana* (Fig. 6C). All these positions are fully conserved among the two species. However, we found differences at position 51 within the □-helical region. This contains aliphatic residues like alanine, valine, or leucine in *A. thaliana* and is fully conserved within the species, containing only non-polar aliphatic residues. The corresponding position in *P. cineraria* contains polar charged/uncharged residues like glutamine, glutamate, and arginine (Fig. 6C; highlighted in black triangle and box). On the other hand, the group VIIIa MSA indicated that the critical residues involved in hydrogen bonding in both *P. cineraria* and *A. thaliana* are R9, G10, R14, R24, R31, Y47, and R55, whereas those exclusively for *P. cineraria* are H/R7, R12, R18, W33; and none found exclusively for *A. thaliana*. In position 49, *P. cineraria* group VIIIa ERFs predominantly contain alanine, but *A. thaliana* has mostly polar residues like threonine, serine, and lysine (Fig. 6E). This variable position, though not directly involved in DNA binding, may have a role in initial DNA search and recognition (Freire-Rios et al. 2020).

### 3.9 Differences in hydrogen bonding profile and charge distribution at the N- and C-terminal tails between *Prosopis cineraria* and *Arabidopsis thaliana* ERFs might be crucial in their differential DNA search and binding

The AP2/ERF from *A. thaliana* and *P. cineraria* have long unstructured tails at the N- and C-terminal ends, having 83% and 75% of the total unstructured region in group VII, respectively. The respective percentages are 70% and 72% in group VIIIa. The group VII ERF from *A. thaliana* has an extended C-terminal tail compared to that of *P. cineraria*. We estimated the electrostatics and hydropathicity of the full-length proteins of *A. thaliana* and *P. cineraria* using the Kyte-Doolittle and Kyte-Woolridge scales, respectively. Although the DBD of group VII ERF from *P. cineraria* has similar hydropathicity, the N- and C-termini are more negatively charged than that of *A. thaliana* group VII ERF (Fig. 7A). Also, *A. thaliana* group VII ERF has an extended unstructured C-terminal region of ∼100 residues, with many consecutive negatively charged residues, as observed from the hydropathy plot. In contrast, the C-terminus of group VIIIa ERF from *P. cineraria* is more negatively charged than that of *A. thaliana.* The same trend was observed from the electrostatic profile of both species from both groups (Fig. 7A). The presence of negatively charged unstructured tails corroborates with a previously reported prevalence of D/E repeats in DNA/RNA binding proteins (Wang et al. 2023). We anticipate that these differential electrostatics in the long unstructured regions along with negatively charged repeats have a differential role in ERF proteins’ effective DNA search mechanisms between tolerant and sensitive species.

**Fig. 7.**
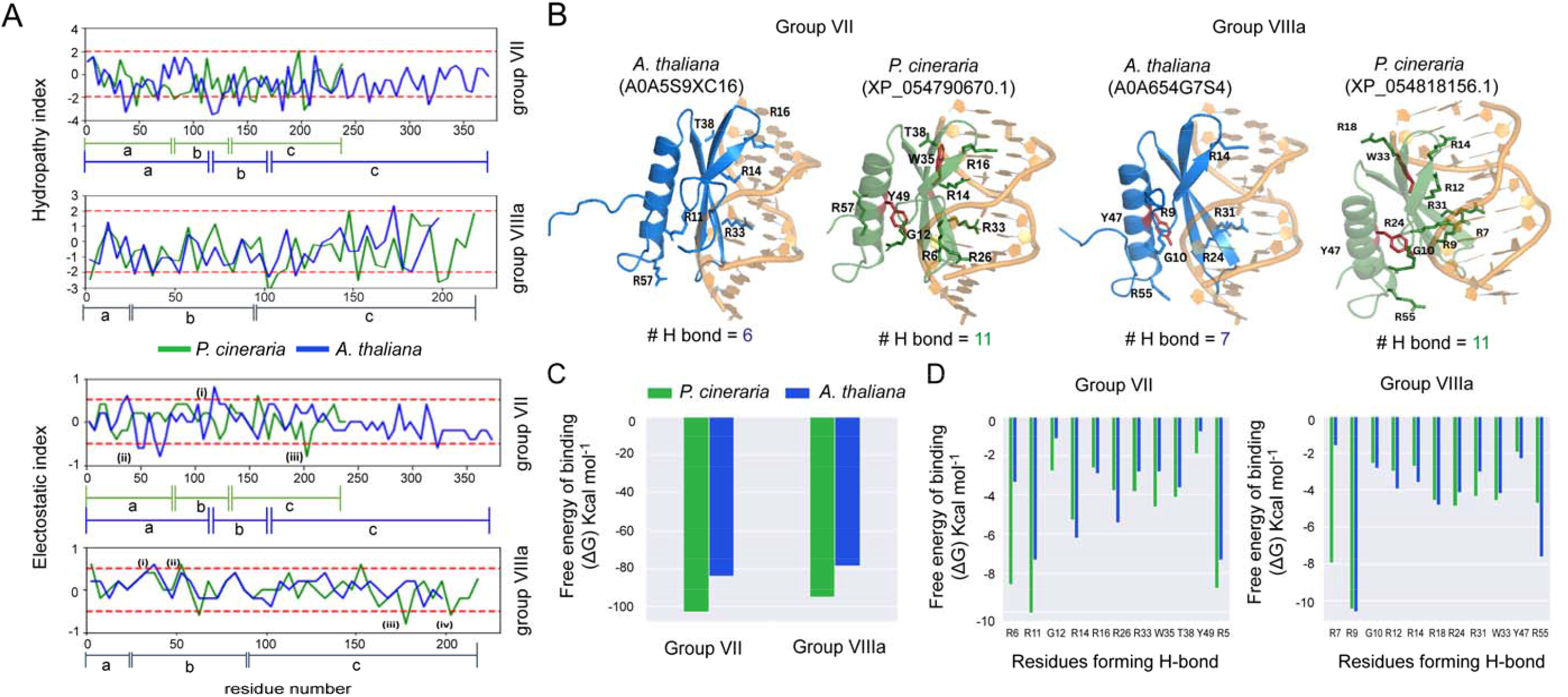
DNA binding interactions of ERF proteins from *Prosopis cineraria* and *Arabidopsis thaliana* determined from all-atom molecular dynamics (MD) simulations. **(A)** Hydropathy (top) and Electrostatics (bottom) profiles for the full-length ERF proteins of group VII and VIIIa from *A. thaliana* and *P. cineraria* showing their local hydrophobicity and charge along the amino acid sequence. For each residue position, the running average score for a window of five residues is plotted (see Method). In Group VII, peak (i) indicates positively charged residues in the DBD of *A. thaliana*, peak (ii) indicates the negatively charged region in the N-terminus of *A. thaliana* and peak (iii) shows a negatively charged region in the C-terminus of *P. cineraria.* In Group VIIIa, peaks (i) and peak (ii) indicate positively charged residues for *A. thaliana* and *P. cineraria* in the DBD region, while peaks (iii) and (iv) show negatively charged region in the C-terminus of *P. cineraria*. **(B)** Stable hydrogen bonds formed by the ERF DBD of *A. thaliana* and *P. cineraria* with the DNA in MD simulations. Protein-DNA complexes are shown in the cartoon, the hydrogen bond-forming residues in the sticks, and aromatic residues in red. The average number of stable hydrogen bonds in each case is shown at the bottom. Hydrogen bonds in each structure are calculated using the HBPLUS program. **(C)** Binding energy of DBD-DNA complexes calculated from MD trajectories using MM/GBSA method in AMBER package (see Method). Average values from 3 independent simulations are plotted in each category. **(D)** Residue-wise binding energy contributions of the DBD-DNA complexes. Only the residues making significant binding energy contributions (also highlighted in the MSA of Fig. 6) are shown.

We also performed three replicates of MD simulations for the four models mentioned above structures (two each from *A. thaliana* and *P. cineraria*) for a timescale of 200 ns to find stable hydrogen bonds and binding affinity. Moreover, we performed hydrogen bonding analysis on 2000 structures extracted from the 200 ns trajectory obtained by MD simulation. We did this for each of the four modeled structures from *A. thaliana* and *P. cineraria*. We found six stable hydrogen bonds present in the DNA binding domain of the representative group VII ERF protein of *A. thaliana,* whereas the representative *P. cineraria* group VII ERF contains eleven stable hydrogen bonds consistently observed in at least two of the three replicates of MD simulations). Like group VII, a significantly higher number of stable hydrogen bonds were also observed in the group VIIIa structure of *P. cineraria* compared to that of *A. thaliana* (11 vs. 7, Fig. 7B). The hydrogen bonding positions are mapped onto the structures, and the details of the hydrogen bonding residues are represented in Table S5. These differences in hydrogen bonding and the chemical nature of the residue positions, particularly position 51 in group VII, may play a critical role in differential DNA binding affinity and specificities between the two species. To further confirm this hypothesis, we performed MM/GBSA to calculate the DNA-binding affinity from the trajectory data of all four models. The results indicated that the ERF of *P. cineraria* from both groups VII and VIIIa have higher DNA binding affinity than that of *A. thaliana,* with a significantly lower free energy of binding ΔG (Fig. 7C). We also conducted residue-wise energy decomposition analysis. The corresponding energy values for critical residues in the DBD are represented in Fig. 7D. Eight of the 11 residues involved in hydrogen bonding have higher contributions to the binding energy in ERF group VII of *P. cineraria.* In contrast, four residues in ERF group VIIIa showed higher contributions than corresponding residues of *A. thaliana*.

### 3.10 DREB proteins have similar structures in drought-tolerant and drought-sensitive species

We extended our analyses to study the DREB1 TF for DT and DS species. Without structural data for DREB1 in PDB, we modeled the structures from eight species, four from DT and DS groups, using AlphaFold2 (Fig. S3). All these structures were superimposed on the *P. cineraria* ERF1 structure (PDB: 1GCC) to understand their structural similarities. Remarkably, the DNA-binding region showed the same fold containing the three-strand β-sheet and the □-helix with the maximum RMSD of 0.7Å among all eight species. The sequences were compared similarly by conducting MSA. Interestingly, we found a distinction in MSA between cereals and non-cereals in the β-sheet region of the DBD. Cereals like sorghum, foxtail millet, and rice had a conserved region (GRRG) depicted by a red box (Fig. S3). In contrast, the corresponding region in non-cereals exhibited a stronger polar/charged character, represented by a black box (NKN/KK/T). AlphaFold2 models of the DREB2 for all eight species also resulted in the same fold for the DBDs (maximum RMSD of 0.8 Å among all eight species; Fig. S4). However, a DREB2 MSA demonstrated that DREB2 proteins’ DBD is conserved across tolerant and sensitive species with no significant difference in the critical DNA-binding residues.

## 4. DISCUSSION

*P. cineraria*, a vital component of the Indian Thar desert, is a leguminous tree that exhibits high resistance to abiotic stresses such as drought, heat, salt, cold, and nutrient deficiency (Rai et al. 2021). The recent sequencing of the whole genome of *P. cineraria* (Sudalaimuthuasari et al. 2022) paves the way for discovering its hitherto unknown molecular mechanisms of stress tolerance. This study aims to fill this knowledge gap with a drought-specific transcriptome analysis and a comprehensive genome-wide analysis of the *AP2/ERF* genes in the Indian cultivar of *P. cineraria*.

The transcriptome analysis of *P. cineraria* seedlings under drought stress showed an abundance of up-regulated genes compared to down-regulated genes. The functional enrichment analysis of the up-regulated genes revealed their roles in cell cycle regulation, possibly affecting growth under drought (Qi and Zhnag 2019) and the olefinic compound metabolic process, a drought stress metabolic marker (Noleto-Dias et al. 2023). Pathway enrichment analysis revealed that the up-regulated genes primarily functioned in phytohormone and MAPK signaling pathways, signatures of drought responses. The PPI networks confirmed that the developmental, defense-related genes were down-regulated, and the induced genes belonged to the stress-responsive TFs and signaling proteins. Several ribosomal proteins constituting the protein synthesis machinery were identified as hub genes (Table S3). These proteins play a pivotal role in stress adaptation by participating in regulatory mechanisms, often targeting the antioxidation machinery (Saha et al. 2017; Shiraku et al. 2021; Zhang et al. 2023). In addition, histone H3.2 and H4, and heat shock proteins also emerged as hub genes in the up-regulated gene network. Previously, these proteins were differentially expressed under drought stress in various plants (Li et al. 2016; Pan et al. 2018). While histones are involved in reprogramming gene expression under drought (Xin et al. 2021), heat shock proteins protect other cellular proteins from misfolding under stress conditions (Guo et al. 2023).

Orthologs of the genes induced by drought in *P. cineraria* had reported roles in drought tolerance (Table S1). For example, late embryogenesis abundant (LEA) family proteins *embryonic protein DC-8*, *ECP63*, and *LEA2-like* proteins were strongly up-regulated. The *A. thaliana* ortholog of *ECP63* is involved in the desiccation tolerance of seeds (Yang et al. 1997). The induction of *LEA* genes in adult plants is the hallmark of drought, and these proteins protect the cytoplasm from desiccation (Hong-Bo, Zong-Suo, and Ming-An, 2005). A *P. cineraria* ortholog of *DREB1A* was highly up-regulated, which interacts with the DRE sequence of the promoter of dehydration-responsive *rd29A* gene (Qiang et al. 1998), having a pivotal role in drought tolerance (Wang et al. 2022). *MOTHER of FT and TFL1-like* (*MFT*)’s Arabidopsis ortholog promoted the ABA-induced genes (Vaistij et al. 2018) regulating ABA signaling (Wanyan et al. 2010). Orthologs of *lipoxygenase 3* induced in *P. cineraria* are expressed highly under drought (Gigon et al. 2004). Drought affects the transport of carbohydrates and phloem loading rate (Salmon et al. 2019). In this context, a couple of carbohydrate transporters like *galactinol-sucrose galactosyltransferase 5*, whose ortholog was also up-regulated in response to drought in wheat (Wang et al. 2019), and *sieve element occlusion B*-like were induced in our study. Another metabolic enzyme, *glutathione S-transferase*, was also highly induced, which reduces cellular damage by drought-induced ROS in rice by enhancing glutathione binding to electrophilic compounds (Li et al. 2003). An ERF TF, *ABR1-like*, induced in *P. cineraria*, negatively regulates ABA responses (Pandey et al. 2005) but improves drought tolerance by enabling soluble sugar accumulation from starch degradation (Zhang et al. 2023). The *A. thaliana* ortholog of *ERF053-like* is induced early in drought, and its overexpression leads to drought tolerance (Cheng et al. 2012, Hsieh et al. 2013). The overexpression of a *1-aminocyclopropane-1-carboxylate oxidase-like* ortholog, functioning in ethylene biosynthesis, led to an elevated abiotic stress tolerance in *Petunia hybrida* (Naing et al. 2021). Another induced gene, *trehalose-phosphate phosphatase C*, responsible for the conversion of trehalose-6-phosphate to trehalose, when overexpressed, improved drought tolerance in Arabidopsis as well as rice (Lin et al. 2019; Garg et al. 2002). *Short-chain dehydrogenase reductase 3b* was strongly up-regulated in *P. cineraria* and is involved in the ABA biosynthesis pathway (Cheng et al. 2002). *Cysteine proteinase inhibitor B-like* orthologs are induced in response to drought (Abdel-Ghany et al. 2020) and are under direct transcriptional control of DREB TFs. Their overexpression improves drought tolerance by suppressing programmed cell death due to oxidative stress under drought (Zhang et al. 2008). *9-cis-epoxycarotenoid dioxygenase 1* (*NCED1*), up-regulated in our study, is involved in the biosynthesis of ABA, conferring drought tolerance (Iuchi et al. 2001; Kalladan et al. 2019; He et al. 2018). A *cytochrome P450 94C1-like* gene was also highly induced in *P. cineraria*, functioning in seed germination, root elongation, and possibly in drought response in cotton (Gu et al. 2023).

Among the down-regulated genes (Table S1), *short-chain aldehyde dehydrogenase 1-like* belongs to the family of NADP-dependent oxidoreductase (Moummou et al. 2012) and is expressed differentially in response to drought stress in garlic (Twaji et al. 2023). Receptor-like protein kinase FERONIA was also highly down-regulated in our study, which is a positive regulator of the auxin signaling pathway, thereby inhibiting ABA signaling through *ABI2* in Arabidopsis (Yu et al. 2012). Arabidopsis *xyloglucan endotransglucosylase/hydrolase protein 6* has a time-dependent differential expression pattern in response to drought (Han et al. 2023, Tenhaken 2015), whose ortholog was also down-regulated in our study. Certain *P. cineraria* paralogs of *Cytosolic sulfotransferase* were highly induced, and some were down-regulated. Orthologs of these genes have a tissue-specific and differential expression pattern under drought stress in *Brassica rapa L*. (Jin et al. 2019) and protect plants from oxidative damage during drought.

The up-regulation of 54 and down-regulation of six *AP2/ERF* genes in *P. cineraria* under drought stress prompted us to conduct a comprehensive genome-wide analysis of this desert species. Based on the classification in *A.thaliana*, *AP2*/*ERF* genes of *P. cineraria* were phylogenetically classified into 15 groups (Nakano et al. 2006; Li et al. 2021). The ML method led to a more accurate clustering of *P. cineraria* AP2/ERF proteins with their *A. thaliana* orthologs than other methods of phylogenetic tree construction, closely following the grouping described earlier (Sakuma et al. 2002; Nakano et al. 2006). The ML method has been used earlier for genome-wide analysis of AP2/ERF proteins in other plants (Lakhwani et al. 2016; Xing et al. 2021; He et al. 2023). Earlier, the ML method showed more accuracy than the neighbor-joining maximum composite likelihood (NJ-MCL) method in tree construction when the sequences are of the order of a few hundred (Som 2009). We found an absence of RAV genes in *P. cineraria* similar to another desert legume, *E. songoricum* (Zhao et al. 2022). The number of *ERF* genes was higher than the *DREB* genes, similar to other species (Agarwal et al. 2016).

We conducted a qPCR analysis of five genes selected from groups III (A-1), IV (A-2), III (A-4), Xb (B-3), and AP2 to study the time-dependent drought stress responses. This revealed different *AP2*, *DREB*, and *ERF* gene expression patterns under drought (Fig. 3A). The early-induced expression of genes of groups III (A-1) and AP2 were suppressed under long-term treatment. *DREB* and *ERF* genes are promptly induced in response to drought across all higher plants within a few hours of stress exposure, and the expression levels of different paralogs undergo differential spatiotemporal regulation (Dossa et al. 2016; Guo et al. 2016; Cui et al. 2021). Hence, the different expression patterns of various subgroups of *P. cineraria AP2/ERF* genes indicate their complex regulation for early and late responses to drought. Several *cis*-acting elements for response to various abiotic stresses were identified in the promoters of *P. cineraria AP2/ERF* genes (Fig. 3B). The roles of ABRE and DRE/CRT in ABA-dependent and -independent gene expression of DREBs in response to abiotic stress are well known (Shinozaki and Yamaguchi-Shinozaki, 2007). *Cis*-elements indicate that groups I, IIa, VII, and AP2 were under feedback regulation by DREBs, while groups I, IIa, III, IV, and VII were under ERF regulation. Other *cis*-elements, including MYB and MYC, indicate the complex regulation of *P. cineraria* AP2/ERFs by different families of TFs under drought stress (Abe et al. 2003). Moreover, we found that the promoter regions of *AP2/ERF* genes contained elements related to light response, such as the G-box and GATA motif (Luo et al. 2010; Schröder et al. 2023). This indicates that these genes might be under regulation by light-dependent signaling and cross-talks with high-light stress response pathways. Additionally, defense-related elements, such as TCA and TGA, were identified in *P. cineraria AP2/ERF* genes. This indicates the cross-talk of *AP2/ERF* genes with biotic stress response pathways. Previously, *AP2/ERF-DREB* family genes were found to be involved in biotic stress resistance in potatoes (Chacón-Cerdas et al. 2020).

The gene structures and conserved protein motifs were analyzed to gain further insights into the evolutionary relationships of *AP2/ERF* gene families. Our analysis indicated that *P. cineraria* DREBs contained 15 (3.79%), and the ERFs contained 380 (96.20%) out of the total number of 395 introns in all the genes of the AP2/ERF superfamily (Fig. 4). Additionally, group III had the highest number of introns for DREBs, i.e., 8 out of 15 (53.33%). Similarly, Group VII consisted of a maximum number of introns for *ERF* genes, including 17 of the 50 (34%). The DNA-binding AP2 domain was ubiquitous in all the DREB and ERF proteins of all the groups, suggesting that it was highly conserved during the evolution of *P. cineraria* (Fig. 5; Fig. S2). Two AP2 domains were present in the members of the AP2 family, like that of *A. thaliana*. A negative regulatory domain (NRD) present in DREB2A just downstream to the AP2 domain contains a PEST sequence with phosphorylation sites, which target these proteins for degradation under non-stressful conditions and whose removal leads to constitutively active forms of the DREB proteins (Sakuma et al. 2006; Lata and Prasad 2011; Sadhukhan et al. 2014). We detected PEST sequences in most AP2/ERF proteins of *P. cineraria*. PEST sequence-mediated regulation of dehydrin proteins was also observed in the cactus *Opuntia streptacantha* (Salazar-Retana et al. 2019). Our results indicate protein degradation as a regulation strategy of AP2/ERF proteins in *P. cineraria,* like other plants. On the other hand, the absence of PEST sequences in group VII and the majority of group V and X members indicate a lack of such regulation in some ERF subgroups of *P. cineraria*. In addition, four members of group IX contain conserved structural features of dehydrins at the C-terminal region comprising a polyserine region followed by charged residues and a lysine-rich region involved in protein-lipid interaction (Close et al. 1989). Dehydrins belong to the group II LEA protein family, which protects membranes during drought and extreme temperature stresses (Puhakainen et al. 2004; Hundertmark and Hincha 2008; Hand et al. 2011). This suggests additional protective functions of the four group IX ERFs of *P. cineraria* under abiotic stress, other than TF activity.

The desert regions of Arabia and the Indian Thar are diverse in rainfall and climate. Hence, considerable genetic diversity in the Arabian and Indian *P. cineraria* genomes is expected. We looked at the nucleotide sequence level variations in 60 *AP2/ERF* genes differentially expressed under drought in our transcriptome, revealing various SNPs, MNPs, and INDELs between the two cultivars. Some of these polymorphisms translated into amino acid changes (Table 2), suggesting the possible involvement of these proteins in the drought tolerance of *P. cineraria*. Natural variations in *DREB* and *ERF* genes within a species have been occasionally associated with drought tolerance. Recently, natural variations in *DREB* genes were examined in 368 varieties of maize, including SNPs in *ZmDREB2.7* and their relationship with drought resistance (Liu et al. 2023). Moreover, an allele-specific marker (ASM) was developed through SNP in *SiDREB2* with drought tolerance in foxtail millet, for marker-aided drought tolerance breeding of foxtail millet (Lata et al. 2011). However, the functional implications of the polymorphisms between *AP2/ERFs* genes from the Arabian and Indian cultivars of *P. cineraria* need to be explored in the future.

Nature employs a dual strategy of CNVs and accumulation of protein mutations, leading to structural differences enabling the evolution of genomic complexity (Keel, Lindholm-Perry, and Snelling 2016; Jayaraman et al. 2022). CNVs of various genes have been linked to different plant phenotypes, including stress tolerance (Żmieńko et al. 2014). CNV analysis of *AP2/ERF* genes in DT and DS species revealed the highest copies in DT diploid legumes, including cowpea and *P. cineraria* (Fig. 5). Due to genome duplication events, even higher copies of AP2/ERFs can be observed in legumes of higher ploidy, viz., partially diploidized tetraploid soybean (Singh and Hymowitz 1988) and segmental allotetraploid groundnut (Bertioli et al. 2019). Similarly, tetraploid cotton species *G. barbadense* and *G. hirsutum* contain 213 and 220 *AP2/ERF* genes, more than diploid cotton (Zafar et al. 2022; Zhao et al. 2022). DT plants have a wide variation in the copy numbers of AP2/ERFs, suggesting that drought tolerance does not solely depend on the copy number of *AP2/ERF* genes. The DT bryophyte had the least number of AP2/ERF proteins, suggesting the copy number of AP2/ERFs is a function of organismal complexity. However, DT legumes possessed higher copies of AP2/ERFs than DS legumes, indicating an important role of these TF families in the drought tolerance of legumes. Hence, higher copies of AP2/ERFs partly explain the higher drought tolerance of *P. cineraria*. The number of ERF family members was the highest in *P. cineraria*, followed by DREBs and other minor families, similar to other species (Zhao et al. 2022).

We modeled the 3D structures of DREB and ERF proteins of DT *P. cineraria* and compared them with orthologs in DS *A. thaliana* to explore any structural modifications of AP2/ERF TFs in DT species. We found that the DBD of DREBs and ERFs are well conserved at the sequence and structural levels among *P. cineraria* and *A.thaliana*. The structures fold similarly, consisting of three beta-strands and one alpha-helix across species. However, differences exist in electrostatics at some positions of the DBD in groups VII and VIIIa ERFs between DS *A. thaliana* and DT *P. cineraria*. The asymmetrical electrostatic charge distribution has a critical role in binding site recognition among transcription factors (Pal et al. 2020). Also, there exists ∼70-80% disorderedness in the N- and C-terminals of the ERF proteins in both groups, especially the group VII ERFs of *A.thaliana* have an extended unstructured negatively-charged tail, that is earlier known to influence the DNA search mechanism among DNA-binding proteins (Wang et al. 2023). Unstructured tails are widely found across proteomes and are known to have important regulatory roles in improving the stability and binding specificity of transcription factors (Bigman et al. 2022, Zaharias et al. 2021). Further analysis from MD simulation indicated more hydrogen bonding interactions between the beta-strand region of the ERF DBD and DNA in DT *P. cineraria.* This characteristic of additional interactions is observed in both the ERF groups VII and VIIIa. Hydrogen bonds are generally known to play a critical role in DNA-binding proteins’ assembly formation (Dey et al. 2012). Freire-Rios et al. (2020) reported that a single extra hydrogen bonding generally leads to a higher binding affinity between auxin-response transcription factors (ARFs) and auxin-responsive promoter *cis*-elements, validated by biochemical and *in vivo* functional experiments. In our case, from the simulation data, we also found a higher binding affinity of *P. cineraria* group VII and VIIIa ERFs to the GCC box DNA *cis-*element than that of *A.thaliana,* most likely due to the additional 4-5 hydrogen bonds, corroborating with earlier reports in plants. Future experimental validation of these important *in-silico* observations will throw light on the evolution of the AP2/ERF TFs in extremophile desert species like *P. cineraria*. In conclusion, our results indicate copy number expansion and structural modifications in ERF proteins as two evolutionary strategies for acquiring drought tolerance in plants. Our results will aid in improving drought tolerance in *P. cineraria* and provide tools for engineering drought tolerance in sensitive crop species.

## Declaration statements

## Data availability statement

The raw transcriptome sequences are available at NCBI Sequence Read Archive with a BioProject accession number PRJNA1023008.

## Author Contributions

AS conceived the study. VD conducted most of the primary bioinformatic sequence analysis. DM conducted stress experiments, transcriptome sequencing, and qPCR. RSS, AKS, and PY analyzed the transcriptome data. AP and SD conceived and performed computational experiments using structures. AP, A, and SD analyzed the structural bioinformatics data. AS and SD acquired funding. PY, AP, SD, and AS supervised the work. AD assisted in bioinformatic sequence analysis. VD, DM, A, AP, SD, and AS wrote the manuscript with input from the rest of the authors.

## Acknowledgments

VD and DM thank the Ministry of Education, Government of India, for their doctoral fellowships. A acknowledges the Council of Scientific & Industrial Research for her doctoral fellowship.

## Supplementary Data

**Supplementary Fig. S1**: Structures of DNA-binding domain of ERF from *Arabidopsis thaliana* and *Prosopis cineraria.* (A, B) Cartoon representation of the DNA binding ERF (1GCC, NMR structure, in grey) and AP2/ERF domain (5WX9, x-ray structure, in grey) from Arabidopsis in complex with its target DNA fragment (in orange). Residues involved in specific interactions with DNA are shown in blue (positively charged) and red (aromatic) sticks.

**Supplementary Fig. S2.** Multiple sequence alignments of AP2/ERF superfamily proteins of *Prosopis cineraria* and *Arabidopsis thaliana*.

**Supplementary Fig. S3.** Structural comparison of DREB1 proteins from drought tolerant and sensitive species.

**Supplementary Fig. S4.** Structural comparison of DREB2 proteins from drought tolerant and sensitive species.

**Supplementary Table S1.** Genes differentially expressed in *Prosopis cineraria* under drought stress.

**Supplementary Table S2**. Functional enrichment analysis of the gene networks of *Prosopis cineraria* under drought.

**Supplementary Table S3.** Hub gene identification in *Prosopis cineraria* under drought stress.

**Supplementary Table S4.** Oligonucleotides used in the quantitative real-time PCR analysis.

**Supplementary Table S5.** Statistics of stable hydrogen bonds between the DNA binding domain of ERF group VII and VIIIa from *Arabidopsis thaliana* and *Prosopis cineraria* and the DNA in all-atom MD simulations.

