## Supplemental files for "AP2/ERF transcription factors enriched in the drought-response transcriptome of the Thar desert tree *Prosopis cineraria* show higher copy number and greater DNA-binding affinity than orthologs in drought-sensitive species"

### Supplementary figure 1

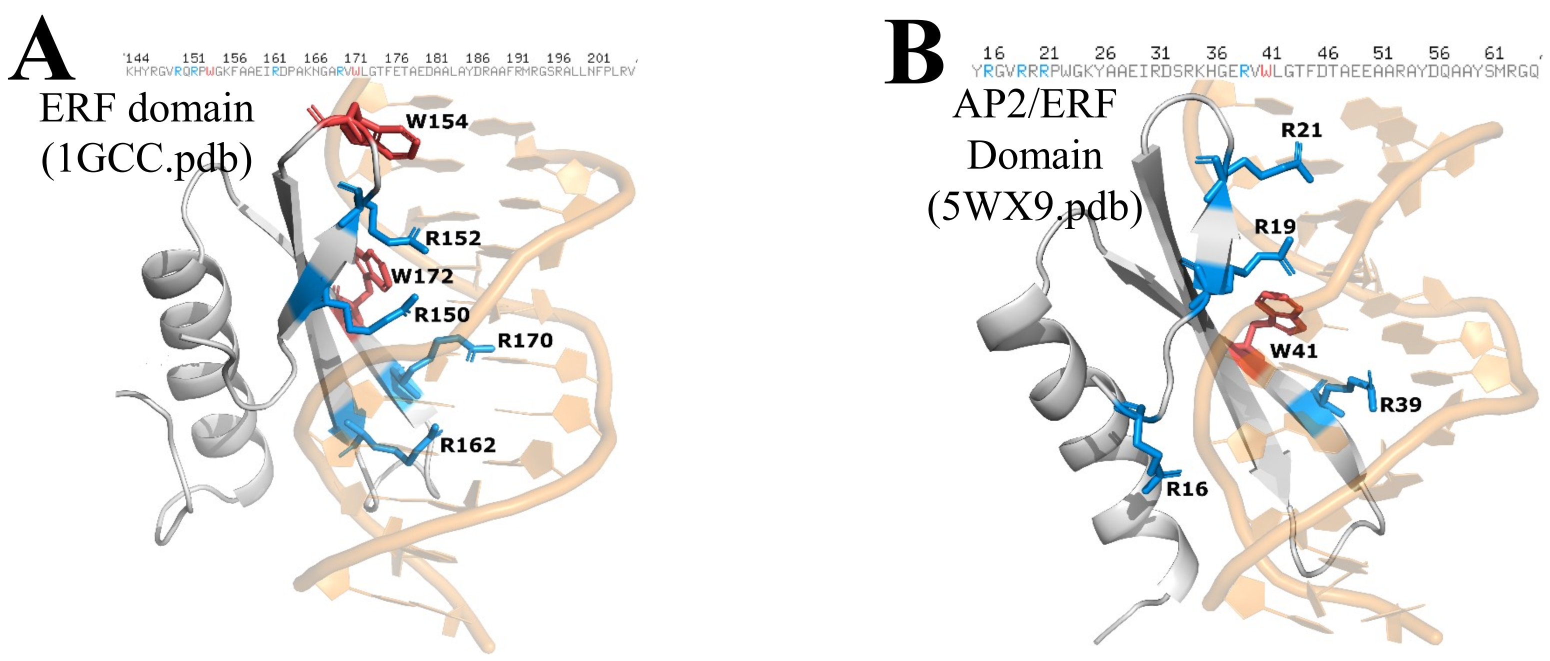


Figure S1: Structures of DNA-binding domain of ERF from *Arabidopsis thaliana* and *Prosopis cineraria*. (A, B) Cartoon representation of the DNA binding ERF (1GCC, NMR structure, in grey) and AP2/ERF domain (5WX9, x-ray structure, in grey) from *Arabidopsis* in complex with its target DNA fragment (in orange). Residues involved in specific interaction with DNA are shown in blue (positively charged) and red (aromatic) sticks.

### Figure S2

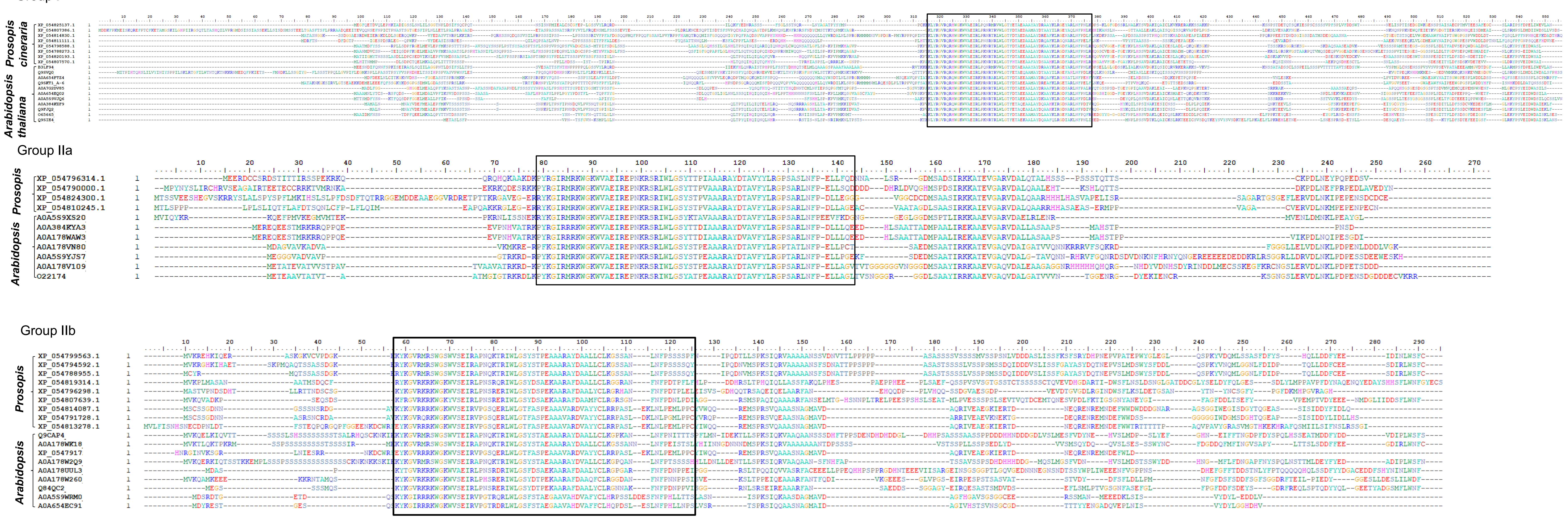


Figure S2 Contd..

Group III

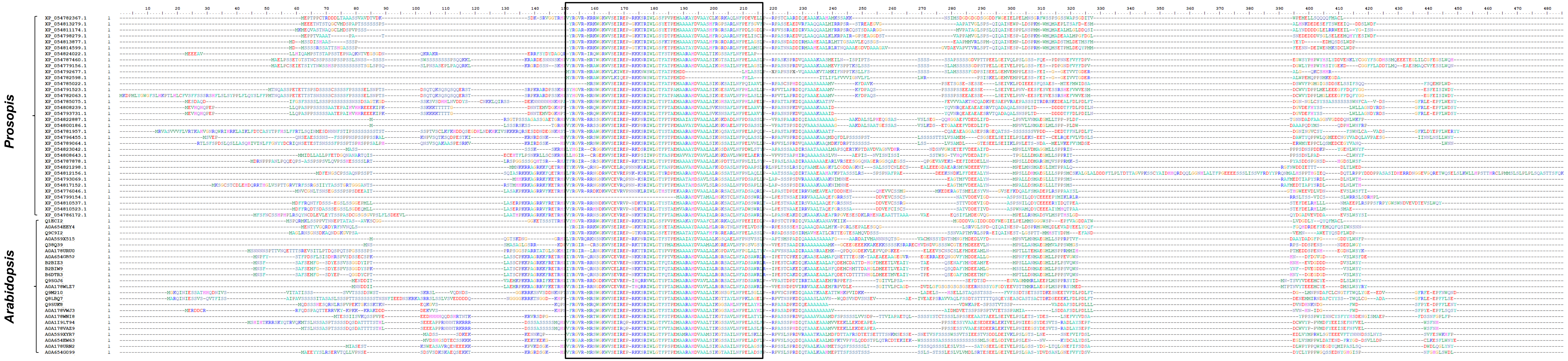


Figure S2 Contd..

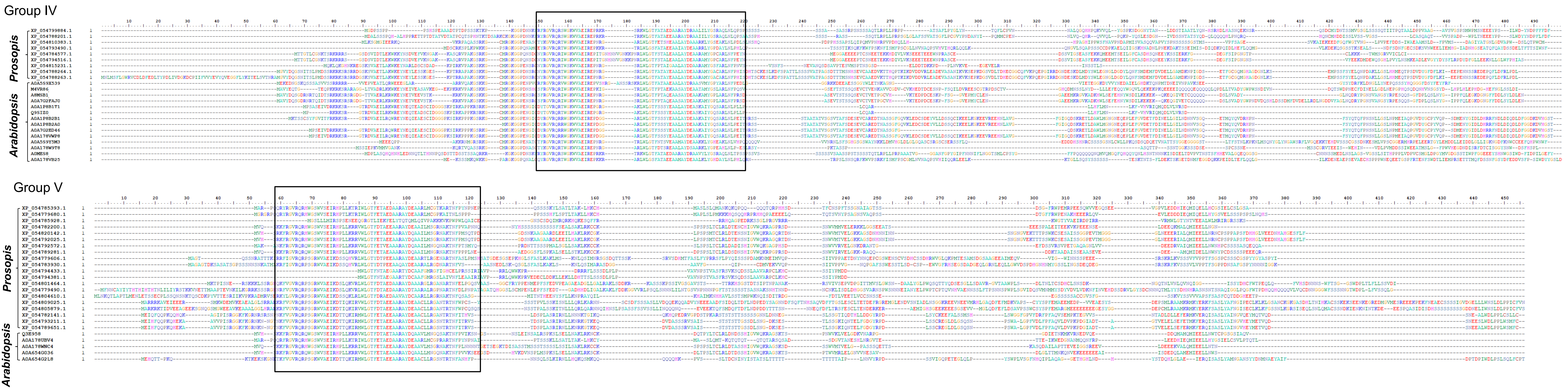


Figure S2 Contd..

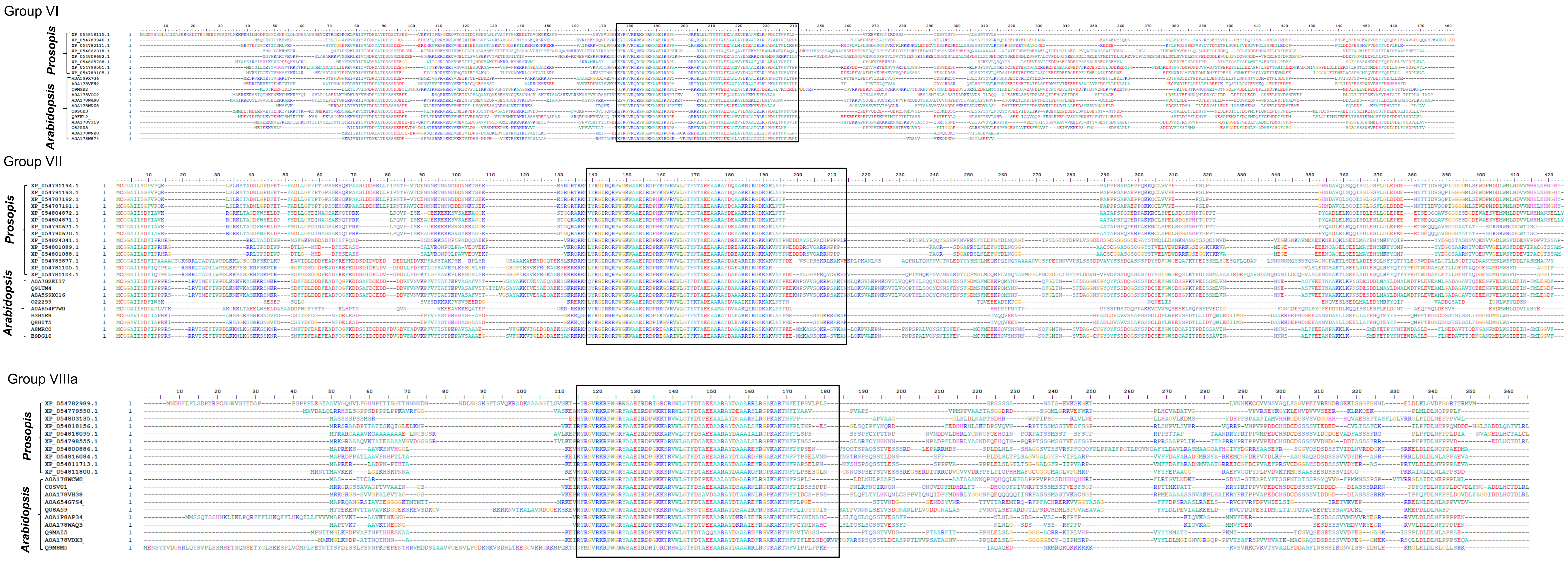


Figure S4 Contd..

Group VIIIb

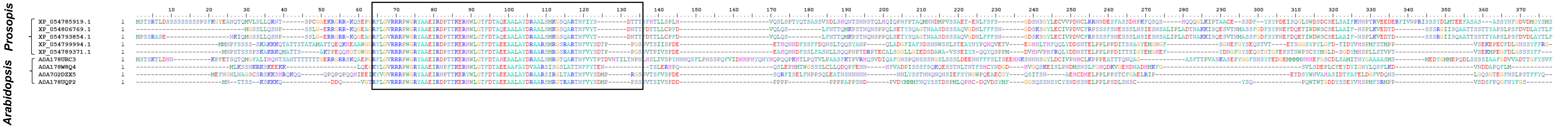


Group IX

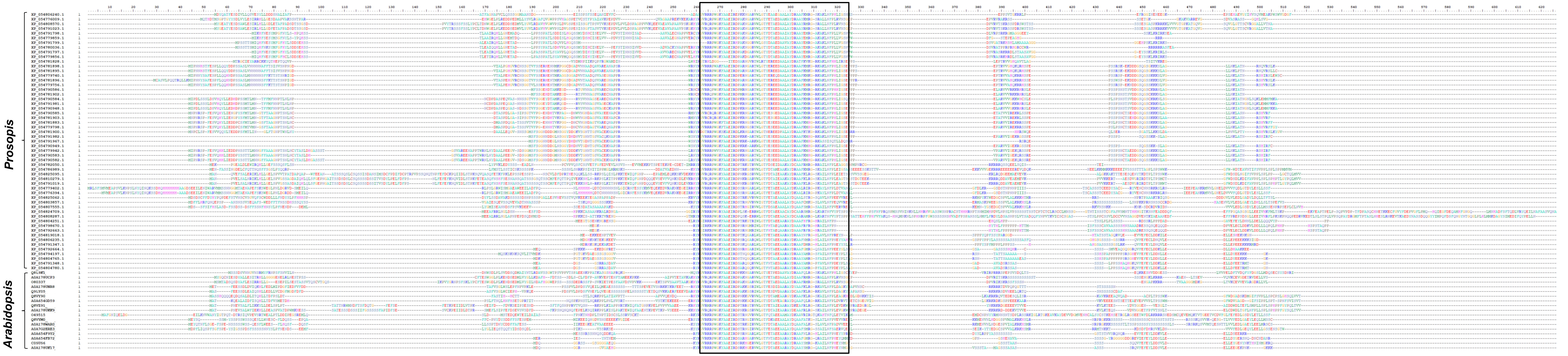


Figure S2 Contd..

Group AP2

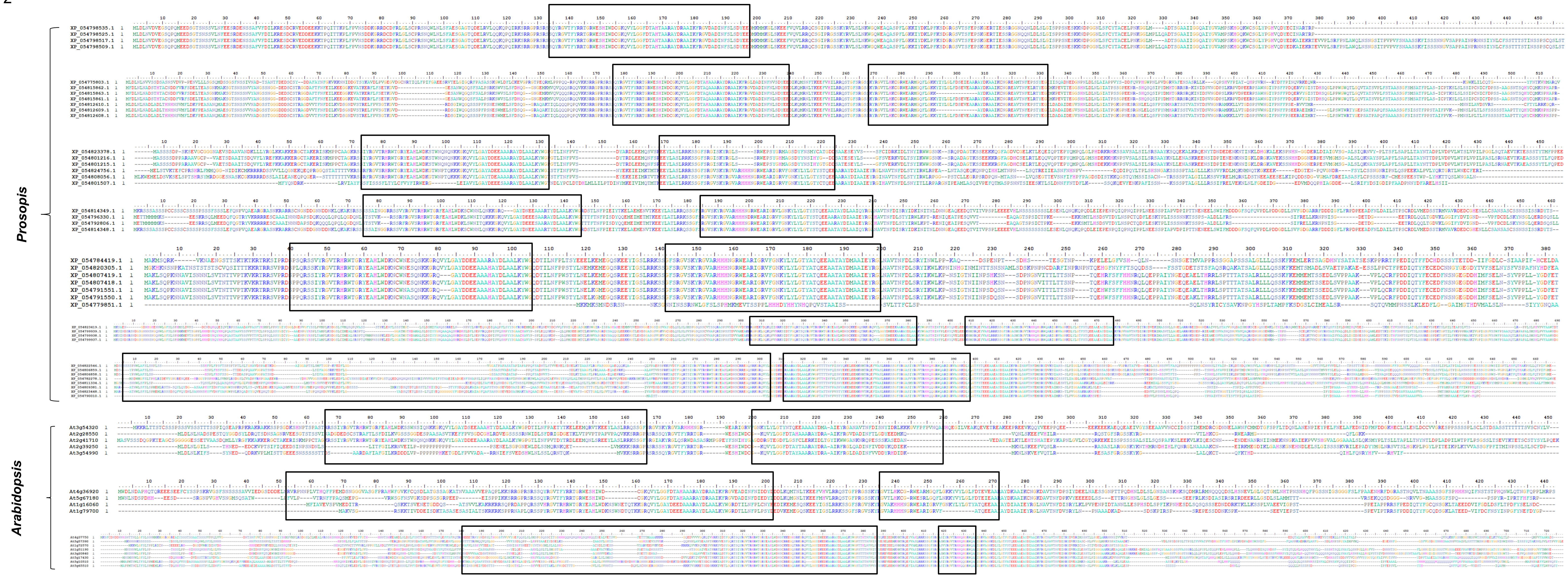


Figure S4 Contd..

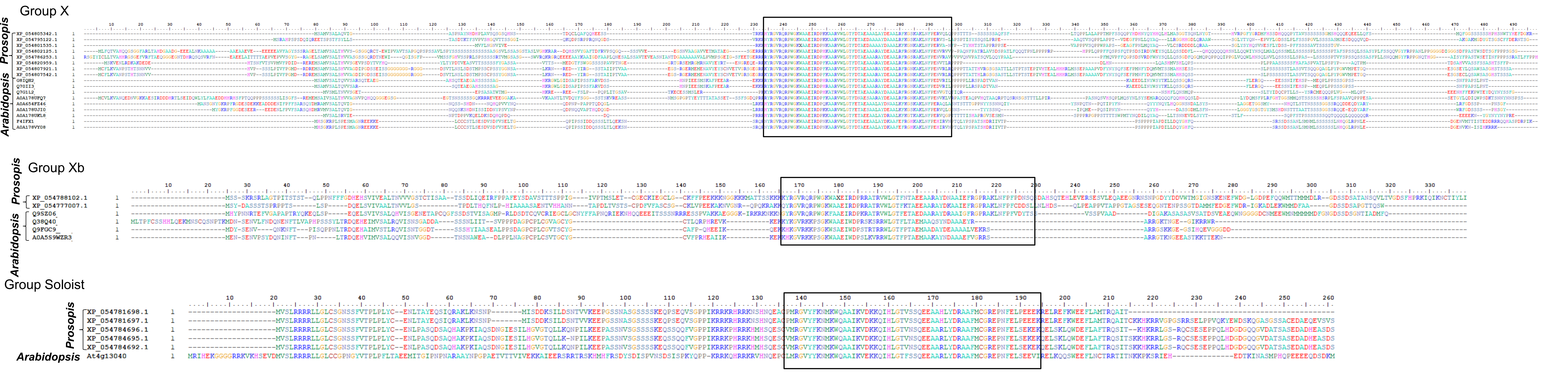


**Figure S2 Multiple sequence alignments of AP2/ERF superfamily proteins of *Prosopis cineraria* and *Arabidopsis thaliana*.** The alignments were conducted by CLUSTALW in BioEdit software, separately for each group of the phylogenetic tree (Figure 2). The black boxes indicate AP2 domains.

Figure S3

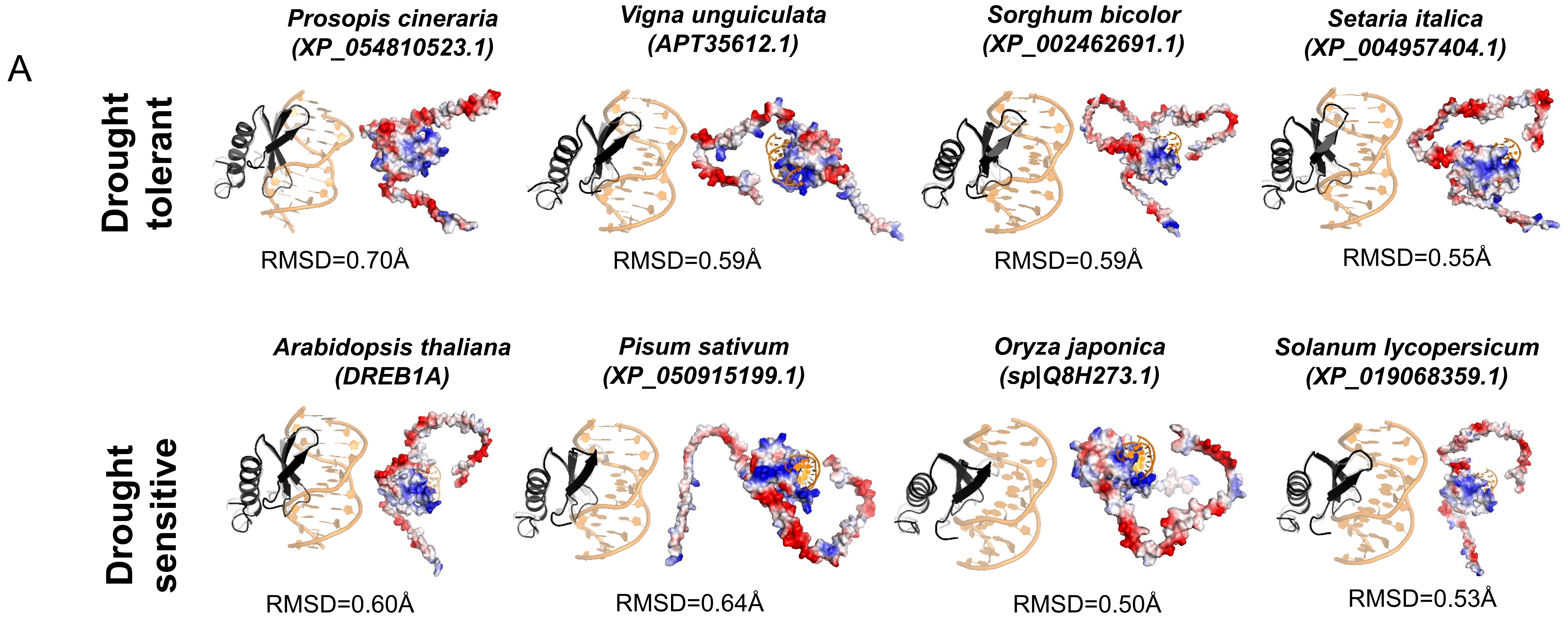


Figure S3 Contd..

Figure S3

B

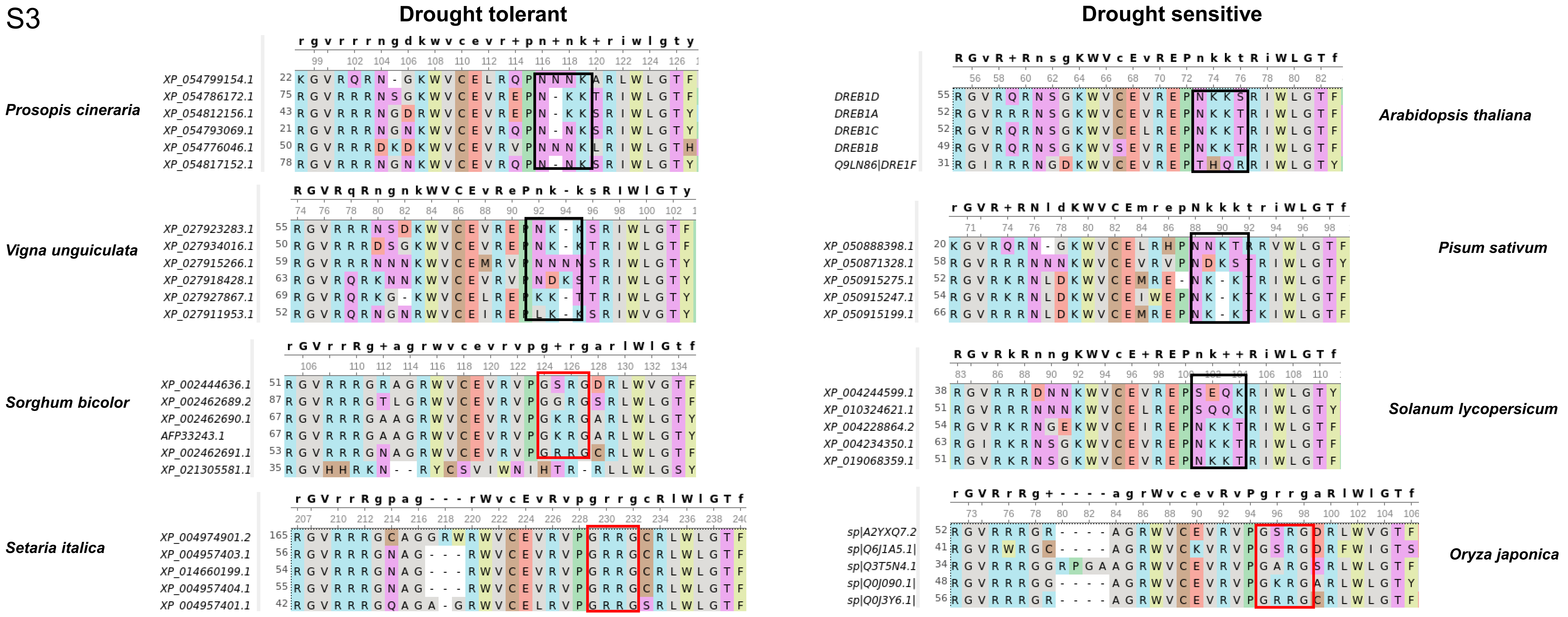

Drought tolerant
Drought sensitive
Prosopis cineraria
XP_054799154.1
XP_054786172.1
XP_054812156.1
XP_054793069.1
XP_054776046.1
XP_054817152.1
Vigna unguiculata
XP_027923283.1
XP_027934016.1
XP_027915266.1
XP_027918428.1
XP_027927867.1
XP_027911953.1
Sorghum bicolor
XP_002444636.1
XP_002462689.2
XP_002462690.1
AFP33243.1
XP_002462691.1
XP_021305581.1
Setaria italica
XP_004974901.2
XP_004957403.1
XP_014660199.1
XP_004957404.1
XP_004957401.1
Arabidopsis thaliana
DREB1D
DREB1A
DREB1C
DREB1B
Q9LN86|DRE1F
Pisum sativum
XP_050888398.1
XP_050871328.1
XP_050915275.1
XP_050915247.1
XP_050915199.1
Solanum lycopersicum
XP_004244599.1
XP_010324621.1
XP_004228864.2
XP_004234350.1
XP_019068359.1
Oryza japonica
sp|A2YXQ7.2
sp|Q6J1A5.1|
sp|Q3T5N4.1
sp|Q0J090.1|
sp|Q0J3Y6.1|

**Figure S3 Structural comparison of DREB1 proteins from drought tolerant and sensitive species.** Modeled full-length structure of dehydration-responsive element binding protein 1 (DREB1) from drought tolerant (top panel) and drought sensitive (bottom panel) species using AlphaFold2. Same representation schemes as that of Figure 1 are used. For each species, the left pane shows only the DNA-binding domain (in black cartoon) superposed on 1GCC.pdb (grey cartoon) whereas the right panel shows the electrostatic surface of the full-length protein in which blue and red corresponds to the positive and negatively charged surfaces, respectively. DNA is shown in orange cartoon. (B) Sequence alignments of DREB1 proteins from four drought-sensitive and tolerant species are shown.

Figure S4

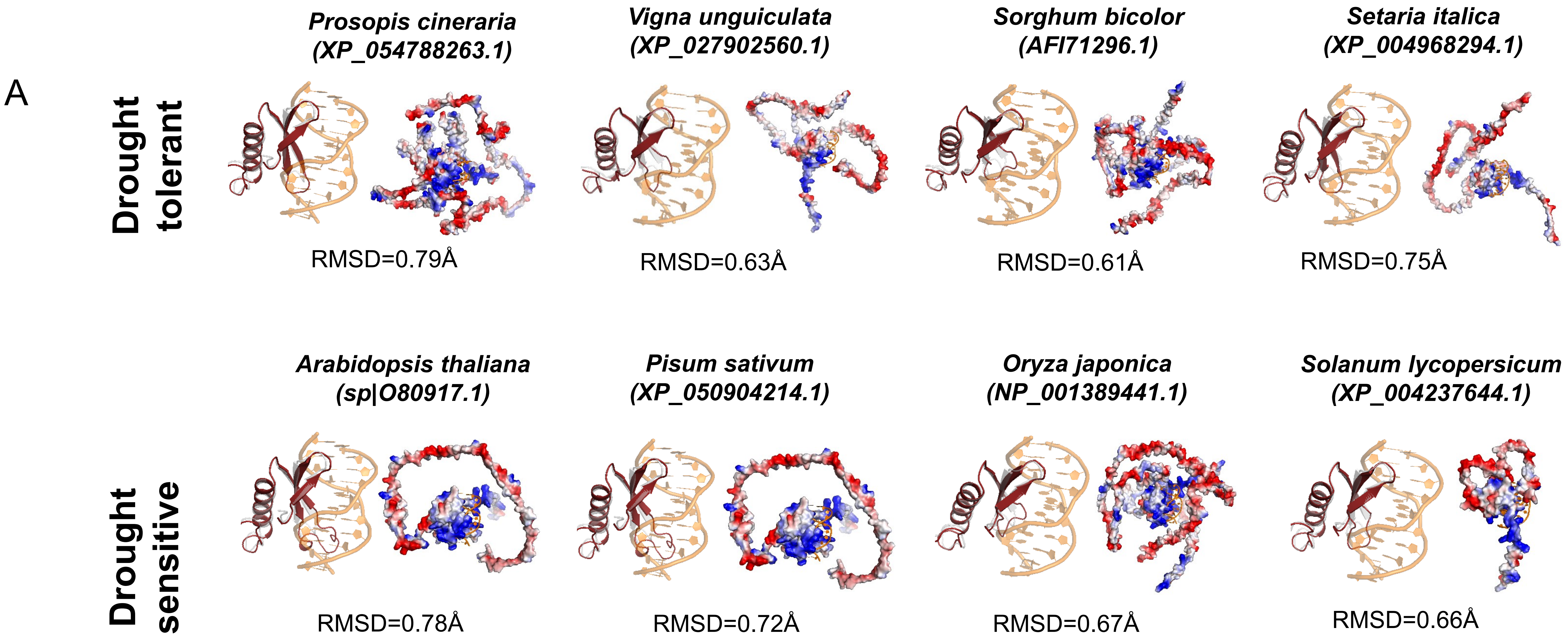


Figure S3 Contd..

Figure S4

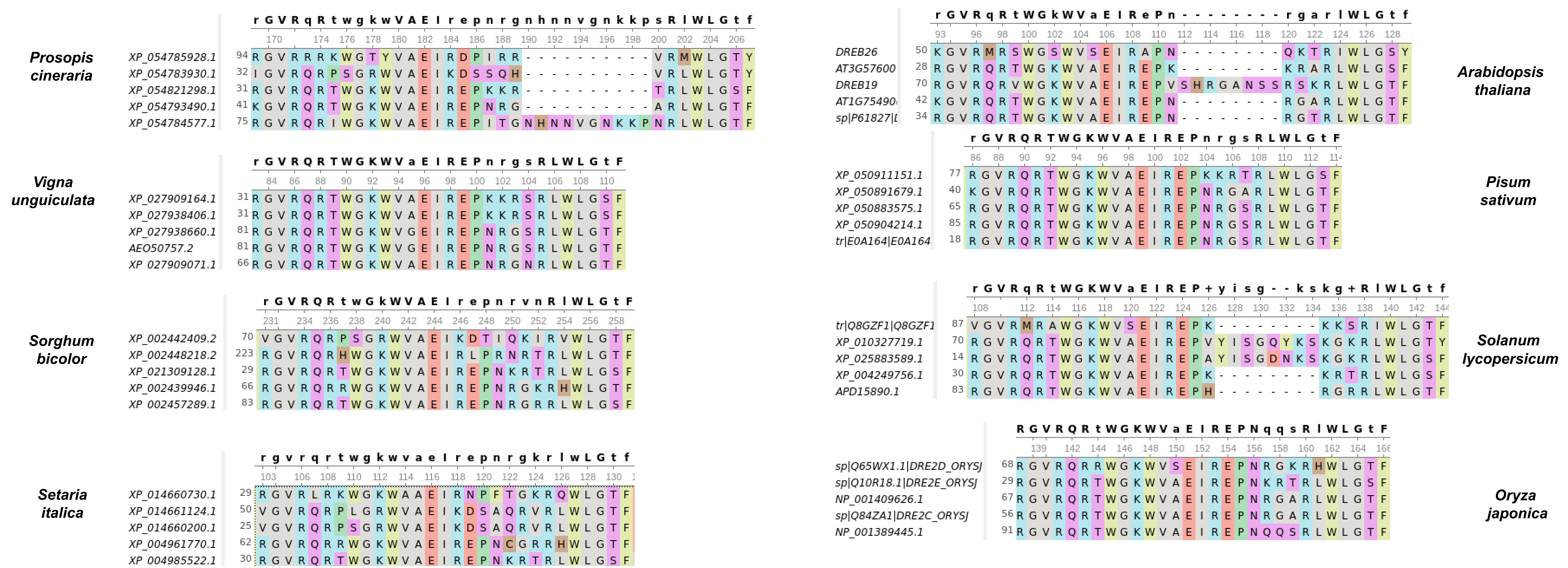
B
Drought tolerant
Drought sensitive
Prosopis cineraria
XP_054785928.1
XP_054783930.1
XP_054821298.1
XP_054793490.1
XP_054784577.1
Vigna unguiculata
XP_027909164.1
XP_027938406.1
XP_027938660.1
AEO50757.2
XP_027909071.1
Sorghum bicolor
XP_002442409.2
XP_002448218.2
XP_021309128.1
XP_002439946.1
XP_002457289.1
Setaria italica
XP_014660730.1
XP_014661124.1
XP_014660200.1
XP_004961770.1
XP_004985522.1
Arabidopsis thaliana
DREB26
AT3G57600
DREB19
AT1G75490
Pisum sativum
XP_050911151.1
XP_050891679.1
XP_050883575.1
XP_050904214.1
tr|E0A164|E0A164
Solanum lycopersicum
tr|Q8GZF1|Q8GZF1
XP_010327719.1
XP_025883589.1
XP_004249756.1
APD15890.1
Oryza japonica
sp|Q65WX1.1|DRE2D_ORYSJ
sp|Q10R18.1|DRE2E_ORYSJ
NP_001409626.1
sp|Q84ZA1|DRE2C_ORYSJ
NP_001389445.1

**Figure S4 Structural comparison of DREB2 proteins from drought tolerant and sensitive species.** (A) Modeled full-length structure of dehydration-responsive element binding protein 2 (DREB2) from drought tolerant (top panel) and drought sensitive (bottom panel) species using AlphaFold2. Same representation schemes as that of Figure 1 are used. For each species, the left pane shows only the DNA-binding domain (in black cartoon) superposed on 1GCC.pdb (grey cartoon) whereas the right panel shows the electrostatic surface of the full-length protein in which blue and red corresponds to the positive and negatively charged surfaces, respectively. DNA is shown in orange cartoon. (B) Sequence alignments of DREB2 proteins from four drought-sensitive and tolerant species are shown.

**Table S1. Genes differentially expressed in *Prosopis cineraria* under drought stress**

| Gene Symbol | Gene ID | Gene Description | $log_2$(Fold Change) | FDR adjusted *P*-value |
|---|---|---|---|---|
| LOC129319211 | 129319211 | uncharacterized protein | 14.4 | 2.79E-24 |
| LOC129307406 | 129307406 | 18 kDa seed maturation protein | 12.2 | 3.04E-29 |
| LOC129287088 | 129287088 | late embryogenesis abundant protein ECP63-like | 11.3 | 3.68E-52 |
| LOC129307452 | 129307452 | embryonic protein DC-8-like | 11.0 | 3.15E-32 |
| LOC129318966 | 129318966 | CSC1-like protein At4g02900 | 10.9 | 2.37E-63 |
| LOC129312011 | 129312011 | dehydration-responsive element-binding protein 1A-like | 10.7 | 5.97E-13 |
| LOC129304596 | 129304596 | protein MOTHER of FT and TFL1-like | 10.4 | 2.99E-13 |
| LOC129298163 | 129298163 | late embryogenesis abundant protein 2-like | 10.2 | 1.86E-37 |
| LOC129287708 | 129287708 | late embryogenesis abundant protein 2-like | 10.0 | 4.77E-36 |
| LOC129296500 | 129296500 | U2 spliceosomal RNA | 10.0 | 7.07E-09 |
| LOC129285393 | 129285393 | lipoxygenase 3, chloroplastic-like | 9.9 | 2.86E-04 |
| LOC129285284 | 129285284 | protein DOWNY MILDEW RESISTANCE 6-like | 9.8 | 4.36E-10 |
| LOC129297881 | 129297881 | protein MOTHER of FT and TFL1-like | 9.8 | 4.23E-11 |
| LOC129321003 | 129321003 | protein SIEVE ELEMENT OCCLUSION B-like | 9.8 | 9.15E-04 |
| LOC129288655 | 129288655 | probable galactinol--sucrose galactosyltransferase 5 | 9.7 | 1.13E-48 |
| LOC129312494 | 129312494 | ethylene-responsive transcription factor ERF053-like | 9.6 | 1.35E-10 |
| LOC129318452 | 129318452 | 1-aminocyclopropane-1-carboxylate oxidase-like | 9.6 | 2.11E-08 |
| LOC129318180 | 129318180 | neutral ceramidase 2-like | 9.6 | 2.10E-07 |
| LOC129307691 | 129307691 | putative calcium-binding protein CML19 | 9.6 | 1.28E-39 |
| LOC129294768 | 129294768 | uncharacterized acetyltransferase At3g50280-like | 9.5 | 8.55E-10 |
| LOC129315378 | 129315378 | uncharacterized LOC129315378 | 9.4 | 2.46E-08 |
| LOC129301921 | 129301921 | probable trehalose-phosphate phosphatase C | 9.4 | 3.40E-11 |
| LOC129294798 | 129294798 | uncharacterized LOC129294798 | 9.3 | 5.58E-10 |
| LOC129309494 | 129309494 | U-box domain-containing protein 19-like | 9.2 | 2.44E-42 |
| LOC129285444 | 129285444 | lipoxygenase 3, chloroplastic-like | 9.0 | 2.36E-07 |
| LOC129316046 | 129316046 | uncharacterized LOC129316046 | 9.0 | 5.41E-08 |
| LOC129306021 | 129306021 | short-chain dehydrogenase reductase 3b-like | 9.0 | 3.32E-13 |
| LOC129284532 | 129284532 | uncharacterized LOC129284532 | 8.9 | 1.26E-31 |
| LOC129287530 | 129287530 | histidine-containing phosphotransfer protein 4-like | 8.9 | 2.16E-07 |
| LOC129293970 | 129293970 | ethylene-responsive transcription factor ERF109 | 8.9 | 4.02E-09 |
| LOC129311887 | 129311887 | protein phosphatase 2C 51-like | 8.8 | 1.90E-37 |
| LOC129301375 | 129301375 | benzyl alcohol O-benzoyltransferase-like | 8.7 | 1.32E-06 |
| LOC129292679 | 129292679 | ethylene-responsive transcription factor ABR1-like | 8.7 | 1.03E-36 |
| LOC129321115 | 129321115 | U2 spliceosomal RNA | 8.6 | 1.34E-06 |
| LOC129318578 | 129318578 | translocator protein homolog | 8.6 | 4.25E-38 |
| LOC129291635 | 129291635 | uncharacterized LOC129291635 | 8.5 | 4.48E-15 |
| LOC129293754 | 129293754 | gibberellin-regulated protein 14 | 8.5 | 1.45E-05 |
| LOC129317042 | 129317042 | senescence associated gene 20-like | 8.5 | 1.61E-05 |
| LOC129301426 | 129301426 | cysteine proteinase inhibitor B-like | 8.4 | 1.04E-38 |
| LOC129321116 | 129321116 | U2 spliceosomal RNA | 8.4 | 7.31E-05 |
| LOC129304544 | 129304544 | F-box protein At1g61340-like | 8.4 | 1.21E-07 |
| LOC129284582 | 129284582 | PAMP-induced secreted peptide 2-like | 8.4 | 1.09E-06 |
| LOC129321157 | 129321157 | U2 spliceosomal RNA | 8.4 | 1.20E-09 |
| LOC129296507 | 129296507 | U2 spliceosomal RNA | 8.3 | 5.58E-09 |

| | | | | |
|---|---|---|---|---|
| LOC129314943 | 129314943 | 9-cis-epoxycarotenoid dioxygenase NCED1, chloroplastic-like | 8.3 | 1.69E-71 |
| LOC129314304 | 129314304 | cytochrome P450 94C1-like | 8.3 | 1.39E-08 |
| LOC129319293 | 129319293 | methyl-CpG-binding domain-containing protein 11-like | 8.2 | 6.13E-05 |
| LOC129318948 | 129318948 | uncharacterized LOC129318948 | 8.2 | 2.27E-05 |
| LOC129306125 | 129306125 | benzyl alcohol O-benzoyltransferase-like | 8.2 | 2.59E-13 |
| LOC129317041 | 129317041 | probable glutathione S-transferase | 8.1 | 1.50E-07 |
| LOC129307443 | 129307443 | transcription factor UPBEAT1 | 8.1 | 8.28E-16 |
| LOC129302142 | 129302142 | uncharacterized LOC129302142 | 8.1 | 5.57E-06 |
| LOC129305469 | 129305469 | uncharacterized LOC129305469 | 8.1 | 1.67E-05 |
| LOC129314460 | 129314460 | ethylene-responsive transcription factor ERF021-like | 8.0 | 1.42E-12 |
| LOC129294619 | 129294619 | transcription factor bHLH93-like | 8.0 | 3.46E-05 |
| LOC129298132 | 129298132 | U2 spliceosomal RNA | 8.0 | 1.42E-04 |
| LOC129297639 | 129297639 | late embryogenesis abundant protein D-34-like | 7.9 | 2.17E-05 |
| LOC129311054 | 129311054 | ATP-dependent zinc metalloprotease FTSH 6, chloroplastic | 7.9 | 8.46E-18 |
| LOC129296377 | 129296377 | serine/threonine-protein kinase SAPK3-like | 7.9 | 8.30E-21 |
| LOC129310886 | 129310886 | long-chain-alcohol oxidase FAO4A-like | 7.9 | 1.18E-13 |
| LOC129293381 | 129293381 | serine decarboxylase 1-like | 7.9 | 6.06E-06 |
| LOC129312172 | 129312172 | homeobox-leucine zipper protein ATHB-40-like | 7.9 | 3.23E-29 |
| LOC129286074 | 129286074 | uncharacterized LOC129286074 | 7.9 | 1.53E-06 |
| LOC129310160 | 129310160 | transcription factor MUTE | 7.8 | 4.76E-05 |
| LOC129297119 | 129297119 | cysteine proteinase inhibitor B-like | 7.8 | 1.81E-33 |
| LOC129301424 | 129301424 | cation/H(+) antiporter 28-like | 7.8 | 1.86E-06 |
| LOC129305247 | 129305247 | galactinol synthase 2 | 7.8 | 1.11E-30 |
| LOC129306743 | 129306743 | uncharacterized LOC129306743 | 7.7 | 5.43E-06 |
| LOC129285450 | 129285450 | lipoxygenase 3, chloroplastic-like | 7.7 | 6.98E-11 |
| LOC129293478 | 129293478 | uncharacterized LOC129293478 | 7.7 | 4.93E-06 |
| LOC129320519 | 129320519 | uncharacterized LOC129320519 | 7.7 | 8.07E-59 |
| LOC129289365 | 129289365 | G-type lectin S-receptor-like serine/threonine-protein kinase At5 | 7.7 | 4.99E-39 |
| LOC129294533 | 129294533 | uncharacterized LOC129294533 | 7.6 | 1.26E-05 |
| LOC129287048 | 129287048 | probable BOI-related E3 ubiquitin-protein ligase 3 | 7.6 | 3.86E-12 |
| LOC129321005 | 129321005 | uncharacterized LOC129321005 | 7.6 | 6.33E-05 |
| LOC129315166 | 129315166 | NDR1/HIN1-like protein 6 | 7.6 | 6.00E-06 |
| LOC129304422 | 129304422 | uncharacterized LOC129304422 | 7.6 | 1.49E-07 |
| LOC129315053 | 129315053 | protein ADP-ribosyltransferase PARP3 | 7.6 | 3.66E-66 |
| LOC129286895 | 129286895 | late embryogenesis abundant protein | 7.6 | 9.59E-08 |
| LOC129292814 | 129292814 | non-specific lipid-transfer protein 8-like | 7.5 | 2.43E-05 |
| LOC129287725 | 129287725 | 17.4 kDa class III heat shock protein | 7.5 | 1.38E-18 |
| LOC129303754 | 129303754 | aminotransferase ALD1, chloroplastic-like | 7.4 | 1.65E-106 |
| LOC129311289 | 129311289 | probable carboxylesterase 15 | 7.4 | 1.91E-05 |
| LOC129303697 | 129303697 | dehydration-responsive element-binding protein 1D-like | 7.4 | 1.47E-06 |
| LOC129311940 | 129311940 | zinc finger AN1 domain-containing stress-associated protein 12- | 7.4 | 4.58E-15 |
| LOC129320460 | 129320460 | cytochrome P450 736A117-like | 7.4 | 1.30E-08 |
| LOC129305657 | 129305657 | protein RTF1 homolog | 7.4 | 1.42E-08 |
| LOC129300707 | 129300707 | uncharacterized LOC129300707 | 7.3 | 5.66E-05 |
| LOC129317624 | 129317624 | transcription factor HEC2-like | 7.3 | 7.86E-05 |
| LOC129311911 | 129311911 | dehydration-responsive element-binding protein 1E-like | 7.3 | 8.07E-13 |
| LOC129297867 | 129297867 | cation/H(+) antiporter 28-like | 7.3 | 5.31E-05 |
| LOC129292480 | 129292480 | serine/threonine-protein kinase BIK1-like | 7.2 | 3.56E-04 |

| | | | | |
|---|---|---|---|---|
| LOC129314340 | 129314340 | uncharacterized LOC129314340 | 7.2 | 4.34E-06 |
| LOC129300041 | 129300041 | LEAF RUST 10 DISEASE-RESISTANCE LOCUS RECEPTO | 7.2 | 1.14E-04 |
| LOC129305159 | 129305159 | protein TIFY 5A | 7.2 | 1.08E-07 |
| LOC129286068 | 129286068 | uncharacterized LOC129286068 | 7.1 | 6.58E-05 |
| LOC129316267 | 129316267 | uncharacterized LOC129316267 | 7.1 | 4.38E-05 |
| LOC129305301 | 129305301 | subtilisin-like protease SBT1.2 | 7.1 | 6.92E-08 |
| LOC129290919 | 129290919 | hydroquinone glucosyltransferase-like | 7.1 | 1.05E-03 |
| LOC129319656 | 129319656 | late embryogenesis abundant protein D-34-like | 7.1 | 2.13E-03 |
| LOC129318552 | 129318552 | NAD(P)H-dependent 6'-deoxychalcone synthase-like | 7.1 | 1.27E-04 |
| LOC129320087 | 129320087 | protein MOTHER of FT and TFL1 | 7.1 | 1.75E-34 |
| LOC129297529 | 129297529 | 17.3 kDa class I heat shock protein-like | 7.1 | 1.14E-04 |
| LOC129318973 | 129318973 | EID1-like F-box protein 3 | 7.1 | 1.11E-62 |
| LOC129307322 | 129307322 | auxin-responsive protein SAUR21-like | 7.0 | 1.59E-04 |
| LOC129309902 | 129309902 | transcription factor bHLH92 | 7.0 | 1.45E-05 |
| LOC129299714 | 129299714 | UPF0496 protein At1g20180-like | 7.0 | 9.44E-05 |
| LOC129307867 | 129307867 | ethylene-responsive transcription factor ERF096-like | 7.0 | 6.41E-04 |
| LOC129305096 | 129305096 | polygalacturonase inhibitor-like | 7.0 | 9.63E-03 |
| LOC129319223 | 129319223 | axial regulator YABBY 5-like | 7.0 | 4.38E-04 |
| LOC129295966 | 129295966 | glycine-rich protein 23-like | 7.0 | 5.93E-09 |
| LOC129322101 | 129322101 | scopoletin glucosyltransferase-like | 7.0 | 1.17E-20 |
| LOC129316816 | 129316816 | NAC transcription factor 29-like | 7.0 | 8.47E-04 |
| LOC129311949 | 129311949 | UDP-glycosyltransferase 83A1-like | 7.0 | 5.18E-04 |
| LOC129293403 | 129293403 | CEN-like protein 1 | 6.9 | 4.72E-12 |
| LOC129307395 | 129307395 | precursor of CEP14-like | 6.9 | 5.53E-07 |
| LOC129299448 | 129299448 | 5.8S ribosomal RNA | 6.9 | 1.37E-02 |
| LOC129315367 | 129315367 | uncharacterized LOC129315367 | 6.9 | 1.28E-03 |
| LOC129291359 | 129291359 | 1-aminocyclopropane-1-carboxylate synthase-like | 6.9 | 2.15E-04 |
| LOC129319775 | 129319775 | probable E3 ubiquitin-protein ligase XERICO | 6.9 | 7.25E-06 |
| LOC129285803 | 129285803 | stachyose synthase-like | 6.9 | 5.77E-14 |
| LOC129290810 | 129290810 | uncharacterized LOC129290810 | 6.9 | 3.71E-04 |
| LOC129292011 | 129292011 | Bowman-Birk type proteinase inhibitor-like | 6.9 | 4.52E-03 |
| LOC129288324 | 129288324 | E3 ubiquitin-protein ligase CHIP-like | 6.9 | 5.39E-04 |
| LOC129291752 | 129291752 | 26.5 kDa heat shock protein, mitochondrial-like | 6.8 | 2.70E-04 |
| LOC129319998 | 129319998 | putative calcium-transporting ATPase 13, plasma membrane-ty | 6.8 | 1.13E-10 |
| LOC129290187 | 129290187 | uncharacterized LOC129290187 | 6.8 | 1.14E-04 |
| LOC129313228 | 129313228 | B-box domain protein 31-like | 6.8 | 4.40E-04 |
| LOC129289517 | 129289517 | jasmonate ZIM domain-containing protein 1-like | 6.8 | 2.80E-10 |
| LOC129295242 | 129295242 | polygalacturonase-like | 6.8 | 4.48E-04 |
| LOC129313527 | 129313527 | NAC domain-containing protein 72-like | 6.8 | 7.27E-22 |
| LOC129307807 | 129307807 | probable N-acetyltransferase HLS1 | 6.8 | 1.60E-02 |
| LOC129303947 | 129303947 | protodermal factor 1 | 6.8 | 5.94E-04 |
| LOC129297118 | 129297118 | cation/H(+) antiporter 28-like | 6.8 | 5.68E-04 |
| LOC129284477 | 129284477 | scarecrow-like protein 32 | 6.8 | 1.69E-02 |
| LOC129316211 | 129316211 | aquaporin TIP1-2-like | 6.8 | 1.24E-17 |
| LOC129295510 | 129295510 | abscisic acid 8'-hydroxylase CYP707A2-like | 6.8 | 1.04E-06 |
| LOC129284608 | 129284608 | uncharacterized LOC129284608 | 6.7 | 9.19E-12 |
| LOC129320117 | 129320117 | dehydration-responsive element-binding protein 2F | 6.7 | 1.06E-03 |
| LOC129308555 | 129308555 | abscisic stress-ripening protein 2-like | 6.7 | 6.73E-04 |

| | | | | |
|---|---|---|---|---|
| LOC129309841 | 129309841 | ethylene-responsive transcription factor 1B | 6.7 | 9.34E-13 |
| LOC129301731 | 129301731 | ethylene-responsive transcription factor ERF017-like | 6.7 | 8.87E-05 |
| LOC129307454 | 129307454 | uncharacterized LOC129307454 | 6.7 | 1.46E-09 |
| LOC129315348 | 129315348 | exocyst complex component EXO70H1-like | 6.7 | 3.62E-16 |
| LOC129293982 | 129293982 | serine carboxypeptidase-like 9 | 6.7 | 2.94E-02 |
| LOC129294442 | 129294442 | uncharacterized LOC129294442 | 6.7 | 1.42E-03 |
| LOC129289691 | 129289691 | ankyrin repeat-containing protein At5g02620-like | 6.7 | 2.30E-02 |
| LOC129298432 | 129298432 | uncharacterized LOC129298432 | 6.7 | 7.14E-04 |
| LOC129307860 | 129307860 | ethylene-responsive transcription factor 14-like | 6.7 | 4.56E-04 |
| LOC129284534 | 129284534 | probable indole-3-acetic acid-amido synthetase GH3.1 | 6.7 | 4.21E-03 |
| LOC129309993 | 129309993 | HVA22-like protein e | 6.6 | 2.45E-26 |
| LOC129306001 | 129306001 | uncharacterized LOC129306001 | 6.6 | 3.96E-08 |
| LOC129292389 | 129292389 | uncharacterized LOC129292389 | 6.6 | 1.97E-03 |
| LOC129308943 | 129308943 | probable aquaporin TIP-type alpha | 6.6 | 3.51E-04 |
| LOC129293660 | 129293660 | abscisic acid 8'-hydroxylase 4 | 6.6 | 8.30E-15 |
| LOC129302602 | 129302602 | ATP-dependent Clp protease ATP-binding subunit ClpA homol | 6.6 | 2.91E-04 |
| LOC129306019 | 129306019 | uncharacterized LOC129306019 | 6.6 | 2.31E-03 |
| LOC129314917 | 129314917 | calcium-binding protein KRP1-like | 6.6 | 1.13E-03 |
| LOC129319320 | 129319320 | cysteine proteinase inhibitor B-like | 6.6 | 1.28E-03 |
| LOC129322388 | 129322388 | 14-3-3-like protein GF14 iota | 6.6 | 7.71E-04 |
| LOC129309878 | 129309878 | 3-ketoacyl-CoA synthase 10-like | 6.6 | 4.66E-03 |
| LOC129304226 | 129304226 | protein S40-7-like | 6.6 | 4.28E-04 |
| LOC129316270 | 129316270 | uncharacterized LOC129316270 | 6.6 | 1.12E-03 |
| LOC129305052 | 129305052 | uncharacterized LOC129305052 | 6.6 | 2.52E-03 |
| LOC129297831 | 129297831 | 28S ribosomal RNA | 6.5 | 8.79E-03 |
| LOC129284556 | 129284556 | dehydration-responsive element-binding protein 1E | 6.5 | 1.34E-07 |
| LOC129299228 | 129299228 | probable glutathione S-transferase | 6.5 | 5.52E-03 |
| LOC129310497 | 129310497 | uncharacterized LOC129310497 | 6.5 | 2.92E-03 |
| LOC129314968 | 129314968 | expansin-A12 | 6.5 | 1.41E-03 |
| LOC129316042 | 129316042 | transcription factor MYB41-like | 6.5 | 1.68E-04 |
| LOC129292617 | 129292617 | dehydration-responsive element-binding protein 1A-like | 6.5 | 4.84E-07 |
| LOC129288731 | 129288731 | laccase-7-like | 6.5 | 2.84E-03 |
| LOC129307056 | 129307056 | desiccation protectant protein Lea14 homolog | 6.5 | 6.99E-17 |
| LOC129307635 | 129307635 | protein UNIFOLIATA | 6.5 | 1.05E-04 |
| LOC129294430 | 129294430 | abietadienol/abietadienal oxidase | 6.5 | 2.49E-03 |
| LOC129312905 | 129312905 | uncharacterized LOC129312905 | 6.5 | 1.52E-03 |
| LOC129307834 | 129307834 | B3 domain-containing protein At2g33720-like | 6.5 | 1.53E-04 |
| LOC129287808 | 129287808 | uncharacterized LOC129287808 | 6.5 | 7.04E-03 |
| LOC129307412 | 129307412 | late embryogenesis abundant protein 31-like | 6.5 | 6.95E-03 |
| LOC129304024 | 129304024 | protein phosphatase 2C 51-like | 6.5 | 2.67E-49 |
| LOC129291619 | 129291619 | transcription factor MYB78-like | 6.4 | 1.85E-07 |
| LOC129299935 | 129299935 | uncharacterized LOC129299935 | 6.4 | 4.20E-04 |
| LOC129306140 | 129306140 | probable galacturonosyltransferase-like 10 | 6.4 | 2.52E-03 |
| LOC129296521 | 129296521 | U2 spliceosomal RNA | 6.4 | 7.51E-05 |
| LOC129306880 | 129306880 | dormancy-associated protein 1 | 6.4 | 8.38E-20 |
| LOC129291736 | 129291736 | LIM domain-containing protein PLIM2c-like | 6.4 | 2.40E-03 |
| LOC129315267 | 129315267 | ethylene-responsive transcription factor ERF053-like | 6.4 | 1.52E-66 |
| LOC129300232 | 129300232 | polygalacturonase inhibitor-like | 6.4 | 3.86E-02 |

| | | | | |
|---|---|---|---|---|
| LOC129299616 | 129299616 | cyclin-dependent protein kinase inhibitor SMR6-like | 6.4 | 1.38E-04 |
| LOC129308331 | 129308331 | uncharacterized LOC129308331 | 6.4 | 3.52E-02 |
| LOC129294188 | 129294188 | probable carboxylesterase 17 | 6.4 | 2.32E-03 |
| LOC129294786 | 129294786 | fasciclin-like arabinogalactan protein 3 | 6.4 | 2.59E-03 |
| LOC129317857 | 129317857 | uncharacterized LOC129317857 | 6.4 | 1.26E-02 |
| LOC129321145 | 129321145 | U2 spliceosomal RNA | 6.4 | 4.42E-03 |
| LOC129285371 | 129285371 | ethylene-responsive transcription factor ERF109-like | 6.4 | 6.48E-03 |
| LOC129296338 | 129296338 | 40S ribosomal protein S17-like | 6.4 | 3.63E-04 |
| LOC129295419 | 129295419 | GABA transporter 1-like | 6.4 | 1.34E-02 |
| LOC129306858 | 129306858 | WAT1-related protein At1g25270-like | 6.3 | 2.46E-10 |
| LOC129300203 | 129300203 | sugar transport protein 13-like | 6.3 | 4.96E-14 |
| LOC129309165 | 129309165 | uncharacterized LOC129309165 | 6.3 | 3.12E-03 |
| LOC129291669 | 129291669 | universal stress protein PHOS32-like | 6.3 | 1.33E-03 |
| LOC129286907 | 129286907 | protein DMR6-LIKE OXYGENASE 2-like | 6.3 | 9.15E-03 |
| LOC129291821 | 129291821 | uncharacterized LOC129291821 | 6.3 | 7.92E-16 |
| LOC129301915 | 129301915 | vicilin Cor a 11.0101-like | 6.3 | 5.06E-03 |
| LOC129300710 | 129300710 | F-box/kelch-repeat protein At3g23880-like | 6.3 | 1.54E-02 |
| LOC129294239 | 129294239 | ferritin-3, chloroplastic-like | 6.3 | 4.19E-02 |
| LOC129309617 | 129309617 | class-10 pathogenesis-related protein 1-like | 6.3 | 1.08E-02 |
| LOC129296515 | 129296515 | U2 spliceosomal RNA | 6.3 | 4.36E-07 |
| LOC129312632 | 129312632 | uncharacterized LOC129312632 | 6.3 | 1.57E-05 |
| LOC129304238 | 129304238 | high mobility group B protein 7-like | 6.3 | 3.93E-04 |
| LOC129288203 | 129288203 | flavanone 3-dioxygenase 3-like | 6.2 | 1.47E-09 |
| LOC129294894 | 129294894 | photosystem II D2 protein-like | 6.2 | 1.43E-02 |
| LOC129294497 | 129294497 | E3 ubiquitin-protein ligase SINAT5-like | 6.2 | 8.88E-03 |
| LOC129305061 | 129305061 | endonuclease 1-like | 6.2 | 5.10E-03 |
| LOC129300007 | 129300007 | uncharacterized LOC129300007 | 6.2 | 1.09E-03 |
| LOC129304967 | 129304967 | polyphenol oxidase, chloroplastic-like | 6.2 | 5.52E-03 |
| LOC129289624 | 129289624 | uncharacterized LOC129289624 | 6.2 | 9.60E-03 |
| LOC129287918 | 129287918 | homeobox-leucine zipper protein ATHB-12-like | 6.2 | 5.24E-57 |
| LOC129316188 | 129316188 | cytochrome P450 71A9-like | 6.2 | 8.57E-14 |
| LOC129307696 | 129307696 | uncharacterized LOC129307696 | 6.2 | 8.29E-03 |
| LOC129304582 | 129304582 | calcium-binding protein PBP1-like | 6.2 | 2.45E-03 |
| LOC129302054 | 129302054 | probable membrane-associated kinase regulator 4 | 6.1 | 1.22E-02 |
| LOC129298726 | 129298726 | uncharacterized LOC129298726 | 6.1 | 8.50E-03 |
| LOC129323052 | 129323052 | putative RING-H2 finger protein ATL21A | 6.1 | 1.20E-02 |
| LOC129321383 | 129321383 | PI-PLC X domain-containing protein At5g67130-like | 6.1 | 8.26E-11 |
| LOC129290659 | 129290659 | homeobox-leucine zipper protein ATHB-12-like | 6.1 | 8.62E-03 |
| LOC129310852 | 129310852 | protein DMP8-like | 6.1 | 1.79E-03 |
| LOC129292379 | 129292379 | uncharacterized LOC129292379 | 6.1 | 9.43E-04 |
| LOC129307387 | 129307387 | probable glycerol-3-phosphate acyltransferase 3 | 6.1 | 1.56E-03 |
| LOC129322757 | 129322757 | probable mannitol dehydrogenase | 6.1 | 1.37E-03 |
| LOC129303029 | 129303029 | uncharacterized LOC129303029 | 6.1 | 2.48E-04 |
| LOC129320066 | 129320066 | protein MANNAN SYNTHESIS-RELATED-like | 6.0 | 1.26E-06 |
| LOC129319236 | 129319236 | uncharacterized LOC129319236 | 6.0 | 8.82E-09 |
| LOC129297953 | 129297953 | uncharacterized LOC129297953 | 6.0 | 3.03E-02 |
| LOC129319109 | 129319109 | U2 spliceosomal RNA | 6.0 | 2.94E-02 |
| LOC129299725 | 129299725 | uncharacterized LOC129299725 | 6.0 | 9.52E-03 |

| | | | | |
|---|---|---|---|---|
| LOC129322643 | 129322643 | uncharacterized LOC129322643 | 6.0 | 3.02E-18 |
| LOC129299002 | 129299002 | elongation of fatty acids protein 3-like | 6.0 | 3.41E-02 |
| LOC129305707 | 129305707 | inorganic phosphate transporter 1-4-like | 6.0 | 1.72E-04 |
| LOC129303992 | 129303992 | subtilisin-like protease SBT4.3 | 6.0 | 1.24E-02 |
| LOC129319232 | 129319232 | FCS-Like Zinc finger 15-like | 6.0 | 2.29E-03 |
| LOC129306855 | 129306855 | probable WRKY transcription factor 75 | 6.0 | 1.13E-02 |
| LOC129313042 | 129313042 | putative receptor-like protein kinase At3g47110 | 5.9 | 1.23E-17 |
| LOC129301613 | 129301613 | ethylene-responsive transcription factor ERF113-like | 5.9 | 5.13E-03 |
| LOC129320261 | 129320261 | uncharacterized LOC129320261 | 5.9 | 1.47E-11 |
| LOC129322132 | 129322132 | endoglucanase-like | 5.9 | 2.84E-02 |
| LOC129284410 | 129284410 | protein LURP-one-related 17 | 5.9 | 2.18E-05 |
| LOC129308891 | 129308891 | ethylene-responsive transcription factor ERF095-like | 5.9 | 3.15E-02 |
| LOC129291998 | 129291998 | multiple organellar RNA editing factor 3, mitochondrial-like | 5.9 | 1.51E-02 |
| LOC129306873 | 129306873 | uncharacterized LOC129306873 | 5.9 | 2.96E-04 |
| LOC129309936 | 129309936 | beta-amylase 1, chloroplastic-like | 5.9 | 3.08E-08 |
| LOC129306310 | 129306310 | protein ABSCISIC ACID-INSENSITIVE 5-like | 5.9 | 6.31E-23 |
| LOC129315008 | 129315008 | probable protein phosphatase 2C 24 | 5.9 | 4.71E-13 |
| LOC129302223 | 129302223 | polygalacturonase-like | 5.9 | 2.03E-02 |
| LOC129302104 | 129302104 | agamous-like MADS-box protein AGL80 | 5.9 | 3.47E-02 |
| LOC129313220 | 129313220 | protein GL2-INTERACTING REPRESSOR 2 | 5.9 | 1.44E-02 |
| LOC129312756 | 129312756 | receptor-like protein EIX1 | 5.9 | 2.09E-02 |
| LOC129319930 | 129319930 | peptidyl-prolyl cis-trans isomerase CYP95-like | 5.9 | 3.03E-02 |
| LOC129318186 | 129318186 | dehydrin ERD14-like | 5.8 | 7.20E-21 |
| LOC129294722 | 129294722 | probable calcium-binding protein CML44 | 5.8 | 8.66E-16 |
| LOC129301939 | 129301939 | CBL-interacting serine/threonine-protein kinase 5-like | 5.8 | 4.64E-10 |
| LOC129301094 | 129301094 | uncharacterized LOC129301094 | 5.8 | 4.59E-04 |
| LOC129293054 | 129293054 | probable amidase At4g34880 | 5.8 | 6.57E-04 |
| LOC129311945 | 129311945 | abscisic acid 8'-hydroxylase CYP707A2-like | 5.8 | 1.14E-09 |
| LOC129310021 | 129310021 | nuclear transcription factor Y subunit B-7 | 5.8 | 2.20E-02 |
| LOC129299021 | 129299021 | uncharacterized LOC129299021 | 5.8 | 2.25E-02 |
| LOC129319269 | 129319269 | oleosin Cor a 15-like | 5.8 | 8.61E-03 |
| LOC129297723 | 129297723 | probable histone H2A.3 | 5.8 | 3.40E-02 |
| LOC129316593 | 129316593 | germin-like protein subfamily 1 member 13 | 5.8 | 2.31E-03 |
| LOC129318840 | 129318840 | probable glutathione S-transferase | 5.8 | 8.00E-07 |
| LOC129318891 | 129318891 | HMG1/2-like protein | 5.8 | 3.41E-02 |
| LOC129291100 | 129291100 | uncharacterized LOC129291100 | 5.8 | 2.10E-13 |
| LOC129322415 | 129322415 | nodulation receptor kinase-like | 5.8 | 2.52E-11 |
| LOC129314571 | 129314571 | uncharacterized LOC129314571 | 5.7 | 3.11E-18 |
| LOC129310209 | 129310209 | U1 spliceosomal RNA | 5.7 | 3.46E-02 |
| LOC129306864 | 129306864 | transcription factor DIVARICATA | 5.7 | 4.31E-28 |
| LOC129317294 | 129317294 | uncharacterized LOC129317294 | 5.7 | 2.45E-02 |
| LOC129294107 | 129294107 | trifunctional UDP-glucose 4,6-dehydratase/UDP-4-keto-6-deoxy | 5.7 | 2.19E-02 |
| LOC129303674 | 129303674 | metal transporter Nramp5-like | 5.7 | 2.04E-02 |
| LOC129293664 | 129293664 | serine carboxypeptidase-like 18 | 5.7 | 5.40E-05 |
| LOC129285504 | 129285504 | phenolic glucoside malonyltransferase 2-like | 5.7 | 2.03E-08 |
| LOC129319433 | 129319433 | uncharacterized LOC129319433 | 5.7 | 4.26E-03 |
| LOC129292124 | 129292124 | uncharacterized LOC129292124 | 5.7 | 3.53E-02 |
| LOC129310574 | 129310574 | probable phospholipid hydroperoxide glutathione peroxidase | 5.7 | 4.78E-09 |

| | | | | |
|---|---|---|---|---|
| LOC129291145 | 129291145 | cytochrome P450 86A8 | 5.7 | 2.77E-16 |
| LOC129300759 | 129300759 | nudix hydrolase 16, mitochondrial-like | 5.7 | 1.41E-15 |
| LOC129320263 | 129320263 | heat shock 70 kDa protein | 5.7 | 7.14E-26 |
| LOC129296488 | 129296488 | U1 spliceosomal RNA | 5.7 | 1.30E-02 |
| LOC129321516 | 129321516 | reticulon-like protein B13 | 5.7 | 2.99E-02 |
| LOC129304605 | 129304605 | UDP-glycosyltransferase 90A1-like | 5.7 | 2.49E-02 |
| LOC129321438 | 129321438 | ninja-family protein AFP1-like | 5.7 | 1.92E-28 |
| LOC129307119 | 129307119 | uncharacterized LOC129307119 | 5.7 | 6.18E-03 |
| LOC129293497 | 129293497 | protein JINGUBANG-like | 5.7 | 3.70E-12 |
| LOC129290574 | 129290574 | protein MAIN-LIKE 1-like | 5.7 | 1.04E-03 |
| LOC129309871 | 129309871 | uncharacterized LOC129309871 | 5.6 | 1.15E-02 |
| LOC129291375 | 129291375 | uncharacterized LOC129291375 | 5.6 | 1.21E-18 |
| LOC129322408 | 129322408 | F-box/kelch-repeat protein At1g23390-like | 5.6 | 1.02E-05 |
| LOC129310740 | 129310740 | putative expansin-A30 | 5.6 | 2.99E-02 |
| LOC129312158 | 129312158 | uncharacterized LOC129312158 | 5.6 | 4.26E-27 |
| LOC129319815 | 129319815 | late embryogenesis abundant protein D-29 | 5.6 | 7.54E-43 |
| LOC129321243 | 129321243 | UDP-glucose 4-epimerase GEPI48 | 5.6 | 4.16E-37 |
| LOC129299536 | 129299536 | beta-amylase 1, chloroplastic-like | 5.6 | 9.89E-11 |
| LOC129299987 | 129299987 | seed biotin-containing protein SBP65-like | 5.6 | 4.57E-04 |
| LOC129315881 | 129315881 | inositol-3-phosphate synthase-like | 5.6 | 1.38E-18 |
| LOC129314597 | 129314597 | COBRA-like protein 10 | 5.6 | 4.02E-02 |
| LOC129293323 | 129293323 | heavy metal-associated isoprenylated plant protein 39-like | 5.6 | 4.15E-08 |
| LOC129305962 | 129305962 | cystinosin homolog | 5.6 | 2.28E-16 |
| LOC129292442 | 129292442 | receptor-like protein EIX1 | 5.6 | 4.35E-02 |
| LOC129299376 | 129299376 | UDP-glycosyltransferase 87A1-like | 5.6 | 3.57E-02 |
| LOC129307920 | 129307920 | GDSL esterase/lipase At5g08460-like | 5.6 | 3.76E-02 |
| LOC129319213 | 129319213 | transcription factor WER-like | 5.6 | 3.60E-02 |
| LOC129294469 | 129294469 | protein ABSCISIC ACID-INSENSITIVE 5 | 5.5 | 8.97E-03 |
| LOC129304646 | 129304646 | piriformospora indica-insensitive protein 2-like | 5.5 | 4.78E-02 |
| LOC129287452 | 129287452 | cyclin-dependent protein kinase inhibitor SMR4 | 5.5 | 2.12E-10 |
| LOC129298057 | 129298057 | dehydration-responsive element-binding protein 1A-like | 5.5 | 2.92E-09 |
| LOC129288229 | 129288229 | heat stress transcription factor B-2b-like | 5.5 | 5.28E-11 |
| LOC129315560 | 129315560 | uncharacterized LOC129315560 | 5.5 | 1.25E-23 |
| LOC129304528 | 129304528 | protein RGF1 INDUCIBLE TRANSCRIPTION FACTOR 1 | 5.5 | 4.90E-02 |
| LOC129310579 | 129310579 | probable phospholipid hydroperoxide glutathione peroxidase | 5.5 | 1.69E-09 |
| LOC129287001 | 129287001 | uncharacterized LOC129287001 | 5.5 | 2.28E-03 |
| LOC129307935 | 129307935 | sugar transport protein 13-like | 5.5 | 9.35E-11 |
| LOC129296170 | 129296170 | 2-oxoglutarate-dependent dioxygenase DAO-like | 5.5 | 3.64E-02 |
| LOC129290068 | 129290068 | protein indeterminate-domain 12-like | 5.5 | 5.20E-04 |
| LOC129321595 | 129321595 | probable methyltransferase PMT27 | 5.5 | 4.13E-02 |
| LOC129320699 | 129320699 | putative ETHYLENE INSENSITIVE 3-like 4 protein | 5.5 | 3.72E-02 |
| LOC129316078 | 129316078 | polygalacturonase-like | 5.5 | 1.54E-02 |
| LOC129290344 | 129290344 | phospholipase A(1) DAD1, chloroplastic-like | 5.5 | 4.61E-02 |
| LOC129298884 | 129298884 | NAC domain-containing protein 72-like | 5.5 | 5.48E-14 |
| LOC129318851 | 129318851 | myb family transcription factor PHL8-like | 5.5 | 4.03E-04 |
| LOC129290575 | 129290575 | non-specific lipid-transfer protein 1-like | 5.5 | 1.27E-12 |
| LOC129303580 | 129303580 | homeobox protein knotted-1-like 2 | 5.5 | 9.40E-03 |
| LOC129302736 | 129302736 | protein EXORDIUM-like 4 | 5.5 | 9.56E-04 |

| | | | | |
|---|---|---|---|---|
| LOC129293640 | 129293640 | beta-glucosidase 46-like | 5.5 | 8.43E-04 |
| LOC129321021 | 129321021 | uncharacterized LOC129321021 | 5.5 | 7.98E-03 |
| LOC129318834 | 129318834 | uncharacterized LOC129318834 | 5.4 | 1.15E-03 |
| LOC129287764 | 129287764 | SNF1-related protein kinase regulatory subunit gamma-like PV4 | 5.4 | 4.71E-02 |
| LOC129300006 | 129300006 | uncharacterized LOC129300006 | 5.4 | 1.70E-06 |
| LOC129301839 | 129301839 | uncharacterized LOC129301839 | 5.4 | 3.15E-03 |
| LOC129308081 | 129308081 | codeine O-demethylase-like | 5.4 | 7.37E-04 |
| LOC129319200 | 129319200 | uncharacterized LOC129319200 | 5.4 | 1.43E-04 |
| LOC129305692 | 129305692 | subtilisin-like protease SBT4.15 | 5.4 | 4.68E-09 |
| LOC129287584 | 129287584 | NAC domain-containing protein 83-like | 5.4 | 1.19E-04 |
| LOC129318589 | 129318589 | uncharacterized LOC129318589 | 5.4 | 1.96E-04 |
| LOC129318101 | 129318101 | 17.3 kDa class II heat shock protein-like | 5.4 | 6.18E-08 |
| LOC129316290 | 129316290 | NAC domain-containing protein 72-like | 5.4 | 1.74E-16 |
| LOC129306016 | 129306016 | uncharacterized LOC129306016 | 5.4 | 1.26E-13 |
| LOC129305658 | 129305658 | protein RTF1 homolog | 5.4 | 6.38E-05 |
| LOC129313638 | 129313638 | transcription factor MYB102-like | 5.4 | 2.17E-07 |
| LOC129295771 | 129295771 | F-box protein SKIP27-like | 5.4 | 6.46E-23 |
| LOC129313242 | 129313242 | protein REDOX 2-like | 5.4 | 1.61E-10 |
| LOC129320318 | 129320318 | U-box domain-containing protein 19-like | 5.4 | 5.11E-35 |
| LOC129312010 | 129312010 | MYB-like transcription factor EOBI | 5.4 | 8.01E-16 |
| LOC129299712 | 129299712 | expansin-A11-like | 5.4 | 4.12E-03 |
| LOC129287423 | 129287423 | ethylene-responsive transcription factor 3-like | 5.4 | 1.13E-20 |
| LOC129288546 | 129288546 | ABC transporter G family member STR-like | 5.3 | 5.24E-05 |
| LOC129286731 | 129286731 | type III polyketide synthase B | 5.3 | 1.64E-08 |
| LOC129311914 | 129311914 | uncharacterized LOC129311914 | 5.3 | 3.39E-06 |
| LOC129303271 | 129303271 | abscisic acid 8'-hydroxylase 4-like | 5.3 | 9.63E-03 |
| LOC129321631 | 129321631 | leucoanthocyanidin reductase-like | 5.3 | 5.37E-04 |
| LOC129307165 | 129307165 | F-box protein SKIP27-like | 5.3 | 5.67E-23 |
| LOC129292606 | 129292606 | low-temperature-induced 65 kDa protein-like | 5.3 | 4.60E-28 |
| LOC129296130 | 129296130 | CASP-like protein 1E2 | 5.3 | 2.26E-03 |
| LOC129310197 | 129310197 | U1 spliceosomal RNA | 5.3 | 1.54E-03 |
| LOC129299371 | 129299371 | UDP-glycosyltransferase 87A1-like | 5.3 | 4.27E-02 |
| LOC129304431 | 129304431 | uncharacterized LOC129304431 | 5.3 | 3.90E-03 |
| LOC129309749 | 129309749 | uncharacterized LOC129309749 | 5.3 | 4.87E-09 |
| LOC129303874 | 129303874 | 17.9 kDa class II heat shock protein-like | 5.3 | 2.40E-08 |
| LOC129321967 | 129321967 | uncharacterized LOC129321967 | 5.3 | 2.01E-06 |
| LOC129309118 | 129309118 | class V chitinase-like | 5.3 | 1.67E-20 |
| LOC129291828 | 129291828 | probable N-acetyltransferase HLS1 | 5.3 | 4.58E-04 |
| LOC129312865 | 129312865 | receptor-like serine/threonine-protein kinase At1g78530 | 5.3 | 3.36E-19 |
| LOC129314557 | 129314557 | uncharacterized LOC129314557 | 5.3 | 5.86E-28 |
| LOC129294691 | 129294691 | uncharacterized LOC129294691 | 5.3 | 1.77E-02 |
| LOC129303671 | 129303671 | glucan endo-1,3-beta-glucosidase 12-like | 5.3 | 1.49E-14 |
| LOC129293838 | 129293838 | arabinogalactan protein 22-like | 5.3 | 1.55E-14 |
| LOC129290140 | 129290140 | uncharacterized LOC129290140 | 5.2 | 4.38E-02 |
| LOC129301890 | 129301890 | uncharacterized LOC129301890 | 5.2 | 1.32E-02 |
| LOC129307735 | 129307735 | uncharacterized LOC129307735 | 5.2 | 3.08E-08 |
| LOC129309856 | 129309856 | dehydrin DHN1-like | 5.2 | 3.22E-14 |
| LOC129291444 | 129291444 | probable WRKY transcription factor 40 | 5.2 | 1.09E-12 |

| | | | | |
|---|---|---|---|---|
| LOC129316232 | 129316232 | squamosa promoter-binding protein 1-like | 5.2 | 1.75E-10 |
| LOC129285703 | 129285703 | ABC transporter G family member 25 | 5.2 | 6.41E-21 |
| LOC129285688 | 129285688 | uncharacterized LOC129285688 | 5.2 | 2.56E-02 |
| LOC129310311 | 129310311 | 21 kDa protein-like | 5.2 | 3.24E-02 |
| LOC129287055 | 129287055 | uncharacterized LOC129287055 | 5.2 | 6.39E-13 |
| LOC129305095 | 129305095 | nudix hydrolase 16, mitochondrial-like | 5.2 | 1.66E-22 |
| LOC129295666 | 129295666 | protodermal factor 1-like | 5.2 | 1.93E-02 |
| LOC129306429 | 129306429 | NAC transcription factor 25 | 5.2 | 3.66E-09 |
| LOC129310032 | 129310032 | uncharacterized LOC129310032 | 5.2 | 2.16E-10 |
| LOC129317662 | 129317662 | probable nucleoredoxin 2 | 5.2 | 2.89E-29 |
| LOC129323135 | 129323135 | transcription factor MYB62-like | 5.1 | 1.04E-08 |
| LOC129304541 | 129304541 | nuclear envelope-associated protein 2-like | 5.1 | 1.21E-11 |
| LOC129312742 | 129312742 | isoliquiritigenin 2'-O-methyltransferase-like | 5.1 | 2.52E-04 |
| LOC129288217 | 129288217 | uncharacterized LOC129288217 | 5.1 | 3.23E-02 |
| LOC129316167 | 129316167 | GRAS family protein TF80-like | 5.1 | 3.84E-03 |
| LOC129285268 | 129285268 | serine carboxypeptidase-like 7 | 5.1 | 4.38E-04 |
| LOC129311585 | 129311585 | dof zinc finger protein DOF3.4-like | 5.1 | 3.22E-18 |
| LOC129295812 | 129295812 | proteinaceous RNase P 1, chloroplastic/mitochondrial-like | 5.1 | 3.08E-02 |
| LOC129304448 | 129304448 | calcium-dependent protein kinase 32-like | 5.1 | 1.23E-16 |
| LOC129300031 | 129300031 | HMG1/2-like protein | 5.1 | 1.15E-05 |
| LOC129296931 | 129296931 | high affinity nitrate transporter 2.7-like | 5.1 | 3.15E-03 |
| LOC129291842 | 129291842 | GRF1-interacting factor 1-like | 5.1 | 1.06E-03 |
| LOC129317334 | 129317334 | GDSL esterase/lipase At1g71250-like | 5.1 | 2.42E-02 |
| LOC129285747 | 129285747 | protein STRUBBELIG-RECEPTOR FAMILY 2 | 5.1 | 1.48E-05 |
| LOC129304445 | 129304445 | probable calcium-binding protein CML46 | 5.1 | 4.26E-06 |
| LOC129306042 | 129306042 | uncharacterized LOC129306042 | 5.1 | 8.40E-06 |
| LOC129307107 | 129307107 | uncharacterized LOC129307107 | 5.1 | 2.79E-12 |
| LOC129298657 | 129298657 | 26.5 kDa heat shock protein, mitochondrial-like | 5.1 | 9.20E-03 |
| LOC129314363 | 129314363 | uncharacterized LOC129314363 | 5.1 | 2.81E-03 |
| LOC129309155 | 129309155 | uncharacterized LOC129309155 | 5.1 | 4.35E-02 |
| LOC129321246 | 129321246 | WUSCHEL-related homeobox 6-like | 5.1 | 3.18E-02 |
| LOC129293213 | 129293213 | cationic amino acid transporter 7, chloroplastic-like | 5.0 | 1.02E-07 |
| LOC129319514 | 129319514 | uncharacterized LOC129319514 | 5.0 | 6.00E-06 |
| LOC129318735 | 129318735 | UPF0496 protein At1g20180-like | 5.0 | 2.49E-02 |
| LOC129305589 | 129305589 | probable F-box protein At5g04010 | 5.0 | 1.03E-15 |
| LOC129295635 | 129295635 | uncharacterized LOC129295635 | 5.0 | 9.57E-03 |
| LOC129320833 | 129320833 | heat stress transcription factor A-2-like | 5.0 | 8.51E-21 |
| LOC129295485 | 129295485 | uncharacterized LOC129295485 | 5.0 | 4.67E-04 |
| LOC129299928 | 129299928 | zinc finger protein ZAT11-like | 5.0 | 2.57E-04 |
| LOC129296549 | 129296549 | U4 spliceosomal RNA | 5.0 | 1.03E-02 |
| LOC129298869 | 129298869 | probable bifunctional TENA-E protein | 5.0 | 2.34E-02 |
| LOC129284594 | 129284594 | uncharacterized protein At2g34160 | 5.0 | 1.35E-04 |
| LOC129299400 | 129299400 | ethylene-responsive transcription factor ERF086-like | 5.0 | 3.08E-02 |
| LOC129322339 | 129322339 | uncharacterized LOC129322339 | 5.0 | 1.48E-03 |
| LOC129322937 | 129322937 | sugar transport protein 10-like | 5.0 | 2.52E-02 |
| LOC129295285 | 129295285 | zinc-finger homeodomain protein 4-like | 5.0 | 3.70E-12 |
| LOC129293855 | 129293855 | photosystem II CP43 reaction center protein-like | 5.0 | 2.09E-02 |
| LOC129315291 | 129315291 | inactive beta-amylase 9-like | 5.0 | 6.68E-13 |

| | | | | |
|---|---|---|---|---|
| LOC129319470 | 129319470 | putative calcium-transporting ATPase 13, plasma membrane-typ | 5.0 | 7.16E-16 |
| LOC129293834 | 129293834 | allene oxide synthase 3-like | 5.0 | 1.45E-07 |
| LOC129318738 | 129318738 | 3-oxo-Delta(4,5)-steroid 5-beta-reductase-like | 5.0 | 1.83E-05 |
| LOC129306929 | 129306929 | high affinity sulfate transporter 2-like | 5.0 | 6.78E-03 |
| LOC129289627 | 129289627 | LOB domain-containing protein 25-like | 5.0 | 2.68E-04 |
| LOC129302134 | 129302134 | oleosin Ara h 15.0101-like | 5.0 | 2.87E-02 |
| LOC129319663 | 129319663 | probable membrane-associated kinase regulator 1 | 5.0 | 4.52E-02 |
| LOC129291061 | 129291061 | protein POLYCHOME | 4.9 | 4.14E-03 |
| LOC129322510 | 129322510 | probable calcium-binding protein CML46 | 4.9 | 5.22E-03 |
| LOC129292791 | 129292791 | uncharacterized LOC129292791 | 4.9 | 3.42E-02 |
| LOC129300835 | 129300835 | 2-oxoglutarate-dependent dioxygenase DAO-like | 4.9 | 3.98E-06 |
| LOC129314688 | 129314688 | albumin-2-like | 4.9 | 1.52E-02 |
| LOC129320003 | 129320003 | probable protein phosphatase 2C 6 | 4.9 | 2.59E-27 |
| LOC129315391 | 129315391 | uncharacterized LOC129315391 | 4.9 | 1.10E-02 |
| LOC129301757 | 129301757 | putative clathrin assembly protein At4g40080 | 4.9 | 8.12E-10 |
| LOC129290542 | 129290542 | putative glucose-6-phosphate 1-epimerase | 4.9 | 6.36E-11 |
| LOC129292022 | 129292022 | uncharacterized LOC129292022 | 4.9 | 3.26E-05 |
| LOC129320904 | 129320904 | auxin-responsive protein SAUR32-like | 4.9 | 5.74E-12 |
| LOC129308049 | 129308049 | MYB-like transcription factor EOBI | 4.9 | 1.21E-14 |
| LOC129308717 | 129308717 | uncharacterized LOC129308717 | 4.9 | 1.06E-20 |
| LOC129317753 | 129317753 | ethylene-responsive transcription factor 3-like | 4.9 | 1.39E-26 |
| LOC129309471 | 129309471 | protein TIFY 10A | 4.9 | 6.77E-13 |
| LOC129306059 | 129306059 | uncharacterized LOC129306059 | 4.9 | 5.41E-04 |
| LOC129303872 | 129303872 | probable inactive receptor kinase At2g26730 | 4.9 | 3.07E-03 |
| LOC129299301 | 129299301 | alcohol dehydrogenase 1-like | 4.8 | 6.06E-24 |
| LOC129313815 | 129313815 | uncharacterized LOC129313815 | 4.8 | 3.04E-29 |
| LOC129309848 | 129309848 | ethylene-responsive transcription factor ERF060-like | 4.8 | 1.03E-06 |
| LOC129318259 | 129318259 | cytochrome P450 86A1 | 4.8 | 3.32E-13 |
| LOC129317641 | 129317641 | transcriptional regulator STERILE APETALA-like | 4.8 | 4.43E-02 |
| LOC129309953 | 129309953 | putative glycine-rich cell wall structural protein 1 | 4.8 | 4.91E-02 |
| LOC129294877 | 129294877 | photosystem II protein D1-like | 4.8 | 1.36E-03 |
| LOC129308949 | 129308949 | BAG family molecular chaperone regulator 6-like | 4.8 | 2.08E-11 |
| LOC129311258 | 129311258 | uncharacterized LOC129311258 | 4.8 | 2.43E-04 |
| LOC129307494 | 129307494 | uncharacterized LOC129307494 | 4.8 | 6.59E-05 |
| LOC129313217 | 129313217 | methylecgonone reductase-like | 4.8 | 3.67E-15 |
| LOC129291177 | 129291177 | centromere protein C-like | 4.8 | 6.79E-06 |
| LOC129304425 | 129304425 | vacuolar iron transporter homolog 2-like | 4.8 | 3.74E-04 |
| LOC129299769 | 129299769 | uncharacterized LOC129299769 | 4.8 | 2.89E-02 |
| LOC129300577 | 129300577 | ethylene-responsive transcription factor ERF113-like | 4.8 | 7.02E-03 |
| LOC129293390 | 129293390 | auxin-responsive protein SAUR21-like | 4.8 | 2.25E-06 |
| LOC129287245 | 129287245 | mitogen-activated protein kinase kinase kinase 17-like | 4.8 | 1.40E-08 |
| LOC129309164 | 129309164 | uncharacterized LOC129309164 | 4.8 | 8.28E-21 |
| LOC129295184 | 129295184 | serine/threonine-protein phosphatase PP2A-2 catalytic subunit-li | 4.8 | 8.45E-04 |
| LOC129298312 | 129298312 | protein JINGUBANG-like | 4.8 | 8.14E-17 |
| LOC129304314 | 129304314 | small nucleolar RNA U3 | 4.8 | 4.66E-02 |
| LOC129318932 | 129318932 | NEP1-interacting protein-like 2 | 4.8 | 6.26E-06 |
| LOC129296389 | 129296389 | uncharacterized LOC129296389 | 4.8 | 1.28E-02 |
| LOC129308590 | 129308590 | 17.3 kDa class I heat shock protein-like | 4.7 | 1.53E-03 |

| | | | | |
|---|---|---|---|---|
| LOC129303737 | 129303737 | fatty acid desaturase 4, chloroplastic-like | 4.7 | 3.80E-02 |
| LOC129306664 | 129306664 | probable galacturonosyltransferase-like 4 | 4.7 | 8.51E-03 |
| LOC129305568 | 129305568 | uncharacterized LOC129305568 | 4.7 | 3.56E-20 |
| LOC129296255 | 129296255 | protein DEHYDRATION-INDUCED 19 homolog 4-like | 4.7 | 3.60E-04 |
| LOC129303730 | 129303730 | dof zinc finger protein DOF1.2-like | 4.7 | 6.52E-08 |
| LOC129320392 | 129320392 | WAT1-related protein At5g64700-like | 4.7 | 4.02E-09 |
| LOC129294620 | 129294620 | TPR repeat-containing thioredoxin TTL1 | 4.7 | 1.33E-15 |
| LOC129310859 | 129310859 | probable serine/threonine-protein kinase PBL11 | 4.7 | 2.60E-05 |
| LOC129319216 | 129319216 | uncharacterized LOC129319216 | 4.7 | 2.64E-02 |
| LOC129304241 | 129304241 | uncharacterized LOC129304241 | 4.7 | 2.49E-04 |
| LOC129293962 | 129293962 | kunitz-type serine protease inhibitor DrTI-like | 4.7 | 7.82E-03 |
| LOC129294116 | 129294116 | aspartic proteinase-like protein 1 | 4.7 | 9.95E-25 |
| LOC129296409 | 129296409 | uncharacterized LOC129296409 | 4.7 | 2.98E-03 |
| LOC129321340 | 129321340 | U-box domain-containing protein 19 | 4.7 | 1.07E-20 |
| LOC129285872 | 129285872 | protein TIFY 6B-like | 4.7 | 2.42E-07 |
| LOC129286685 | 129286685 | WRKY transcription factor 28-like | 4.7 | 2.15E-04 |
| LOC129293847 | 129293847 | salt stress-induced hydrophobic peptide ESI3 | 4.7 | 6.13E-08 |
| LOC129312019 | 129312019 | dehydration-responsive element-binding protein 1E-like | 4.7 | 1.46E-04 |
| LOC129310136 | 129310136 | protein LURP-one-related 6 | 4.7 | 9.59E-04 |
| LOC129290999 | 129290999 | uncharacterized protein C594.04c | 4.7 | 1.48E-11 |
| LOC129318668 | 129318668 | protein GRAVITROPIC IN THE LIGHT 1 | 4.7 | 1.05E-04 |
| LOC129287262 | 129287262 | auxin-responsive protein SAUR71-like | 4.7 | 2.65E-06 |
| LOC129321877 | 129321877 | GDSL esterase/lipase EXL3-like | 4.6 | 9.14E-03 |
| LOC129291477 | 129291477 | auxin-responsive protein IAA1-like | 4.6 | 3.06E-12 |
| LOC129297691 | 129297691 | uncharacterized LOC129297691 | 4.6 | 5.10E-03 |
| LOC129292490 | 129292490 | glycine-rich cell wall structural protein 2-like | 4.6 | 2.16E-02 |
| LOC129298022 | 129298022 | uncharacterized LOC129298022 | 4.6 | 5.06E-04 |
| LOC129314441 | 129314441 | defensin-like protein 1 | 4.6 | 2.13E-02 |
| LOC129295534 | 129295534 | uncharacterized LOC129295534 | 4.6 | 2.86E-04 |
| LOC129305131 | 129305131 | WRKY transcription factor 44-like | 4.6 | 3.62E-04 |
| LOC129287542 | 129287542 | acid phosphatase 1-like | 4.6 | 1.77E-06 |
| LOC129288868 | 129288868 | B-box zinc finger protein 32 | 4.6 | 4.60E-07 |
| LOC129308076 | 129308076 | transcription factor bHLH35 | 4.6 | 5.29E-07 |
| LOC129322368 | 129322368 | probable mannitol dehydrogenase | 4.6 | 2.06E-03 |
| LOC129301790 | 129301790 | disease resistance protein RUN1-like | 4.6 | 9.91E-05 |
| LOC129293151 | 129293151 | subtilisin-like protease SBT2.4 | 4.6 | 1.25E-02 |
| LOC129321112 | 129321112 | malate dehydrogenase, chloroplastic-like | 4.6 | 5.64E-14 |
| LOC129290539 | 129290539 | BAHD acyltransferase DCR-like | 4.6 | 6.80E-09 |
| LOC129312938 | 129312938 | ethylene-responsive transcription factor 4-like | 4.6 | 2.48E-11 |
| LOC129323089 | 129323089 | bidirectional sugar transporter SWEET4-like | 4.5 | 2.51E-02 |
| LOC129292010 | 129292010 | Bowman-Birk type proteinase inhibitor-like | 4.5 | 4.24E-02 |
| LOC129300429 | 129300429 | probable inactive poly [ADP-ribose] polymerase SRO5 | 4.5 | 4.19E-02 |
| LOC129312141 | 129312141 | protein LURP-one-related 17-like | 4.5 | 1.06E-04 |
| LOC129313319 | 129313319 | putative 12-oxophytodienoate reductase 11 | 4.5 | 7.30E-06 |
| LOC129312034 | 129312034 | uncharacterized LOC129312034 | 4.5 | 3.91E-02 |
| LOC129295583 | 129295583 | protein EARLY RESPONSIVE TO DEHYDRATION 15-like | 4.5 | 2.04E-05 |
| LOC129321090 | 129321090 | defensin-like protein 1 | 4.5 | 1.39E-02 |
| LOC129295987 | 129295987 | UDP-glycosyltransferase 87A1-like | 4.5 | 1.48E-04 |

| | | | | |
|---|---|---|---|---|
| LOC129299469 | 129299469 | trihelix transcription factor ASIL2-like | 4.5 | 5.69E-04 |
| LOC129294486 | 129294486 | polcalcin Nic t 1-like | 4.5 | 2.83E-03 |
| LOC129301571 | 129301571 | transcription factor ICE1-like | 4.5 | 6.65E-07 |
| LOC129314535 | 129314535 | uncharacterized LOC129314535 | 4.5 | 9.56E-27 |
| LOC129307638 | 129307638 | 11-beta-hydroxysteroid dehydrogenase A-like | 4.5 | 2.19E-02 |
| LOC129294571 | 129294571 | putative clathrin assembly protein At4g40080 | 4.5 | 1.87E-07 |
| LOC129297422 | 129297422 | probable serine/threonine-protein kinase At1g54610 | 4.5 | 4.84E-04 |
| LOC129320815 | 129320815 | uncharacterized LOC129320815 | 4.5 | 2.81E-02 |
| LOC129320644 | 129320644 | temperature-induced lipocalin-1-like | 4.5 | 8.99E-07 |
| LOC129305498 | 129305498 | probable starch synthase 4, chloroplastic/amyloplastic | 4.5 | 3.91E-11 |
| LOC129308687 | 129308687 | uncharacterized LOC129308687 | 4.5 | 3.08E-05 |
| LOC129295759 | 129295759 | eukaryotic translation initiation factor 1A-like | 4.5 | 1.22E-04 |
| LOC129293534 | 129293534 | uncharacterized LOC129293534 | 4.5 | 1.22E-02 |
| LOC129319981 | 129319981 | uncharacterized LOC129319981 | 4.4 | 2.99E-02 |
| LOC129299027 | 129299027 | uncharacterized LOC129299027 | 4.4 | 2.39E-03 |
| LOC129313227 | 129313227 | uncharacterized LOC129313227 | 4.4 | 4.76E-03 |
| LOC129294450 | 129294450 | peroxidase P7 | 4.4 | 1.39E-04 |
| LOC129295818 | 129295818 | ferritin-3, chloroplastic-like | 4.4 | 1.56E-22 |
| LOC129309325 | 129309325 | probable WRKY transcription factor 48 | 4.4 | 5.94E-13 |
| LOC129299592 | 129299592 | uncharacterized LOC129299592 | 4.4 | 2.79E-03 |
| LOC129295557 | 129295557 | uncharacterized LOC129295557 | 4.4 | 1.13E-10 |
| LOC129315005 | 129315005 | putative RING-H2 finger protein ATL69 | 4.4 | 2.90E-03 |
| LOC129303287 | 129303287 | uncharacterized LOC129303287 | 4.4 | 4.05E-02 |
| LOC129307661 | 129307661 | MYB-like transcription factor EOBII | 4.4 | 4.48E-05 |
| LOC129311263 | 129311263 | mitotic checkpoint protein BUB3.3 | 4.4 | 1.26E-02 |
| LOC129323001 | 129323001 | homeobox-leucine zipper protein ATHB-40 | 4.4 | 4.24E-06 |
| LOC129314211 | 129314211 | uncharacterized LOC129314211 | 4.4 | 7.38E-06 |
| LOC129322763 | 129322763 | uncharacterized LOC129322763 | 4.4 | 1.17E-13 |
| LOC129288722 | 129288722 | uncharacterized LOC129288722 | 4.4 | 7.30E-10 |
| LOC129316974 | 129316974 | U1 spliceosomal RNA | 4.4 | 3.12E-02 |
| LOC129286089 | 129286089 | UDP-glycosyltransferase 83A1-like | 4.4 | 2.76E-07 |
| LOC129302328 | 129302328 | protein IQ-DOMAIN 6 | 4.4 | 4.54E-15 |
| LOC129311700 | 129311700 | transcription factor HEC1-like | 4.4 | 1.56E-09 |
| LOC129320951 | 129320951 | protein SAR DEFICIENT 1-like | 4.4 | 2.30E-03 |
| LOC129284887 | 129284887 | uncharacterized LOC129284887 | 4.4 | 5.88E-04 |
| LOC129321711 | 129321711 | protein DETOXIFICATION 49-like | 4.4 | 5.41E-08 |
| LOC129310890 | 129310890 | uncharacterized LOC129310890 | 4.4 | 4.92E-02 |
| LOC129322401 | 129322401 | classical arabinogalactan protein 4-like | 4.4 | 3.79E-33 |
| LOC129317745 | 129317745 | peptidyl-prolyl cis-trans isomerase | 4.4 | 1.24E-11 |
| LOC129285432 | 129285432 | uncharacterized LOC129285432 | 4.4 | 5.48E-14 |
| LOC129287692 | 129287692 | uncharacterized LOC129287692 | 4.3 | 2.50E-24 |
| LOC129318444 | 129318444 | bZIP transcription factor TRAB1-like | 4.3 | 2.20E-50 |
| LOC129291703 | 129291703 | stemmadenine O-acetyltransferase-like | 4.3 | 5.81E-03 |
| LOC129287802 | 129287802 | delta-1-pyrroline-5-carboxylate synthase-like | 4.3 | 4.51E-19 |
| LOC129307244 | 129307244 | monothiol glutaredoxin-S6-like | 4.3 | 1.82E-04 |
| LOC129312680 | 129312680 | ubiquitin-like protein 5 | 4.3 | 1.55E-04 |
| LOC129305540 | 129305540 | uncharacterized LOC129305540 | 4.3 | 3.10E-05 |
| LOC129310132 | 129310132 | cystathionine beta-lyase, chloroplastic-like | 4.3 | 1.56E-03 |

| | | | | |
|---|---|---|---|---|
| LOC129295600 | 129295600 | 26S proteasome non-ATPase regulatory subunit 14 homolog | 4.3 | 2.69E-04 |
| LOC129306537 | 129306537 | protein trichome birefringence-like | 4.3 | 4.67E-44 |
| LOC129293297 | 129293297 | vacuolar iron transporter homolog 1-like | 4.3 | 9.60E-03 |
| LOC129305963 | 129305963 | uncharacterized LOC129305963 | 4.3 | 5.01E-08 |
| LOC129315239 | 129315239 | uncharacterized LOC129315239 | 4.3 | 1.12E-02 |
| LOC129307161 | 129307161 | glutaredoxin-C9-like | 4.3 | 2.60E-07 |
| LOC129286952 | 129286952 | CBL-interacting serine/threonine-protein kinase 20 | 4.3 | 4.48E-09 |
| LOC129309947 | 129309947 | chorismate synthase, chloroplastic-like | 4.3 | 9.68E-04 |
| LOC129290471 | 129290471 | cyclin-A2-4 | 4.2 | 1.19E-02 |
| LOC129299683 | 129299683 | glyoxylase I 4-like | 4.2 | 6.99E-08 |
| LOC129289573 | 129289573 | cytochrome P450 76C2-like | 4.2 | 3.06E-02 |
| LOC129320230 | 129320230 | uncharacterized LOC129320230 | 4.2 | 9.78E-10 |
| LOC129292286 | 129292286 | 5'-adenylylsulfate reductase 3, chloroplastic-like | 4.2 | 2.11E-09 |
| LOC129314928 | 129314928 | zinc finger protein ZAT11-like | 4.2 | 1.69E-02 |
| LOC129287767 | 129287767 | uncharacterized LOC129287767 | 4.2 | 1.66E-02 |
| LOC129305072 | 129305072 | remorin 4.1-like | 4.2 | 5.35E-15 |
| LOC129285727 | 129285727 | serine carboxypeptidase-like 16 | 4.2 | 7.35E-03 |
| LOC129315726 | 129315726 | disease resistance protein RUN1-like | 4.2 | 1.16E-02 |
| LOC129322730 | 129322730 | ethylene-responsive transcription factor 1B-like | 4.2 | 6.54E-07 |
| LOC129297001 | 129297001 | multiprotein-bridging factor 1c-like | 4.2 | 4.18E-02 |
| LOC129307027 | 129307027 | ethylene-responsive transcription factor ERF094-like | 4.2 | 1.05E-04 |
| LOC129309920 | 129309920 | uncharacterized LOC129309920 | 4.2 | 2.31E-22 |
| LOC129318707 | 129318707 | probable ribosome biogenesis protein RLP24 | 4.2 | 4.45E-02 |
| LOC129289868 | 129289868 | cell number regulator 8-like | 4.2 | 2.12E-03 |
| LOC129301542 | 129301542 | cellulose synthase-like protein E6 | 4.2 | 1.87E-04 |
| LOC129292558 | 129292558 | glucose-6-phosphate/phosphate translocator 2, chloroplastic-like | 4.2 | 4.09E-02 |
| LOC129313102 | 129313102 | calcium-dependent protein kinase 24 | 4.2 | 4.07E-12 |
| LOC129286644 | 129286644 | bZIP transcription factor 12-like | 4.2 | 8.79E-10 |
| LOC129294572 | 129294572 | putative clathrin assembly protein At4g40080 | 4.2 | 9.33E-08 |
| LOC129317968 | 129317968 | shewanella-like protein phosphatase 1 | 4.2 | 5.26E-07 |
| LOC129292274 | 129292274 | xyloglucan galactosyltransferase XLT2 | 4.2 | 2.47E-12 |
| LOC129292691 | 129292691 | uncharacterized protein At4g06744-like | 4.2 | 2.01E-02 |
| LOC129304449 | 129304449 | monothiol glutaredoxin-S6-like | 4.2 | 3.72E-03 |
| LOC129309989 | 129309989 | stress enhanced protein 2, chloroplastic-like | 4.2 | 4.01E-03 |
| LOC129313103 | 129313103 | aquaporin TIP1-2-like | 4.2 | 9.93E-13 |
| LOC129317581 | 129317581 | transcription repressor OFP1-like | 4.2 | 5.21E-10 |
| LOC129305013 | 129305013 | ethylene-responsive transcription factor 4-like | 4.2 | 6.12E-09 |
| LOC129303890 | 129303890 | protein REVEILLE 1 | 4.1 | 5.76E-16 |
| LOC129294082 | 129294082 | probable carboxylesterase 17 | 4.1 | 4.67E-02 |
| LOC129309799 | 129309799 | uncharacterized LOC129309799 | 4.1 | 6.35E-11 |
| LOC129321027 | 129321027 | PLAT domain-containing protein 3-like | 4.1 | 1.14E-05 |
| LOC129286883 | 129286883 | probable histone H2A.3 | 4.1 | 2.57E-02 |
| LOC129300654 | 129300654 | condensin-1 complex subunit CAP-D2-like | 4.1 | 3.07E-02 |
| LOC129294269 | 129294269 | NAC domain-containing protein 2-like | 4.1 | 1.09E-05 |
| LOC129308561 | 129308561 | abscisic stress-ripening protein 2-like | 4.1 | 1.27E-03 |
| LOC129312904 | 129312904 | external alternative NAD(P)H-ubiquinone oxidoreductase B2, m | 4.1 | 9.08E-06 |
| LOC129290494 | 129290494 | uncharacterized LOC129290494 | 4.1 | 3.74E-03 |
| LOC129321299 | 129321299 | probable CoA ligase CCL5 | 4.1 | 4.19E-02 |

| | | | | |
|---|---|---|---|---|
| LOC129284370 | 129284370 | zinc finger protein JAGGED-like | 4.1 | 8.57E-07 |
| LOC129317191 | 129317191 | zinc finger protein ZAT11-like | 4.1 | 2.01E-05 |
| LOC129297696 | 129297696 | uncharacterized LOC129297696 | 4.1 | 1.74E-08 |
| LOC129292516 | 129292516 | REF/SRPP-like protein At1g67360 | 4.1 | 1.63E-04 |
| LOC129300282 | 129300282 | F-box protein PP2-A12-like | 4.1 | 1.77E-09 |
| LOC129314334 | 129314334 | uncharacterized LOC129314334 | 4.1 | 2.15E-02 |
| LOC129317661 | 129317661 | RNA pseudouridine synthase 2, chloroplastic | 4.1 | 3.58E-18 |
| LOC129301102 | 129301102 | 60S ribosomal protein L18a-like protein | 4.1 | 1.61E-03 |
| LOC129309751 | 129309751 | CDT1-like protein a, chloroplastic | 4.1 | 5.57E-07 |
| LOC129304232 | 129304232 | uncharacterized LOC129304232 | 4.1 | 5.10E-03 |
| LOC129315315 | 129315315 | two-component response regulator ORR9-like | 4.1 | 4.94E-02 |
| LOC129305305 | 129305305 | uncharacterized LOC129305305 | 4.1 | 4.84E-07 |
| LOC129290800 | 129290800 | uncharacterized LOC129290800 | 4.1 | 3.27E-05 |
| LOC129305862 | 129305862 | probable F-box protein At2g36090 | 4.1 | 3.23E-03 |
| LOC129316263 | 129316263 | protein JINGUBANG-like | 4.1 | 9.59E-04 |
| LOC129287922 | 129287922 | serine/threonine-protein kinase OXI1-like | 4.1 | 1.13E-06 |
| LOC129305094 | 129305094 | transcription factor BEE 1 | 4.0 | 1.15E-03 |
| LOC129313649 | 129313649 | protein SOSEKI 5 | 4.0 | 7.03E-08 |
| LOC129304834 | 129304834 | gibberellin 2-beta-dioxygenase-like | 4.0 | 3.40E-15 |
| LOC129311202 | 129311202 | transcription factor MYB3-like | 4.0 | 1.10E-02 |
| LOC129294860 | 129294860 | uncharacterized protein ORF91 | 4.0 | 1.09E-02 |
| LOC129316789 | 129316789 | trans-resveratrol di-O-methyltransferase-like | 4.0 | 1.18E-04 |
| LOC129311134 | 129311134 | allene oxide cyclase, chloroplastic-like | 4.0 | 4.28E-05 |
| LOC129301492 | 129301492 | GDSL esterase/lipase At4g10955-like | 4.0 | 1.66E-02 |
| LOC129304616 | 129304616 | transcription factor bHLH162-like | 4.0 | 2.86E-03 |
| LOC129291316 | 129291316 | uncharacterized protein At5g48480 | 4.0 | 1.53E-16 |
| LOC129306499 | 129306499 | protein SINE1-like | 4.0 | 1.82E-05 |
| LOC129287606 | 129287606 | histone-lysine N-methyltransferase ASHH3 | 4.0 | 1.15E-02 |
| LOC129303547 | 129303547 | serine acetyltransferase 1, chloroplastic-like | 4.0 | 1.28E-04 |
| LOC129321260 | 129321260 | protein LURP-one-related 4-like | 4.0 | 3.67E-03 |
| LOC129307168 | 129307168 | alpha carbonic anhydrase 7-like | 4.0 | 2.67E-02 |
| LOC129305245 | 129305245 | heat stress transcription factor B-3 | 4.0 | 2.96E-04 |
| LOC129286110 | 129286110 | heat stress transcription factor C-1-like | 4.0 | 2.08E-14 |
| LOC129321259 | 129321259 | uncharacterized LOC129321259 | 4.0 | 4.63E-02 |
| LOC129320649 | 129320649 | protein phosphatase 2C 50-like | 4.0 | 3.67E-15 |
| LOC129310104 | 129310104 | ATP sulfurylase 1, chloroplastic-like | 4.0 | 4.82E-10 |
| LOC129290649 | 129290649 | PR5-like receptor kinase | 4.0 | 3.88E-02 |
| LOC129316127 | 129316127 | squamosa promoter-binding protein 1-like | 4.0 | 1.34E-14 |
| LOC129318846 | 129318846 | zinc finger CCCH domain-containing protein 20-like | 4.0 | 3.95E-12 |
| LOC129285819 | 129285819 | enoyl-CoA delta isomerase 1, peroxisomal | 4.0 | 8.40E-09 |
| LOC129304006 | 129304006 | probable calcium-binding protein CML27 | 4.0 | 6.01E-15 |
| LOC129318090 | 129318090 | cinnamoyl-CoA reductase 1-like | 4.0 | 3.79E-13 |
| LOC129305733 | 129305733 | xyloglucan endotransglucosylase/hydrolase 1 | 4.0 | 3.24E-03 |
| LOC129314454 | 129314454 | uncharacterized LOC129314454 | 4.0 | 2.36E-04 |
| LOC129288805 | 129288805 | CASP-like protein 3A1 | 4.0 | 1.62E-02 |
| LOC129292513 | 129292513 | CBS domain-containing protein CBSX1, chloroplastic-like | 4.0 | 9.78E-07 |
| LOC129313588 | 129313588 | DNA topoisomerase 2 | 4.0 | 2.30E-04 |
| LOC129304547 | 129304547 | putative cyclin-D6-1 | 4.0 | 2.00E-07 |

| | | | | |
|---|---|---|---|---|
| LOC129289652 | 129289652 | glyoxylase I 4-like | 4.0 | 1.04E-04 |
| LOC129302173 | 129302173 | uncharacterized LOC129302173 | 4.0 | 1.22E-12 |
| LOC129309366 | 129309366 | rRNA-processing protein fcf2-like | 3.9 | 2.16E-02 |
| LOC129322893 | 129322893 | uncharacterized LOC129322893 | 3.9 | 1.36E-09 |
| LOC129313014 | 129313014 | nuclear transcription factor Y subunit A-1-like | 3.9 | 1.27E-15 |
| LOC129301188 | 129301188 | membrane-anchored ubiquitin-fold protein 3-like | 3.9 | 2.73E-02 |
| LOC129295988 | 129295988 | cinnamoyl-CoA reductase 1-like | 3.9 | 1.12E-08 |
| LOC129295515 | 129295515 | uncharacterized LOC129295515 | 3.9 | 4.82E-04 |
| LOC129307081 | 129307081 | sugar transport protein 1-like | 3.9 | 3.80E-02 |
| LOC129287260 | 129287260 | protein TIFY 5A-like | 3.9 | 1.09E-04 |
| LOC129322367 | 129322367 | probable mannitol dehydrogenase | 3.9 | 6.60E-03 |
| LOC129303843 | 129303843 | putative lipid-transfer protein DIR1 | 3.9 | 3.22E-03 |
| LOC129288674 | 129288674 | clathrin light chain 3-like | 3.9 | 1.56E-12 |
| LOC129321078 | 129321078 | probable WRKY transcription factor 75 | 3.9 | 1.88E-08 |
| LOC129318810 | 129318810 | probable protein phosphatase 2C 25 | 3.9 | 4.10E-06 |
| LOC129312105 | 129312105 | uncharacterized LOC129312105 | 3.9 | 8.33E-03 |
| LOC129297026 | 129297026 | uncharacterized LOC129297026 | 3.9 | 2.09E-02 |
| LOC129292208 | 129292208 | uncharacterized LOC129292208 | 3.9 | 7.17E-17 |
| LOC129291807 | 129291807 | F-box/kelch-repeat protein At1g80440 | 3.9 | 5.54E-09 |
| LOC129291688 | 129291688 | auxin-responsive protein IAA20-like | 3.9 | 1.09E-05 |
| LOC129304113 | 129304113 | protein phosphatase 2C 37-like | 3.9 | 5.82E-16 |
| LOC129322271 | 129322271 | cysteine proteinase COT44-like | 3.9 | 3.78E-05 |
| LOC129295898 | 129295898 | uncharacterized LOC129295898 | 3.9 | 7.59E-03 |
| LOC129284826 | 129284826 | uncharacterized LOC129284826 | 3.9 | 3.48E-02 |
| LOC129310113 | 129310113 | uncharacterized LOC129310113 | 3.9 | 1.80E-17 |
| LOC129294717 | 129294717 | heavy metal-associated isoprenylated plant protein 9-like | 3.9 | 9.99E-04 |
| LOC129299893 | 129299893 | seed biotin-containing protein SBP65-like | 3.9 | 2.53E-10 |
| LOC129288637 | 129288637 | COP9 signalosome complex subunit 1-like | 3.9 | 2.54E-02 |
| LOC129320358 | 129320358 | inositol transporter 1-like | 3.9 | 7.89E-03 |
| LOC129298080 | 129298080 | F-box protein SKP2A-like | 3.9 | 2.69E-09 |
| LOC129308634 | 129308634 | S-protein homolog 2-like | 3.9 | 6.04E-03 |
| LOC129320972 | 129320972 | uncharacterized LOC129320972 | 3.9 | 2.87E-06 |
| LOC129301555 | 129301555 | ATP synthase small subunit 6-A, mitochondrial-like | 3.9 | 3.40E-02 |
| LOC129301058 | 129301058 | 28S ribosomal RNA | 3.8 | 1.05E-02 |
| LOC129318556 | 129318556 | protein translation factor SUI1 homolog 2 | 3.8 | 1.35E-06 |
| LOC129309344 | 129309344 | protein CDC73 homolog | 3.8 | 1.73E-07 |
| LOC129310277 | 129310277 | glycosyltransferase 6-like | 3.8 | 5.11E-03 |
| LOC129311232 | 129311232 | uncharacterized LOC129311232 | 3.8 | 1.48E-02 |
| LOC129301609 | 129301609 | two-component response regulator ARR17-like | 3.8 | 1.39E-02 |
| LOC129323036 | 129323036 | uncharacterized LOC129323036 | 3.8 | 1.36E-09 |
| LOC129301096 | 129301096 | uncharacterized LOC129301096 | 3.8 | 3.35E-03 |
| LOC129311571 | 129311571 | cysteine-rich receptor-like protein kinase 3 | 3.8 | 2.14E-06 |
| LOC129286885 | 129286885 | serine/threonine-protein kinase Aurora-3 | 3.8 | 3.09E-05 |
| LOC129288169 | 129288169 | probable glutathione S-transferase | 3.8 | 1.85E-02 |
| LOC129302482 | 129302482 | auxin response factor 6-like | 3.8 | 3.43E-03 |
| LOC129321524 | 129321524 | protein DETOXIFICATION 29-like | 3.8 | 8.33E-03 |
| LOC129317893 | 129317893 | histone H4 | 3.8 | 1.59E-03 |
| LOC129310920 | 129310920 | putative SNAP25 homologous protein SNAP30 | 3.8 | 2.07E-02 |

| | | | | |
|---|---|---|---|---|
| LOC129302191 | 129302191 | uncharacterized LOC129302191 | 3.8 | 1.02E-02 |
| LOC129299871 | 129299871 | uncharacterized LOC129299871 | 3.8 | 3.58E-04 |
| LOC129301005 | 129301005 | probable membrane-associated kinase regulator 1 | 3.8 | 4.92E-02 |
| LOC129302475 | 129302475 | chaperonin-like RbcX protein 2, chloroplastic | 3.8 | 3.94E-10 |
| LOC129303408 | 129303408 | uncharacterized LOC129303408 | 3.8 | 4.58E-10 |
| LOC129319024 | 129319024 | uncharacterized LOC129319024 | 3.8 | 6.33E-03 |
| LOC129321768 | 129321768 | phosphatidylinositol 4-kinase gamma 2 | 3.8 | 7.45E-03 |
| LOC129322672 | 129322672 | 1-aminocyclopropane-1-carboxylate synthase | 3.8 | 1.24E-06 |
| LOC129312125 | 129312125 | heat shock factor protein HSF24-like | 3.8 | 1.19E-08 |
| LOC129306339 | 129306339 | uncharacterized LOC129306339 | 3.8 | 1.70E-10 |
| LOC129293273 | 129293273 | zinc finger protein ZAT10-like | 3.8 | 8.67E-11 |
| LOC129320677 | 129320677 | protein ALP1-like | 3.8 | 3.03E-06 |
| LOC129293414 | 129293414 | rac-like GTP-binding protein ARAC7 | 3.8 | 3.80E-02 |
| LOC129321355 | 129321355 | actin-interacting protein 1-2-like | 3.8 | 8.08E-03 |
| LOC129309204 | 129309204 | hsp70-Hsp90 organizing protein 3-like | 3.8 | 1.11E-03 |
| LOC129295258 | 129295258 | uncharacterized LOC129295258 | 3.8 | 4.48E-02 |
| LOC129318047 | 129318047 | V-type proton ATPase subunit a3-like | 3.8 | 3.07E-04 |
| LOC129294235 | 129294235 | uncharacterized LOC129294235 | 3.8 | 6.36E-03 |
| LOC129311000 | 129311000 | uncharacterized LOC129311000 | 3.8 | 6.62E-07 |
| LOC129304546 | 129304546 | LRR receptor-like serine/threonine-protein kinase GSO1 | 3.8 | 2.71E-02 |
| LOC129293261 | 129293261 | allene oxide synthase 2-like | 3.8 | 7.26E-07 |
| LOC129296265 | 129296265 | probable choline kinase 2 | 3.7 | 4.16E-02 |
| LOC129296187 | 129296187 | 28S ribosomal RNA | 3.7 | 2.41E-02 |
| LOC129321668 | 129321668 | probable lysophospholipase BODYGUARD 1 | 3.7 | 1.03E-02 |
| LOC129311963 | 129311963 | E3 ubiquitin-protein ligase RHA2A-like | 3.7 | 7.76E-14 |
| LOC129314270 | 129314270 | neoxanthin synthase, chloroplastic-like | 3.7 | 6.14E-04 |
| LOC129290545 | 129290545 | putative disease resistance protein At5g47280 | 3.7 | 1.06E-02 |
| LOC129299574 | 129299574 | G2/mitotic-specific cyclin S13-7-like | 3.7 | 2.95E-05 |
| LOC129301834 | 129301834 | gibberellin 2-beta-dioxygenase 2 | 3.7 | 2.40E-06 |
| LOC129318106 | 129318106 | beta-amyrin synthase-like | 3.7 | 1.07E-03 |
| LOC129296448 | 129296448 | U1 spliceosomal RNA | 3.7 | 2.83E-03 |
| LOC129295615 | 129295615 | phosphatidylinositol 4-phosphate 5-kinase 9-like | 3.7 | 7.59E-03 |
| LOC129310114 | 129310114 | uncharacterized protein At4g08330, chloroplastic-like | 3.7 | 6.15E-04 |
| LOC129299220 | 129299220 | 9-cis-epoxycarotenoid dioxygenase NCED2, chloroplastic-like | 3.7 | 3.23E-02 |
| LOC129298135 | 129298135 | zinc finger CCCH domain-containing protein 20-like | 3.7 | 8.81E-15 |
| LOC129308668 | 129308668 | chalcone synthase 1-like | 3.7 | 1.77E-02 |
| LOC129299681 | 129299681 | receptor-like serine/threonine-protein kinase SD1-8 | 3.7 | 2.84E-04 |
| LOC129288421 | 129288421 | uncharacterized LOC129288421 | 3.7 | 8.37E-03 |
| LOC129299386 | 129299386 | trihelix transcription factor ENAP1-like | 3.7 | 2.93E-02 |
| LOC129314857 | 129314857 | carboxyl-terminal-processing peptidase 3, chloroplastic | 3.7 | 9.05E-08 |
| LOC129284324 | 129284324 | kinetochore protein SPC25 homolog | 3.7 | 1.90E-02 |
| LOC129287809 | 129287809 | UPF0481 protein At3g47200-like | 3.7 | 3.80E-02 |
| LOC129292060 | 129292060 | uncharacterized LOC129292060 | 3.7 | 3.24E-05 |
| LOC129313009 | 129313009 | ethylene-responsive transcription factor 4 | 3.7 | 6.60E-14 |
| LOC129304892 | 129304892 | protein C2-DOMAIN ABA-RELATED 11-like | 3.7 | 2.63E-16 |
| LOC129291495 | 129291495 | cell division control protein 2 homolog C | 3.7 | 1.28E-02 |
| LOC129309599 | 129309599 | S-adenosylmethionine synthase 2-like | 3.7 | 5.21E-10 |
| LOC129319657 | 129319657 | serine/threonine-protein kinase D6PK-like | 3.7 | 1.46E-02 |

| | | | | |
|---|---|---|---|---|
| LOC129288991 | 129288991 | uncharacterized LOC129288991 | 3.7 | 1.51E-07 |
| LOC129294810 | 129294810 | expansin-A8-like | 3.7 | 1.36E-04 |
| LOC129303577 | 129303577 | heavy metal-associated isoprenylated plant protein 9-like | 3.7 | 7.45E-08 |
| LOC129291018 | 129291018 | two-component response regulator-like APRR9 | 3.7 | 6.36E-06 |
| LOC129291225 | 129291225 | protein SULFUR DEFICIENCY-INDUCED 1-like | 3.7 | 1.24E-11 |
| LOC129304096 | 129304096 | chaperone protein dnaJ 11, chloroplastic-like | 3.7 | 4.34E-10 |
| LOC129306856 | 129306856 | protein S40-1-like | 3.7 | 5.20E-10 |
| LOC129309506 | 129309506 | galactinol--sucrose galactosyltransferase-like | 3.7 | 1.41E-03 |
| LOC129293770 | 129293770 | probable choline kinase 2 | 3.7 | 4.78E-03 |
| LOC129284434 | 129284434 | F-box/LRR-repeat protein 4 | 3.7 | 1.35E-03 |
| LOC129320631 | 129320631 | calcium-binding allergen Ole e 8-like | 3.6 | 1.26E-11 |
| LOC129295660 | 129295660 | putative RING-H2 finger protein ATL69 | 3.6 | 3.38E-03 |
| LOC129303087 | 129303087 | probable amino acid permease 7 | 3.6 | 7.62E-12 |
| LOC129304036 | 129304036 | heterodimeric geranylgeranyl pyrophosphate synthase small sub | 3.6 | 4.70E-04 |
| LOC129298519 | 129298519 | uncharacterized LOC129298519 | 3.6 | 8.11E-04 |
| LOC129300392 | 129300392 | aquaporin PIP2-2 | 3.6 | 2.70E-07 |
| LOC129319019 | 129319019 | beta-carotene hydroxylase 2, chloroplastic-like | 3.6 | 3.21E-09 |
| LOC129296232 | 129296232 | photosystem II core complex proteins psbY, chloroplastic-like | 3.6 | 1.87E-02 |
| LOC129310680 | 129310680 | ethylene-responsive transcription factor 1A-like | 3.6 | 1.05E-04 |
| LOC129291456 | 129291456 | squamosa promoter-binding-like protein 7 | 3.6 | 5.66E-06 |
| LOC129296440 | 129296440 | uncharacterized LOC129296440 | 3.6 | 4.47E-09 |
| LOC129286740 | 129286740 | uncharacterized LOC129286740 | 3.6 | 4.18E-08 |
| LOC129285770 | 129285770 | universal stress protein PHOS32 | 3.6 | 3.98E-06 |
| LOC129311990 | 129311990 | protein MIZU-KUSSEI 1 | 3.6 | 9.60E-03 |
| LOC129299869 | 129299869 | PRA1 family protein B4-like | 3.6 | 2.95E-07 |
| LOC129284926 | 129284926 | uncharacterized LOC129284926 | 3.6 | 1.20E-03 |
| LOC129284849 | 129284849 | cationic amino acid transporter 5-like | 3.6 | 1.85E-07 |
| LOC129287683 | 129287683 | uncharacterized LOC129287683 | 3.6 | 1.21E-02 |
| LOC129303822 | 129303822 | heavy metal-associated isoprenylated plant protein 24 | 3.6 | 1.14E-03 |
| LOC129322715 | 129322715 | probable nucleoredoxin 1 | 3.6 | 1.03E-02 |
| LOC129302540 | 129302540 | cell wall / vacuolar inhibitor of fructosidase 2-like | 3.6 | 7.01E-03 |
| LOC129300554 | 129300554 | GTP-binding protein YPTM2 | 3.6 | 4.18E-02 |
| LOC129303654 | 129303654 | glycosyltransferase BC10 | 3.6 | 1.21E-07 |
| LOC129287693 | 129287693 | uncharacterized LOC129287693 | 3.6 | 3.08E-08 |
| LOC129310657 | 129310657 | plasmodesmata-located protein 6-like | 3.6 | 1.30E-08 |
| LOC129305075 | 129305075 | NDR1/HIN1-like protein 1 | 3.6 | 9.40E-12 |
| LOC129315492 | 129315492 | myosin-12 | 3.6 | 1.80E-10 |
| LOC129321086 | 129321086 | calcium-binding protein CML37-like | 3.6 | 6.36E-06 |
| LOC129308637 | 129308637 | mitochondrial phosphate carrier protein 3, mitochondrial-like | 3.6 | 8.55E-04 |
| LOC129297418 | 129297418 | metallothionein-like protein 2 | 3.6 | 8.40E-04 |
| LOC129318630 | 129318630 | uncharacterized LOC129318630 | 3.6 | 5.56E-04 |
| LOC129294375 | 129294375 | uncharacterized LOC129294375 | 3.6 | 4.57E-02 |
| LOC129321627 | 129321627 | putative pentatricopeptide repeat-containing protein At1g26500 | 3.6 | 1.66E-02 |
| LOC129302427 | 129302427 | formin-like protein 20 | 3.6 | 1.02E-03 |
| LOC129294974 | 129294974 | probable galacturonosyltransferase-like 9 | 3.6 | 3.90E-03 |
| LOC129293861 | 129293861 | uncharacterized LOC129293861 | 3.6 | 4.21E-03 |
| LOC129318598 | 129318598 | protein CYSTEINE-RICH TRANSMEMBRANE MODULE 9- | 3.6 | 8.23E-04 |
| LOC129311259 | 129311259 | ninja-family protein AFP2-like | 3.6 | 7.20E-10 |

| | | | | |
|---|---|---|---|---|
| LOC129322656 | 129322656 | auxin-responsive protein IAA9-like | 3.6 | 2.07E-03 |
| LOC129299630 | 129299630 | uncharacterized protein At4g28440-like | 3.6 | 8.64E-03 |
| LOC129309686 | 129309686 | spermidine synthase 1-like | 3.6 | 5.03E-05 |
| LOC129310994 | 129310994 | protein IQ-DOMAIN 22 | 3.6 | 1.32E-04 |
| LOC129311922 | 129311922 | uncharacterized LOC129311922 | 3.6 | 1.69E-06 |
| LOC129292614 | 129292614 | glycine-rich cell wall structural protein 1.8-like | 3.6 | 8.01E-15 |
| LOC129307308 | 129307308 | histone H3.2 | 3.6 | 2.74E-04 |
| LOC129301123 | 129301123 | calcineurin B-like protein 1 | 3.6 | 2.04E-08 |
| LOC129311541 | 129311541 | non-specific lipid transfer protein GPI-anchored 7-like | 3.6 | 2.17E-02 |
| LOC129289001 | 129289001 | protein LURP-one-related 12-like | 3.6 | 8.15E-08 |
| LOC129315936 | 129315936 | ethylene-responsive transcription factor 4 | 3.6 | 3.46E-09 |
| LOC129312159 | 129312159 | uncharacterized LOC129312159 | 3.5 | 1.69E-02 |
| LOC129306850 | 129306850 | F-box protein At1g10780 | 3.5 | 3.77E-03 |
| LOC129306456 | 129306456 | thaumatin-like protein 1b | 3.5 | 4.48E-04 |
| LOC129322393 | 129322393 | uncharacterized LOC129322393 | 3.5 | 5.93E-08 |
| LOC129296348 | 129296348 | F-box/kelch-repeat protein At1g23390-like | 3.5 | 4.22E-08 |
| LOC129293007 | 129293007 | uncharacterized LOC129293007 | 3.5 | 2.06E-04 |
| LOC129298282 | 129298282 | uncharacterized LOC129298282 | 3.5 | 1.27E-02 |
| LOC129322931 | 129322931 | protein DETOXIFICATION 27-like | 3.5 | 9.62E-03 |
| LOC129302469 | 129302469 | uncharacterized LOC129302469 | 3.5 | 1.44E-03 |
| LOC129300048 | 129300048 | neutral/alkaline invertase 3, chloroplastic-like | 3.5 | 9.91E-04 |
| LOC129304828 | 129304828 | U-box domain-containing protein 8-like | 3.5 | 5.88E-10 |
| LOC129285304 | 129285304 | cysteine-rich receptor-like protein kinase 42 | 3.5 | 4.19E-09 |
| LOC129294639 | 129294639 | proteasome subunit beta type-3-A-like | 3.5 | 3.93E-02 |
| LOC129306007 | 129306007 | BAG family molecular chaperone regulator 2 | 3.5 | 5.74E-04 |
| LOC129320572 | 129320572 | 17.5 kDa class I heat shock protein-like | 3.5 | 1.20E-04 |
| LOC129304942 | 129304942 | pentatricopeptide repeat-containing protein At5g06540-like | 3.5 | 1.03E-14 |
| LOC129313282 | 129313282 | dynamin-2B-like | 3.5 | 9.41E-05 |
| LOC129297549 | 129297549 | uncharacterized LOC129297549 | 3.5 | 2.04E-02 |
| LOC129291600 | 129291600 | trihelix transcription factor GT-3b-like | 3.5 | 9.15E-03 |
| LOC129285712 | 129285712 | syntaxin-121-like | 3.5 | 6.35E-07 |
| LOC129320576 | 129320576 | myb-related protein 306-like | 3.5 | 1.80E-08 |
| LOC129300145 | 129300145 | protein CDC73 homolog | 3.5 | 3.72E-03 |
| LOC129308395 | 129308395 | early nodulin-75-like | 3.5 | 1.22E-05 |
| LOC129307880 | 129307880 | pollen receptor-like kinase 3 | 3.5 | 1.25E-05 |
| LOC129320593 | 129320593 | RING-H2 finger protein ATL2-like | 3.5 | 1.04E-04 |
| LOC129290560 | 129290560 | D-3-phosphoglycerate dehydrogenase 3, chloroplastic-like | 3.5 | 2.29E-02 |
| LOC129292673 | 129292673 | two-component response regulator-like APRR7 | 3.5 | 2.50E-03 |
| LOC129306271 | 129306271 | kinesin-like protein KIN-5C | 3.5 | 8.97E-03 |
| LOC129309444 | 129309444 | patellin-3-like | 3.5 | 2.63E-02 |
| LOC129317559 | 129317559 | uncharacterized LOC129317559 | 3.5 | 1.65E-05 |
| LOC129323145 | 129323145 | uncharacterized LOC129323145 | 3.5 | 1.48E-03 |
| LOC129294673 | 129294673 | ethylene-responsive transcription factor CRF4-like | 3.5 | 9.77E-03 |
| LOC129314785 | 129314785 | ABSCISIC ACID-INSENSITIVE 5-like protein 1 | 3.5 | 5.34E-03 |
| LOC129284419 | 129284419 | uncharacterized LOC129284419 | 3.4 | 5.58E-04 |
| LOC129318649 | 129318649 | disease resistance protein UNI-like | 3.4 | 4.67E-02 |
| LOC129307492 | 129307492 | rho GTPase-activating protein 2-like | 3.4 | 8.42E-03 |
| LOC129303006 | 129303006 | uncharacterized LOC129303006 | 3.4 | 6.04E-03 |

| | | | | |
|---|---|---|---|---|
| LOC129306305 | 129306305 | kinesin-like protein KIN-14S | 3.4 | 1.21E-03 |
| LOC129318659 | 129318659 | NDR1/HIN1-like protein 13 | 3.4 | 1.72E-05 |
| LOC129288965 | 129288965 | probable 2-oxoglutarate-dependent dioxygenase AOP1 | 3.4 | 1.26E-04 |
| LOC129284521 | 129284521 | ethylene-responsive transcription factor 1A-like | 3.4 | 8.79E-05 |
| LOC129293873 | 129293873 | NAC domain-containing protein 6-like | 3.4 | 1.62E-04 |
| LOC129322622 | 129322622 | ankyrin repeat-containing protein NPR4-like | 3.4 | 2.09E-02 |
| LOC129316236 | 129316236 | mitotic checkpoint serine/threonine-protein kinase BUB1 | 3.4 | 2.06E-02 |
| LOC129320723 | 129320723 | uncharacterized LOC129320723 | 3.4 | 4.02E-05 |
| LOC129320884 | 129320884 | uncharacterized LOC129320884 | 3.4 | 6.39E-06 |
| LOC129322687 | 129322687 | NAC domain-containing protein 2-like | 3.4 | 4.03E-03 |
| LOC129312466 | 129312466 | tetraspanin-3-like | 3.4 | 3.14E-17 |
| LOC129287838 | 129287838 | cytochrome b5-like | 3.4 | 5.11E-06 |
| LOC129319667 | 129319667 | probable WRKY transcription factor 48 | 3.4 | 5.28E-06 |
| LOC129306995 | 129306995 | vacuolar iron transporter homolog 4-like | 3.4 | 1.69E-03 |
| LOC129293960 | 129293960 | uncharacterized LOC129293960 | 3.4 | 8.61E-12 |
| LOC129308725 | 129308725 | ABC transporter G family member 6-like | 3.4 | 3.08E-03 |
| LOC129311651 | 129311651 | probable pectinesterase 68 | 3.4 | 2.41E-02 |
| LOC129294751 | 129294751 | DDT domain-containing protein DDR4 | 3.4 | 4.89E-02 |
| LOC129295202 | 129295202 | uncharacterized LOC129295202 | 3.4 | 6.52E-08 |
| LOC129306729 | 129306729 | auxin-induced protein 22D-like | 3.4 | 2.30E-11 |
| LOC129319833 | 129319833 | uncharacterized LOC129319833 | 3.4 | 3.07E-09 |
| LOC129304380 | 129304380 | RHOMBOID-like protein 3 | 3.4 | 3.86E-12 |
| LOC129320209 | 129320209 | TORTIFOLIA1-like protein 3 | 3.4 | 4.35E-03 |
| LOC129295864 | 129295864 | uncharacterized LOC129295864 | 3.4 | 5.02E-06 |
| LOC129291817 | 129291817 | uncharacterized LOC129291817 | 3.4 | 5.24E-05 |
| LOC129311889 | 129311889 | uncharacterized LOC129311889 | 3.4 | 4.33E-12 |
| LOC129286003 | 129286003 | patatin-like protein 6 | 3.4 | 5.46E-08 |
| LOC129304988 | 129304988 | alternative oxidase 3, mitochondrial-like | 3.4 | 7.83E-06 |
| LOC129305193 | 129305193 | uncharacterized LOC129305193 | 3.4 | 4.80E-09 |
| LOC129289696 | 129289696 | thaumatin-like protein | 3.4 | 2.02E-02 |
| LOC129303329 | 129303329 | dormancy-associated protein homolog 3 | 3.4 | 1.33E-07 |
| LOC129298510 | 129298510 | putative disease resistance RPP13-like protein 3 | 3.4 | 1.43E-04 |
| LOC129310384 | 129310384 | uncharacterized LOC129310384 | 3.4 | 1.10E-02 |
| LOC129310357 | 129310357 | uncharacterized LOC129310357 | 3.4 | 6.79E-03 |
| LOC129295035 | 129295035 | uncharacterized LOC129295035 | 3.4 | 4.40E-19 |
| LOC129312187 | 129312187 | uncharacterized LOC129312187 | 3.3 | 1.54E-02 |
| LOC129312882 | 129312882 | probable histone H2B.3 | 3.3 | 8.02E-03 |
| LOC129307218 | 129307218 | galactan beta-1,4-galactosyltransferase GALS3 | 3.3 | 9.30E-12 |
| LOC129306557 | 129306557 | serine carboxypeptidase-like 31 | 3.3 | 9.12E-03 |
| LOC129302456 | 129302456 | 17.5 kDa class I heat shock protein-like | 3.3 | 6.44E-06 |
| LOC129294709 | 129294709 | FCS-Like Zinc finger 10-like | 3.3 | 4.69E-04 |
| LOC129317828 | 129317828 | uncharacterized LOC129317828 | 3.3 | 2.98E-09 |
| LOC129286265 | 129286265 | serine carboxypeptidase-like 18 | 3.3 | 3.41E-02 |
| LOC129315347 | 129315347 | P-loop NTPase domain-containing protein LPA1 homolog 1 | 3.3 | 2.12E-02 |
| LOC129285182 | 129285182 | classical arabinogalactan protein 9-like | 3.3 | 3.22E-11 |
| LOC129313501 | 129313501 | hydrophobic protein RCI2B | 3.3 | 7.25E-06 |
| LOC129306677 | 129306677 | protein PHLOEM PROTEIN 2-LIKE A9-like | 3.3 | 1.01E-02 |
| LOC129302820 | 129302820 | proline-rich extensin-like protein EPR1 | 3.3 | 5.05E-03 |

| | | | | |
|---|---|---|---|---|
| LOC129309308 | 129309308 | probable methyltransferase At1g27930 | 3.3 | 4.54E-10 |
| LOC129306740 | 129306740 | 18.5 kDa class I heat shock protein | 3.3 | 1.69E-03 |
| LOC129290534 | 129290534 | uncharacterized LOC129290534 | 3.3 | 1.93E-02 |
| LOC129307152 | 129307152 | pathogenesis-related genes transcriptional activator PTI5-like | 3.3 | 8.49E-04 |
| LOC129290933 | 129290933 | uncharacterized LOC129290933 | 3.3 | 3.58E-02 |
| LOC129294898 | 129294898 | maturase K-like | 3.3 | 3.73E-02 |
| LOC129301702 | 129301702 | probably inactive leucine-rich repeat receptor-like protein kinase | 3.3 | 2.20E-02 |
| LOC129287233 | 129287233 | cysteine-rich and transmembrane domain-containing protein WI | 3.3 | 1.98E-02 |
| LOC129321523 | 129321523 | homeobox-leucine zipper protein HOX20-like | 3.3 | 8.29E-11 |
| LOC129291212 | 129291212 | putative lipid-transfer protein DIR1 | 3.3 | 3.68E-08 |
| LOC129299133 | 129299133 | coatomer subunit delta-like | 3.3 | 3.15E-03 |
| LOC129292495 | 129292495 | G2/mitotic-specific cyclin-2-like | 3.3 | 4.54E-02 |
| LOC129312191 | 129312191 | fatty acid desaturase 4, chloroplastic-like | 3.3 | 4.91E-02 |
| LOC129304259 | 129304259 | putative defensin-like protein 128 | 3.3 | 1.99E-04 |
| LOC129320141 | 129320141 | probable protein phosphatase 2C 14 | 3.3 | 5.77E-03 |
| LOC129297415 | 129297415 | kinesin-like protein KIN-10B | 3.3 | 1.26E-02 |
| LOC129320304 | 129320304 | uncharacterized LOC129320304 | 3.3 | 1.32E-02 |
| LOC129306518 | 129306518 | syntaxin-124-like | 3.3 | 1.01E-04 |
| LOC129294075 | 129294075 | uncharacterized LOC129294075 | 3.3 | 4.26E-04 |
| LOC129307054 | 129307054 | cytochrome P450 CYP736A12-like | 3.3 | 4.74E-06 |
| LOC129310589 | 129310589 | transcription factor TGA2.3-like | 3.2 | 6.75E-04 |
| LOC129313791 | 129313791 | zinc finger protein CONSTANS-LIKE 12-like | 3.2 | 1.06E-04 |
| LOC129299305 | 129299305 | 60S ribosomal protein L24-like | 3.2 | 1.49E-02 |
| LOC129323137 | 129323137 | protein trichome birefringence-like 2 | 3.2 | 4.26E-10 |
| LOC129321073 | 129321073 | histone H3.2 | 3.2 | 7.97E-03 |
| LOC129293581 | 129293581 | brassinosteroid LRR receptor kinase BRL1-like | 3.2 | 9.86E-04 |
| LOC129308594 | 129308594 | kinesin-like protein KIN-10A | 3.2 | 1.39E-08 |
| LOC129306334 | 129306334 | UDP-glucuronic acid decarboxylase 2 | 3.2 | 2.06E-14 |
| LOC129298638 | 129298638 | 22.7 kDa class IV heat shock protein-like | 3.2 | 4.86E-02 |
| LOC129286715 | 129286715 | LOB domain-containing protein 4 | 3.2 | 2.59E-04 |
| LOC129303355 | 129303355 | 17.5 kDa class I heat shock protein-like | 3.2 | 8.17E-05 |
| LOC129305855 | 129305855 | BRCT domain-containing protein At4g02110 | 3.2 | 5.21E-10 |
| LOC129300397 | 129300397 | protein NTM1-like 9 | 3.2 | 1.62E-02 |
| LOC129313772 | 129313772 | uncharacterized LOC129313772 | 3.2 | 8.23E-06 |
| LOC129309833 | 129309833 | ethylene-responsive transcription factor ABR1-like | 3.2 | 2.89E-06 |
| LOC129297936 | 129297936 | 60S ribosomal protein L11-like | 3.2 | 2.34E-02 |
| LOC129320200 | 129320200 | protein MIZU-KUSSEI 1 | 3.2 | 1.40E-05 |
| LOC129308757 | 129308757 | histone H3.2 | 3.2 | 2.82E-02 |
| LOC129313551 | 129313551 | sodium/calcium exchanger NCL-like | 3.2 | 1.89E-09 |
| LOC129309148 | 129309148 | NAC domain-containing protein 2-like | 3.2 | 1.47E-03 |
| LOC129312079 | 129312079 | SNF1-related protein kinase regulatory subunit gamma-1-like | 3.2 | 3.64E-05 |
| LOC129303413 | 129303413 | ABC transporter G family member 23 | 3.2 | 1.80E-03 |
| LOC129313207 | 129313207 | protein REDOX 2-like | 3.2 | 8.92E-13 |
| LOC129292069 | 129292069 | uncharacterized LOC129292069 | 3.2 | 9.09E-03 |
| LOC129295493 | 129295493 | ethylene-responsive transcription factor RAP2-1-like | 3.2 | 3.84E-06 |
| LOC129305916 | 129305916 | protein root UVB sensitive 6-like | 3.2 | 2.35E-11 |
| LOC129300751 | 129300751 | uncharacterized LOC129300751 | 3.2 | 4.07E-03 |
| LOC129314960 | 129314960 | arogenate dehydratase/prephenate dehydratase 6, chloroplastic-li | 3.2 | 8.90E-11 |

| | | | | |
|---|---|---|---|---|
| LOC129317127 | 129317127 | protein DMP2-like | 3.2 | 3.39E-03 |
| LOC129287063 | 129287063 | dirigent protein 23-like | 3.2 | 3.36E-03 |
| LOC129298522 | 129298522 | uncharacterized LOC129298522 | 3.2 | 7.53E-05 |
| LOC129293238 | 129293238 | histone H2A variant 1-like | 3.2 | 9.62E-03 |
| LOC129287069 | 129287069 | protein POLYCHOME-like | 3.2 | 3.78E-03 |
| LOC129307010 | 129307010 | dof zinc finger protein DOF1.5-like | 3.2 | 9.67E-07 |
| LOC129308718 | 129308718 | uncharacterized LOC129308718 | 3.2 | 1.56E-02 |
| LOC129318526 | 129318526 | probable linoleate 9S-lipoxygenase 5 | 3.2 | 1.21E-04 |
| LOC129321010 | 129321010 | HVA22-like protein e | 3.2 | 1.33E-07 |
| LOC129287512 | 129287512 | UPF0481 protein At3g47200-like | 3.2 | 2.85E-06 |
| LOC129302216 | 129302216 | uncharacterized LOC129302216 | 3.2 | 1.18E-02 |
| LOC129294725 | 129294725 | uncharacterized LOC129294725 | 3.2 | 7.31E-03 |
| LOC129306875 | 129306875 | GDSL esterase/lipase At5g03610-like | 3.2 | 4.72E-02 |
| LOC129294073 | 129294073 | remorin 4.2-like | 3.2 | 3.33E-02 |
| LOC129293942 | 129293942 | protein kinase STUNTED | 3.2 | 1.17E-06 |
| LOC129294739 | 129294739 | protein DUF642 L-GALACTONO-1,4-LACTONE-RESPONSI | 3.2 | 1.88E-02 |
| LOC129310096 | 129310096 | ethylene-responsive transcription factor ABR1-like | 3.2 | 4.48E-03 |
| LOC129303693 | 129303693 | probable inorganic phosphate transporter 1-7 | 3.2 | 2.43E-04 |
| LOC129312053 | 129312053 | uncharacterized LOC129312053 | 3.2 | 1.49E-06 |
| LOC129304002 | 129304002 | bZIP transcription factor TRAB1-like | 3.2 | 3.71E-11 |
| LOC129293316 | 129293316 | ethylene-responsive transcription factor RAP2-3-like | 3.2 | 5.99E-07 |
| LOC129290497 | 129290497 | uncharacterized LOC129290497 | 3.1 | 8.64E-05 |
| LOC129298701 | 129298701 | ribulose bisphosphate carboxylase large chain | 3.1 | 3.25E-03 |
| LOC129301652 | 129301652 | ubiquitin-conjugating enzyme E2 20-like | 3.1 | 3.84E-03 |
| LOC129285744 | 129285744 | kinesin-like protein KIN-14C | 3.1 | 1.28E-02 |
| LOC129309577 | 129309577 | protein phosphatase 2C 53-like | 3.1 | 2.47E-15 |
| LOC129294778 | 129294778 | uncharacterized LOC129294778 | 3.1 | 1.94E-05 |
| LOC129291510 | 129291510 | quinone-oxidoreductase homolog, chloroplastic-like | 3.1 | 1.54E-02 |
| LOC129315803 | 129315803 | pentatricopeptide repeat-containing protein At1g34160-like | 3.1 | 1.56E-02 |
| LOC129298702 | 129298702 | ribulose bisphosphate carboxylase large chain | 3.1 | 3.29E-04 |
| LOC129303590 | 129303590 | phospholipase A1-Igamma3, chloroplastic | 3.1 | 2.19E-06 |
| LOC129304072 | 129304072 | uncharacterized LOC129304072 | 3.1 | 4.93E-03 |
| LOC129295720 | 129295720 | uncharacterized LOC129295720 | 3.1 | 1.33E-05 |
| LOC129291656 | 129291656 | uncharacterized LOC129291656 | 3.1 | 1.11E-02 |
| LOC129293460 | 129293460 | uncharacterized LOC129293460 | 3.1 | 2.18E-06 |
| LOC129289489 | 129289489 | uncharacterized LOC129289489 | 3.1 | 3.80E-03 |
| LOC129309353 | 129309353 | uncharacterized LOC129309353 | 3.1 | 8.49E-09 |
| LOC129306627 | 129306627 | NDR1/HIN1-like protein 3 | 3.1 | 7.89E-06 |
| LOC129310611 | 129310611 | NAC domain-containing protein 2-like | 3.1 | 2.68E-03 |
| LOC129319360 | 129319360 | CBL-interacting serine/threonine-protein kinase 4-like | 3.1 | 5.67E-07 |
| LOC129306439 | 129306439 | uncharacterized LOC129306439 | 3.1 | 9.57E-06 |
| LOC129294465 | 129294465 | cytochrome P450 714A1-like | 3.1 | 3.26E-12 |
| LOC129290704 | 129290704 | plastidic ATP/ADP-transporter | 3.1 | 2.21E-14 |
| LOC129322952 | 129322952 | uncharacterized LOC129322952 | 3.1 | 3.74E-03 |
| LOC129312080 | 129312080 | bZIP transcription factor 53-like | 3.1 | 1.05E-04 |
| LOC129315181 | 129315181 | structural maintenance of chromosomes protein 2-1-like | 3.1 | 2.50E-02 |
| LOC129308589 | 129308589 | 17.3 kDa class I heat shock protein | 3.1 | 3.62E-06 |
| LOC129298728 | 129298728 | probable cinnamyl alcohol dehydrogenase 1 | 3.1 | 4.96E-03 |

| | | | | |
|---|---|---|---|---|
| LOC129309948 | 129309948 | uncharacterized LOC129309948 | 3.1 | 4.15E-03 |
| LOC129314666 | 129314666 | protein LNK3-like | 3.1 | 1.30E-08 |
| LOC129320447 | 129320447 | heat shock factor protein HSF30-like | 3.1 | 1.13E-06 |
| LOC129306513 | 129306513 | WRKY transcription factor 28-like | 3.1 | 1.29E-03 |
| LOC129304731 | 129304731 | probable methyltransferase At1g29790 | 3.1 | 2.30E-04 |
| LOC129299232 | 129299232 | aspartic proteinase PCS1-like | 3.1 | 9.58E-08 |
| LOC129309479 | 129309479 | protein QUIRKY | 3.1 | 4.95E-04 |
| LOC129306212 | 129306212 | type IV inositol polyphosphate 5-phosphatase 6-like | 3.1 | 1.99E-08 |
| LOC129292252 | 129292252 | protein DEHYDRATION-INDUCED 19 homolog 4-like | 3.1 | 4.56E-03 |
| LOC129321693 | 129321693 | high mobility group B protein 6 | 3.1 | 2.12E-04 |
| LOC129317168 | 129317168 | basic leucine zipper 4-like | 3.1 | 2.85E-02 |
| LOC129318817 | 129318817 | uncharacterized LOC129318817 | 3.1 | 3.69E-04 |
| LOC129285179 | 129285179 | serine/threonine-protein kinase haspin homolog | 3.1 | 4.80E-02 |
| LOC129303393 | 129303393 | probable E3 ubiquitin-protein ligase XERICO | 3.1 | 1.83E-09 |
| LOC129298970 | 129298970 | stress enhanced protein 2, chloroplastic-like | 3.1 | 7.47E-07 |
| LOC129316068 | 129316068 | zinc finger protein CONSTANS-LIKE 1-like | 3.1 | 8.27E-07 |
| LOC129306786 | 129306786 | remorin 4.1-like | 3.1 | 1.42E-04 |
| LOC129292035 | 129292035 | Bowman-Birk type proteinase inhibitor-like | 3.0 | 2.85E-03 |
| LOC129323006 | 129323006 | uncharacterized LOC129323006 | 3.0 | 1.26E-05 |
| LOC129293721 | 129293721 | serine/threonine protein phosphatase 2A 57 kDa regulatory subu | 3.0 | 2.12E-04 |
| LOC129298703 | 129298703 | ATP synthase subunit beta, chloroplastic-like | 3.0 | 4.94E-02 |
| LOC129290302 | 129290302 | uncharacterized LOC129290302 | 3.0 | 7.12E-06 |
| LOC129317029 | 129317029 | uncharacterized LOC129317029 | 3.0 | 3.50E-04 |
| LOC129297550 | 129297550 | uncharacterized LOC129297550 | 3.0 | 8.10E-06 |
| LOC129307972 | 129307972 | AAA-ATPase At5g57480-like | 3.0 | 1.54E-04 |
| LOC129285413 | 129285413 | probable cinnamyl alcohol dehydrogenase 1 | 3.0 | 2.69E-03 |
| LOC129320876 | 129320876 | glycine-rich cell wall structural protein-like | 3.0 | 1.25E-02 |
| LOC129316330 | 129316330 | uncharacterized LOC129316330 | 3.0 | 2.24E-07 |
| LOC129306718 | 129306718 | uncharacterized LOC129306718 | 3.0 | 2.07E-04 |
| LOC129293421 | 129293421 | F-box/LRR-repeat protein At4g29420 | 3.0 | 1.39E-08 |
| LOC129286764 | 129286764 | protein JINGUBANG-like | 3.0 | 4.15E-09 |
| LOC129291527 | 129291527 | nuclear envelope-associated protein 2 | 3.0 | 4.12E-03 |
| LOC129298079 | 129298079 | VQ motif-containing protein 4-like | 3.0 | 2.78E-05 |
| LOC129314316 | 129314316 | polyamine oxidase 1-like | 3.0 | 4.15E-07 |
| LOC129292943 | 129292943 | F-box protein PP2-B15-like | 3.0 | 1.18E-02 |
| LOC129309846 | 129309846 | calcium-binding protein CP1-like | 3.0 | 2.53E-04 |
| LOC129301701 | 129301701 | uncharacterized LOC129301701 | 3.0 | 3.79E-02 |
| LOC129293283 | 129293283 | cytochrome P450 78A5-like | 3.0 | 7.92E-07 |
| LOC129303420 | 129303420 | uncharacterized LOC129303420 | 3.0 | 4.64E-05 |
| LOC129306604 | 129306604 | probable aspartic proteinase GIP2 | 3.0 | 9.24E-03 |
| LOC129308462 | 129308462 | uncharacterized LOC129308462 | 3.0 | 1.63E-02 |
| LOC129286994 | 129286994 | uncharacterized LOC129286994 | 3.0 | 2.48E-04 |
| LOC129320179 | 129320179 | uncharacterized LOC129320179 | 3.0 | 1.67E-04 |
| LOC129291805 | 129291805 | cytosolic sulfotransferase 12-like | 3.0 | 5.85E-03 |
| LOC129303676 | 129303676 | E3 ubiquitin-protein ligase At4g11680-like | 3.0 | 2.58E-06 |
| LOC129311899 | 129311899 | nudix hydrolase 17, mitochondrial-like | 3.0 | 2.27E-03 |
| LOC129305708 | 129305708 | actin-depolymerizing factor | 3.0 | 6.00E-07 |
| LOC129294883 | 129294883 | NADH-ubiquinone oxidoreductase chain 3 | 3.0 | 1.25E-02 |

| | | | | |
|---|---|---|---|---|
| LOC129295487 | 129295487 | chaperone protein dnaJ 10-like | 3.0 | 4.35E-03 |
| LOC129284862 | 129284862 | disease resistance protein SUMM2-like | 3.0 | 1.11E-03 |
| LOC129303086 | 129303086 | probable aquaporin PIP-type 7a | 3.0 | 4.14E-08 |
| LOC129290804 | 129290804 | reticulon-like protein B1 | 3.0 | 2.86E-20 |
| LOC129284382 | 129284382 | uncharacterized LOC129284382 | 2.9 | 3.92E-10 |
| LOC129309940 | 129309940 | uncharacterized LOC129309940 | 2.9 | 3.06E-02 |
| LOC129301684 | 129301684 | uncharacterized LOC129301684 | 2.9 | 3.17E-08 |
| LOC129302063 | 129302063 | putative lipid-transfer protein DIR1 | 2.9 | 1.54E-02 |
| LOC129298450 | 129298450 | uncharacterized LOC129298450 | 2.9 | 1.34E-02 |
| LOC129294752 | 129294752 | uncharacterized LOC129294752 | 2.9 | 6.93E-05 |
| LOC129291878 | 129291878 | histidine-containing phosphotransfer protein 4 | 2.9 | 2.89E-02 |
| LOC129317670 | 129317670 | CBL-interacting serine/threonine-protein kinase 11 | 2.9 | 3.50E-08 |
| LOC129285842 | 129285842 | ATP-dependent zinc metalloprotease FTSH, chloroplastic-like | 2.9 | 1.44E-05 |
| LOC129309435 | 129309435 | uncharacterized LOC129309435 | 2.9 | 2.26E-03 |
| LOC129284988 | 129284988 | auxin transporter-like protein 5 | 2.9 | 2.29E-03 |
| LOC129302034 | 129302034 | glutaredoxin-C10-like | 2.9 | 9.50E-03 |
| LOC129308477 | 129308477 | indole-3-acetic acid-induced protein ARG2-like | 2.9 | 6.61E-06 |
| LOC129294846 | 129294846 | putative ATP synthase protein YMF19 | 2.9 | 1.47E-02 |
| LOC129312880 | 129312880 | septum-promoting GTP-binding protein 1 | 2.9 | 3.92E-06 |
| LOC129297770 | 129297770 | ATP synthase subunit a, chloroplastic | 2.9 | 1.32E-02 |
| LOC129287146 | 129287146 | potassium channel AKT2/3-like | 2.9 | 2.97E-02 |
| LOC129291086 | 129291086 | uncharacterized LOC129291086 | 2.9 | 2.21E-03 |
| LOC129314253 | 129314253 | uncharacterized LOC129314253 | 2.9 | 4.63E-02 |
| LOC129287842 | 129287842 | uncharacterized LOC129287842 | 2.9 | 1.74E-02 |
| LOC129312475 | 129312475 | uncharacterized LOC129312475 | 2.9 | 3.96E-02 |
| LOC129300418 | 129300418 | uncharacterized LOC129300418 | 2.9 | 1.22E-04 |
| LOC129291441 | 129291441 | gibberellin receptor GID1B-like | 2.9 | 6.54E-06 |
| LOC129309367 | 129309367 | chaperone protein ClpD, chloroplastic | 2.9 | 1.03E-02 |
| LOC129293961 | 129293961 | uncharacterized LOC129293961 | 2.9 | 5.47E-10 |
| LOC129304247 | 129304247 | cinnamoyl-CoA reductase CAD2-like | 2.9 | 1.50E-04 |
| LOC129304139 | 129304139 | kinesin-like protein KIN-6 | 2.9 | 8.50E-05 |
| LOC129303598 | 129303598 | alpha-amylase-like | 2.9 | 5.85E-05 |
| LOC129304506 | 129304506 | NAC domain-containing protein 87-like | 2.9 | 6.34E-08 |
| LOC129315938 | 129315938 | uncharacterized LOC129315938 | 2.9 | 2.88E-02 |
| LOC129317802 | 129317802 | ABC transporter B family member 4-like | 2.9 | 2.61E-04 |
| LOC129314477 | 129314477 | glutaredoxin-C9-like | 2.9 | 9.03E-05 |
| LOC129299739 | 129299739 | probable CoA ligase CCL5 | 2.9 | 2.13E-04 |
| LOC129309928 | 129309928 | probable alpha,alpha-trehalose-phosphate synthase [UDP-formir | 2.9 | 3.53E-09 |
| LOC129294840 | 129294840 | ribosomal protein S3, mitochondrial-like | 2.9 | 1.86E-05 |
| LOC129285804 | 129285804 | uncharacterized LOC129285804 | 2.9 | 1.54E-09 |
| LOC129304438 | 129304438 | NAC domain-containing protein 1-like | 2.9 | 1.15E-05 |
| LOC129316040 | 129316040 | probable carboxylesterase 6 | 2.9 | 1.32E-03 |
| LOC129292632 | 129292632 | uncharacterized LOC129292632 | 2.9 | 1.29E-03 |
| LOC129318720 | 129318720 | ethylene-responsive transcription factor RAP2-1-like | 2.9 | 6.75E-04 |
| LOC129304198 | 129304198 | putative receptor protein kinase ZmPK1 | 2.9 | 1.16E-06 |
| LOC129313178 | 129313178 | 65-kDa microtubule-associated protein 3-like | 2.9 | 3.40E-02 |
| LOC129307026 | 129307026 | probable cytochrome c oxidase subunit 5C-3 | 2.9 | 2.44E-07 |
| LOC129320029 | 129320029 | alkaline/neutral invertase A, mitochondrial-like | 2.9 | 5.57E-06 |

| | | | | |
|---|---|---|---|---|
| LOC129284827 | 129284827 | receptor-like protein kinase 7 | 2.9 | 6.95E-09 |
| LOC129294801 | 129294801 | molybdate transporter 1-like | 2.9 | 9.43E-05 |
| LOC129319756 | 129319756 | trihelix transcription factor ASR3-like | 2.9 | 3.33E-03 |
| LOC129294587 | 129294587 | uncharacterized LOC129294587 | 2.9 | 7.90E-09 |
| LOC129303899 | 129303899 | uncharacterized LOC129303899 | 2.9 | 6.24E-03 |
| LOC129285224 | 129285224 | polyol transporter 5-like | 2.9 | 2.93E-03 |
| LOC129313297 | 129313297 | uncharacterized LOC129313297 | 2.9 | 1.78E-02 |
| LOC129323186 | 129323186 | methyltransferase FGSG_00040 | 2.9 | 2.96E-02 |
| LOC129303916 | 129303916 | zinc finger protein 1 | 2.9 | 7.39E-03 |
| LOC129315451 | 129315451 | uncharacterized LOC129315451 | 2.8 | 4.13E-06 |
| LOC129300468 | 129300468 | transcription factor MYB3R-1-like | 2.8 | 9.23E-03 |
| LOC129315034 | 129315034 | uncharacterized LOC129315034 | 2.8 | 1.88E-04 |
| LOC129311429 | 129311429 | putative pentatricopeptide repeat-containing protein At3g23330 | 2.8 | 2.82E-06 |
| LOC129293479 | 129293479 | heavy metal-associated isoprenylated plant protein 26 | 2.8 | 1.14E-06 |
| LOC129286713 | 129286713 | uncharacterized LOC129286713 | 2.8 | 5.19E-03 |
| LOC129295081 | 129295081 | uncharacterized LOC129295081 | 2.8 | 1.53E-05 |
| LOC129323148 | 129323148 | small heat shock protein, chloroplastic-like | 2.8 | 3.08E-05 |
| LOC129296545 | 129296545 | U12 minor spliceosomal RNA | 2.8 | 2.18E-03 |
| LOC129286689 | 129286689 | glucan endo-1,3-beta-glucosidase 5-like | 2.8 | 2.17E-04 |
| LOC129284716 | 129284716 | VAN3-binding protein-like | 2.8 | 4.54E-02 |
| LOC129310014 | 129310014 | serine/threonine-protein kinase RUNKEL | 2.8 | 4.42E-03 |
| LOC129288686 | 129288686 | uncharacterized LOC129288686 | 2.8 | 3.14E-02 |
| LOC129297942 | 129297942 | classical arabinogalactan protein 1-like | 2.8 | 2.61E-02 |
| LOC129321351 | 129321351 | protein NRT1/ PTR FAMILY 3.1-like | 2.8 | 3.82E-04 |
| LOC129296075 | 129296075 | geranylgeranyl pyrophosphate synthase, chloroplastic-like | 2.8 | 1.69E-03 |
| LOC129291531 | 129291531 | vestitone reductase-like | 2.8 | 4.12E-03 |
| LOC129299163 | 129299163 | uncharacterized LOC129299163 | 2.8 | 3.62E-02 |
| LOC129304566 | 129304566 | external alternative NAD(P)H-ubiquinone oxidoreductase B2, m | 2.8 | 7.65E-04 |
| LOC129322636 | 129322636 | ACT domain-containing protein ACR8-like | 2.8 | 1.35E-06 |
| LOC129292511 | 129292511 | protein ASPARTIC PROTEASE IN GUARD CELL 1-like | 2.8 | 5.15E-09 |
| LOC129284358 | 129284358 | uncharacterized LOC129284358 | 2.8 | 1.29E-02 |
| LOC129322611 | 129322611 | MLP-like protein 34 | 2.8 | 1.43E-02 |
| LOC129298741 | 129298741 | NAC domain-containing protein 83-like | 2.8 | 8.03E-04 |
| LOC129285001 | 129285001 | cytochrome b561 and DOMON domain-containing protein At4g | 2.8 | 3.08E-03 |
| LOC129292698 | 129292698 | protein SRC1-like | 2.8 | 3.27E-05 |
| LOC129288076 | 129288076 | uncharacterized LOC129288076 | 2.8 | 1.03E-02 |
| LOC129286951 | 129286951 | disease resistance protein RUN1-like | 2.8 | 4.04E-05 |
| LOC129303778 | 129303778 | transcription factor MYB20-like | 2.8 | 3.10E-05 |
| LOC129321642 | 129321642 | kinesin-like protein NACK1 | 2.8 | 5.65E-03 |
| LOC129323163 | 129323163 | microtubule-associated protein RP/EB family member 1C | 2.8 | 3.51E-02 |
| LOC129306500 | 129306500 | thioredoxin-like protein YLS8 | 2.8 | 3.02E-03 |
| LOC129320731 | 129320731 | TSL-kinase interacting protein 1 | 2.8 | 7.81E-03 |
| LOC129295366 | 129295366 | temperature-induced lipocalin-1-like | 2.8 | 2.07E-07 |
| LOC129293192 | 129293192 | tetraspanin-2-like | 2.8 | 4.45E-02 |
| LOC129299515 | 129299515 | uncharacterized LOC129299515 | 2.8 | 9.00E-03 |
| LOC129320028 | 129320028 | proline-rich receptor-like protein kinase PERK10 | 2.8 | 2.23E-04 |
| LOC129293663 | 129293663 | uncharacterized LOC129293663 | 2.8 | 9.81E-07 |
| LOC129294264 | 129294264 | protein kinase STUNTED-like | 2.8 | 5.19E-08 |

| | | | | |
|---|---|---|---|---|
| LOC129316003 | 129316003 | gibberellin 2-beta-dioxygenase 1 | 2.8 | 1.89E-07 |
| LOC129315840 | 129315840 | protein GRAVITROPIC IN THE LIGHT 1 | 2.8 | 2.11E-08 |
| LOC129301698 | 129301698 | fimbrin-1-like | 2.8 | 3.01E-03 |
| LOC129300569 | 129300569 | transcription repressor OFP13-like | 2.8 | 2.83E-02 |
| LOC129311871 | 129311871 | pectinesterase | 2.8 | 3.01E-04 |
| LOC129291534 | 129291534 | auxin-responsive protein SAUR32 | 2.8 | 1.47E-04 |
| LOC129316114 | 129316114 | DNA-directed RNA polymerases II, IV and V subunit 12-like | 2.8 | 8.87E-04 |
| LOC129315559 | 129315559 | non-specific lipid transfer protein GPI-anchored 14-like | 2.8 | 2.02E-02 |
| LOC129288804 | 129288804 | auxin-responsive protein IAA17-like | 2.8 | 7.19E-10 |
| LOC129309736 | 129309736 | uncharacterized LOC129309736 | 2.8 | 1.57E-02 |
| LOC129303904 | 129303904 | protein DETOXIFICATION 48-like | 2.8 | 1.93E-02 |
| LOC129315864 | 129315864 | myb-related protein 2-like | 2.8 | 1.51E-02 |
| LOC129312699 | 129312699 | acid phosphatase 1-like | 2.7 | 1.93E-06 |
| LOC129290402 | 129290402 | uncharacterized LOC129290402 | 2.7 | 1.83E-09 |
| LOC129319575 | 129319575 | chorismate synthase, chloroplastic-like | 2.7 | 1.77E-02 |
| LOC129312611 | 129312611 | FT-interacting protein 3 | 2.7 | 2.07E-03 |
| LOC129306625 | 129306625 | universal stress protein PHOS34-like | 2.7 | 1.98E-11 |
| LOC129315923 | 129315923 | zinc finger A20 and AN1 domain-containing stress-associated p | 2.7 | 9.37E-06 |
| LOC129310006 | 129310006 | auxin-induced protein 6B-like | 2.7 | 1.27E-03 |
| LOC129305143 | 129305143 | probable protein phosphatase 2C 38 | 2.7 | 1.31E-03 |
| LOC129298911 | 129298911 | nematode resistance protein-like HSPRO2 | 2.7 | 7.60E-09 |
| LOC129306441 | 129306441 | transcription factor bHLH147-like | 2.7 | 7.82E-05 |
| LOC129294142 | 129294142 | dehydration-responsive element-binding protein 2C-like | 2.7 | 1.38E-05 |
| LOC129321530 | 129321530 | cation/calcium exchanger 2-like | 2.7 | 1.75E-13 |
| LOC129323080 | 129323080 | transcription factor LHW | 2.7 | 3.36E-03 |
| LOC129322184 | 129322184 | protein IQ-DOMAIN 20 | 2.7 | 2.11E-02 |
| LOC129320446 | 129320446 | heat shock factor protein HSF30-like | 2.7 | 1.40E-04 |
| LOC129306139 | 129306139 | uncharacterized LOC129306139 | 2.7 | 1.63E-03 |
| LOC129302237 | 129302237 | uncharacterized LOC129302237 | 2.7 | 1.47E-04 |
| LOC129295511 | 129295511 | transcription factor bHLH93-like | 2.7 | 6.73E-05 |
| LOC129301629 | 129301629 | NAC domain-containing protein 2-like | 2.7 | 1.43E-06 |
| LOC129295384 | 129295384 | uncharacterized LOC129295384 | 2.7 | 1.27E-06 |
| LOC129304038 | 129304038 | probable receptor-like protein kinase At5g38990 | 2.7 | 3.95E-03 |
| LOC129311566 | 129311566 | uncharacterized LOC129311566 | 2.7 | 5.84E-12 |
| LOC129290233 | 129290233 | uncharacterized LOC129290233 | 2.7 | 2.89E-10 |
| LOC129314505 | 129314505 | probable aminotransferase TAT2 | 2.7 | 3.10E-03 |
| LOC129291657 | 129291657 | uncharacterized LOC129291657 | 2.7 | 6.65E-03 |
| LOC129306724 | 129306724 | syntaxin-121-like | 2.7 | 3.68E-02 |
| LOC129315833 | 129315833 | nuclear transcription factor Y subunit C-2-like | 2.7 | 2.45E-05 |
| LOC129310290 | 129310290 | LOB domain-containing protein 40-like | 2.7 | 3.06E-04 |
| LOC129291369 | 129291369 | meiotic nuclear division protein 1 homolog | 2.7 | 1.69E-02 |
| LOC129321616 | 129321616 | TORTIFOLIA1-like protein 4 | 2.7 | 1.09E-04 |
| LOC129312596 | 129312596 | 65-kDa microtubule-associated protein 5 | 2.7 | 5.33E-03 |
| LOC129315148 | 129315148 | uncharacterized LOC129315148 | 2.7 | 7.14E-16 |
| LOC129294584 | 129294584 | protein ZW2-like | 2.7 | 1.81E-04 |
| LOC129318968 | 129318968 | peptide methionine sulfoxide reductase-like | 2.7 | 1.58E-02 |
| LOC129312147 | 129312147 | nudix hydrolase 1-like | 2.7 | 6.16E-03 |
| LOC129310547 | 129310547 | lysine histidine transporter-like 6 | 2.7 | 3.47E-06 |

| | | | | |
|---|---|---|---|---|
| LOC129285303 | 129285303 | ABSCISIC ACID-INSENSITIVE 5-like protein 7 | 2.7 | 1.39E-12 |
| LOC129312602 | 129312602 | indole-3-acetic acid-amido synthetase GH3.6-like | 2.7 | 3.86E-02 |
| LOC129307688 | 129307688 | KIN14B-interacting protein At4g14310 | 2.7 | 1.06E-02 |
| LOC129322781 | 129322781 | expansin-A10-like | 2.7 | 3.33E-07 |
| LOC129318404 | 129318404 | CBL-interacting serine/threonine-protein kinase 10-like | 2.7 | 5.72E-10 |
| LOC129309424 | 129309424 | uncharacterized LOC129309424 | 2.7 | 1.59E-03 |
| LOC129318733 | 129318733 | uncharacterized LOC129318733 | 2.7 | 2.61E-04 |
| LOC129305051 | 129305051 | bifunctional 3-dehydroquinate dehydratase/shikimate dehydroge | 2.7 | 4.17E-02 |
| LOC129295349 | 129295349 | uncharacterized LOC129295349 | 2.7 | 2.40E-03 |
| LOC129299559 | 129299559 | E3 ubiquitin-protein ligase SGR9, amyloplastic-like | 2.7 | 3.03E-05 |
| LOC129308417 | 129308417 | 1-aminocyclopropane-1-carboxylate oxidase 1 | 2.7 | 1.17E-04 |
| LOC129303261 | 129303261 | probable protein phosphatase 2C 25 | 2.7 | 1.18E-03 |
| LOC129303038 | 129303038 | cysteine-rich receptor-like protein kinase 2 | 2.7 | 4.29E-02 |
| LOC129307210 | 129307210 | uncharacterized LOC129307210 | 2.7 | 2.35E-02 |
| LOC129322667 | 129322667 | phosphatidylinositol/phosphatidylcholine transfer protein SFH12 | 2.7 | 1.65E-02 |
| LOC129309733 | 129309733 | uncharacterized LOC129309733 | 2.7 | 4.91E-04 |
| LOC129307364 | 129307364 | uncharacterized LOC129307364 | 2.7 | 8.29E-03 |
| LOC129318432 | 129318432 | WRKY transcription factor 71-like | 2.7 | 2.13E-04 |
| LOC129322725 | 129322725 | probable xyloglucan galactosyltransferase GT17 | 2.6 | 2.12E-03 |
| LOC129294394 | 129294394 | uncharacterized WD repeat-containing protein C2A9.03 | 2.6 | 2.04E-05 |
| LOC129304233 | 129304233 | RPM1 interacting protein 13-like | 2.6 | 4.14E-03 |
| LOC129291318 | 129291318 | 18.1 kDa class I heat shock protein-like | 2.6 | 4.96E-03 |
| LOC129303713 | 129303713 | uncharacterized LOC129303713 | 2.6 | 9.41E-05 |
| LOC129294837 | 129294837 | NADH-ubiquinone oxidoreductase chain 5-like | 2.6 | 3.39E-04 |
| LOC129290945 | 129290945 | allene oxide synthase-like | 2.6 | 2.56E-02 |
| LOC129311804 | 129311804 | ethylene-responsive transcription factor ERF011-like | 2.6 | 4.14E-07 |
| LOC129300004 | 129300004 | uncharacterized LOC129300004 | 2.6 | 1.63E-02 |
| LOC129311750 | 129311750 | D-aminoacyl-tRNA deacylase-like | 2.6 | 8.65E-03 |
| LOC129307236 | 129307236 | uncharacterized LOC129307236 | 2.6 | 2.96E-02 |
| LOC129321763 | 129321763 | homeobox-leucine zipper protein HAT22-like | 2.6 | 2.71E-05 |
| LOC129296263 | 129296263 | protein GIGANTEA-like | 2.6 | 3.26E-02 |
| LOC129303110 | 129303110 | amino acid permease 6-like | 2.6 | 1.16E-03 |
| LOC129306555 | 129306555 | protein WVD2-like 7 | 2.6 | 4.57E-02 |
| LOC129302908 | 129302908 | uncharacterized LOC129302908 | 2.6 | 3.84E-04 |
| LOC129284466 | 129284466 | cyclin-dependent protein kinase inhibitor SMR8-like | 2.6 | 7.43E-05 |
| LOC129318495 | 129318495 | polyubiquitin | 2.6 | 1.41E-05 |
| LOC129294849 | 129294849 | cytochrome b | 2.6 | 3.36E-02 |
| LOC129313478 | 129313478 | uncharacterized LOC129313478 | 2.6 | 2.54E-04 |
| LOC129303739 | 129303739 | G2/mitotic-specific cyclin S13-7-like | 2.6 | 2.04E-02 |
| LOC129309616 | 129309616 | uncharacterized LOC129309616 | 2.6 | 1.16E-02 |
| LOC129311332 | 129311332 | purple acid phosphatase 3 | 2.6 | 2.17E-03 |
| LOC129320744 | 129320744 | uncharacterized LOC129320744 | 2.6 | 2.47E-04 |
| LOC129293939 | 129293939 | probable chlorophyll(ide) b reductase NYC1, chloroplastic | 2.6 | 2.57E-06 |
| LOC129311621 | 129311621 | zinc-finger homeodomain protein 11-like | 2.6 | 1.88E-13 |
| LOC129316414 | 129316414 | uncharacterized LOC129316414 | 2.6 | 9.15E-03 |
| LOC129293697 | 129293697 | outer envelope pore protein 16-2, chloroplastic-like | 2.6 | 1.13E-06 |
| LOC129308824 | 129308824 | pathogen-associated molecular patterns-induced protein A70-lik | 2.6 | 9.00E-03 |
| LOC129306599 | 129306599 | structural maintenance of chromosomes protein 4 | 2.6 | 1.62E-03 |

| | | | | |
|---|---|---|---|---|
| LOC129285221 | 129285221 | ATP-dependent DNA helicase Q-like 4A | 2.6 | 2.50E-02 |
| LOC129319888 | 129319888 | ethylene-responsive transcription factor ERF110-like | 2.6 | 3.32E-03 |
| LOC129285125 | 129285125 | abscisic acid 8'-hydroxylase 4-like | 2.6 | 3.74E-02 |
| LOC129313705 | 129313705 | dynamin-related protein 5A | 2.6 | 2.24E-03 |
| LOC129310675 | 129310675 | uncharacterized LOC129310675 | 2.6 | 1.12E-03 |
| LOC129302141 | 129302141 | F-box protein GID2-like | 2.6 | 2.04E-05 |
| LOC129294884 | 129294884 | ribosomal protein S12, mitochondrial | 2.6 | 2.12E-02 |
| LOC129309274 | 129309274 | uncharacterized LOC129309274 | 2.6 | 2.01E-02 |
| LOC129314341 | 129314341 | probable inactive purple acid phosphatase 27 | 2.6 | 1.69E-02 |
| LOC129305969 | 129305969 | cytokinin dehydrogenase 9 | 2.6 | 3.05E-02 |
| LOC129317030 | 129317030 | uncharacterized LOC129317030 | 2.6 | 1.06E-02 |
| LOC129291195 | 129291195 | NADPH-dependent aldo-keto reductase, chloroplastic-like | 2.6 | 1.26E-05 |
| LOC129288947 | 129288947 | uncharacterized LOC129288947 | 2.6 | 5.58E-03 |
| LOC129313552 | 129313552 | uncharacterized LOC129313552 | 2.6 | 5.41E-04 |
| LOC129296222 | 129296222 | metallothionein-like protein 2 | 2.6 | 3.65E-03 |
| LOC129317633 | 129317633 | U-box domain-containing protein 4 | 2.6 | 9.86E-04 |
| LOC129293613 | 129293613 | zinc finger protein ZAT12-like | 2.6 | 2.47E-02 |
| LOC129312750 | 129312750 | aspartic proteinase NANA, chloroplast | 2.6 | 3.13E-06 |
| LOC129301877 | 129301877 | serine/threonine-protein kinase MPS1 | 2.6 | 4.74E-02 |
| LOC129306859 | 129306859 | uncharacterized LOC129306859 | 2.6 | 2.18E-08 |
| LOC129318824 | 129318824 | protein REVERSION-TO-ETHYLENE SENSITIVITY1 | 2.6 | 3.03E-06 |
| LOC129304932 | 129304932 | ninja-family protein AFP3 | 2.6 | 3.63E-07 |
| LOC129308787 | 129308787 | ethylene receptor 2 | 2.6 | 3.64E-06 |
| LOC129323012 | 129323012 | PLASMODESMATA CALLOSE-BINDING PROTEIN 3-like | 2.6 | 4.35E-03 |
| LOC129311363 | 129311363 | uncharacterized LOC129311363 | 2.6 | 3.36E-05 |
| LOC129315822 | 129315822 | peptide methionine sulfoxide reductase B1, chloroplastic | 2.6 | 3.71E-04 |
| LOC129321507 | 129321507 | phytosulfokine receptor 2 | 2.6 | 6.25E-05 |
| LOC129294839 | 129294839 | ATP synthase subunit a | 2.6 | 1.55E-02 |
| LOC129298667 | 129298667 | uncharacterized LOC129298667 | 2.6 | 9.83E-05 |
| LOC129310665 | 129310665 | uncharacterized LOC129310665 | 2.6 | 4.54E-03 |
| LOC129313291 | 129313291 | kinesin-like protein KIN-14Q | 2.6 | 3.06E-02 |
| LOC129294570 | 129294570 | lysine-rich arabinogalactan protein 19-like | 2.6 | 2.77E-03 |
| LOC129321499 | 129321499 | cysteine-rich receptor-like protein kinase 43 | 2.6 | 1.43E-02 |
| LOC129299176 | 129299176 | protein SMALL AUXIN UP-REGULATED RNA 12-like | 2.6 | 3.71E-02 |
| LOC129303782 | 129303782 | adenine phosphoribosyltransferase 5-like | 2.6 | 4.34E-02 |
| LOC129293080 | 129293080 | oxalate--CoA ligase | 2.6 | 7.14E-03 |
| LOC129320828 | 129320828 | uncharacterized LOC129320828 | 2.6 | 1.34E-03 |
| LOC129322202 | 129322202 | ABC transporter G family member 21-like | 2.6 | 2.58E-05 |
| LOC129311818 | 129311818 | uncharacterized protein At4g22758 | 2.6 | 2.88E-05 |
| LOC129303369 | 129303369 | cryptochrome DASH, chloroplastic/mitochondrial | 2.6 | 2.03E-05 |
| LOC129322915 | 129322915 | protein TIFY 10b-like | 2.6 | 9.94E-09 |
| LOC129292720 | 129292720 | F-box protein At4g00755-like | 2.6 | 6.85E-03 |
| LOC129286929 | 129286929 | xyloglucan galactosyltransferase MUR3-like | 2.6 | 6.13E-06 |
| LOC129314424 | 129314424 | cytochrome c oxidase copper chaperone 1-like | 2.6 | 1.02E-03 |
| LOC129295642 | 129295642 | ubiquitin-conjugating enzyme E2 22-like | 2.6 | 9.95E-04 |
| LOC129311837 | 129311837 | LEAF RUST 10 DISEASE-RESISTANCE LOCUS RECEPTOI | 2.6 | 9.43E-05 |
| LOC129305928 | 129305928 | ethylene-responsive transcription factor ABR1-like | 2.5 | 4.41E-03 |
| LOC129309890 | 129309890 | DNA repair protein RAD51 homolog | 2.5 | 2.23E-02 |

| | | | | |
|---|---|---|---|---|
| LOC129305727 | 129305727 | monothiol glutaredoxin-S6-like | 2.5 | 1.74E-02 |
| LOC129308593 | 129308593 | protein NETWORKED 3A-like | 2.5 | 4.37E-02 |
| LOC129292051 | 129292051 | uncharacterized LOC129292051 | 2.5 | 3.15E-02 |
| LOC129303559 | 129303559 | F-box/kelch-repeat protein At5g26960-like | 2.5 | 3.20E-03 |
| LOC129286106 | 129286106 | uncharacterized LOC129286106 | 2.5 | 1.94E-02 |
| LOC129309503 | 129309503 | E2F transcription factor-like E2FE | 2.5 | 1.13E-02 |
| LOC129314209 | 129314209 | protein FAF-like, chloroplastic | 2.5 | 9.88E-04 |
| LOC129311905 | 129311905 | zinc finger protein GIS3-like | 2.5 | 5.53E-03 |
| LOC129299019 | 129299019 | uncharacterized calcium-binding protein At1g02270-like | 2.5 | 5.88E-03 |
| LOC129310543 | 129310543 | probable protein phosphatase 2C 49 | 2.5 | 1.56E-08 |
| LOC129306077 | 129306077 | cytochrome P450 94B3-like | 2.5 | 1.96E-02 |
| LOC129306395 | 129306395 | cytochrome P450 94A2-like | 2.5 | 4.85E-04 |
| LOC129305243 | 129305243 | xylan glycosyltransferase MUCI21-like | 2.5 | 1.06E-03 |
| LOC129318489 | 129318489 | putative cyclin-A3-1 | 2.5 | 2.29E-03 |
| LOC129311933 | 129311933 | uncharacterized LOC129311933 | 2.5 | 2.68E-03 |
| LOC129289472 | 129289472 | uncharacterized LOC129289472 | 2.5 | 3.38E-03 |
| LOC129289459 | 129289459 | axial regulator YABBY 1 | 2.5 | 2.99E-02 |
| LOC129316765 | 129316765 | uncharacterized LOC129316765 | 2.5 | 1.45E-04 |
| LOC129303851 | 129303851 | cytochrome P450 82A3-like | 2.5 | 2.29E-04 |
| LOC129301525 | 129301525 | polyamine oxidase 2-like | 2.5 | 1.02E-03 |
| LOC129293022 | 129293022 | uncharacterized LOC129293022 | 2.5 | 1.21E-08 |
| LOC129285888 | 129285888 | glutaredoxin-C11-like | 2.5 | 5.10E-03 |
| LOC129289612 | 129289612 | mitogen-activated protein kinase kinase 5-like | 2.5 | 7.80E-10 |
| LOC129300785 | 129300785 | uncharacterized LOC129300785 | 2.5 | 3.98E-02 |
| LOC129292598 | 129292598 | uncharacterized LOC129292598 | 2.5 | 4.42E-02 |
| LOC129291905 | 129291905 | histone H1-like | 2.5 | 1.34E-02 |
| LOC129310605 | 129310605 | LEAF RUST 10 DISEASE-RESISTANCE LOCUS RECEPTO | 2.5 | 1.62E-03 |
| LOC129303508 | 129303508 | pirin-like protein | 2.5 | 6.88E-04 |
| LOC129294741 | 129294741 | nitrate regulatory gene2 protein | 2.5 | 2.39E-02 |
| LOC129313661 | 129313661 | uncharacterized LOC129313661 | 2.5 | 1.05E-02 |
| LOC129321952 | 129321952 | iridoid oxidase | 2.5 | 4.24E-03 |
| LOC129300743 | 129300743 | nuclear envelope-associated protein 2-like | 2.5 | 1.48E-02 |
| LOC129308737 | 129308737 | inactive beta-amylase 9 | 2.5 | 4.34E-05 |
| LOC129318718 | 129318718 | uncharacterized LOC129318718 | 2.5 | 3.58E-04 |
| LOC129311987 | 129311987 | defensin-like protein 2 | 2.5 | 1.55E-02 |
| LOC129312948 | 129312948 | ras-related protein RABA4d-like | 2.5 | 2.69E-07 |
| LOC129292834 | 129292834 | heat shock protein 83 | 2.5 | 1.07E-04 |
| LOC129300330 | 129300330 | receptor-like serine/threonine-protein kinase At2g45590 | 2.5 | 3.84E-03 |
| LOC129298495 | 129298495 | protein SUPPRESSOR OF MAX2 1-like | 2.5 | 1.32E-02 |
| LOC129294905 | 129294905 | uncharacterized mitochondrial protein ymf1-like | 2.5 | 3.45E-02 |
| LOC129295315 | 129295315 | uncharacterized LOC129295315 | 2.5 | 1.94E-02 |
| LOC129303246 | 129303246 | uridine nucleosidase 1 | 2.5 | 7.78E-07 |
| LOC129321458 | 129321458 | ethylene response sensor 1-like | 2.5 | 2.30E-04 |
| LOC129294881 | 129294881 | NADH-ubiquinone oxidoreductase chain 2 | 2.5 | 1.50E-04 |
| LOC129314792 | 129314792 | probable protein phosphatase 2C 58 | 2.5 | 2.30E-05 |
| LOC129287218 | 129287218 | BAG family molecular chaperone regulator 2-like | 2.5 | 9.56E-04 |
| LOC129287539 | 129287539 | BTB/POZ domain-containing protein At3g08570 | 2.5 | 1.84E-02 |
| LOC129301733 | 129301733 | probable receptor-like protein kinase At5g38990 | 2.5 | 1.51E-02 |

| | | | | |
|---|---|---|---|---|
| LOC129303419 | 129303419 | long chain base biosynthesis protein 1-like | 2.5 | 4.23E-02 |
| LOC129294382 | 129294382 | receptor-like protein kinase 7 | 2.5 | 4.19E-04 |
| LOC129305853 | 129305853 | fra a 1-associated protein | 2.5 | 5.24E-05 |
| LOC129309529 | 129309529 | uncharacterized LOC129309529 | 2.5 | 4.84E-02 |
| LOC129300782 | 129300782 | PRA1 family protein F2-like | 2.5 | 3.48E-04 |
| LOC129286609 | 129286609 | uncharacterized LOC129286609 | 2.5 | 1.06E-05 |
| LOC129317285 | 129317285 | L-ascorbate peroxidase 2, cytosolic | 2.5 | 2.32E-02 |
| LOC129291205 | 129291205 | extradiol ring-cleavage dioxygenase-like | 2.5 | 1.49E-03 |
| LOC129310495 | 129310495 | galactomannan galactosyltransferase 1-like | 2.5 | 1.15E-06 |
| LOC129311251 | 129311251 | leucoanthocyanidin reductase-like | 2.5 | 2.43E-04 |
| LOC129285390 | 129285390 | putative disease resistance RPP13-like protein 3 | 2.5 | 3.99E-02 |
| LOC129321723 | 129321723 | uncharacterized LOC129321723 | 2.5 | 5.50E-03 |
| LOC129290614 | 129290614 | uncharacterized LOC129290614 | 2.4 | 5.31E-03 |
| LOC129308664 | 129308664 | guanine nucleotide-binding protein subunit gamma 3-like | 2.4 | 1.04E-02 |
| LOC129293929 | 129293929 | LEAF RUST 10 DISEASE-RESISTANCE LOCUS RECEPTO | 2.4 | 4.39E-02 |
| LOC129310276 | 129310276 | scarecrow-like protein 28 | 2.4 | 9.62E-03 |
| LOC129297666 | 129297666 | DNA repair protein XRCC3 homolog | 2.4 | 1.63E-03 |
| LOC129299983 | 129299983 | transcription factor GTE4-like | 2.4 | 2.70E-02 |
| LOC129311211 | 129311211 | uncharacterized LOC129311211 | 2.4 | 4.70E-03 |
| LOC129296434 | 129296434 | E3 ubiquitin-protein ligase RING1-like | 2.4 | 9.57E-08 |
| LOC129297461 | 129297461 | uncharacterized LOC129297461 | 2.4 | 1.10E-03 |
| LOC129288488 | 129288488 | G-box-binding factor 3-like | 2.4 | 2.62E-07 |
| LOC129297587 | 129297587 | uncharacterized LOC129297587 | 2.4 | 4.93E-06 |
| LOC129303610 | 129303610 | ras-related protein RABA6b-like | 2.4 | 6.57E-04 |
| LOC129323168 | 129323168 | uncharacterized LOC129323168 | 2.4 | 4.18E-03 |
| LOC129306863 | 129306863 | uncharacterized LOC129306863 | 2.4 | 2.18E-02 |
| LOC129303680 | 129303680 | U-box domain-containing protein 4 | 2.4 | 3.16E-10 |
| LOC129322176 | 129322176 | uncharacterized LOC129322176 | 2.4 | 6.55E-05 |
| LOC129286774 | 129286774 | uncharacterized LOC129286774 | 2.4 | 4.81E-09 |
| LOC129309762 | 129309762 | probable arabinosyltransferase ARAD1 | 2.4 | 4.82E-02 |
| LOC129303818 | 129303818 | rho GDP-dissociation inhibitor 1-like | 2.4 | 2.45E-03 |
| LOC129318957 | 129318957 | uncharacterized LOC129318957 | 2.4 | 1.44E-02 |
| LOC129311067 | 129311067 | thioredoxin H2-like | 2.4 | 4.02E-03 |
| LOC129290488 | 129290488 | PRA1 family protein B4-like | 2.4 | 6.77E-05 |
| LOC129294847 | 129294847 | NADH-ubiquinone oxidoreductase chain 1 | 2.4 | 3.86E-02 |
| LOC129321858 | 129321858 | probable WRKY transcription factor 4 | 2.4 | 1.21E-03 |
| LOC129285662 | 129285662 | transcription factor MYB14-like | 2.4 | 2.77E-02 |
| LOC129295655 | 129295655 | uncharacterized protein At3g61260-like | 2.4 | 1.86E-04 |
| LOC129294689 | 129294689 | uncharacterized protein At5g65660-like | 2.4 | 2.29E-04 |
| LOC129285769 | 129285769 | phenylcoumaran benzylic ether reductase POP1-like | 2.4 | 3.57E-03 |
| LOC129286545 | 129286545 | uncharacterized LOC129286545 | 2.4 | 8.90E-04 |
| LOC129284416 | 129284416 | probable calcium-binding protein CML25 | 2.4 | 1.86E-02 |
| LOC129288180 | 129288180 | B3 domain-containing transcription factor NGA1-like | 2.4 | 8.07E-11 |
| LOC129306311 | 129306311 | iron-sulfur assembly protein IscA-like 1, mitochondrial | 2.4 | 4.38E-04 |
| LOC129322777 | 129322777 | glycine-rich protein A3-like | 2.4 | 1.68E-04 |
| LOC129322987 | 129322987 | transcription factor HHO5-like | 2.4 | 2.36E-08 |
| LOC129313760 | 129313760 | AT-hook motif nuclear-localized protein 9-like | 2.4 | 4.66E-02 |
| LOC129309264 | 129309264 | cytochrome P450 71A1-like | 2.4 | 9.48E-03 |

| | | | | |
|---|---|---|---|---|
| LOC129294582 | 129294582 | pathogenesis-related thaumatin-like protein 3.5 | 2.4 | 1.98E-03 |
| LOC129288162 | 129288162 | GATA transcription factor 2 | 2.4 | 1.25E-04 |
| LOC129311764 | 129311764 | uncharacterized LOC129311764 | 2.4 | 3.18E-02 |
| LOC129293536 | 129293536 | caffeoyl-CoA O-methyltransferase-like | 2.4 | 1.31E-02 |
| LOC129287781 | 129287781 | uncharacterized LOC129287781 | 2.4 | 6.52E-05 |
| LOC129318879 | 129318879 | protein EIN6 ENHANCER-like | 2.4 | 1.38E-02 |
| LOC129317842 | 129317842 | uncharacterized LOC129317842 | 2.4 | 6.06E-06 |
| LOC129287898 | 129287898 | jasmonate-induced oxygenase 2-like | 2.4 | 8.97E-03 |
| LOC129315939 | 129315939 | uncharacterized protein At5g39865-like | 2.4 | 1.39E-03 |
| LOC129293570 | 129293570 | protein ELF4-LIKE 3 | 2.4 | 5.66E-07 |
| LOC129297137 | 129297137 | peroxidase 42-like | 2.4 | 2.89E-02 |
| LOC129291524 | 129291524 | WEB family protein At2g38370-like | 2.4 | 9.38E-04 |
| LOC129316293 | 129316293 | cyclin-dependent kinases regulatory subunit 1 | 2.4 | 2.70E-05 |
| LOC129315989 | 129315989 | la-related protein 6B-like | 2.4 | 2.83E-02 |
| LOC129313143 | 129313143 | histone H3.2 | 2.4 | 5.95E-03 |
| LOC129306998 | 129306998 | receptor-like serine/threonine-protein kinase SD1-8 | 2.4 | 2.81E-05 |
| LOC129288771 | 129288771 | glycine-rich protein A3-like | 2.4 | 1.44E-02 |
| LOC129292768 | 129292768 | probable serine/threonine-protein kinase At1g54610 | 2.4 | 7.57E-03 |
| LOC129308596 | 129308596 | CRIB domain-containing protein RIC10 | 2.4 | 1.17E-03 |
| LOC129319842 | 129319842 | protein REVEILLE 7-like | 2.4 | 4.35E-06 |
| LOC129311815 | 129311815 | UDP-glycosyltransferase 89A2-like | 2.4 | 4.92E-03 |
| LOC129320355 | 129320355 | cysteine-rich receptor-like protein kinase 10 | 2.3 | 4.11E-02 |
| LOC129306203 | 129306203 | xyloglucan O-acetyltransferase 3-like | 2.3 | 7.58E-03 |
| LOC129298050 | 129298050 | uncharacterized LOC129298050 | 2.3 | 1.23E-02 |
| LOC129320202 | 129320202 | uncharacterized LOC129320202 | 2.3 | 1.89E-02 |
| LOC129298271 | 129298271 | uncharacterized LOC129298271 | 2.3 | 5.95E-05 |
| LOC129315097 | 129315097 | putative cyclin-B3-1 | 2.3 | 5.52E-06 |
| LOC129310008 | 129310008 | uncharacterized LOC129310008 | 2.3 | 1.24E-02 |
| LOC129299139 | 129299139 | proteasome subunit beta type-6 | 2.3 | 4.81E-02 |
| LOC129288747 | 129288747 | uncharacterized LOC129288747 | 2.3 | 5.80E-04 |
| LOC129320802 | 129320802 | chaperone protein dnaJ 20, chloroplastic-like | 2.3 | 4.69E-04 |
| LOC129285474 | 129285474 | uncharacterized LOC129285474 | 2.3 | 4.94E-02 |
| LOC129290301 | 129290301 | vestitone reductase-like | 2.3 | 2.30E-03 |
| LOC129295626 | 129295626 | eukaryotic translation initiation factor 5-like | 2.3 | 8.98E-03 |
| LOC129320152 | 129320152 | probable protein phosphatase 2C 40 | 2.3 | 3.19E-04 |
| LOC129288630 | 129288630 | F-box protein At3g07870-like | 2.3 | 3.08E-02 |
| LOC129284559 | 129284559 | non-specific lipid transfer protein GPI-anchored 15 | 2.3 | 3.39E-02 |
| LOC129309255 | 129309255 | autophagy-related protein 8i-like | 2.3 | 4.78E-04 |
| LOC129305960 | 129305960 | uncharacterized LOC129305960 | 2.3 | 7.12E-03 |
| LOC129308575 | 129308575 | oil body-associated protein 1A-like | 2.3 | 1.55E-03 |
| LOC129286131 | 129286131 | ATP-dependent 6-phosphofructokinase 2 | 2.3 | 5.42E-05 |
| LOC129309045 | 129309045 | uncharacterized LOC129309045 | 2.3 | 3.06E-02 |
| LOC129319379 | 129319379 | aspartic proteinase Asp1-like | 2.3 | 3.26E-03 |
| LOC129320841 | 129320841 | CBL-interacting serine/threonine-protein kinase 6-like | 2.3 | 6.69E-05 |
| LOC129318014 | 129318014 | uncharacterized LOC129318014 | 2.3 | 1.18E-03 |
| LOC129315904 | 129315904 | protein DETOXIFICATION 40-like | 2.3 | 3.99E-02 |
| LOC129317918 | 129317918 | protein DOUBLE-STRAND BREAK FORMATION | 2.3 | 1.08E-03 |
| LOC129313382 | 129313382 | protein NRT1/ PTR FAMILY 5.10-like | 2.3 | 8.33E-03 |

| | | | | |
|---|---|---|---|---|
| LOC129290658 | 129290658 | uncharacterized membrane protein At1g16860-like | 2.3 | 3.03E-02 |
| LOC129311401 | 129311401 | isoflavone reductase homolog | 2.3 | 2.94E-02 |
| LOC129312024 | 129312024 | protein SRC2 homolog | 2.3 | 5.51E-03 |
| LOC129291757 | 129291757 | uncharacterized LOC129291757 | 2.3 | 2.47E-04 |
| LOC129311390 | 129311390 | calcium uniporter protein 2, mitochondrial-like | 2.3 | 9.46E-05 |
| LOC129284947 | 129284947 | protein EARLY-RESPONSIVE TO DEHYDRATION 7, chloro | 2.3 | 1.68E-06 |
| LOC129306925 | 129306925 | probable plastidic glucose transporter 2 | 2.3 | 3.99E-07 |
| LOC129288305 | 129288305 | LOB domain-containing protein 40-like | 2.3 | 9.61E-03 |
| LOC129317721 | 129317721 | kinesin-like protein KIN-12F | 2.3 | 4.98E-02 |
| LOC129295083 | 129295083 | transcription repressor OFP13-like | 2.3 | 3.79E-03 |
| LOC129313394 | 129313394 | MYB-like transcription factor ETC1 | 2.3 | 2.62E-06 |
| LOC129314421 | 129314421 | zinc finger CCCH domain-containing protein 61-like | 2.3 | 4.19E-02 |
| LOC129300978 | 129300978 | 26S proteasome non-ATPase regulatory subunit 13 homolog A | 2.3 | 2.67E-03 |
| LOC129286928 | 129286928 | uncharacterized LOC129286928 | 2.3 | 6.55E-03 |
| LOC129302103 | 129302103 | U-box domain-containing protein 21-like | 2.3 | 8.98E-04 |
| LOC129303174 | 129303174 | probable galactinol--sucrose galactosyltransferase 1 | 2.3 | 2.22E-06 |
| LOC129316910 | 129316910 | probable E3 ubiquitin-protein ligase BAH1-like 1 | 2.3 | 2.80E-05 |
| LOC129309103 | 129309103 | uncharacterized LOC129309103 | 2.3 | 1.93E-04 |
| LOC129319315 | 129319315 | uncharacterized LOC129319315 | 2.3 | 1.62E-03 |
| LOC129286930 | 129286930 | uncharacterized LOC129286930 | 2.3 | 3.54E-03 |
| LOC129298512 | 129298512 | uncharacterized LOC129298512 | 2.3 | 5.80E-04 |
| LOC129320752 | 129320752 | 60S ribosomal protein L39-like | 2.3 | 3.47E-04 |
| LOC129301884 | 129301884 | AAA-ATPase At2g46620-like | 2.3 | 3.02E-04 |
| LOC129292545 | 129292545 | allene oxide synthase 1, chloroplastic-like | 2.3 | 8.99E-04 |
| LOC129291845 | 129291845 | mitochondrial arginine transporter BAC2-like | 2.3 | 2.72E-04 |
| LOC129290308 | 129290308 | trimethyltridecatetraene synthase-like | 2.3 | 3.20E-02 |
| LOC129298467 | 129298467 | polyubiquitin-like | 2.3 | 3.13E-04 |
| LOC129313739 | 129313739 | serine/threonine-protein kinase D6PK-like | 2.3 | 7.25E-07 |
| LOC129320783 | 129320783 | trihelix transcription factor DF1-like | 2.3 | 2.15E-07 |
| LOC129307367 | 129307367 | uncharacterized LOC129307367 | 2.3 | 3.03E-02 |
| LOC129285575 | 129285575 | seipin-1 | 2.3 | 1.82E-04 |
| LOC129297904 | 129297904 | uncharacterized LOC129297904 | 2.3 | 3.91E-02 |
| LOC129311011 | 129311011 | phragmoplastin DRP1C-like | 2.3 | 2.83E-03 |
| LOC129301678 | 129301678 | nuclear transcription factor Y subunit B-3 | 2.2 | 1.21E-04 |
| LOC129320637 | 129320637 | cyclic dof factor 3 | 2.2 | 6.09E-05 |
| LOC129309917 | 129309917 | uncharacterized LOC129309917 | 2.2 | 2.80E-03 |
| LOC129286834 | 129286834 | cysteine proteinase inhibitor 5 | 2.2 | 1.41E-02 |
| LOC129314210 | 129314210 | zinc finger protein CONSTANS-LIKE 4-like | 2.2 | 1.93E-05 |
| LOC129298166 | 129298166 | uncharacterized LOC129298166 | 2.2 | 8.88E-03 |
| LOC129307945 | 129307945 | ethylene-responsive transcription factor RAP2-3-like | 2.2 | 3.60E-04 |
| LOC129285644 | 129285644 | serine/threonine-protein kinase STY46-like | 2.2 | 3.28E-05 |
| LOC129323034 | 129323034 | zinc-finger homeodomain protein 2-like | 2.2 | 2.30E-02 |
| LOC129292187 | 129292187 | uncharacterized LOC129292187 | 2.2 | 5.82E-03 |
| LOC129292603 | 129292603 | uncharacterized LOC129292603 | 2.2 | 1.62E-02 |
| LOC129286024 | 129286024 | disease resistance protein RPV1-like | 2.2 | 9.15E-06 |
| LOC129306402 | 129306402 | dolichyl-diphosphooligosaccharide--protein glycosyltransferase s | 2.2 | 1.90E-02 |
| LOC129293304 | 129293304 | uncharacterized LOC129293304 | 2.2 | 6.72E-07 |
| LOC129306954 | 129306954 | uncharacterized LOC129306954 | 2.2 | 5.19E-03 |

| | | | | |
|---|---|---|---|---|
| LOC129287712 | 129287712 | kinesin-like protein KIN-7O | 2.2 | 1.12E-02 |
| LOC129311591 | 129311591 | benzaldehyde dehydrogenase, mitochondrial | 2.2 | 2.81E-03 |
| LOC129319952 | 129319952 | DNA repair protein XRCC3 homolog | 2.2 | 4.17E-02 |
| LOC129287505 | 129287505 | uncharacterized protein At3g61260-like | 2.2 | 8.25E-03 |
| LOC129284837 | 129284837 | probable choline kinase 1 | 2.2 | 2.80E-03 |
| LOC129307992 | 129307992 | uncharacterized LOC129307992 | 2.2 | 2.34E-04 |
| LOC129291564 | 129291564 | DEAD-box ATP-dependent RNA helicase 31-like | 2.2 | 4.94E-02 |
| LOC129305087 | 129305087 | UDP-glycosyltransferase 73C3-like | 2.2 | 2.27E-02 |
| LOC129322599 | 129322599 | uncharacterized LOC129322599 | 2.2 | 6.24E-03 |
| LOC129311660 | 129311660 | uncharacterized LOC129311660 | 2.2 | 2.67E-02 |
| LOC129296141 | 129296141 | uncharacterized LOC129296141 | 2.2 | 2.08E-10 |
| LOC129314244 | 129314244 | F-box protein At2g27310 | 2.2 | 1.93E-06 |
| LOC129290794 | 129290794 | ferritin-3, chloroplastic-like | 2.2 | 1.52E-06 |
| LOC129311747 | 129311747 | TMV resistance protein N-like | 2.2 | 4.57E-02 |
| LOC129302414 | 129302414 | uncharacterized LOC129302414 | 2.2 | 6.53E-08 |
| LOC129305912 | 129305912 | uncharacterized LOC129305912 | 2.2 | 2.98E-06 |
| LOC129322320 | 129322320 | methyl jasmonate esterase 1-like | 2.2 | 2.10E-03 |
| LOC129285138 | 129285138 | nuclear transcription factor Y subunit B-10-like | 2.2 | 2.38E-07 |
| LOC129309239 | 129309239 | F-box protein SKIP14 | 2.2 | 1.77E-06 |
| LOC129294681 | 129294681 | homeobox-leucine zipper protein ATHB-12-like | 2.2 | 2.93E-04 |
| LOC129302105 | 129302105 | zinc-finger homeodomain protein 2-like | 2.2 | 4.95E-07 |
| LOC129312087 | 129312087 | probable NAD(P)H dehydrogenase (quinone) FQR1-like 2 | 2.2 | 1.57E-03 |
| LOC129313557 | 129313557 | heat shock cognate 70 kDa protein 2-like | 2.2 | 4.35E-03 |
| LOC129294277 | 129294277 | kinesin-like protein KIN-4C | 2.2 | 3.85E-02 |
| LOC129290611 | 129290611 | uncharacterized LOC129290611 | 2.2 | 3.90E-03 |
| LOC129291525 | 129291525 | uncharacterized LOC129291525 | 2.2 | 1.12E-02 |
| LOC129285787 | 129285787 | cytochrome b561, DM13 and DOMON domain-containing prot | 2.2 | 1.11E-04 |
| LOC129319843 | 129319843 | galactinol--sucrose galactosyltransferase-like | 2.2 | 1.46E-02 |
| LOC129311739 | 129311739 | uncharacterized LOC129311739 | 2.2 | 2.15E-04 |
| LOC129322308 | 129322308 | uncharacterized LOC129322308 | 2.2 | 7.44E-05 |
| LOC129297277 | 129297277 | nuclear transcription factor Y subunit C-2-like | 2.2 | 7.40E-13 |
| LOC129306370 | 129306370 | thioredoxin-like 1-1, chloroplastic | 2.2 | 1.51E-08 |
| LOC129317106 | 129317106 | 40S ribosomal protein S30 | 2.2 | 4.64E-02 |
| LOC129303105 | 129303105 | zinc finger CCCH domain-containing protein 20-like | 2.2 | 2.78E-07 |
| LOC129303504 | 129303504 | homeobox-leucine zipper protein HOX15-like | 2.2 | 4.59E-03 |
| LOC129306927 | 129306927 | uncharacterized LOC129306927 | 2.2 | 5.29E-04 |
| LOC129318568 | 129318568 | uncharacterized LOC129318568 | 2.2 | 3.23E-02 |
| LOC129294867 | 129294867 | cytochrome c biogenesis CcmF C-terminal-like mitochondrial p | 2.2 | 2.01E-02 |
| LOC129295989 | 129295989 | uncharacterized LOC129295989 | 2.2 | 3.50E-02 |
| LOC129284533 | 129284533 | caffeoylshikimate esterase-like | 2.2 | 5.80E-04 |
| LOC129323161 | 129323161 | probable copper-transporting ATPase HMA5 | 2.2 | 8.48E-06 |
| LOC129319939 | 129319939 | uncharacterized LOC129319939 | 2.2 | 4.33E-03 |
| LOC129302362 | 129302362 | DIS3-like exonuclease 2 | 2.2 | 1.41E-06 |
| LOC129289298 | 129289298 | probable disease resistance protein At1g61300 | 2.2 | 4.89E-02 |
| LOC129294987 | 129294987 | mediator of RNA polymerase II transcription subunit 19a-like | 2.2 | 9.18E-03 |
| LOC129311512 | 129311512 | F-box protein PP2-A12-like | 2.2 | 1.66E-04 |
| LOC129307415 | 129307415 | guanylate kinase 2 | 2.2 | 1.47E-03 |
| LOC129302378 | 129302378 | 60S ribosomal protein L39 | 2.1 | 2.69E-03 |

| | | | | |
|---|---|---|---|---|
| LOC129314257 | 129314257 | vegetative cell wall protein gp1 | 2.1 | 1.93E-03 |
| LOC129315683 | 129315683 | ABC transporter F family member 1 | 2.1 | 1.29E-05 |
| LOC129303939 | 129303939 | uncharacterized LOC129303939 | 2.1 | 3.95E-02 |
| LOC129304743 | 129304743 | CSC1-like protein ERD4 | 2.1 | 1.33E-07 |
| LOC129292900 | 129292900 | S-adenosylmethionine decarboxylase proenzyme-like | 2.1 | 7.43E-04 |
| LOC129313569 | 129313569 | neutral/alkaline invertase 3, chloroplastic-like | 2.1 | 1.63E-03 |
| LOC129285073 | 129285073 | dynamin-related protein 4C-like | 2.1 | 3.64E-02 |
| LOC129316892 | 129316892 | cytochrome c | 2.1 | 2.13E-04 |
| LOC129306733 | 129306733 | glutaredoxin-C13-like | 2.1 | 3.53E-03 |
| LOC129320044 | 129320044 | uncharacterized LOC129320044 | 2.1 | 2.14E-05 |
| LOC129311868 | 129311868 | RING-H2 finger protein ATL80-like | 2.1 | 1.85E-05 |
| LOC129291689 | 129291689 | uncharacterized LOC129291689 | 2.1 | 2.86E-02 |
| LOC129305027 | 129305027 | probable microtubule-binding protein TANGLED | 2.1 | 4.22E-02 |
| LOC129303799 | 129303799 | AT-hook motif nuclear-localized protein 15-like | 2.1 | 1.07E-04 |
| LOC129322496 | 129322496 | uncharacterized LOC129322496 | 2.1 | 1.21E-04 |
| LOC129290107 | 129290107 | tropinone reductase homolog | 2.1 | 4.37E-03 |
| LOC129314426 | 129314426 | uncharacterized LOC129314426 | 2.1 | 2.51E-02 |
| LOC129308143 | 129308143 | PLASMODESMATA CALLOSE-BINDING PROTEIN 3-like | 2.1 | 3.92E-02 |
| LOC129311715 | 129311715 | adenine phosphoribosyltransferase 3-like | 2.1 | 2.55E-02 |
| LOC129306454 | 129306454 | uncharacterized LOC129306454 | 2.1 | 3.78E-04 |
| LOC129302033 | 129302033 | basic leucine zipper 9-like | 2.1 | 1.55E-03 |
| LOC129302264 | 129302264 | protein CYPRO4 | 2.1 | 2.66E-02 |
| LOC129306569 | 129306569 | thiamine pyrophosphokinase 1-like | 2.1 | 1.06E-04 |
| LOC129293706 | 129293706 | uncharacterized LOC129293706 | 2.1 | 9.64E-04 |
| LOC129300297 | 129300297 | serine/threonine-protein kinase Aurora-2 | 2.1 | 5.52E-06 |
| LOC129285755 | 129285755 | 4-hydroxy-3-methylbut-2-enyl diphosphate reductase, chloroplas | 2.1 | 1.34E-02 |
| LOC129293292 | 129293292 | probable indole-3-acetic acid-amido synthetase GH3.1 | 2.1 | 3.61E-02 |
| LOC129318613 | 129318613 | ABC transporter G family member 22-like | 2.1 | 3.87E-05 |
| LOC129306020 | 129306020 | probable protein phosphatase 2C 49 | 2.1 | 2.59E-06 |
| LOC129308540 | 129308540 | polyubiquitin 11 | 2.1 | 4.94E-03 |
| LOC129294887 | 129294887 | ATP synthase subunit 9, mitochondrial | 2.1 | 4.01E-03 |
| LOC129320312 | 129320312 | uncharacterized LOC129320312 | 2.1 | 2.26E-02 |
| LOC129311994 | 129311994 | uncharacterized LOC129311994 | 2.1 | 4.70E-03 |
| LOC129300604 | 129300604 | serine/threonine-protein kinase STY13-like | 2.1 | 3.87E-02 |
| LOC129288931 | 129288931 | cytochrome c oxidase subunit 5C | 2.1 | 3.65E-03 |
| LOC129309552 | 129309552 | 40S ribosomal protein S21 | 2.1 | 4.23E-02 |
| LOC129312194 | 129312194 | brassinosteroid-responsive RING protein 1-like | 2.1 | 3.11E-02 |
| LOC129321242 | 129321242 | probable isoprenylcysteine alpha-carbonyl methylesterase ICME | 2.1 | 5.50E-03 |
| LOC129300073 | 129300073 | probable mannitol dehydrogenase | 2.1 | 1.16E-02 |
| LOC129306232 | 129306232 | phosphatidylinositol 4-kinase gamma 7-like | 2.1 | 2.20E-03 |
| LOC129293413 | 129293413 | cystinosin homolog | 2.1 | 4.84E-02 |
| LOC129318279 | 129318279 | heat shock factor protein HSF30 | 2.1 | 7.16E-03 |
| LOC129310938 | 129310938 | CDP-diacylglycerol--serine O-phosphatidyltransferase 1-like | 2.1 | 8.11E-05 |
| LOC129288518 | 129288518 | cytochrome P450 78A3-like | 2.1 | 6.87E-03 |
| LOC129301816 | 129301816 | uncharacterized LOC129301816 | 2.1 | 1.58E-04 |
| LOC129286661 | 129286661 | probable inositol transporter 2 | 2.1 | 6.42E-03 |
| LOC129313666 | 129313666 | uncharacterized LOC129313666 | 2.1 | 5.92E-03 |
| LOC129305835 | 129305835 | protein EARLY RESPONSIVE TO DEHYDRATION 15-like | 2.1 | 2.09E-02 |

| | | | | |
|---|---|---|---|---|
| LOC129285831 | 129285831 | protein DMR6-LIKE OXYGENASE 2-like | 2.1 | 3.03E-02 |
| LOC129288574 | 129288574 | AAA-ATPase At2g46620 | 2.1 | 1.51E-02 |
| LOC129287984 | 129287984 | ACT domain-containing protein ACR6 | 2.1 | 2.81E-03 |
| LOC129297814 | 129297814 | 26S proteasome non-ATPase regulatory subunit 13 homolog A | 2.1 | 3.23E-02 |
| LOC129285584 | 129285584 | tyrosine-protein phosphatase DSP5 | 2.1 | 1.53E-04 |
| LOC129300834 | 129300834 | uncharacterized LOC129300834 | 2.1 | 9.30E-03 |
| LOC129309060 | 129309060 | ribonucleoside-diphosphate reductase large subunit | 2.1 | 4.54E-03 |
| LOC129295617 | 129295617 | serine/threonine protein phosphatase 2A 57 kDa regulatory subu | 2.1 | 4.63E-03 |
| LOC129286327 | 129286327 | glutamyl-tRNA reductase 2, chloroplastic-like | 2.1 | 7.33E-04 |
| LOC129288862 | 129288862 | zinc finger CCCH domain-containing protein 62-like | 2.1 | 3.79E-02 |
| LOC129287723 | 129287723 | trihelix transcription factor ASIL2-like | 2.1 | 5.06E-07 |
| LOC129309113 | 129309113 | uncharacterized LOC129309113 | 2.1 | 2.04E-04 |
| LOC129290587 | 129290587 | uncharacterized LOC129290587 | 2.1 | 5.38E-03 |
| LOC129287197 | 129287197 | secoisolariciresinol dehydrogenase-like | 2.1 | 4.27E-02 |
| LOC129296953 | 129296953 | UDP-arabinose 4-epimerase 1-like | 2.1 | 2.81E-04 |
| LOC129307964 | 129307964 | AAA-ATPase At5g57480-like | 2.1 | 1.56E-02 |
| LOC129292514 | 129292514 | uncharacterized LOC129292514 | 2.1 | 8.34E-05 |
| LOC129290907 | 129290907 | histone H3-like centromeric protein CENH3 | 2.1 | 4.57E-02 |
| LOC129306612 | 129306612 | two-pore potassium channel 3-like | 2.1 | 1.72E-03 |
| LOC129291302 | 129291302 | uncharacterized protein At4g00950-like | 2.1 | 4.38E-03 |
| LOC129289403 | 129289403 | probable GTP diphosphokinase RSH3, chloroplastic | 2.1 | 8.12E-04 |
| LOC129318559 | 129318559 | BTB/POZ domain-containing protein At1g55760-like | 2.1 | 5.20E-04 |
| LOC129289693 | 129289693 | uncharacterized LOC129289693 | 2.1 | 6.49E-03 |
| LOC129288689 | 129288689 | uncharacterized LOC129288689 | 2.1 | 3.41E-03 |
| LOC129306445 | 129306445 | auxin response factor 1-like | 2.0 | 7.54E-04 |
| LOC129315849 | 129315849 | uncharacterized LOC129315849 | 2.0 | 4.89E-03 |
| LOC129313428 | 129313428 | uncharacterized LOC129313428 | 2.0 | 1.18E-02 |
| LOC129305964 | 129305964 | uncharacterized LOC129305964 | 2.0 | 2.62E-10 |
| LOC129319428 | 129319428 | putative glycerol-3-phosphate transporter 5 | 2.0 | 2.09E-03 |
| LOC129306035 | 129306035 | ABC transporter B family member 19 | 2.0 | 3.24E-02 |
| LOC129295327 | 129295327 | 12-oxophytodienoate reductase 3-like | 2.0 | 4.51E-02 |
| LOC129288179 | 129288179 | uncharacterized LOC129288179 | 2.0 | 2.42E-04 |
| LOC129288787 | 129288787 | auxin-responsive protein IAA17-like | 2.0 | 2.03E-03 |
| LOC129313141 | 129313141 | uncharacterized LOC129313141 | 2.0 | 2.47E-04 |
| LOC129322708 | 129322708 | PLASMODESMATA CALLOSE-BINDING PROTEIN 3-like | 2.0 | 5.02E-05 |
| LOC129301964 | 129301964 | protein transport protein Sec61 subunit gamma-like | 2.0 | 5.80E-03 |
| LOC129285089 | 129285089 | histone H2AX | 2.0 | 1.25E-02 |
| LOC129306818 | 129306818 | calmodulin-binding protein 25-like | 2.0 | 1.85E-03 |
| LOC129301913 | 129301913 | 60S ribosomal protein L10-like | 2.0 | 9.63E-03 |
| LOC129296493 | 129296493 | acireductone dioxygenase 2-like | 2.0 | 5.45E-04 |
| LOC129299342 | 129299342 | exocyst complex component EXO70B1-like | 2.0 | 2.86E-04 |
| LOC129289788 | 129289788 | uncharacterized LOC129289788 | 2.0 | 3.32E-04 |
| LOC129311356 | 129311356 | glycosyltransferase BC10-like | 2.0 | 4.16E-04 |
| LOC129315925 | 129315925 | aquaporin PIP2-2 | 2.0 | 1.48E-04 |
| LOC129289610 | 129289610 | uncharacterized LOC129289610 | 2.0 | 2.16E-02 |
| LOC129292525 | 129292525 | zinc finger CCCH domain-containing protein 20-like | 2.0 | 7.65E-04 |
| LOC129286612 | 129286612 | uncharacterized LOC129286612 | 2.0 | 6.77E-04 |
| LOC129306715 | 129306715 | uncharacterized LOC129306715 | 2.0 | 2.51E-03 |

| | | | | |
|---|---|---|---|---|
| LOC129309185 | 129309185 | zinc finger protein CONSTANS-LIKE 5 | 2.0 | 5.22E-04 |
| LOC129293661 | 129293661 | calmodulin-like protein 30 | 2.0 | 2.76E-02 |
| LOC129318467 | 129318467 | probable serine/threonine-protein kinase WNK11 | 2.0 | 4.18E-02 |
| LOC129292561 | 129292561 | receptor-like serine/threonine-protein kinase At4g25390 | 2.0 | 1.05E-03 |
| LOC129318594 | 129318594 | uncharacterized protein At5g39865 | 2.0 | 4.85E-03 |
| LOC129294728 | 129294728 | uncharacterized LOC129294728 | 2.0 | 2.32E-04 |
| LOC129321379 | 129321379 | 1-aminocyclopropane-1-carboxylate oxidase homolog 4-like | 2.0 | 3.31E-02 |
| LOC129290789 | 129290789 | NAC domain-containing protein 30-like | 2.0 | 1.09E-03 |
| LOC129300647 | 129300647 | fruit protein pKIWI502-like | 2.0 | 8.48E-03 |
| LOC129299945 | 129299945 | uncharacterized LOC129299945 | 2.0 | 2.09E-02 |
| LOC129319087 | 129319087 | scarecrow-like transcription factor PAT1 | 2.0 | 1.93E-06 |
| LOC129319327 | 129319327 | uncharacterized LOC129319327 | 2.0 | 1.38E-02 |
| LOC129290369 | 129290369 | cationic amino acid transporter 9, chloroplastic-like | 2.0 | 1.38E-03 |
| LOC129284911 | 129284911 | filament-like plant protein 7 | 2.0 | 1.20E-02 |
| LOC129312724 | 129312724 | O-fucosyltransferase 9 | 2.0 | 2.13E-02 |
| LOC129291030 | 129291030 | common plant regulatory factor 1 | 2.0 | 8.96E-09 |
| LOC129303225 | 129303225 | calcineurin B-like protein 3 | 2.0 | 3.27E-02 |
| LOC129305153 | 129305153 | ethylene-responsive transcription factor RAP2-3-like | 2.0 | 1.80E-02 |
| LOC129311578 | 129311578 | protein DETOXIFICATION 14-like | 2.0 | 1.89E-02 |
| LOC129320320 | 129320320 | protein DSS1 HOMOLOG ON CHROMOSOME V-like | 2.0 | 1.53E-03 |
| LOC129323164 | 129323164 | acyl-CoA-binding domain-containing protein 3-like | 2.0 | 1.04E-02 |
| LOC129317573 | 129317573 | uncharacterized LOC129317573 | 2.0 | 6.80E-05 |
| LOC129285424 | 129285424 | protein LNK1-like | 2.0 | 5.50E-03 |
| LOC129318134 | 129318134 | serine/arginine-rich splicing factor SR45a-like | 2.0 | 5.42E-05 |
| LOC129294386 | 129294386 | bet1-like protein At4g14600 | 2.0 | 6.79E-03 |
| LOC129309560 | 129309560 | heavy metal-associated isoprenylated plant protein 7 | 2.0 | 1.69E-02 |
| LOC129306974 | 129306974 | protein PHLOEM PROTEIN 2-LIKE A9-like | 2.0 | 2.70E-02 |
| LOC129285023 | 129285023 | protein kinase PINOID | 2.0 | 4.48E-04 |
| LOC129300244 | 129300244 | splicing factor Cactin-like | 2.0 | 2.10E-02 |
| LOC129309969 | 129309969 | phospholipid:diacylglycerol acyltransferase 1-like | 2.0 | 1.11E-03 |
| LOC129295892 | 129295892 | small heat shock protein, chloroplastic-like | 2.0 | 1.64E-02 |
| LOC129293988 | 129293988 | uncharacterized LOC129293988 | 2.0 | 3.37E-02 |
| LOC129306989 | 129306989 | polygalacturonase QRT3 | 2.0 | 7.17E-05 |
| LOC129314480 | 129314480 | E3 ubiquitin-protein ligase DIS1-like | 2.0 | 2.08E-03 |
| LOC129298843 | 129298843 | uncharacterized LOC129298843 | 2.0 | 1.28E-03 |
| LOC129311744 | 129311744 | probable aspartyl protease At4g16563 | 2.0 | 3.01E-03 |
| LOC129297767 | 129297767 | ATP synthase subunit alpha, chloroplastic | 2.0 | 1.63E-02 |
| LOC129292073 | 129292073 | type IV inositol polyphosphate 5-phosphatase 3 | 2.0 | 2.36E-03 |
| LOC129323153 | 129323153 | basic leucine zipper 4 | 2.0 | 1.42E-03 |
| LOC129296446 | 129296446 | ethylene-responsive transcription factor 1A-like | 2.0 | 2.62E-04 |
| LOC129305142 | 129305142 | pathogenesis-related thaumatin-like protein 3.5 | 2.0 | 3.26E-02 |
| LOC129308201 | 129308201 | protein SENSITIVE TO PROTON RHIZOTOXICITY 1-like | 2.0 | 2.85E-04 |
| LOC129284914 | 129284914 | protein NRT1/ PTR FAMILY 4.3-like | 2.0 | 7.36E-04 |
| LOC129301515 | 129301515 | probable CoA ligase CCL5 | 2.0 | 7.83E-08 |
| LOC129321219 | 129321219 | uncharacterized membrane protein At3g27390 | 2.0 | 4.37E-02 |
| LOC129315103 | 129315103 | (R,S)-reticuline 7-O-methyltransferase-like | 2.0 | 2.88E-03 |
| LOC129301306 | 129301306 | probable RNA 3'-terminal phosphate cyclase-like protein | 2.0 | 7.57E-03 |
| LOC129318286 | 129318286 | 60S ribosomal protein L39-like | 2.0 | 2.15E-02 |

| | | | | |
|---|---|---|---|---|
| LOC129314980 | 129314980 | E2F transcription factor-like E2FF | 2.0 | 1.24E-02 |
| LOC129303870 | 129303870 | uncharacterized LOC129303870 | 2.0 | 8.99E-04 |
| LOC129288379 | 129288379 | uncharacterized LOC129288379 | 2.0 | 1.80E-04 |
| LOC129294669 | 129294669 | universal stress protein A-like protein | 2.0 | 3.66E-04 |
| LOC129320336 | 129320336 | EG45-like domain containing protein | 2.0 | 4.32E-02 |
| LOC129294896 | 129294896 | NADH dehydrogenase [ubiquinone] iron-sulfur protein 2-like | 2.0 | 5.81E-03 |
| LOC129293924 | 129293924 | glucan endo-1,3-beta-glucosidase-like | 2.0 | 4.42E-03 |
| LOC129309143 | 129309143 | uncharacterized LOC129309143 | 1.9 | 9.56E-06 |
| LOC129295477 | 129295477 | zinc finger A20 and AN1 domain-containing stress-associated p | 1.9 | 1.07E-02 |
| LOC129317683 | 129317683 | uncharacterized LOC129317683 | 1.9 | 3.06E-02 |
| LOC129320466 | 129320466 | uncharacterized LOC129320466 | 1.9 | 6.61E-03 |
| LOC129295602 | 129295602 | uncharacterized LOC129295602 | 1.9 | 3.08E-02 |
| LOC129321497 | 129321497 | uncharacterized LOC129321497 | 1.9 | 1.87E-04 |
| LOC129322748 | 129322748 | IRK-interacting protein-like | 1.9 | 4.24E-02 |
| LOC129312078 | 129312078 | E3 ubiquitin-protein ligase MPSR1-like | 1.9 | 4.49E-03 |
| LOC129309768 | 129309768 | GATA transcription factor 15 | 1.9 | 3.39E-03 |
| LOC129294148 | 129294148 | ethylene-responsive transcription factor RAP2-4 | 1.9 | 5.02E-03 |
| LOC129286913 | 129286913 | protein PSK SIMULATOR 1-like | 1.9 | 1.77E-04 |
| LOC129289577 | 129289577 | LRR receptor-like serine/threonine-protein kinase GSO1 | 1.9 | 2.16E-02 |
| LOC129318872 | 129318872 | probable long-chain-alcohol O-fatty-acyltransferase 5 | 1.9 | 4.19E-02 |
| LOC129312152 | 129312152 | uncharacterized LOC129312152 | 1.9 | 2.23E-04 |
| LOC129321578 | 129321578 | AP2-like ethylene-responsive transcription factor At2g41710 | 1.9 | 2.24E-06 |
| LOC129310608 | 129310608 | protein RGF1 INDUCIBLE TRANSCRIPTION FACTOR 1-lik | 1.9 | 7.47E-04 |
| LOC129321343 | 129321343 | protein EARLY RESPONSIVE TO DEHYDRATION 15-like | 1.9 | 1.22E-03 |
| LOC129298407 | 129298407 | lactoylglutathione lyase GLX1-like | 1.9 | 1.77E-02 |
| LOC129305244 | 129305244 | AP2-like ethylene-responsive transcription factor At2g41710 | 1.9 | 1.74E-02 |
| LOC129311015 | 129311015 | calvin cycle protein CP12-2, chloroplastic-like | 1.9 | 6.72E-03 |
| LOC129319984 | 129319984 | pectinesterase 3-like | 1.9 | 3.79E-03 |
| LOC129292148 | 129292148 | protein DMR6-LIKE OXYGENASE 1-like | 1.9 | 9.42E-03 |
| LOC129290836 | 129290836 | uncharacterized LOC129290836 | 1.9 | 5.43E-03 |
| LOC129285495 | 129285495 | uncharacterized LOC129285495 | 1.9 | 2.29E-02 |
| LOC129302257 | 129302257 | inositol-tetrakisphosphate 1-kinase 3-like | 1.9 | 2.63E-06 |
| LOC129316154 | 129316154 | zinc finger A20 and AN1 domain-containing stress-associated p | 1.9 | 1.46E-04 |
| LOC129291076 | 129291076 | ultraviolet-B receptor UVR8-like | 1.9 | 4.33E-03 |
| LOC129309983 | 129309983 | FCS-Like Zinc finger 15-like | 1.9 | 1.31E-02 |
| LOC129290265 | 129290265 | mitochondrial import inner membrane translocase subunit TIM1 | 1.9 | 1.46E-02 |
| LOC129311583 | 129311583 | homeobox-leucine zipper protein HAT22 | 1.9 | 9.99E-04 |
| LOC129315024 | 129315024 | early nodulin-like protein 2 | 1.9 | 6.47E-05 |
| LOC129284984 | 129284984 | peptidyl-prolyl cis-trans isomerase-like | 1.9 | 9.91E-04 |
| LOC129322885 | 129322885 | protein EARLY-RESPONSIVE TO DEHYDRATION 7, chloro | 1.9 | 4.56E-02 |
| LOC129296939 | 129296939 | protein CANDIDATE G-PROTEIN COUPLED RECEPTOR 7- | 1.9 | 2.91E-02 |
| LOC129311528 | 129311528 | FCS-Like Zinc finger 1-like | 1.9 | 1.61E-02 |
| LOC129317966 | 129317966 | proliferating cell nuclear antigen | 1.9 | 5.62E-04 |
| LOC129295106 | 129295106 | calmodulin-binding receptor kinase CaMRLK-like | 1.9 | 4.03E-02 |
| LOC129313148 | 129313148 | protein S40-4-like | 1.9 | 2.08E-03 |
| LOC129314667 | 129314667 | protein LNK4-like | 1.9 | 7.47E-04 |
| LOC129309461 | 129309461 | TORTIFOLIA1-like protein 3 | 1.9 | 2.68E-03 |
| LOC129288821 | 129288821 | auxin-induced in root cultures protein 12 | 1.9 | 1.12E-03 |

| | | | | |
|---|---|---|---|---|
| LOC129313698 | 129313698 | protein LYK5 | 1.9 | 2.71E-03 |
| LOC129312314 | 129312314 | putative 1-phosphatidylinositol-3-phosphate 5-kinase FAB1D | 1.9 | 9.98E-05 |
| LOC129317866 | 129317866 | cellulose synthase-like protein E1 | 1.9 | 7.06E-03 |
| LOC129302507 | 129302507 | peroxidase 46-like | 1.9 | 3.11E-04 |
| LOC129290833 | 129290833 | BAG family molecular chaperone regulator 6 | 1.9 | 3.75E-03 |
| LOC129308232 | 129308232 | probable E3 ubiquitin-protein ligase XERICO | 1.9 | 3.06E-03 |
| LOC129291381 | 129291381 | uncharacterized LOC129291381 | 1.9 | 1.69E-05 |
| LOC129304785 | 129304785 | GRF1-interacting factor 1 | 1.9 | 6.48E-05 |
| LOC129305176 | 129305176 | SAGA-associated factor 29 homolog A-like | 1.9 | 2.85E-06 |
| LOC129313594 | 129313594 | B-box zinc finger protein 22-like | 1.9 | 2.19E-03 |
| LOC129294411 | 129294411 | uncharacterized LOC129294411 | 1.9 | 8.75E-03 |
| LOC129311985 | 129311985 | classical arabinogalactan protein 9-like | 1.9 | 9.63E-03 |
| LOC129297021 | 129297021 | protein NUCLEAR FUSION DEFECTIVE 4-like | 1.9 | 3.17E-02 |
| LOC129288353 | 129288353 | protein HOTHEAD | 1.9 | 4.74E-02 |
| LOC129318451 | 129318451 | uncharacterized LOC129318451 | 1.9 | 8.12E-05 |
| LOC129313717 | 129313717 | protein TPX2 | 1.9 | 3.78E-02 |
| LOC129296613 | 129296613 | linoleate 13S-lipoxygenase 3-1, chloroplastic-like | 1.9 | 3.30E-02 |
| LOC129293392 | 129293392 | 40S ribosomal protein S16-like | 1.9 | 4.37E-03 |
| LOC129315495 | 129315495 | ethylene receptor | 1.9 | 4.26E-04 |
| LOC129319854 | 129319854 | ubiquitin-conjugating enzyme E2-23 kDa-like | 1.9 | 4.91E-02 |
| LOC129288707 | 129288707 | uncharacterized LOC129288707 | 1.9 | 5.08E-03 |
| LOC129291338 | 129291338 | AT-hook motif nuclear-localized protein 7 | 1.9 | 2.46E-08 |
| LOC129295484 | 129295484 | uncharacterized LOC129295484 | 1.9 | 4.42E-02 |
| LOC129303259 | 129303259 | uncharacterized LOC129303259 | 1.9 | 1.19E-02 |
| LOC129293382 | 129293382 | uncharacterized LOC129293382 | 1.9 | 7.35E-03 |
| LOC129285440 | 129285440 | metal tolerance protein 11 | 1.9 | 4.48E-04 |
| LOC129321498 | 129321498 | uncharacterized LOC129321498 | 1.9 | 1.14E-03 |
| LOC129320957 | 129320957 | protein MHF1 homolog | 1.9 | 3.24E-02 |
| LOC129294198 | 129294198 | pentatricopeptide repeat-containing protein DOT4, chloroplastic | 1.9 | 1.34E-03 |
| LOC129293645 | 129293645 | probable 1-deoxy-D-xylulose-5-phosphate synthase, chloroplasti | 1.9 | 1.26E-03 |
| LOC129318625 | 129318625 | E3 ubiquitin-protein ligase ATL23 | 1.9 | 1.19E-06 |
| LOC129303387 | 129303387 | WD repeat-containing protein LWD1-like | 1.9 | 7.92E-03 |
| LOC129314936 | 129314936 | uncharacterized LOC129314936 | 1.9 | 4.35E-03 |
| LOC129310083 | 129310083 | bHLH transcription factor RHL1-like | 1.9 | 3.41E-04 |
| LOC129311986 | 129311986 | uncharacterized LOC129311986 | 1.9 | 3.91E-02 |
| LOC129320915 | 129320915 | uncharacterized LOC129320915 | 1.9 | 4.30E-02 |
| LOC129291208 | 129291208 | rop guanine nucleotide exchange factor 7-like | 1.9 | 1.99E-02 |
| LOC129312568 | 129312568 | pheophytinase, chloroplastic-like | 1.9 | 1.26E-02 |
| LOC129308400 | 129308400 | uncharacterized LOC129308400 | 1.9 | 2.83E-02 |
| LOC129292966 | 129292966 | probable isoprenylcysteine alpha-carbonyl methylesterase ICME | 1.9 | 2.48E-02 |
| LOC129290799 | 129290799 | kinetochore protein SPC24 homolog | 1.9 | 8.65E-03 |
| LOC129298009 | 129298009 | AP-1 complex subunit mu-2 | 1.9 | 1.49E-02 |
| LOC129292320 | 129292320 | uncharacterized LOC129292320 | 1.9 | 4.33E-02 |
| LOC129311723 | 129311723 | cyclin-dependent kinase inhibitor 7-like | 1.9 | 1.76E-03 |
| LOC129305783 | 129305783 | chlorophyll a-b binding protein of LHCII type 1-like | 1.9 | 1.27E-02 |
| LOC129309380 | 129309380 | EIN3-binding F-box protein 1-like | 1.9 | 1.13E-06 |
| LOC129313578 | 129313578 | condensin complex subunit 2 | 1.9 | 4.87E-02 |
| LOC129285513 | 129285513 | extradiol ring-cleavage dioxygenase-like | 1.9 | 4.02E-02 |

| | | | | |
|---|---|---|---|---|
| LOC129318768 | 129318768 | VQ motif-containing protein 22-like | 1.9 | 3.99E-02 |
| LOC129302290 | 129302290 | cyclin-A1-1-like | 1.9 | 9.10E-06 |
| LOC129297120 | 129297120 | uncharacterized LOC129297120 | 1.9 | 4.47E-02 |
| LOC129310677 | 129310677 | calcineurin B-like protein 1 | 1.9 | 3.59E-06 |
| LOC129319976 | 129319976 | uncharacterized LOC129319976 | 1.9 | 3.69E-03 |
| LOC129303403 | 129303403 | uncharacterized LOC129303403 | 1.9 | 1.05E-02 |
| LOC129307754 | 129307754 | uncharacterized LOC129307754 | 1.9 | 1.56E-03 |
| LOC129294850 | 129294850 | ribosomal protein S4, mitochondrial | 1.9 | 8.31E-03 |
| LOC129310002 | 129310002 | uncharacterized LOC129310002 | 1.9 | 2.87E-02 |
| LOC129292245 | 129292245 | glycine-rich RNA-binding protein 2-like | 1.9 | 1.72E-02 |
| LOC129309397 | 129309397 | crossover junction endonuclease EME1B | 1.8 | 1.12E-02 |
| LOC129309481 | 129309481 | plant intracellular Ras-group-related LRR protein 7-like | 1.8 | 4.26E-02 |
| LOC129311631 | 129311631 | putative F-box protein At1g47790 | 1.8 | 5.31E-05 |
| LOC129313508 | 129313508 | uncharacterized LOC129313508 | 1.8 | 4.92E-02 |
| LOC129311891 | 129311891 | phosphatidylinositol 4-kinase gamma 5-like | 1.8 | 4.44E-07 |
| LOC129291293 | 129291293 | vestitone reductase-like | 1.8 | 3.21E-02 |
| LOC129292757 | 129292757 | cysteine-rich and transmembrane domain-containing protein WI | 1.8 | 1.75E-02 |
| LOC129322825 | 129322825 | uncharacterized LOC129322825 | 1.8 | 1.04E-03 |
| LOC129304768 | 129304768 | uncharacterized LOC129304768 | 1.8 | 1.50E-03 |
| LOC129289177 | 129289177 | probable mediator of RNA polymerase II transcription subunit 2 | 1.8 | 1.19E-04 |
| LOC129292830 | 129292830 | cyclase-associated protein 1-like | 1.8 | 1.14E-04 |
| LOC129305794 | 129305794 | serine/arginine-rich splicing factor SR30-like | 1.8 | 6.09E-05 |
| LOC129313561 | 129313561 | CBS domain-containing protein CBSX5 | 1.8 | 4.08E-02 |
| LOC129321611 | 129321611 | protein SRC2-like | 1.8 | 6.85E-04 |
| LOC129303556 | 129303556 | homeobox-leucine zipper protein HAT5-like | 1.8 | 1.43E-02 |
| LOC129315553 | 129315553 | YTH domain-containing protein ECT3 | 1.8 | 5.16E-03 |
| LOC129309574 | 129309574 | uncharacterized LOC129309574 | 1.8 | 3.42E-02 |
| LOC129315412 | 129315412 | uncharacterized LOC129315412 | 1.8 | 2.03E-02 |
| LOC129298662 | 129298662 | pyruvate dehydrogenase (acetyl-transferring) kinase, mitochondı | 1.8 | 2.29E-02 |
| LOC129304673 | 129304673 | probable LRR receptor-like serine/threonine-protein kinase RKF | 1.8 | 1.93E-02 |
| LOC129320487 | 129320487 | calcium-dependent protein kinase 10-like | 1.8 | 2.75E-02 |
| LOC129310471 | 129310471 | bax inhibitor 1-like | 1.8 | 2.50E-03 |
| LOC129285947 | 129285947 | cardiolipin synthase (CMP-forming), mitochondrial | 1.8 | 9.93E-07 |
| LOC129309570 | 129309570 | histone H4 | 1.8 | 2.37E-02 |
| LOC129321295 | 129321295 | alkaline/neutral invertase A, mitochondrial-like | 1.8 | 8.53E-03 |
| LOC129284878 | 129284878 | F-box protein SKIP19-like | 1.8 | 1.39E-04 |
| LOC129305109 | 129305109 | uncharacterized LOC129305109 | 1.8 | 1.85E-02 |
| LOC129322714 | 129322714 | lysine-specific demethylase JMJ13-like | 1.8 | 6.27E-05 |
| LOC129316038 | 129316038 | sm-like protein LSM8 | 1.8 | 1.84E-03 |
| LOC129294897 | 129294897 | NADH dehydrogenase [ubiquinone] iron-sulfur protein 2-like | 1.8 | 2.38E-02 |
| LOC129303343 | 129303343 | ethylene-responsive transcription factor RAP2-13-like | 1.8 | 6.18E-08 |
| LOC129322909 | 129322909 | mitochondrial import inner membrane translocase subunit TIM1 | 1.8 | 7.57E-03 |
| LOC129316373 | 129316373 | soluble inorganic pyrophosphatase 4 | 1.8 | 7.59E-03 |
| LOC129313411 | 129313411 | ABC transporter C family member 5-like | 1.8 | 1.23E-02 |
| LOC129314875 | 129314875 | glyceraldehyde-3-phosphate dehydrogenase GAPCP1, chloropla | 1.8 | 1.87E-02 |
| LOC129309112 | 129309112 | probable protein phosphatase 2C 33 | 1.8 | 1.54E-05 |
| LOC129316159 | 129316159 | NAC domain-containing protein 90-like | 1.8 | 2.30E-02 |
| LOC129292813 | 129292813 | polyubiquitin-like | 1.8 | 2.87E-04 |

| | | | | |
|---|---|---|---|---|
| LOC129285828 | 129285828 | uncharacterized LOC129285828 | 1.8 | 2.27E-04 |
| LOC129294286 | 129294286 | VQ motif-containing protein 9-like | 1.8 | 2.04E-02 |
| LOC129314902 | 129314902 | uncharacterized LOC129314902 | 1.8 | 1.98E-02 |
| LOC129300346 | 129300346 | uncharacterized LOC129300346 | 1.8 | 2.92E-02 |
| LOC129310330 | 129310330 | transcription factor bHLH149-like | 1.8 | 1.63E-03 |
| LOC129290588 | 129290588 | probable serine/threonine-protein kinase PBL1 | 1.8 | 1.72E-03 |
| LOC129295976 | 129295976 | alpha-soluble NSF attachment protein 2-like | 1.8 | 4.03E-02 |
| LOC129309609 | 129309609 | uncharacterized LOC129309609 | 1.8 | 1.15E-05 |
| LOC129311970 | 129311970 | uncharacterized LOC129311970 | 1.8 | 9.69E-03 |
| LOC129310057 | 129310057 | protein HEAT INTOLERANT 4 | 1.8 | 2.08E-04 |
| LOC129301423 | 129301423 | la-related protein 1C-like | 1.8 | 3.86E-02 |
| LOC129320886 | 129320886 | uncharacterized LOC129320886 | 1.8 | 1.10E-03 |
| LOC129322793 | 129322793 | mannan endo-1,4-beta-mannosidase 7 | 1.8 | 3.06E-03 |
| LOC129310063 | 129310063 | uncharacterized LOC129310063 | 1.8 | 4.27E-02 |
| LOC129322142 | 129322142 | calcium-transporting ATPase 2, plasma membrane-type-like | 1.8 | 1.64E-05 |
| LOC129304053 | 129304053 | E3 ubiquitin-protein ligase WAV3 | 1.8 | 1.12E-04 |
| LOC129318505 | 129318505 | 2-alkenal reductase (NADP(+)-dependent)-like | 1.8 | 2.14E-02 |
| LOC129311534 | 129311534 | vesicle-associated protein 2-1-like | 1.8 | 1.47E-03 |
| LOC129323053 | 129323053 | FCS-Like Zinc finger 1 | 1.8 | 1.01E-04 |
| LOC129320437 | 129320437 | uncharacterized LOC129320437 | 1.8 | 5.10E-03 |
| LOC129318260 | 129318260 | uncharacterized LOC129318260 | 1.8 | 4.97E-03 |
| LOC129311757 | 129311757 | uncharacterized LOC129311757 | 1.8 | 2.81E-02 |
| LOC129302059 | 129302059 | uncharacterized LOC129302059 | 1.8 | 2.75E-03 |
| LOC129312104 | 129312104 | histone H2A-like | 1.8 | 2.84E-02 |
| LOC129289382 | 129289382 | C-terminal binding protein AN-like | 1.8 | 1.33E-02 |
| LOC129316010 | 129316010 | uncharacterized LOC129316010 | 1.8 | 3.00E-03 |
| LOC129294126 | 129294126 | growth-regulating factor 7-like | 1.8 | 1.40E-02 |
| LOC129321301 | 129321301 | pectinesterase-like | 1.8 | 2.50E-02 |
| LOC129292843 | 129292843 | cystinosin homolog | 1.8 | 1.72E-02 |
| LOC129316212 | 129316212 | ATP synthase small subunit 6-A, mitochondrial-like | 1.8 | 2.82E-02 |
| LOC129291893 | 129291893 | LIMR family protein At5g01460 | 1.8 | 1.53E-02 |
| LOC129312399 | 129312399 | uncharacterized LOC129312399 | 1.8 | 1.50E-03 |
| LOC129298001 | 129298001 | uncharacterized LOC129298001 | 1.8 | 1.66E-02 |
| LOC129318445 | 129318445 | F-box protein At2g26850-like | 1.8 | 7.68E-05 |
| LOC129313629 | 129313629 | probable BOI-related E3 ubiquitin-protein ligase 3 | 1.8 | 4.24E-03 |
| LOC129315546 | 129315546 | UDP-glycosyltransferase 92A1 | 1.8 | 1.07E-04 |
| LOC129314689 | 129314689 | senescence-associated carboxylesterase 101-like | 1.8 | 2.89E-03 |
| LOC129294219 | 129294219 | uncharacterized LOC129294219 | 1.7 | 3.84E-02 |
| LOC129288483 | 129288483 | O-fucosyltransferase 19 | 1.7 | 9.09E-05 |
| LOC129309023 | 129309023 | uncharacterized LOC129309023 | 1.7 | 1.36E-02 |
| LOC129303729 | 129303729 | uncharacterized LOC129303729 | 1.7 | 1.02E-03 |
| LOC129293969 | 129293969 | scarecrow-like protein 8 | 1.7 | 2.70E-03 |
| LOC129313446 | 129313446 | uncharacterized LOC129313446 | 1.7 | 1.21E-07 |
| LOC129294197 | 129294197 | uncharacterized LOC129294197 | 1.7 | 2.44E-03 |
| LOC129316124 | 129316124 | uncharacterized LOC129316124 | 1.7 | 1.03E-03 |
| LOC129319278 | 129319278 | sulfite exporter TauE/SafE family protein 4 | 1.7 | 1.30E-03 |
| LOC129321456 | 129321456 | reticulon-like protein B2 | 1.7 | 8.79E-05 |
| LOC129305685 | 129305685 | uncharacterized LOC129305685 | 1.7 | 2.28E-03 |

| | | | | |
|---|---|---|---|---|
| LOC129319652 | 129319652 | homocysteine S-methyltransferase 3-like | 1.7 | 2.34E-04 |
| LOC129313819 | 129313819 | mediator of RNA polymerase II transcription subunit 22a-like | 1.7 | 1.84E-04 |
| LOC129298906 | 129298906 | chorismate synthase, chloroplastic-like | 1.7 | 2.49E-02 |
| LOC129321406 | 129321406 | protein CDI-like | 1.7 | 2.63E-02 |
| LOC129316148 | 129316148 | glutaredoxin-C9-like | 1.7 | 1.54E-02 |
| LOC129304984 | 129304984 | NADH--cytochrome b5 reductase 1-like | 1.7 | 2.24E-02 |
| LOC129315693 | 129315693 | myosin-2 | 1.7 | 1.98E-02 |
| LOC129308751 | 129308751 | uncharacterized LOC129308751 | 1.7 | 3.69E-02 |
| LOC129313482 | 129313482 | transcription factor E2FB-like | 1.7 | 2.13E-04 |
| LOC129292786 | 129292786 | cold-regulated 413 plasma membrane protein 1 | 1.7 | 4.63E-02 |
| LOC129322233 | 129322233 | zinc finger A20 and AN1 domain-containing stress-associated p | 1.7 | 1.57E-03 |
| LOC129308669 | 129308669 | probable magnesium transporter NIPA1 | 1.7 | 8.62E-04 |
| LOC129290395 | 129290395 | vestitone reductase-like | 1.7 | 1.12E-03 |
| LOC129293401 | 129293401 | gallate 1-beta-glucosyltransferase-like | 1.7 | 3.23E-02 |
| LOC129309299 | 129309299 | F-box/kelch-repeat protein SKIP6 | 1.7 | 1.02E-02 |
| LOC129284317 | 129284317 | disease resistance protein RPV1-like | 1.7 | 5.50E-03 |
| LOC129301693 | 129301693 | jasmonate-induced oxygenase 4 | 1.7 | 7.28E-03 |
| LOC129313153 | 129313153 | chaperone protein dnaJ C76, chloroplastic | 1.7 | 1.15E-02 |
| LOC129303910 | 129303910 | indole-3-acetic acid-induced protein ARG7 | 1.7 | 9.98E-03 |
| LOC129319073 | 129319073 | splicing factor 3B subunit 6-like protein | 1.7 | 4.12E-02 |
| LOC129314263 | 129314263 | small polypeptide DEVIL 10 | 1.7 | 4.35E-03 |
| LOC129294229 | 129294229 | uncharacterized LOC129294229 | 1.7 | 1.07E-03 |
| LOC129309220 | 129309220 | nuclear transcription factor Y subunit C-1-like | 1.7 | 9.74E-04 |
| LOC129320462 | 129320462 | manganese-dependent ADP-ribose/CDP-alcohol diphosphatase | 1.7 | 4.95E-02 |
| LOC129285710 | 129285710 | uncharacterized LOC129285710 | 1.7 | 1.38E-02 |
| LOC129294184 | 129294184 | uncharacterized LOC129294184 | 1.7 | 1.05E-02 |
| LOC129317592 | 129317592 | cellulose synthase-like protein E6 | 1.7 | 1.66E-02 |
| LOC129292780 | 129292780 | methylthioribose kinase-like | 1.7 | 1.40E-02 |
| LOC129307647 | 129307647 | transcription factor MYB1-like | 1.7 | 1.11E-02 |
| LOC129295105 | 129295105 | transcription factor MTB1-like | 1.7 | 1.01E-02 |
| LOC129314686 | 129314686 | uncharacterized LOC129314686 | 1.7 | 1.31E-04 |
| LOC129293437 | 129293437 | histone H4 | 1.7 | 2.06E-02 |
| LOC129319803 | 129319803 | putative clathrin assembly protein At4g40080 | 1.7 | 3.26E-03 |
| LOC129294109 | 129294109 | F-box protein SKP2A-like | 1.7 | 1.89E-05 |
| LOC129296569 | 129296569 | probable GTP diphosphokinase RSH3, chloroplastic | 1.7 | 5.63E-03 |
| LOC129321245 | 129321245 | acid beta-fructofuranosidase-like | 1.7 | 2.37E-02 |
| LOC129309634 | 129309634 | uncharacterized LOC129309634 | 1.7 | 4.69E-02 |
| LOC129306990 | 129306990 | probable inositol transporter 2 | 1.7 | 1.14E-03 |
| LOC129316690 | 129316690 | uncharacterized LOC129316690 | 1.7 | 5.65E-03 |
| LOC129321644 | 129321644 | probable xyloglucan endotransglucosylase/hydrolase protein 30 | 1.7 | 1.29E-02 |
| LOC129294401 | 129294401 | serine/threonine-protein kinase CTR1-like | 1.7 | 1.54E-02 |
| LOC129315262 | 129315262 | uncharacterized protein At4g28440-like | 1.7 | 2.95E-02 |
| LOC129303425 | 129303425 | uncharacterized LOC129303425 | 1.7 | 3.98E-05 |
| LOC129309869 | 129309869 | kinesin-like protein KIN-12B | 1.7 | 2.52E-02 |
| LOC129318500 | 129318500 | nuclear transcription factor Y subunit A-1-like | 1.7 | 3.01E-06 |
| LOC129293258 | 129293258 | U-box domain-containing protein 45-like | 1.7 | 5.52E-03 |
| LOC129285231 | 129285231 | trihelix transcription factor DF1 | 1.7 | 3.45E-04 |
| LOC129303222 | 129303222 | protein kinase STUNTED-like | 1.7 | 1.10E-02 |

| | | | | |
|---|---|---|---|---|
| LOC129301826 | 129301826 | pathogenesis-related thaumatin-like protein 3.5 | 1.7 | 4.05E-02 |
| LOC129302346 | 129302346 | uncharacterized LOC129302346 | 1.7 | 5.42E-03 |
| LOC129322180 | 129322180 | aspartyl protease family protein 1 | 1.7 | 7.35E-03 |
| LOC129319511 | 129319511 | anaphase-promoting complex subunit 8 | 1.7 | 1.13E-04 |
| LOC129292259 | 129292259 | vacuolar protein sorting-associated protein 2 homolog 1-like | 1.7 | 3.47E-03 |
| LOC129303539 | 129303539 | cyclin-D4-1-like | 1.7 | 2.18E-02 |
| LOC129285069 | 129285069 | receptor-like serine/threonine-protein kinase At2g45590 | 1.7 | 1.26E-02 |
| LOC129315265 | 129315265 | vicilin-like seed storage protein At2g18540 | 1.7 | 3.63E-04 |
| LOC129321677 | 129321677 | uncharacterized LOC129321677 | 1.7 | 2.93E-02 |
| LOC129310999 | 129310999 | putative phospholipid-transporting ATPase 9 | 1.7 | 8.09E-04 |
| LOC129322637 | 129322637 | DNA repair endonuclease UVH1 | 1.7 | 4.66E-03 |
| LOC129304180 | 129304180 | scarecrow-like protein 3 | 1.7 | 1.82E-05 |
| LOC129305628 | 129305628 | uncharacterized LOC129305628 | 1.7 | 2.99E-02 |
| LOC129295888 | 129295888 | growth-regulating factor 5-like | 1.7 | 2.30E-02 |
| LOC129316255 | 129316255 | B-box zinc finger protein 22 | 1.7 | 3.27E-02 |
| LOC129305255 | 129305255 | GDSL esterase/lipase At5g45950 | 1.7 | 4.64E-02 |
| LOC129306509 | 129306509 | uncharacterized protein At5g19025-like | 1.7 | 3.92E-02 |
| LOC129284444 | 129284444 | protein WVD2-like 1 | 1.7 | 4.10E-03 |
| LOC129285835 | 129285835 | probable calcium-binding protein CML35 | 1.7 | 6.07E-03 |
| LOC129313635 | 129313635 | protein CYSTEINE-RICH TRANSMEMBRANE MODULE 7- | 1.7 | 8.19E-03 |
| LOC129305551 | 129305551 | succinate dehydrogenase [ubiquinone] iron-sulfur subunit 2, mit | 1.7 | 1.51E-02 |
| LOC129306115 | 129306115 | adenylosuccinate synthetase 2, chloroplastic-like | 1.7 | 1.09E-03 |
| LOC129320714 | 129320714 | very-long-chain aldehyde decarbonylase GL1-1-like | 1.7 | 4.72E-05 |
| LOC129312787 | 129312787 | transcription factor bHLH153 | 1.7 | 2.20E-04 |
| LOC129293571 | 129293571 | protein S40-5 | 1.7 | 1.06E-02 |
| LOC129316714 | 129316714 | probable E3 ubiquitin-protein ligase RHC2A | 1.7 | 3.60E-04 |
| LOC129297773 | 129297773 | 50S ribosomal protein L2, chloroplastic | 1.7 | 2.41E-02 |
| LOC129304944 | 129304944 | uncharacterized protein At5g39570-like | 1.7 | 3.32E-03 |
| LOC129291386 | 129291386 | phosphatidylcholine:diacylglycerol cholinephosphotransferase 1- | 1.7 | 3.07E-02 |
| LOC129314993 | 129314993 | protein DJ-1 homolog B-like | 1.7 | 4.70E-04 |
| LOC129317806 | 129317806 | uncharacterized LOC129317806 | 1.7 | 5.05E-04 |
| LOC129315061 | 129315061 | uncharacterized LOC129315061 | 1.7 | 1.24E-02 |
| LOC129305302 | 129305302 | auxin-responsive protein IAA11-like | 1.7 | 2.15E-03 |
| LOC129315520 | 129315520 | mitogen-activated protein kinase 19-like | 1.7 | 6.06E-03 |
| LOC129303142 | 129303142 | dihydroflavonol 4-reductase-like | 1.7 | 4.20E-02 |
| LOC129289889 | 129289889 | uncharacterized LOC129289889 | 1.7 | 2.23E-04 |
| LOC129303997 | 129303997 | polyamine oxidase 2-like | 1.7 | 3.82E-02 |
| LOC129318326 | 129318326 | UDP-glycosyltransferase 87A1-like | 1.7 | 4.20E-02 |
| LOC129294880 | 129294880 | ATP synthase subunit alpha, mitochondrial | 1.7 | 2.67E-02 |
| LOC129307895 | 129307895 | uncharacterized LOC129307895 | 1.7 | 2.14E-02 |
| LOC129312575 | 129312575 | zinc finger A20 and AN1 domain-containing stress-associated p | 1.7 | 1.64E-04 |
| LOC129309214 | 129309214 | probable methyltransferase PMT26 | 1.7 | 3.73E-03 |
| LOC129293160 | 129293160 | eukaryotic peptide chain release factor subunit 1-3-like | 1.7 | 2.77E-03 |
| LOC129291685 | 129291685 | uncharacterized LOC129291685 | 1.7 | 1.87E-02 |
| LOC129322770 | 129322770 | uncharacterized LOC129322770 | 1.7 | 9.27E-03 |
| LOC129286653 | 129286653 | glutelin type-A 2-like | 1.6 | 4.08E-02 |
| LOC129291265 | 129291265 | ESCRT-related protein CHMP1B | 1.6 | 1.92E-02 |
| LOC129310818 | 129310818 | CLAVATA3/ESR (CLE)-related protein 43-like | 1.6 | 1.71E-02 |

| | | | | |
|---|---|---|---|---|
| LOC129313449 | 129313449 | uncharacterized LOC129313449 | 1.6 | 2.19E-02 |
| LOC129309673 | 129309673 | myb-related protein 306-like | 1.6 | 1.56E-02 |
| LOC129306315 | 129306315 | UDP-arabinose 4-epimerase 1-like | 1.6 | 2.84E-02 |
| LOC129303511 | 129303511 | PHD finger-like domain-containing protein 5A | 1.6 | 4.14E-03 |
| LOC129322941 | 129322941 | granule-bound starch synthase 1, chloroplastic/amyloplastic-like | 1.6 | 7.01E-03 |
| LOC129311105 | 129311105 | fra a 1-associated protein-like | 1.6 | 8.54E-03 |
| LOC129288007 | 129288007 | UDP-glycosyltransferase 74G1-like | 1.6 | 4.86E-02 |
| LOC129319723 | 129319723 | UDP-arabinopyranose mutase 3-like | 1.6 | 2.08E-03 |
| LOC129304356 | 129304356 | chloride channel protein CLC-c-like | 1.6 | 5.30E-04 |
| LOC129292289 | 129292289 | probable transcriptional regulator SLK2 | 1.6 | 2.39E-08 |
| LOC129294853 | 129294853 | NADH-ubiquinone oxidoreductase chain 4-like | 1.6 | 3.97E-02 |
| LOC129320925 | 129320925 | uncharacterized LOC129320925 | 1.6 | 1.16E-03 |
| LOC129320681 | 129320681 | membrane-anchored ubiquitin-fold protein 3-like | 1.6 | 2.16E-02 |
| LOC129322772 | 129322772 | calmodulin-binding protein 25-like | 1.6 | 2.94E-03 |
| LOC129312844 | 129312844 | sm-like protein LSM7 | 1.6 | 1.46E-02 |
| LOC129286771 | 129286771 | metacaspase-3-like | 1.6 | 4.52E-02 |
| LOC129312427 | 129312427 | protein STRUBBELIG-RECEPTOR FAMILY 3-like | 1.6 | 3.80E-03 |
| LOC129291427 | 129291427 | vestitone reductase-like | 1.6 | 8.28E-03 |
| LOC129315819 | 129315819 | uncharacterized LOC129315819 | 1.6 | 1.22E-03 |
| LOC129305950 | 129305950 | protein NETWORKED 3A-like | 1.6 | 2.53E-03 |
| LOC129314287 | 129314287 | 60S ribosomal protein L12-like | 1.6 | 5.85E-03 |
| LOC129320898 | 129320898 | AP2-like ethylene-responsive transcription factor AIL5 | 1.6 | 2.64E-05 |
| LOC129292612 | 129292612 | homeobox-leucine zipper protein ANTHOCYANINLESS 2-like | 1.6 | 3.09E-02 |
| LOC129303455 | 129303455 | uncharacterized LOC129303455 | 1.6 | 7.25E-03 |
| LOC129296005 | 129296005 | uncharacterized LOC129296005 | 1.6 | 9.42E-03 |
| LOC129313365 | 129313365 | phenylalanine ammonia-lyase-like | 1.6 | 4.31E-02 |
| LOC129323099 | 129323099 | uncharacterized LOC129323099 | 1.6 | 8.13E-03 |
| LOC129287149 | 129287149 | uncharacterized LOC129287149 | 1.6 | 2.40E-02 |
| LOC129316362 | 129316362 | uncharacterized LOC129316362 | 1.6 | 2.45E-02 |
| LOC129304144 | 129304144 | uncharacterized LOC129304144 | 1.6 | 3.93E-02 |
| LOC129286595 | 129286595 | glycine-rich protein 2-like | 1.6 | 7.80E-03 |
| LOC129322006 | 129322006 | bet1-like SNARE 1-1 | 1.6 | 1.19E-02 |
| LOC129287679 | 129287679 | mitochondrial import inner membrane translocase subunit TIM2 | 1.6 | 1.76E-02 |
| LOC129301668 | 129301668 | strigolactone esterase RMS3 | 1.6 | 3.73E-03 |
| LOC129300419 | 129300419 | plant UBX domain-containing protein 4-like | 1.6 | 4.14E-02 |
| LOC129323094 | 129323094 | uncharacterized LOC129323094 | 1.6 | 2.32E-02 |
| LOC129295588 | 129295588 | uncharacterized LOC129295588 | 1.6 | 5.21E-03 |
| LOC129322353 | 129322353 | probable polygalacturonase | 1.6 | 1.05E-03 |
| LOC129318639 | 129318639 | uncharacterized membrane protein At3g27390 | 1.6 | 1.38E-02 |
| LOC129309819 | 129309819 | RING-H2 finger protein ATL2-like | 1.6 | 3.67E-03 |
| LOC129303734 | 129303734 | HMG-Y-related protein A | 1.6 | 3.38E-03 |
| LOC129287302 | 129287302 | uncharacterized LOC129287302 | 1.6 | 6.43E-03 |
| LOC129318536 | 129318536 | uncharacterized LOC129318536 | 1.6 | 1.17E-05 |
| LOC129304502 | 129304502 | uncharacterized LOC129304502 | 1.6 | 2.39E-02 |
| LOC129298853 | 129298853 | DYRK-family kinase pom1-like | 1.6 | 2.66E-02 |
| LOC129306297 | 129306297 | peroxisomal adenine nucleotide carrier 1 | 1.6 | 4.79E-02 |
| LOC129295871 | 129295871 | uncharacterized LOC129295871 | 1.6 | 2.39E-02 |
| LOC129304363 | 129304363 | probable polyol transporter 4 | 1.6 | 1.51E-03 |

| | | | | |
|---|---|---|---|---|
| LOC129319005 | 129319005 | cytochrome P450 83B1-like | 1.6 | 4.77E-02 |
| LOC129318307 | 129318307 | WRKY transcription factor WRKY24-like | 1.6 | 3.96E-02 |
| LOC129322286 | 129322286 | PI-PLC X domain-containing protein At5g67130-like | 1.6 | 7.38E-03 |
| LOC129286776 | 129286776 | uncharacterized LOC129286776 | 1.6 | 1.47E-02 |
| LOC129296302 | 129296302 | wall-associated receptor kinase-like 22 | 1.6 | 4.24E-02 |
| LOC129285743 | 129285743 | probable zinc metallopeptidase EGY3, chloroplastic | 1.6 | 8.90E-04 |
| LOC129290797 | 129290797 | GATA transcription factor 16-like | 1.6 | 3.39E-02 |
| LOC129296517 | 129296517 | staphylococcal-like nuclease CAN2 | 1.6 | 2.57E-04 |
| LOC129298535 | 129298535 | zinc finger protein ZAT10-like | 1.6 | 3.07E-03 |
| LOC129309839 | 129309839 | uncharacterized LOC129309839 | 1.6 | 5.78E-03 |
| LOC129288796 | 129288796 | uncharacterized LOC129288796 | 1.6 | 1.43E-02 |
| LOC129303223 | 129303223 | phosphatidylinositol/phosphatidylcholine transfer protein SFH1[illegible] | 1.6 | 1.12E-03 |
| LOC129308786 | 129308786 | pyrophosphate-energized vacuolar membrane proton pump-like | 1.6 | 1.08E-02 |
| LOC129308619 | 129308619 | uncharacterized LOC129308619 | 1.6 | 1.34E-03 |
| LOC129301369 | 129301369 | peroxisomal and mitochondrial division factor 2-like | 1.6 | 1.79E-02 |
| LOC129291102 | 129291102 | clavaminate synthase-like protein At3g21360 | 1.6 | 1.86E-02 |
| LOC129310978 | 129310978 | nodulin-related protein 1-like | 1.6 | 4.45E-03 |
| LOC129296934 | 129296934 | shewanella-like protein phosphatase 2 | 1.6 | 8.06E-03 |
| LOC129296032 | 129296032 | uncharacterized LOC129296032 | 1.6 | 2.08E-02 |
| LOC129317981 | 129317981 | probable magnesium transporter NIPA6 | 1.6 | 1.72E-02 |
| LOC129318577 | 129318577 | putative cyclin-A3-1 | 1.6 | 1.40E-03 |
| LOC129311173 | 129311173 | trihelix transcription factor GT-3b-like | 1.6 | 5.10E-03 |
| LOC129287623 | 129287623 | ras-related protein Rab7 | 1.6 | 1.01E-02 |
| LOC129321801 | 129321801 | ubiquitin carboxyl-terminal hydrolase 21-like | 1.6 | 1.62E-03 |
| LOC129321095 | 129321095 | uncharacterized LOC129321095 | 1.6 | 2.50E-02 |
| LOC129317838 | 129317838 | zinc finger protein CONSTANS-LIKE 4 | 1.6 | 8.95E-03 |
| LOC129294094 | 129294094 | E3 ubiquitin-protein ligase MBR2-like | 1.6 | 4.74E-03 |
| LOC129286877 | 129286877 | myb family transcription factor EFM-like | 1.6 | 8.40E-04 |
| LOC129309394 | 129309394 | serine/threonine-protein kinase AFC2 | 1.6 | 8.40E-06 |
| LOC129314208 | 129314208 | zinc finger A20 and AN1 domain-containing stress-associated p | 1.6 | 3.87E-02 |
| LOC129306458 | 129306458 | ABC transporter G family member 22-like | 1.6 | 1.39E-03 |
| LOC129305737 | 129305737 | actin-101-like | 1.5 | 9.42E-05 |
| LOC129304710 | 129304710 | BTB/POZ and MATH domain-containing protein 2-like | 1.5 | 6.16E-04 |
| LOC129314977 | 129314977 | E3 ubiquitin-protein ligase makorin-like | 1.5 | 8.29E-03 |
| LOC129305251 | 129305251 | transcription factor ICE1-like | 1.5 | 1.30E-02 |
| LOC129318726 | 129318726 | small ubiquitin-related modifier 2 | 1.5 | 1.43E-02 |
| LOC129306803 | 129306803 | vacuolar protein sorting-associated protein 2 homolog 2-like | 1.5 | 4.34E-02 |
| LOC129297689 | 129297689 | protein TIFY 6B-like | 1.5 | 1.94E-03 |
| LOC129295785 | 129295785 | ADP-ribosylation factor 1 | 1.5 | 6.51E-03 |
| LOC129303555 | 129303555 | putative MYST-like histone acetyltransferase 1 | 1.5 | 1.54E-02 |
| LOC129302974 | 129302974 | uncharacterized LOC129302974 | 1.5 | 3.39E-03 |
| LOC129300693 | 129300693 | membrin-11-like | 1.5 | 1.69E-02 |
| LOC129311504 | 129311504 | homeobox-leucine zipper protein HAT5 | 1.5 | 7.16E-04 |
| LOC129311371 | 129311371 | uncharacterized LOC129311371 | 1.5 | 3.72E-02 |
| LOC129319933 | 129319933 | E3 ubiquitin-protein ligase MPSR1-like | 1.5 | 2.47E-02 |
| LOC129302327 | 129302327 | cytochrome P450 711A1 | 1.5 | 1.76E-04 |
| LOC129322392 | 129322392 | kinesin-like protein KIN-14R | 1.5 | 1.49E-03 |
| LOC129306576 | 129306576 | uncharacterized LOC129306576 | 1.5 | 2.03E-03 |

| | | | | |
|---|---|---|---|---|
| LOC129319326 | 129319326 | dof zinc finger protein DOF5.4-like | 1.5 | 1.64E-02 |
| LOC129320203 | 129320203 | uncharacterized LOC129320203 | 1.5 | 4.03E-02 |
| LOC129320260 | 129320260 | enolase | 1.5 | 4.26E-03 |
| LOC129313104 | 129313104 | uncharacterized LOC129313104 | 1.5 | 6.88E-03 |
| LOC129303692 | 129303692 | transmembrane emp24 domain-containing protein p24beta2-like | 1.5 | 4.57E-02 |
| LOC129306981 | 129306981 | cytochrome b-c1 complex subunit 6-1, mitochondrial | 1.5 | 4.75E-03 |
| LOC129313480 | 129313480 | uncharacterized LOC129313480 | 1.5 | 2.25E-02 |
| LOC129294011 | 129294011 | protein DETOXIFICATION 9-like | 1.5 | 2.04E-04 |
| LOC129293193 | 129293193 | peptidyl-prolyl cis-trans isomerase 1 | 1.5 | 1.58E-02 |
| LOC129309440 | 129309440 | phosphatidylinositol 4-kinase gamma 8-like | 1.5 | 1.70E-02 |
| LOC129319413 | 129319413 | uncharacterized LOC129319413 | 1.5 | 2.55E-02 |
| LOC129294769 | 129294769 | uncharacterized LOC129294769 | 1.5 | 1.16E-04 |
| LOC129318678 | 129318678 | 1-aminocyclopropane-1-carboxylate oxidase-like | 1.5 | 2.35E-02 |
| LOC129306391 | 129306391 | nuclear transcription factor Y subunit B-10-like | 1.5 | 9.88E-04 |
| LOC129288881 | 129288881 | uncharacterized LOC129288881 | 1.5 | 1.36E-02 |
| LOC129307064 | 129307064 | uncharacterized LOC129307064 | 1.5 | 1.27E-03 |
| LOC129309260 | 129309260 | calcium-dependent mitochondrial ATP-magnesium/phosphate ca | 1.5 | 6.96E-03 |
| LOC129309316 | 129309316 | probable xyloglucan galactosyltransferase GT14 | 1.5 | 3.83E-02 |
| LOC129309418 | 129309418 | diacylglycerol O-acyltransferase 1A-like | 1.5 | 3.45E-02 |
| LOC129310062 | 129310062 | homeobox-leucine zipper protein HAT5-like | 1.5 | 3.39E-03 |
| LOC129288816 | 129288816 | uncharacterized LOC129288816 | 1.5 | 1.47E-02 |
| LOC129315499 | 129315499 | phosphoenolpyruvate carboxylase, housekeeping isozyme | 1.5 | 4.14E-02 |
| LOC129293626 | 129293626 | mannose-6-phosphate isomerase 1-like | 1.5 | 2.20E-02 |
| LOC129317839 | 129317839 | uncharacterized protein At1g32220, chloroplastic | 1.5 | 2.32E-02 |
| LOC129318571 | 129318571 | CBL-interacting serine/threonine-protein kinase 5-like | 1.5 | 1.48E-02 |
| LOC129304982 | 129304982 | serine decarboxylase-like | 1.5 | 1.03E-02 |
| LOC129319063 | 129319063 | probable receptor-like protein kinase At2g39360 | 1.5 | 3.37E-04 |
| LOC129304591 | 129304591 | ricin B-like lectin EULS3 | 1.5 | 3.81E-04 |
| LOC129293543 | 129293543 | 60S ribosomal protein L36-3-like | 1.5 | 1.41E-02 |
| LOC129304042 | 129304042 | heavy metal-associated isoprenylated plant protein 6-like | 1.5 | 1.90E-04 |
| LOC129290981 | 129290981 | calcineurin B-like protein 10 | 1.5 | 8.29E-03 |
| LOC129290196 | 129290196 | uncharacterized LOC129290196 | 1.5 | 1.31E-04 |
| LOC129307966 | 129307966 | probable protein phosphatase 2C 13 | 1.5 | 1.49E-03 |
| LOC129294357 | 129294357 | protein NTM1-like 9 | 1.5 | 3.58E-04 |
| LOC129320616 | 129320616 | S-adenosylmethionine synthase | 1.5 | 1.33E-02 |
| LOC129316131 | 129316131 | uncharacterized LOC129316131 | 1.5 | 3.72E-02 |
| LOC129314212 | 129314212 | early nodulin-like protein 1 | 1.5 | 3.47E-02 |
| LOC129309076 | 129309076 | uncharacterized LOC129309076 | 1.5 | 7.57E-04 |
| LOC129291759 | 129291759 | uncharacterized LOC129291759 | 1.5 | 8.76E-04 |
| LOC129313813 | 129313813 | uncharacterized protein At1g08160 | 1.5 | 3.23E-02 |
| LOC129288387 | 129288387 | eukaryotic translation initiation factor 1A-like | 1.5 | 2.97E-02 |
| LOC129317847 | 129317847 | strigolactone esterase RMS3-like | 1.5 | 3.90E-03 |
| LOC129291423 | 129291423 | uncharacterized LOC129291423 | 1.5 | 1.97E-02 |
| LOC129323204 | 129323204 | DCD domain-containing protein NRP-B | 1.5 | 1.63E-02 |
| LOC129303202 | 129303202 | ultraviolet-B receptor UVR8 | 1.5 | 1.34E-02 |
| LOC129294177 | 129294177 | uncharacterized LOC129294177 | 1.5 | 1.77E-04 |
| LOC129305901 | 129305901 | probable DNA helicase MCM8 | 1.5 | 2.09E-02 |
| LOC129319252 | 129319252 | GDP-mannose transporter GONST1-like | 1.5 | 6.12E-03 |

| | | | | |
|---|---|---|---|---|
| LOC129285725 | 129285725 | tRNA (guanine-N(7)-)-methyltransferase | 1.5 | 1.95E-02 |
| LOC129303095 | 129303095 | proline transporter 2-like | 1.5 | 4.72E-02 |
| LOC129313502 | 129313502 | autophagy-related protein 8C-like | 1.5 | 2.93E-02 |
| LOC129305021 | 129305021 | putative E3 ubiquitin-protein ligase RING1a | 1.5 | 1.31E-04 |
| LOC129306669 | 129306669 | uncharacterized LOC129306669 | 1.5 | 3.29E-03 |
| LOC129311915 | 129311915 | uncharacterized LOC129311915 | 1.5 | 1.33E-02 |
| LOC129305959 | 129305959 | cystinosin homolog | 1.5 | 1.83E-02 |
| LOC129311142 | 129311142 | protein CHROMATIN REMODELING 19 | 1.5 | 5.53E-03 |
| LOC129287525 | 129287525 | geraniol 8-hydroxylase-like | 1.5 | 4.36E-02 |
| LOC129318581 | 129318581 | protein REVEILLE 1-like | 1.5 | 1.44E-02 |
| LOC129290770 | 129290770 | uncharacterized LOC129290770 | 1.5 | 5.30E-03 |
| LOC129320181 | 129320181 | probable histone H2A variant 3 | 1.5 | 4.81E-03 |
| LOC129306000 | 129306000 | probable phospholipid-transporting ATPase 8 | 1.5 | 2.71E-04 |
| LOC129322876 | 129322876 | upstream activation factor subunit spp27-like | 1.5 | 8.65E-03 |
| LOC129321267 | 129321267 | transcription factor BIM2-like | 1.5 | 2.99E-02 |
| LOC129309221 | 129309221 | probable galactinol--sucrose galactosyltransferase 2 | 1.5 | 1.28E-03 |
| LOC129285699 | 129285699 | calmodulin-binding transcription activator 2-like | 1.5 | 4.91E-04 |
| LOC129290559 | 129290559 | dof zinc finger protein DOF3.1-like | 1.5 | 3.60E-02 |
| LOC129308241 | 129308241 | iron-sulfur assembly protein IscA, chloroplastic | 1.5 | 2.65E-02 |
| LOC129293010 | 129293010 | protein REVEILLE 6 | 1.5 | 3.54E-04 |
| LOC129320341 | 129320341 | ABC transporter I family member 20 | 1.5 | 8.41E-05 |
| LOC129318319 | 129318319 | protein FATTY ACID EXPORT 5-like | 1.5 | 2.51E-02 |
| LOC129292868 | 129292868 | uncharacterized LOC129292868 | 1.4 | 1.33E-02 |
| LOC129303995 | 129303995 | uncharacterized LOC129303995 | 1.4 | 3.68E-02 |
| LOC129312404 | 129312404 | uncharacterized LOC129312404 | 1.4 | 2.25E-04 |
| LOC129306593 | 129306593 | 60S ribosomal protein L34-like | 1.4 | 1.84E-02 |
| LOC129294889 | 129294889 | uncharacterized LOC129294889 | 1.4 | 4.25E-02 |
| LOC129293560 | 129293560 | serine/threonine-protein kinase STY46-like | 1.4 | 2.88E-04 |
| LOC129296943 | 129296943 | calcium-dependent protein kinase 10-like | 1.4 | 1.74E-02 |
| LOC129293219 | 129293219 | myb-related protein 308 | 1.4 | 1.20E-02 |
| LOC129320531 | 129320531 | uncharacterized LOC129320531 | 1.4 | 4.96E-02 |
| LOC129312205 | 129312205 | uncharacterized LOC129312205 | 1.4 | 2.33E-02 |
| LOC129322051 | 129322051 | probable aspartic proteinase GIP1 | 1.4 | 2.26E-02 |
| LOC129315618 | 129315618 | protein FANTASTIC FOUR 3-like | 1.4 | 2.73E-02 |
| LOC129293550 | 129293550 | autophagy-related protein 8C-like | 1.4 | 1.94E-02 |
| LOC129316062 | 129316062 | E3 SUMO-protein ligase MMS21 | 1.4 | 2.64E-02 |
| LOC129306643 | 129306643 | bifunctional riboflavin biosynthesis protein RIBA 1, chloroplasti | 1.4 | 7.82E-03 |
| LOC129290457 | 129290457 | protein NUCLEAR FUSION DEFECTIVE 4-like | 1.4 | 2.84E-02 |
| LOC129287989 | 129287989 | acireductone dioxygenase 1 | 1.4 | 3.72E-02 |
| LOC129305187 | 129305187 | polyubiquitin | 1.4 | 8.47E-03 |
| LOC129293544 | 129293544 | uncharacterized LOC129293544 | 1.4 | 1.69E-02 |
| LOC129291206 | 129291206 | caffeoylshikimate esterase | 1.4 | 4.35E-03 |
| LOC129307855 | 129307855 | WUSCHEL-related homeobox 8-like | 1.4 | 2.81E-03 |
| LOC129321536 | 129321536 | putative glucuronosyltransferase PGSIP8 | 1.4 | 1.17E-02 |
| LOC129320455 | 129320455 | homeobox protein knotted-1-like 3 | 1.4 | 1.35E-03 |
| LOC129301505 | 129301505 | uncharacterized LOC129301505 | 1.4 | 1.27E-03 |
| LOC129318693 | 129318693 | uroporphyrinogen-III synthase, chloroplastic | 1.4 | 2.08E-03 |
| LOC129316142 | 129316142 | trifunctional UDP-glucose 4,6-dehydratase/UDP-4-keto-6-deoxy | 1.4 | 1.17E-04 |

| | | | | |
|---|---|---|---|---|
| LOC129302051 | 129302051 | uncharacterized LOC129302051 | 1.4 | 4.64E-02 |
| LOC129305236 | 129305236 | histone H2A | 1.4 | 1.24E-02 |
| LOC129321622 | 129321622 | inositol-tetrakisphosphate 1-kinase 1-like | 1.4 | 2.38E-02 |
| LOC129310584 | 129310584 | metalloendoproteinase 2-MMP | 1.4 | 2.25E-02 |
| LOC129305493 | 129305493 | ras-related protein RABF1 | 1.4 | 1.02E-02 |
| LOC129322892 | 129322892 | 60S ribosomal protein L22-2 | 1.4 | 2.19E-03 |
| LOC129300331 | 129300331 | tetraspanin-8-like | 1.4 | 4.57E-02 |
| LOC129308022 | 129308022 | translation machinery-associated protein 22-like | 1.4 | 1.28E-02 |
| LOC129320157 | 129320157 | 40S ribosomal protein S21-like | 1.4 | 3.23E-02 |
| LOC129286881 | 129286881 | polyadenylate-binding protein RBP45-like | 1.4 | 3.26E-02 |
| LOC129306792 | 129306792 | 40S ribosomal protein S28-1 | 1.4 | 2.00E-02 |
| LOC129292526 | 129292526 | protein RER1B-like | 1.4 | 1.39E-02 |
| LOC129306662 | 129306662 | zinc-finger homeodomain protein 3-like | 1.4 | 4.93E-02 |
| LOC129311991 | 129311991 | uncharacterized LOC129311991 | 1.4 | 1.51E-03 |
| LOC129313624 | 129313624 | uncharacterized LOC129313624 | 1.4 | 3.09E-02 |
| LOC129313586 | 129313586 | uncharacterized protein At5g39865-like | 1.4 | 1.19E-02 |
| LOC129312800 | 129312800 | E3 ubiquitin-protein ligase RMA1H1 | 1.4 | 3.49E-02 |
| LOC129284497 | 129284497 | heavy metal-associated isoprenylated plant protein 21-like | 1.4 | 4.89E-02 |
| LOC129312627 | 129312627 | protein RGF1 INDUCIBLE TRANSCRIPTION FACTOR 1-lik | 1.4 | 4.57E-02 |
| LOC129317152 | 129317152 | ribonuclease 3-like protein 2 | 1.4 | 7.26E-04 |
| LOC129291764 | 129291764 | protein REVEILLE 3-like | 1.4 | 6.31E-03 |
| LOC129291835 | 129291835 | vesicle-associated membrane protein 722 | 1.4 | 4.60E-02 |
| LOC129291014 | 129291014 | serine/threonine-protein kinase AtPK2/AtPK19-like | 1.4 | 3.67E-02 |
| LOC129313410 | 129313410 | probable protein phosphatase 2C 52 | 1.4 | 2.02E-03 |
| LOC129313794 | 129313794 | F-box/kelch-repeat protein At5g60570-like | 1.4 | 1.74E-03 |
| LOC129303308 | 129303308 | ethylene-responsive transcription factor RAP2-7-like | 1.4 | 1.27E-03 |
| LOC129317048 | 129317048 | tryptophan synthase beta chain 1 | 1.4 | 2.61E-02 |
| LOC129308607 | 129308607 | mitogen-activated protein kinase kinase kinase NPK1 | 1.4 | 4.52E-03 |
| LOC129318147 | 129318147 | WEB family protein At5g55860-like | 1.4 | 3.99E-03 |
| LOC129300645 | 129300645 | CAX-interacting protein 4-like | 1.4 | 1.13E-02 |
| LOC129319219 | 129319219 | ethylene-responsive transcription factor ERF060-like | 1.4 | 1.20E-02 |
| LOC129315428 | 129315428 | histone H2A.6 | 1.4 | 2.87E-02 |
| LOC129287283 | 129287283 | protein DSE2 | 1.4 | 3.71E-03 |
| LOC129301817 | 129301817 | autophagy-related protein 18c-like | 1.4 | 6.51E-03 |
| LOC129318841 | 129318841 | chaperone protein dnaJ 11, chloroplastic | 1.4 | 3.18E-02 |
| LOC129321671 | 129321671 | GDSL esterase/lipase At4g10955-like | 1.4 | 9.25E-04 |
| LOC129322248 | 129322248 | glycerol-3-phosphate acyltransferase, chloroplastic-like | 1.4 | 2.57E-02 |
| LOC129285894 | 129285894 | phenylcoumaran benzylic ether reductase POP1 | 1.4 | 9.37E-03 |
| LOC129307683 | 129307683 | pyruvate dehydrogenase (acetyl-transferring) kinase, mitochondr | 1.4 | 3.86E-02 |
| LOC129288336 | 129288336 | RNA polymerase II C-terminal domain phosphatase-like 2 | 1.4 | 1.78E-04 |
| LOC129294650 | 129294650 | uncharacterized LOC129294650 | 1.4 | 3.33E-02 |
| LOC129318406 | 129318406 | uncharacterized LOC129318406 | 1.4 | 4.18E-03 |
| LOC129288426 | 129288426 | transcription factor bHLH68-like | 1.4 | 1.77E-02 |
| LOC129310520 | 129310520 | uncharacterized LOC129310520 | 1.4 | 1.69E-02 |
| LOC129311438 | 129311438 | 60S ribosomal protein L37a | 1.4 | 1.36E-02 |
| LOC129293579 | 129293579 | protein S-acyltransferase 8-like | 1.4 | 2.61E-03 |
| LOC129317078 | 129317078 | PHD finger protein At1g33420-like | 1.4 | 3.07E-03 |
| LOC129314350 | 129314350 | uncharacterized LOC129314350 | 1.4 | 1.31E-03 |

| | | | | |
|---|---|---|---|---|
| LOC129310649 | 129310649 | uncharacterized LOC129310649 | 1.4 | 3.20E-02 |
| LOC129314645 | 129314645 | transcription initiation factor IIA large subunit-like | 1.4 | 2.93E-03 |
| LOC129306668 | 129306668 | protein HEAT-STRESS-ASSOCIATED 32-like | 1.4 | 1.34E-02 |
| LOC129290403 | 129290403 | protein HEADING DATE REPRESSOR 1 | 1.4 | 2.65E-02 |
| LOC129292216 | 129292216 | V-type proton ATPase 16 kDa proteolipid subunit | 1.4 | 3.66E-02 |
| LOC129290709 | 129290709 | uncharacterized LOC129290709 | 1.4 | 5.22E-04 |
| LOC129290424 | 129290424 | metal transporter Nramp6 | 1.4 | 2.51E-02 |
| LOC129314889 | 129314889 | AP2-like ethylene-responsive transcription factor At1g16060 | 1.4 | 2.63E-02 |
| LOC129299339 | 129299339 | lysine-specific demethylase JMJ14-like | 1.4 | 2.71E-02 |
| LOC129311161 | 129311161 | uncharacterized LOC129311161 | 1.4 | 1.81E-03 |
| LOC129306623 | 129306623 | uncharacterized LOC129306623 | 1.4 | 2.51E-02 |
| LOC129310542 | 129310542 | MLO-like protein 8 | 1.3 | 4.53E-02 |
| LOC129307164 | 129307164 | probable hexosyltransferase MUCI70 | 1.3 | 4.07E-02 |
| LOC129306342 | 129306342 | transcription elongation factor 1 homolog | 1.3 | 3.39E-02 |
| LOC129294796 | 129294796 | basic leucine zipper 9 | 1.3 | 4.24E-02 |
| LOC129300959 | 129300959 | F-box protein SKIP14-like | 1.3 | 3.17E-02 |
| LOC129294249 | 129294249 | probable histone H2A variant 3 | 1.3 | 4.00E-02 |
| LOC129293989 | 129293989 | protein GIGANTEA | 1.3 | 3.79E-03 |
| LOC129322345 | 129322345 | protein phosphatase 2C 29-like | 1.3 | 2.88E-03 |
| LOC129313433 | 129313433 | P-loop NTPase domain-containing protein LPA1 homolog 1-like | 1.3 | 1.38E-02 |
| LOC129308213 | 129308213 | probable bifunctional TENA-E protein | 1.3 | 4.76E-02 |
| LOC129322967 | 129322967 | NEP1-interacting protein 1 | 1.3 | 2.40E-02 |
| LOC129304730 | 129304730 | uncharacterized LOC129304730 | 1.3 | 2.34E-02 |
| LOC129300837 | 129300837 | calmodulin-binding protein 60 F-like | 1.3 | 3.92E-02 |
| LOC129311126 | 129311126 | beta-fructofuranosidase, insoluble isoenzyme CWINV3-like | 1.3 | 1.71E-03 |
| LOC129286658 | 129286658 | transmembrane 9 superfamily member 1-like | 1.3 | 3.55E-02 |
| LOC129306737 | 129306737 | importin subunit alpha-like | 1.3 | 3.65E-03 |
| LOC129290619 | 129290619 | UV-stimulated scaffold protein A homolog | 1.3 | 2.22E-03 |
| LOC129314473 | 129314473 | DNA polymerase II subunit B3-1 | 1.3 | 1.46E-02 |
| LOC129305988 | 129305988 | probable linoleate 9S-lipoxygenase 5 | 1.3 | 4.64E-03 |
| LOC129286000 | 129286000 | uncharacterized LOC129286000 | 1.3 | 8.30E-03 |
| LOC129290850 | 129290850 | mitogen-activated protein kinase homolog MMK2-like | 1.3 | 1.15E-02 |
| LOC129305022 | 129305022 | uncharacterized LOC129305022 | 1.3 | 8.90E-05 |
| LOC129305059 | 129305059 | helicase-like transcription factor CHR28 | 1.3 | 2.63E-02 |
| LOC129309681 | 129309681 | protein OXIDATIVE STRESS 3 LIKE 1-like | 1.3 | 1.38E-03 |
| LOC129288840 | 129288840 | replication protein A 32 kDa subunit B | 1.3 | 3.03E-02 |
| LOC129294613 | 129294613 | gamma-glutamylcyclotransferase 2-1-like | 1.3 | 3.05E-03 |
| LOC129320547 | 129320547 | ras-related protein Rab11A-like | 1.3 | 3.15E-03 |
| LOC129293493 | 129293493 | uncharacterized LOC129293493 | 1.3 | 4.75E-02 |
| LOC129314677 | 129314677 | probable calcium-binding protein CML48 | 1.3 | 4.38E-02 |
| LOC129287751 | 129287751 | uncharacterized LOC129287751 | 1.3 | 9.93E-03 |
| LOC129312721 | 129312721 | hydroxyisourate hydrolase | 1.3 | 1.26E-02 |
| LOC129306744 | 129306744 | 40S ribosomal protein S29 | 1.3 | 3.61E-02 |
| LOC129307991 | 129307991 | uncharacterized LOC129307991 | 1.3 | 1.42E-02 |
| LOC129301523 | 129301523 | mitogen-activated protein kinase 20-like | 1.3 | 4.87E-03 |
| LOC129304397 | 129304397 | probable tyrosine-protein phosphatase DSP4 | 1.3 | 1.34E-02 |
| LOC129287729 | 129287729 | nuclear pore complex protein NUP35-like | 1.3 | 2.89E-02 |
| LOC129309176 | 129309176 | RING-box protein 1a | 1.3 | 2.92E-02 |

| | | | | |
|---|---|---|---|---|
| LOC129315463 | 129315463 | C2 domain-containing protein At1g53590-like | 1.3 | 6.23E-03 |
| LOC129314897 | 129314897 | disease resistance protein Roq1-like | 1.3 | 8.53E-03 |
| LOC129317090 | 129317090 | EID1-like F-box protein 2 | 1.3 | 3.78E-02 |
| LOC129321670 | 129321670 | dynamin-related protein 5A | 1.3 | 6.16E-03 |
| LOC129295316 | 129295316 | E3 ubiquitin-protein ligase MBR2-like | 1.3 | 3.48E-02 |
| LOC129322316 | 129322316 | sucrose transport protein SUC3 | 1.3 | 4.86E-02 |
| LOC129297237 | 129297237 | 3-hydroxy-3-methylglutaryl-coenzyme A reductase 1-like | 1.3 | 2.19E-02 |
| LOC129317028 | 129317028 | uncharacterized LOC129317028 | 1.3 | 2.65E-03 |
| LOC129318572 | 129318572 | uncharacterized LOC129318572 | 1.3 | 1.91E-02 |
| LOC129308754 | 129308754 | DNA repair protein XRCC4 | 1.3 | 1.41E-02 |
| LOC129320812 | 129320812 | uncharacterized LOC129320812 | 1.3 | 4.88E-04 |
| LOC129303759 | 129303759 | uncharacterized LOC129303759 | 1.3 | 4.62E-02 |
| LOC129304820 | 129304820 | histone H2A variant 1 | 1.3 | 2.98E-02 |
| LOC129293262 | 129293262 | uncharacterized LOC129293262 | 1.3 | 1.18E-02 |
| LOC129308563 | 129308563 | formin-like protein 18 | 1.3 | 3.51E-04 |
| LOC129315765 | 129315765 | transcription factor bHLH80 | 1.3 | 2.75E-02 |
| LOC129321026 | 129321026 | NAC domain-containing protein 100-like | 1.3 | 5.47E-03 |
| LOC129307259 | 129307259 | binding partner of ACD11 1 | 1.3 | 6.21E-05 |
| LOC129284830 | 129284830 | probable protein phosphatase 2C 63 | 1.3 | 3.28E-02 |
| LOC129311635 | 129311635 | histone-lysine N-methyltransferase CLF-like | 1.3 | 2.76E-02 |
| LOC129303541 | 129303541 | copper-transporting ATPase PAA2, chloroplastic | 1.3 | 2.38E-03 |
| LOC129310043 | 129310043 | probable mitochondrial adenine nucleotide transporter BTL3 | 1.3 | 6.18E-03 |
| LOC129295620 | 129295620 | uncharacterized LOC129295620 | 1.3 | 1.90E-02 |
| LOC129296433 | 129296433 | F-box/kelch-repeat protein OR23-like | 1.3 | 1.57E-02 |
| LOC129294616 | 129294616 | uncharacterized LOC129294616 | 1.3 | 9.15E-03 |
| LOC129318033 | 129318033 | probable protein phosphatase 2C 27 | 1.3 | 5.39E-03 |
| LOC129294601 | 129294601 | L-arabinokinase-like | 1.3 | 6.11E-03 |
| LOC129313324 | 129313324 | protein OBERON 4 | 1.3 | 2.80E-04 |
| LOC129307989 | 129307989 | mitochondrial import receptor subunit TOM5 homolog | 1.3 | 3.43E-02 |
| LOC129311156 | 129311156 | uncharacterized LOC129311156 | 1.3 | 2.49E-03 |
| LOC129284842 | 129284842 | BTB/POZ domain-containing protein At1g63850-like | 1.3 | 5.66E-03 |
| LOC129320243 | 129320243 | hydroxyproline O-galactosyltransferase GALT6-like | 1.3 | 9.74E-04 |
| LOC129310161 | 129310161 | dual specificity protein phosphatase 1 | 1.3 | 1.88E-02 |
| LOC129302975 | 129302975 | probable sugar phosphate/phosphate translocator At3g11320 | 1.3 | 2.13E-02 |
| LOC129317630 | 129317630 | uncharacterized LOC129317630 | 1.3 | 2.77E-03 |
| LOC129320592 | 129320592 | amino acid permease 4 | 1.3 | 1.45E-02 |
| LOC129321381 | 129321381 | protein XAP5 CIRCADIAN TIMEKEEPER | 1.3 | 1.30E-02 |
| LOC129305929 | 129305929 | uncharacterized LOC129305929 | 1.3 | 5.48E-04 |
| LOC129313388 | 129313388 | uncharacterized LOC129313388 | 1.3 | 1.56E-02 |
| LOC129311300 | 129311300 | uncharacterized LOC129311300 | 1.3 | 4.66E-02 |
| LOC129303458 | 129303458 | probable isoaspartyl peptidase/L-asparaginase 2 | 1.3 | 3.66E-02 |
| LOC129297155 | 129297155 | DEK domain-containing chromatin-associated protein 1-like | 1.2 | 2.98E-03 |
| LOC129294226 | 129294226 | uncharacterized LOC129294226 | 1.2 | 9.99E-04 |
| LOC129307057 | 129307057 | 40S ribosomal protein S13 | 1.2 | 1.57E-02 |
| LOC129303228 | 129303228 | probable transcriptional regulator SLK2 | 1.2 | 1.85E-04 |
| LOC129309377 | 129309377 | proteasome subunit beta type-5 | 1.2 | 3.41E-02 |
| LOC129316583 | 129316583 | formin-like protein 14 | 1.2 | 3.15E-03 |
| LOC129315787 | 129315787 | UDP-D-apiose/UDP-D-xylose synthase 2-like | 1.2 | 2.19E-02 |

| | | | | |
|---|---|---|---|---|
| LOC129310349 | 129310349 | receptor protein kinase-like protein ZAR1 | 1.2 | 1.84E-03 |
| LOC129288754 | 129288754 | uncharacterized LOC129288754 | 1.2 | 4.27E-02 |
| LOC129291554 | 129291554 | probable WRKY transcription factor 70 | 1.2 | 4.07E-02 |
| LOC129323121 | 129323121 | phosphatidylinositol 4-kinase gamma 5-like | 1.2 | 9.95E-03 |
| LOC129313415 | 129313415 | uncharacterized LOC129313415 | 1.2 | 4.48E-04 |
| LOC129320743 | 129320743 | rhamnogalacturonan I rhamnosyltransferase 1-like | 1.2 | 1.16E-02 |
| LOC129305919 | 129305919 | SPX domain-containing protein 4 | 1.2 | 1.80E-02 |
| LOC129304127 | 129304127 | soluble starch synthase 1, chloroplastic/amyloplastic | 1.2 | 4.92E-02 |
| LOC129287642 | 129287642 | protein mago nashi homolog | 1.2 | 7.75E-03 |
| LOC129311553 | 129311553 | uncharacterized LOC129311553 | 1.2 | 2.91E-02 |
| LOC129304534 | 129304534 | putative SWI/SNF-related matrix-associated actin-dependent reg | 1.2 | 3.90E-02 |
| LOC129288151 | 129288151 | protein LATE ELONGATED HYPOCOTYL-like | 1.2 | 2.16E-02 |
| LOC129309256 | 129309256 | 40S ribosomal protein S15a | 1.2 | 4.87E-02 |
| LOC129311348 | 129311348 | E3 ubiquitin-protein ligase RING1 | 1.2 | 2.42E-02 |
| LOC129323104 | 129323104 | uncharacterized LOC129323104 | 1.2 | 1.16E-02 |
| LOC129305325 | 129305325 | calmodulin-binding transcription activator 2-like | 1.2 | 1.86E-02 |
| LOC129306546 | 129306546 | phytochrome A-associated F-box protein-like | 1.2 | 3.25E-02 |
| LOC129321408 | 129321408 | soluble inorganic pyrophosphatase | 1.2 | 3.80E-02 |
| LOC129319382 | 129319382 | NAC domain-containing protein 2-like | 1.2 | 4.18E-02 |
| LOC129315111 | 129315111 | probable galactinol--sucrose galactosyltransferase 2 | 1.2 | 3.15E-03 |
| LOC129308127 | 129308127 | probable small nuclear ribonucleoprotein F | 1.2 | 2.99E-02 |
| LOC129313121 | 129313121 | uncharacterized LOC129313121 | 1.2 | 2.71E-02 |
| LOC129318128 | 129318128 | ethylene response sensor 1-like | 1.2 | 2.79E-02 |
| LOC129287788 | 129287788 | UPF0496 protein At3g19330-like | 1.2 | 3.56E-02 |
| LOC129306641 | 129306641 | MACPF domain-containing protein CAD1 | 1.2 | 3.49E-02 |
| LOC129318449 | 129318449 | V-type proton ATPase subunit e1 | 1.2 | 3.46E-02 |
| LOC129301739 | 129301739 | uncharacterized LOC129301739 | 1.2 | 1.05E-02 |
| LOC129292269 | 129292269 | uncharacterized LOC129292269 | 1.2 | 2.23E-02 |
| LOC129305786 | 129305786 | E3 ubiquitin-protein ligase AIRP2-like | 1.2 | 5.10E-03 |
| LOC129303485 | 129303485 | telomere repeat-binding protein 3 | 1.2 | 2.82E-02 |
| LOC129323212 | 129323212 | SKP1-like protein 1A | 1.2 | 4.65E-02 |
| LOC129306332 | 129306332 | CBL-interacting serine/threonine-protein kinase 12 | 1.2 | 3.32E-02 |
| LOC129286554 | 129286554 | calcium-dependent protein kinase 32 | 1.2 | 6.26E-04 |
| LOC129290397 | 129290397 | uncharacterized LOC129290397 | 1.2 | 2.33E-02 |
| LOC129318189 | 129318189 | uncharacterized LOC129318189 | 1.2 | 1.91E-02 |
| LOC129308721 | 129308721 | uncharacterized LOC129308721 | 1.2 | 2.83E-02 |
| LOC129309286 | 129309286 | co-chaperone protein p23-1-like | 1.2 | 3.87E-02 |
| LOC129292126 | 129292126 | kinesin-like protein KIN-13B | 1.2 | 3.96E-02 |
| LOC129308314 | 129308314 | MA3 DOMAIN-CONTAINING TRANSLATION REGULATC | 1.2 | 1.74E-02 |
| LOC129291057 | 129291057 | DEAD-box ATP-dependent RNA helicase 15-like | 1.2 | 1.98E-02 |
| LOC129313138 | 129313138 | uncharacterized LOC129313138 | 1.2 | 1.08E-02 |
| LOC129315721 | 129315721 | trehalose-phosphate phosphatase A | 1.2 | 1.69E-02 |
| LOC129291278 | 129291278 | uncharacterized protein At2g34160-like | 1.2 | 1.21E-02 |
| LOC129293033 | 129293033 | uncharacterized LOC129293033 | 1.2 | 1.42E-02 |
| LOC129314834 | 129314834 | probable serine/threonine-protein kinase PBL5 | 1.2 | 2.56E-02 |
| LOC129311405 | 129311405 | E3 ubiquitin protein ligase RIE1 | 1.2 | 4.43E-03 |
| LOC129321759 | 129321759 | probable lipid phosphate phosphatase beta | 1.2 | 1.88E-02 |
| LOC129320103 | 129320103 | ACT domain-containing protein ACR12 | 1.2 | 2.90E-02 |

| | | | | |
|---|---|---|---|---|
| LOC129294748 | 129294748 | uncharacterized LOC129294748 | 1.2 | 2.30E-03 |
| LOC129302405 | 129302405 | arogenate dehydrogenase 1, chloroplastic-like | 1.2 | 9.63E-03 |
| LOC129319363 | 129319363 | VIN3-like protein 1 | 1.2 | 1.29E-02 |
| LOC129321984 | 129321984 | pectin acetylesterase 8-like | 1.2 | 2.08E-02 |
| LOC129321375 | 129321375 | uncharacterized LOC129321375 | 1.2 | 6.70E-03 |
| LOC129314567 | 129314567 | uncharacterized LOC129314567 | 1.2 | 3.23E-02 |
| LOC129302395 | 129302395 | BTB/POZ domain-containing protein At2g04740 | 1.2 | 1.06E-02 |
| LOC129291376 | 129291376 | transketolase, chloroplastic-like | 1.2 | 3.04E-02 |
| LOC129307906 | 129307906 | cyclin-dependent kinase inhibitor 4-like | 1.2 | 8.82E-03 |
| LOC129302660 | 129302660 | monodehydroascorbate reductase 4, peroxisomal-like | 1.2 | 2.34E-02 |
| LOC129295362 | 129295362 | auxin response factor 18-like | 1.2 | 4.03E-03 |
| LOC129308041 | 129308041 | acyl carrier protein 1, chloroplastic-like | 1.2 | 8.09E-03 |
| LOC129319779 | 129319779 | uncharacterized LOC129319779 | 1.1 | 2.81E-03 |
| LOC129315810 | 129315810 | L-ascorbate peroxidase, cytosolic | 1.1 | 4.86E-03 |
| LOC129292498 | 129292498 | uncharacterized LOC129292498 | 1.1 | 8.30E-03 |
| LOC129288775 | 129288775 | transcription factor PIF3-like | 1.1 | 1.91E-02 |
| LOC129313520 | 129313520 | protein yippee-like At5g53940 | 1.1 | 4.42E-02 |
| LOC129286845 | 129286845 | uncharacterized LOC129286845 | 1.1 | 3.23E-02 |
| LOC129285158 | 129285158 | 26S proteasome non-ATPase regulatory subunit 8 homolog A | 1.1 | 3.74E-02 |
| LOC129318362 | 129318362 | uncharacterized LOC129318362 | 1.1 | 4.41E-02 |
| LOC129309430 | 129309430 | RNA polymerase II transcriptional coactivator KIWI-like | 1.1 | 3.00E-02 |
| LOC129303147 | 129303147 | probable inactive receptor kinase At1g48480 | 1.1 | 2.60E-02 |
| LOC129293869 | 129293869 | uncharacterized LOC129293869 | 1.1 | 4.07E-02 |
| LOC129305233 | 129305233 | uncharacterized LOC129305233 | 1.1 | 2.70E-03 |
| LOC129298967 | 129298967 | 40S ribosomal protein S21-2 | 1.1 | 3.88E-02 |
| LOC129300289 | 129300289 | FACT complex subunit SSRP1-like | 1.1 | 4.16E-02 |
| LOC129313559 | 129313559 | 60S ribosomal protein L31-like | 1.1 | 4.53E-02 |
| LOC129305866 | 129305866 | vacuolar protein sorting-associated protein 55 homolog | 1.1 | 4.33E-02 |
| LOC129317181 | 129317181 | bifunctional nuclease 1-like | 1.1 | 4.82E-02 |
| LOC129294714 | 129294714 | putative clathrin assembly protein At2g25430 | 1.1 | 1.46E-02 |
| LOC129311249 | 129311249 | F-box/kelch-repeat protein At1g26930-like | 1.1 | 3.93E-02 |
| LOC129322853 | 129322853 | auxin response factor 9 | 1.1 | 3.91E-03 |
| LOC129318360 | 129318360 | mitogen-activated protein kinase 20 | 1.1 | 6.98E-03 |
| LOC129308444 | 129308444 | mavicyanin | 1.1 | 3.17E-02 |
| LOC129302498 | 129302498 | uncharacterized LOC129302498 | 1.1 | 3.16E-02 |
| LOC129306276 | 129306276 | cell division control protein 48 homolog D-like | 1.1 | 1.89E-02 |
| LOC129296982 | 129296982 | probable WRKY transcription factor 4 | 1.1 | 3.70E-02 |
| LOC129310450 | 129310450 | F-box protein SKIP2-like | 1.1 | 8.73E-03 |
| LOC129320491 | 129320491 | uncharacterized LOC129320491 | 1.1 | 3.03E-02 |
| LOC129312875 | 129312875 | uncharacterized LOC129312875 | 1.1 | 1.18E-02 |
| LOC129285120 | 129285120 | protein DEHYDRATION-INDUCED 19-like | 1.1 | 2.99E-02 |
| LOC129285049 | 129285049 | probable receptor-like serine/threonine-protein kinase At4g3450 | 1.1 | 4.26E-02 |
| LOC129311606 | 129311606 | 60S ribosomal protein L27a-3 | 1.1 | 2.82E-02 |
| LOC129290740 | 129290740 | serine/threonine-protein kinase ATG1c | 1.1 | 3.72E-02 |
| LOC129320251 | 129320251 | transcription factor MYB1R1-like | 1.1 | 2.17E-02 |
| LOC129314240 | 129314240 | uncharacterized LOC129314240 | 1.1 | 2.11E-02 |
| LOC129288728 | 129288728 | methyl-CpG-binding domain-containing protein 11-like | 1.1 | 2.38E-02 |
| LOC129289450 | 129289450 | transcription factor MTB1-like | 1.1 | 2.16E-02 |

| | | | | |
|---|---|---|---|---|
| LOC129307774 | 129307774 | probable 60S ribosomal protein L14 | 1.1 | 4.44E-03 |
| LOC129320127 | 129320127 | protein tesmin/TSO1-like CXC 5 | 1.1 | 3.44E-02 |
| LOC129313646 | 129313646 | proton pump-interactor 1-like | 1.1 | 2.25E-02 |
| LOC129301818 | 129301818 | protein DAMAGED DNA-BINDING 2 | 1.1 | 2.48E-02 |
| LOC129308628 | 129308628 | protein KINESIN LIGHT CHAIN-RELATED 3-like | 1.1 | 1.81E-02 |
| LOC129285568 | 129285568 | CSC1-like protein ERD4 | 1.1 | 2.36E-02 |
| LOC129301747 | 129301747 | uncharacterized LOC129301747 | 1.1 | 3.00E-02 |
| LOC129313556 | 129313556 | uncharacterized LOC129313556 | 1.1 | 4.19E-03 |
| LOC129319478 | 129319478 | PWWP domain-containing protein 3-like | 1.1 | 2.99E-02 |
| LOC129317073 | 129317073 | non-specific phospholipase C1 | 1.1 | 1.08E-02 |
| LOC129321029 | 129321029 | lactoylglutathione lyase | 1.1 | 3.06E-02 |
| LOC129287768 | 129287768 | uncharacterized LOC129287768 | 1.1 | 3.21E-02 |
| LOC129307183 | 129307183 | uncharacterized LOC129307183 | 1.1 | 2.76E-02 |
| LOC129309127 | 129309127 | very-long-chain 3-oxoacyl-CoA reductase 1-like | 1.1 | 3.98E-02 |
| LOC129294406 | 129294406 | uncharacterized LOC129294406 | 1.0 | 1.29E-02 |
| LOC129287897 | 129287897 | dnaJ protein ERDJ3A | 1.0 | 2.77E-02 |
| LOC129323167 | 129323167 | KH domain-containing protein HEN4-like | 1.0 | 3.24E-02 |
| LOC129293275 | 129293275 | protein ROH1-like | 1.0 | 2.75E-02 |
| LOC129313799 | 129313799 | protein NUCLEAR FUSION DEFECTIVE 6, mitochondrial-like | 1.0 | 3.39E-02 |
| LOC129321585 | 129321585 | uncharacterized LOC129321585 | 1.0 | 3.67E-02 |
| LOC129298165 | 129298165 | uncharacterized LOC129298165 | 1.0 | 4.32E-02 |
| LOC129313633 | 129313633 | uncharacterized CRM domain-containing protein At3g25440, ch | 1.0 | 4.30E-02 |
| LOC129311601 | 129311601 | probable membrane-associated kinase regulator 1 | 1.0 | 3.52E-02 |
| LOC129320185 | 129320185 | protein SUPPRESSOR OF MAX2 1 | 1.0 | 2.42E-02 |
| LOC129316879 | 129316879 | proteasome subunit alpha type-2-A-like | 1.0 | 4.19E-02 |
| LOC129311374 | 129311374 | O-fucosyltransferase 39 | 1.0 | 3.71E-02 |
| LOC129298862 | 129298862 | beta-1,2-xylosyltransferase-like | 1.0 | 3.83E-02 |
| LOC129307639 | 129307639 | SUMO-activating enzyme subunit 1A-like | 1.0 | 3.07E-02 |
| LOC129306507 | 129306507 | probable small nuclear ribonucleoprotein G | 1.0 | 4.44E-02 |
| LOC129288874 | 129288874 | AT-hook motif nuclear-localized protein 22-like | 1.0 | 3.72E-02 |
| LOC129321476 | 129321476 | transcription initiation factor IIB-2 | 1.0 | 4.65E-02 |
| LOC129304929 | 129304929 | uncharacterized LOC129304929 | 1.0 | 1.33E-02 |
| LOC129308407 | 129308407 | polyadenylate-binding protein 2-like | 1.0 | 3.84E-02 |
| LOC129302215 | 129302215 | protein OXIDATIVE STRESS 3 LIKE 4-like | 1.0 | 1.52E-02 |
| LOC129285941 | 129285941 | chromatin remodeling protein SHL-like | 1.0 | 3.08E-02 |
| LOC129294248 | 129294248 | uncharacterized LOC129294248 | 1.0 | 4.58E-02 |
| LOC129313718 | 129313718 | SWI/SNF complex subunit SWI3A | 1.0 | 1.30E-02 |
| LOC129303532 | 129303532 | protein SMAX1-LIKE 6-like | 1.0 | 1.54E-02 |
| LOC129319182 | 129319182 | zinc finger CCCH domain-containing protein 32-like | -1.0 | 3.37E-02 |
| LOC129291124 | 129291124 | short-chain dehydrogenase TIC 32 B, chloroplastic-like | -1.0 | 3.06E-02 |
| LOC129308626 | 129308626 | dihydroxy-acid dehydratase, chloroplastic | -1.0 | 3.62E-02 |
| LOC129294157 | 129294157 | probable L-ascorbate peroxidase 6, chloroplastic/mitochondrial | -1.0 | 3.14E-02 |
| LOC129319828 | 129319828 | farnesylcysteine lyase | -1.0 | 2.74E-02 |
| LOC129305270 | 129305270 | probable ADP-ribosylation factor GTPase-activating protein AG | -1.0 | 4.23E-02 |
| LOC129304672 | 129304672 | tubulin beta chain | -1.0 | 3.64E-02 |
| LOC129291603 | 129291603 | probable plastid-lipid-associated protein 10, chloroplastic | -1.0 | 2.12E-02 |
| LOC129304838 | 129304838 | two-component response regulator-like APRR7 | -1.0 | 1.89E-02 |
| LOC129288351 | 129288351 | VIN3-like protein 2 | -1.0 | 1.04E-02 |

| | | | | |
|---|---|---|---|---|
| LOC129321373 | 129321373 | probable nucleoredoxin 1 | -1.0 | 2.00E-02 |
| LOC129320301 | 129320301 | S-adenosylmethionine carrier 1, chloroplastic/mitochondrial-like | -1.0 | 3.24E-02 |
| LOC129319895 | 129319895 | alanine--glyoxylate aminotransferase 2 homolog 1, mitochondria | -1.0 | 1.50E-02 |
| LOC129302087 | 129302087 | two-pore potassium channel 1 | -1.0 | 2.78E-02 |
| LOC129285358 | 129285358 | enhancer of mRNA-decapping protein 4-like | -1.0 | 1.84E-02 |
| LOC129319849 | 129319849 | ABC transporter B family member 25-like | -1.0 | 1.38E-02 |
| LOC129300149 | 129300149 | caffeoylshikimate esterase-like | -1.0 | 1.57E-02 |
| LOC129318867 | 129318867 | polyadenylate-binding protein RBP47-like | -1.0 | 2.87E-02 |
| LOC129321405 | 129321405 | pentatricopeptide repeat-containing protein At2g35130 | -1.0 | 2.41E-02 |
| LOC129311261 | 129311261 | adenylate kinase, chloroplastic | -1.0 | 8.29E-03 |
| LOC129322194 | 129322194 | uncharacterized LOC129322194 | -1.0 | 4.63E-02 |
| LOC129309359 | 129309359 | L-type lectin-domain containing receptor kinase S.1 | -1.0 | 6.11E-03 |
| LOC129316411 | 129316411 | topless-related protein 3-like | -1.0 | 1.04E-02 |
| LOC129290181 | 129290181 | putative pentatricopeptide repeat-containing protein At1g02420 | -1.0 | 4.94E-02 |
| LOC129288693 | 129288693 | mechanosensitive ion channel protein 1, mitochondrial | -1.0 | 1.89E-02 |
| LOC129288339 | 129288339 | O-methyltransferase 1, chloroplastic | -1.0 | 3.99E-02 |
| LOC129293671 | 129293671 | probable plastid-lipid-associated protein 13, chloroplastic | -1.0 | 3.32E-02 |
| LOC129306775 | 129306775 | solute carrier family 40 member 3, chloroplastic | -1.0 | 2.97E-02 |
| LOC129320456 | 129320456 | haloacid dehalogenase-like hydrolase domain-containing protein | -1.0 | 3.42E-02 |
| LOC129285103 | 129285103 | alkaline ceramidase | -1.0 | 1.29E-02 |
| LOC129303368 | 129303368 | probable serine/threonine-protein kinase PBL21 | -1.0 | 2.89E-02 |
| LOC129303345 | 129303345 | amino acid transporter AVT6C | -1.0 | 1.68E-02 |
| LOC129304473 | 129304473 | IAA-amino acid hydrolase ILR1-like 9 | -1.0 | 3.40E-02 |
| LOC129302857 | 129302857 | probable pectin methyltransferase QUA2 | -1.0 | 2.54E-02 |
| LOC129303097 | 129303097 | adenylyltransferase and sulfurtransferase MOCS3 | -1.0 | 2.02E-02 |
| LOC129288537 | 129288537 | CBL-interacting serine/threonine-protein kinase 9 | -1.0 | 5.52E-03 |
| LOC129311758 | 129311758 | NAD(P)H-quinone oxidoreductase subunit L, chloroplastic | -1.0 | 4.19E-02 |
| LOC129294807 | 129294807 | L-galactono-1,4-lactone dehydrogenase, mitochondrial | -1.0 | 2.27E-02 |
| LOC129312775 | 129312775 | uncharacterized LOC129312775 | -1.0 | 2.60E-02 |
| LOC129315596 | 129315596 | thiosulfate/3-mercaptopyruvate sulfurtransferase 1, mitochondri | -1.0 | 2.02E-02 |
| LOC129291707 | 129291707 | protein MANNAN SYNTHESIS-RELATED 1-like | -1.0 | 4.77E-02 |
| LOC129316039 | 129316039 | protein WHAT'S THIS FACTOR 1 homolog, chloroplastic-like | -1.0 | 3.88E-02 |
| LOC129317846 | 129317846 | receptor homology region, transmembrane domain- and RING d | -1.0 | 3.39E-02 |
| LOC129292077 | 129292077 | nucleobase-ascorbate transporter 4 | -1.0 | 2.89E-02 |
| LOC129311179 | 129311179 | uncharacterized LOC129311179 | -1.0 | 1.69E-02 |
| LOC129305753 | 129305753 | protein IQ-DOMAIN 2-like | -1.1 | 1.23E-02 |
| LOC129284822 | 129284822 | tyrosyl-DNA phosphodiesterase 1 | -1.1 | 4.27E-02 |
| LOC129295038 | 129295038 | WAT1-related protein At5g40240-like | -1.1 | 2.55E-02 |
| LOC129322177 | 129322177 | uncharacterized LOC129322177 | -1.1 | 3.65E-02 |
| LOC129306256 | 129306256 | receptor-like protein 4 | -1.1 | 6.58E-03 |
| LOC129312503 | 129312503 | transcriptional activator DEMETER | -1.1 | 3.60E-02 |
| LOC129310454 | 129310454 | inorganic pyrophosphatase TTM2-like | -1.1 | 4.75E-02 |
| LOC129285840 | 129285840 | peroxisomal ATPase PEX1 | -1.1 | 2.33E-02 |
| LOC129319385 | 129319385 | IQ domain-containing protein IQM1-like | -1.1 | 9.57E-03 |
| LOC129292672 | 129292672 | BAHD acyltransferase DCR | -1.1 | 3.08E-02 |
| LOC129288708 | 129288708 | abscisic acid receptor PYL4 | -1.1 | 4.53E-02 |
| LOC129309462 | 129309462 | mediator of RNA polymerase II transcription subunit 13 | -1.1 | 2.24E-02 |
| LOC129293703 | 129293703 | CRM-domain containing factor CFM2, chloroplastic | -1.1 | 1.84E-02 |

| | | | | |
|---|---|---|---|---|
| LOC129290813 | 129290813 | U-box domain-containing protein 44-like | -1.1 | 3.14E-02 |
| LOC129314831 | 129314831 | uncharacterized LOC129314831 | -1.1 | 3.72E-02 |
| LOC129314989 | 129314989 | solanesyl diphosphate synthase 1, chloroplastic | -1.1 | 3.00E-02 |
| LOC129308610 | 129308610 | DEAD-box ATP-dependent RNA helicase 58, chloroplastic | -1.1 | 1.98E-02 |
| LOC129311545 | 129311545 | elongation factor G-2, chloroplastic | -1.1 | 4.67E-02 |
| LOC129322218 | 129322218 | glucan endo-1,3-beta-glucosidase 3-like | -1.1 | 3.22E-02 |
| LOC129284603 | 129284603 | trihelix transcription factor DF1-like | -1.1 | 8.65E-03 |
| LOC129288521 | 129288521 | protein PSK SIMULATOR 1-like | -1.1 | 2.82E-02 |
| LOC129318698 | 129318698 | neutral ceramidase 1-like | -1.1 | 1.92E-03 |
| LOC129309232 | 129309232 | choline monooxygenase, chloroplastic | -1.1 | 3.68E-02 |
| LOC129303365 | 129303365 | uncharacterized LOC129303365 | -1.1 | 5.98E-03 |
| LOC129293377 | 129293377 | F-box/kelch-repeat protein At1g67480 | -1.1 | 2.47E-03 |
| LOC129320151 | 129320151 | uncharacterized LOC129320151 | -1.1 | 9.00E-03 |
| LOC129322747 | 129322747 | uncharacterized LOC129322747 | -1.1 | 2.28E-02 |
| LOC129294451 | 129294451 | phosphoinositide phospholipase C 2-like | -1.1 | 3.81E-03 |
| LOC129302875 | 129302875 | uncharacterized LOC129302875 | -1.1 | 5.83E-03 |
| LOC129323007 | 129323007 | inorganic phosphate transporter 2-1, chloroplastic-like | -1.1 | 2.33E-02 |
| LOC129305973 | 129305973 | nudix hydrolase 2-like | -1.1 | 1.23E-02 |
| LOC129284651 | 129284651 | GDP-mannose 3,5-epimerase 2 | -1.1 | 3.12E-02 |
| LOC129317999 | 129317999 | uncharacterized LOC129317999 | -1.1 | 1.91E-02 |
| LOC129321225 | 129321225 | telomere repeat-binding factor 2 | -1.1 | 9.13E-03 |
| LOC129311811 | 129311811 | transcription factor BHLH089-like | -1.1 | 1.15E-02 |
| LOC129286946 | 129286946 | ras-related protein RABA2a | -1.1 | 3.56E-02 |
| LOC129310864 | 129310864 | protein PHOSPHATE STARVATION RESPONSE 1-like | -1.1 | 4.68E-03 |
| LOC129287701 | 129287701 | transcription factor bHLH62-like | -1.1 | 1.62E-02 |
| LOC129289488 | 129289488 | fructose-1,6-bisphosphatase, chloroplastic | -1.1 | 3.90E-02 |
| LOC129321906 | 129321906 | protein transport protein SEC23 A | -1.1 | 3.99E-03 |
| LOC129306892 | 129306892 | uncharacterized LOC129306892 | -1.1 | 4.94E-02 |
| LOC129315449 | 129315449 | BTB/POZ domain-containing protein At3g44820 | -1.1 | 1.95E-02 |
| LOC129292195 | 129292195 | mediator of RNA polymerase II transcription subunit 33A-like | -1.1 | 1.42E-02 |
| LOC129320213 | 129320213 | 1-phosphatidylinositol-3-phosphate 5-kinase FAB1B-like | -1.1 | 8.82E-03 |
| LOC129308614 | 129308614 | peroxisomal adenine nucleotide carrier 1-like | -1.1 | 9.69E-03 |
| LOC129308588 | 129308588 | 5-oxoprolinase 1-like | -1.1 | 1.35E-02 |
| LOC129293023 | 129293023 | glycerate dehydrogenase | -1.1 | 2.68E-02 |
| LOC129304818 | 129304818 | phosphoribosylaminoimidazole carboxylase, chloroplastic-like | -1.1 | 1.89E-02 |
| LOC129294254 | 129294254 | putative pentatricopeptide repeat-containing protein At1g56570 | -1.1 | 1.20E-02 |
| LOC129284633 | 129284633 | probable protein phosphatase 2C 15 | -1.1 | 2.11E-02 |
| LOC129315755 | 129315755 | leucine-rich repeat receptor-like kinase protein HAR1 | -1.1 | 2.73E-02 |
| LOC129303744 | 129303744 | WD repeat-containing protein PCN-like | -1.1 | 4.08E-02 |
| LOC129318900 | 129318900 | chloroplastic group IIA intron splicing facilitator CRS1, chlorop | -1.1 | 1.83E-02 |
| LOC129303284 | 129303284 | UDP-glycosyltransferase 71K2-like | -1.1 | 2.41E-02 |
| LOC129289768 | 129289768 | DNA-repair protein XRCC1 | -1.1 | 2.76E-02 |
| LOC129304440 | 129304440 | peptidyl-prolyl cis-trans isomerase FKBP17-1, chloroplastic | -1.1 | 4.94E-02 |
| LOC129318206 | 129318206 | pentatricopeptide repeat-containing protein At4g38150 | -1.1 | 1.38E-02 |
| LOC129314866 | 129314866 | ankyrin repeat-containing protein At5g02620-like | -1.1 | 3.71E-03 |
| LOC129309925 | 129309925 | pentatricopeptide repeat-containing protein At1g04840 | -1.1 | 3.90E-02 |
| LOC129298404 | 129298404 | 3beta-hydroxysteroid-dehydrogenase/decarboxylase-like | -1.1 | 2.55E-02 |
| LOC129288955 | 129288955 | calcium-transporting ATPase 9, plasma membrane-type-like | -1.1 | 4.38E-03 |

| | | | | |
|---|---|---|---|---|
| LOC129309954 | 129309954 | protein TIME FOR COFFEE-like | -1.1 | 9.30E-03 |
| LOC129320000 | 129320000 | plasma membrane ATPase 4-like | -1.1 | 3.96E-02 |
| LOC129305847 | 129305847 | uncharacterized LOC129305847 | -1.1 | 1.91E-03 |
| LOC129284823 | 129284823 | protein TRIGALACTOSYLDIACYLGLYCEROL 4, chloroplas | -1.1 | 2.94E-02 |
| LOC129304381 | 129304381 | zinc finger protein GAI-ASSOCIATED FACTOR 1-like | -1.1 | 4.93E-02 |
| LOC129296694 | 129296694 | protein REGULATOR OF FATTY ACID COMPOSITION 3, c | -1.1 | 4.55E-02 |
| LOC129311153 | 129311153 | BEL1-like homeodomain protein 4 | -1.1 | 8.79E-03 |
| LOC129322279 | 129322279 | probable cadmium/zinc-transporting ATPase HMA1, chloroplas | -1.1 | 1.25E-02 |
| LOC129322183 | 129322183 | uncharacterized LOC129322183 | -1.1 | 1.23E-03 |
| LOC129306549 | 129306549 | protein IQ-DOMAIN 9-like | -1.1 | 1.89E-02 |
| LOC129306938 | 129306938 | cyclin-dependent kinase F-1 | -1.1 | 1.43E-02 |
| LOC129291161 | 129291161 | uncharacterized LOC129291161 | -1.1 | 3.82E-02 |
| LOC129289581 | 129289581 | probable methyltransferase PMT3 | -1.2 | 7.71E-04 |
| LOC129311419 | 129311419 | zinc finger protein CONSTANS-LIKE 16-like | -1.2 | 1.66E-02 |
| LOC129302040 | 129302040 | uncharacterized LOC129302040 | -1.2 | 3.10E-02 |
| LOC129286819 | 129286819 | sugar carrier protein C-like | -1.2 | 2.10E-03 |
| LOC129285829 | 129285829 | histidinol dehydrogenase, chloroplastic-like | -1.2 | 3.06E-02 |
| LOC129287161 | 129287161 | wall-associated receptor kinase-like 8 | -1.2 | 3.96E-02 |
| LOC129320210 | 129320210 | serine/threonine-protein kinase PBL34-like | -1.2 | 1.25E-02 |
| LOC129313163 | 129313163 | uncharacterized LOC129313163 | -1.2 | 2.49E-02 |
| LOC129305519 | 129305519 | cell morphogenesis protein PAG1 | -1.2 | 4.79E-02 |
| LOC129320877 | 129320877 | protein transport protein SEC23 G | -1.2 | 4.82E-02 |
| LOC129320328 | 129320328 | adagio protein 1-like | -1.2 | 6.96E-03 |
| LOC129315537 | 129315537 | cellulose synthase A catalytic subunit 3 [UDP-forming]-like | -1.2 | 2.09E-02 |
| LOC129309499 | 129309499 | uridylate kinase PUMPKIN, chloroplastic | -1.2 | 4.91E-02 |
| LOC129287956 | 129287956 | fe-S cluster assembly factor HCF101, chloroplastic | -1.2 | 7.59E-03 |
| LOC129304891 | 129304891 | uncharacterized LOC129304891 | -1.2 | 3.07E-02 |
| LOC129319341 | 129319341 | serine/threonine protein phosphatase 2A 57 kDa regulatory subu | -1.2 | 2.54E-02 |
| LOC129309788 | 129309788 | tubulin alpha-3 chain | -1.2 | 3.25E-03 |
| LOC129311654 | 129311654 | uncharacterized LOC129311654 | -1.2 | 9.80E-04 |
| LOC129311848 | 129311848 | branched-chain amino acid aminotransferase 2, chloroplastic-lik | -1.2 | 1.32E-02 |
| LOC129320688 | 129320688 | E3 ubiquitin-protein ligase XBAT32 | -1.2 | 1.14E-02 |
| LOC129318299 | 129318299 | serine/threonine-protein kinase EDR1-like | -1.2 | 4.11E-02 |
| LOC129291403 | 129291403 | ethylene-responsive transcription factor-like protein At4g13040 | -1.2 | 4.33E-02 |
| LOC129321685 | 129321685 | uncharacterized LOC129321685 | -1.2 | 1.08E-02 |
| LOC129303963 | 129303963 | uncharacterized LOC129303963 | -1.2 | 6.51E-03 |
| LOC129311857 | 129311857 | probable enoyl-CoA hydratase 2, mitochondrial | -1.2 | 3.97E-02 |
| LOC129290729 | 129290729 | mediator of RNA polymerase II transcription subunit 15a-like | -1.2 | 2.75E-03 |
| LOC129290376 | 129290376 | cleavage stimulating factor 64-like | -1.2 | 4.57E-02 |
| LOC129303103 | 129303103 | coronatine-insensitive protein 1-like | -1.2 | 2.11E-02 |
| LOC129293634 | 129293634 | photosynthetic NDH subunit of lumenal location 1, chloroplastic | -1.2 | 1.64E-02 |
| LOC129304795 | 129304795 | uncharacterized LOC129304795 | -1.2 | 4.74E-02 |
| LOC129291095 | 129291095 | thioredoxin-like 3-1, chloroplastic | -1.2 | 2.85E-02 |
| LOC129302274 | 129302274 | abscisic-aldehyde oxidase-like | -1.2 | 3.35E-03 |
| LOC129297825 | 129297825 | probable pectin methyltransferase QUA3 | -1.2 | 4.12E-02 |
| LOC129315674 | 129315674 | probable NOT transcription complex subunit VIP2 | -1.2 | 7.71E-03 |
| LOC129302268 | 129302268 | photosystem I chlorophyll a/b-binding protein 5, chloroplastic | -1.2 | 2.96E-02 |
| LOC129316126 | 129316126 | adenosine kinase 2-like | -1.2 | 2.63E-02 |

| | | | | |
|---|---|---|---|---|
| LOC129285706 | 129285706 | KH domain-containing protein At4g18375 | -1.2 | 3.40E-03 |
| LOC129294372 | 129294372 | folylpolyglutamate synthase | -1.2 | 4.58E-02 |
| LOC129309201 | 129309201 | fibrillin protein 5 homolog | -1.2 | 4.17E-02 |
| LOC129322817 | 129322817 | methionine aminopeptidase 1D, chloroplastic/mitochondrial | -1.2 | 4.71E-02 |
| LOC129311272 | 129311272 | arogenate dehydrogenase 1, chloroplastic-like | -1.2 | 4.82E-03 |
| LOC129288277 | 129288277 | AMP deaminase-like | -1.2 | 1.12E-02 |
| LOC129303771 | 129303771 | glucose-6-phosphate isomerase 1, chloroplastic | -1.2 | 2.19E-03 |
| LOC129317787 | 129317787 | cytochrome c oxidase assembly protein COX15 | -1.2 | 1.65E-02 |
| LOC129293018 | 129293018 | putative lipid phosphate phosphatase 3, chloroplastic | -1.2 | 3.81E-02 |
| LOC129284961 | 129284961 | pentatricopeptide repeat-containing protein At2g30100, chloropl | -1.2 | 6.03E-03 |
| LOC129289839 | 129289839 | uncharacterized LOC129289839 | -1.2 | 1.93E-02 |
| LOC129311106 | 129311106 | protein TWIN LOV 1 | -1.2 | 1.19E-02 |
| LOC129320903 | 129320903 | 1-deoxy-D-xylulose 5-phosphate reductoisomerase, chloroplasti | -1.2 | 7.70E-03 |
| LOC129292782 | 129292782 | glutamine synthetase leaf isozyme, chloroplastic | -1.2 | 1.14E-02 |
| LOC129313010 | 129313010 | wax ester synthase/diacylglycerol acyltransferase 4-like | -1.2 | 6.24E-03 |
| LOC129309339 | 129309339 | calcium sensing receptor, chloroplastic | -1.2 | 3.41E-02 |
| LOC129302164 | 129302164 | putative DUF21 domain-containing protein At3g13070, chlorop | -1.2 | 2.92E-03 |
| LOC129322213 | 129322213 | uncharacterized LOC129322213 | -1.2 | 4.57E-03 |
| LOC129315752 | 129315752 | dicarboxylate transporter 1, chloroplastic-like | -1.2 | 4.03E-02 |
| LOC129293423 | 129293423 | sucrose-phosphatase 2-like | -1.2 | 1.10E-02 |
| LOC129322649 | 129322649 | mediator of RNA polymerase II transcription subunit 25-like | -1.2 | 4.91E-02 |
| LOC129322131 | 129322131 | uncharacterized LOC129322131 | -1.2 | 2.99E-02 |
| LOC129292540 | 129292540 | cytochrome P450 89A2-like | -1.2 | 1.77E-02 |
| LOC129306800 | 129306800 | protein RESISTANCE TO PHYTOPHTHORA 1, chloroplastic | -1.2 | 1.65E-02 |
| LOC129292539 | 129292539 | uncharacterized LOC129292539 | -1.2 | 1.09E-02 |
| LOC129322146 | 129322146 | sec-independent protein translocase protein TATC, chloroplastic | -1.2 | 1.04E-02 |
| LOC129322681 | 129322681 | aluminum-activated malate transporter 12-like | -1.2 | 3.06E-02 |
| LOC129318116 | 129318116 | probable amino acid permease 7 | -1.2 | 2.84E-02 |
| LOC129315143 | 129315143 | protein N-terminal asparagine amidohydrolase | -1.2 | 4.85E-02 |
| LOC129317759 | 129317759 | uncharacterized protein ycf20 | -1.2 | 4.18E-02 |
| LOC129313671 | 129313671 | coumaroyl-CoA:anthocyanidin 3-O-glucoside-6"-O-coumaroyltr | -1.2 | 2.48E-02 |
| LOC129285762 | 129285762 | allantoate deiminase 2 | -1.2 | 4.70E-02 |
| LOC129297365 | 129297365 | rhodanese-like domain-containing protein 4, chloroplastic | -1.2 | 2.51E-02 |
| LOC129320824 | 129320824 | peroxisomal membrane protein PEX14-like | -1.2 | 1.65E-02 |
| LOC129291387 | 129291387 | peptide chain release factor PrfB3, chloroplastic | -1.2 | 1.87E-04 |
| LOC129288448 | 129288448 | polygalacturonate 4-alpha-galacturonosyltransferase-like | -1.2 | 3.89E-02 |
| LOC129312788 | 129312788 | uncharacterized LOC129312788 | -1.2 | 1.41E-02 |
| LOC129287054 | 129287054 | elongation factor 1-gamma-like | -1.2 | 4.96E-03 |
| LOC129313484 | 129313484 | patellin-3-like | -1.2 | 1.91E-02 |
| LOC129294621 | 129294621 | uncharacterized LOC129294621 | -1.2 | 3.07E-02 |
| LOC129315909 | 129315909 | uncharacterized LOC129315909 | -1.2 | 1.79E-02 |
| LOC129313134 | 129313134 | uncharacterized LOC129313134 | -1.2 | 2.34E-02 |
| LOC129320096 | 129320096 | probable protein phosphatase 2C 27 | -1.2 | 3.39E-02 |
| LOC129321606 | 129321606 | bifunctional dTDP-4-dehydrorhamnose 3,5-epimerase/dTDP-4- | -1.2 | 4.34E-02 |
| LOC129314376 | 129314376 | uncharacterized LOC129314376 | -1.2 | 1.42E-02 |
| LOC129302404 | 129302404 | protein RETICULATA-RELATED 5, chloroplastic-like | -1.2 | 1.64E-04 |
| LOC129294439 | 129294439 | psbP domain-containing protein 5, chloroplastic | -1.2 | 2.13E-02 |
| LOC129293353 | 129293353 | aminomethyltransferase, mitochondrial | -1.2 | 2.16E-02 |

| | | | | |
|---|---|---|---|---|
| LOC129317986 | 129317986 | glutenin, high molecular weight subunit DX5 | -1.2 | 2.42E-02 |
| LOC129285566 | 129285566 | WRKY transcription factor SUSIBA2-like | -1.2 | 4.40E-02 |
| LOC129302913 | 129302913 | delta-1-pyrroline-5-carboxylate synthase | -1.2 | 3.09E-02 |
| LOC129319504 | 129319504 | uncharacterized LOC129319504 | -1.2 | 4.82E-02 |
| LOC129306044 | 129306044 | nonsense-mediated mRNA decay factor SMG7-like | -1.3 | 3.15E-03 |
| LOC129302963 | 129302963 | probable protein phosphatase 2C 55 | -1.3 | 1.88E-03 |
| LOC129284850 | 129284850 | probable helicase MAGATAMA 3 | -1.3 | 4.13E-03 |
| LOC129292887 | 129292887 | cytochrome P450 89A2-like | -1.3 | 2.48E-04 |
| LOC129293821 | 129293821 | chloride channel protein CLC-e | -1.3 | 9.04E-03 |
| LOC129293041 | 129293041 | serine carboxypeptidase-like 27 | -1.3 | 4.40E-02 |
| LOC129304980 | 129304980 | ACT domain-containing protein ACR10-like | -1.3 | 2.50E-02 |
| LOC129309348 | 129309348 | histone deacetylase 14, chloroplastic | -1.3 | 7.58E-03 |
| LOC129320090 | 129320090 | uncharacterized LOC129320090 | -1.3 | 1.45E-02 |
| LOC129292291 | 129292291 | probable flavin-containing monooxygenase 1 | -1.3 | 3.25E-03 |
| LOC129305192 | 129305192 | putative GTP diphosphokinase RSH1, chloroplastic | -1.3 | 3.59E-03 |
| LOC129294247 | 129294247 | uncharacterized LOC129294247 | -1.3 | 3.80E-03 |
| LOC129316056 | 129316056 | flowering time control protein FY | -1.3 | 5.53E-03 |
| LOC129304786 | 129304786 | potassium transporter 11-like | -1.3 | 1.75E-02 |
| LOC129291055 | 129291055 | syntaxin-71-like | -1.3 | 1.01E-02 |
| LOC129321713 | 129321713 | putative potassium transporter 12 | -1.3 | 8.89E-03 |
| LOC129292856 | 129292856 | NADPH-dependent aldehyde reductase 1, chloroplastic-like | -1.3 | 3.75E-02 |
| LOC129310100 | 129310100 | chlorophyll a-b binding protein 4, chloroplastic | -1.3 | 1.77E-02 |
| LOC129285602 | 129285602 | translocase of chloroplast 159, chloroplastic | -1.3 | 4.35E-03 |
| LOC129306970 | 129306970 | stromal 70 kDa heat shock-related protein, chloroplastic-like | -1.3 | 1.91E-02 |
| LOC129298690 | 129298690 | 3-ketoacyl-CoA synthase 6 | -1.3 | 4.15E-02 |
| LOC129295630 | 129295630 | uncharacterized LOC129295630 | -1.3 | 3.21E-02 |
| LOC129288072 | 129288072 | peptidyl-prolyl cis-trans isomerase FKBP17-2, chloroplastic | -1.3 | 4.40E-02 |
| LOC129300581 | 129300581 | F-box protein GID2-like | -1.3 | 2.06E-02 |
| LOC129301800 | 129301800 | uncharacterized LOC129301800 | -1.3 | 1.10E-02 |
| LOC129307372 | 129307372 | pentatricopeptide repeat-containing protein At5g66520-like | -1.3 | 3.23E-02 |
| LOC129315678 | 129315678 | protein SPA1-RELATED 3 | -1.3 | 3.41E-02 |
| LOC129320135 | 129320135 | lipase-like PAD4 | -1.3 | 2.49E-02 |
| LOC129322236 | 129322236 | very-long-chain (3R)-3-hydroxyacyl-CoA dehydratase PASTICC | -1.3 | 4.39E-03 |
| LOC129318220 | 129318220 | chlorophyll a-b binding protein 7, chloroplastic | -1.3 | 1.29E-02 |
| LOC129306502 | 129306502 | uncharacterized LOC129306502 | -1.3 | 2.60E-02 |
| LOC129285379 | 129285379 | glyoxysomal processing protease, glyoxysomal | -1.3 | 3.06E-03 |
| LOC129318149 | 129318149 | protein RBL | -1.3 | 2.48E-02 |
| LOC129317964 | 129317964 | pleiotropic drug resistance protein 3-like | -1.3 | 1.29E-02 |
| LOC129288510 | 129288510 | AT-hook motif nuclear-localized protein 9-like | -1.3 | 2.65E-02 |
| LOC129298276 | 129298276 | NAC domain-containing protein 17-like | -1.3 | 1.09E-02 |
| LOC129309786 | 129309786 | uncharacterized LOC129309786 | -1.3 | 3.27E-02 |
| LOC129303045 | 129303045 | endo-1,4-beta-xylanase 1-like | -1.3 | 1.37E-02 |
| LOC129311060 | 129311060 | heat shock factor protein HSF8-like | -1.3 | 1.49E-02 |
| LOC129285927 | 129285927 | uncharacterized LOC129285927 | -1.3 | 1.30E-02 |
| LOC129296698 | 129296698 | uncharacterized LOC129296698 | -1.3 | 3.40E-02 |
| LOC129320774 | 129320774 | uncharacterized LOC129320774 | -1.3 | 3.61E-02 |
| LOC129321992 | 129321992 | zinc finger CCCH domain-containing protein 62-like | -1.3 | 6.39E-03 |
| LOC129285886 | 129285886 | potassium transporter 10-like | -1.3 | 1.19E-02 |

| | | | | |
|---|---|---|---|---|
| LOC129319183 | 129319183 | probable clathrin assembly protein At4g32285 | -1.3 | 6.73E-03 |
| LOC129293011 | 129293011 | trihelix transcription factor DF1 | -1.3 | 1.07E-02 |
| LOC129290555 | 129290555 | uncharacterized LOC129290555 | -1.3 | 3.75E-02 |
| LOC129310965 | 129310965 | mitochondrial outer membrane protein porin 2-like | -1.3 | 4.70E-02 |
| LOC129292078 | 129292078 | ACT domain-containing protein ACR9 | -1.3 | 4.27E-02 |
| LOC129304787 | 129304787 | RNA polymerase sigma factor sigF, chloroplastic | -1.3 | 2.18E-02 |
| LOC129315078 | 129315078 | serine carboxypeptidase-like 27 | -1.3 | 2.89E-02 |
| LOC129311185 | 129311185 | pentatricopeptide repeat-containing protein At5g02830, chloropl | -1.3 | 1.28E-02 |
| LOC129305382 | 129305382 | protein PSK SIMULATOR 3 | -1.3 | 2.89E-02 |
| LOC129292521 | 129292521 | probable polyamine transporter At3g19553 | -1.3 | 1.06E-02 |
| LOC129311928 | 129311928 | uncharacterized LOC129311928 | -1.3 | 4.19E-02 |
| LOC129303775 | 129303775 | GDSL esterase/lipase At4g10955-like | -1.3 | 6.51E-03 |
| LOC129322847 | 129322847 | probable carboxylesterase 2 | -1.3 | 4.13E-02 |
| LOC129285799 | 129285799 | transcription factor TCP2-like | -1.3 | 4.64E-03 |
| LOC129305967 | 129305967 | protein SAR DEFICIENT 4 | -1.3 | 9.67E-03 |
| LOC129308056 | 129308056 | probable methyltransferase PMT19 | -1.3 | 1.62E-03 |
| LOC129316444 | 129316444 | receptor-like protein kinase FERONIA | -1.3 | 1.51E-02 |
| LOC129304888 | 129304888 | pentatricopeptide repeat-containing protein At3g57430, chloropl | -1.3 | 3.72E-02 |
| LOC129310493 | 129310493 | uncharacterized LOC129310493 | -1.3 | 3.18E-02 |
| LOC129288208 | 129288208 | uncharacterized LOC129288208 | -1.3 | 1.01E-02 |
| LOC129320908 | 129320908 | heparanase-like protein 1 | -1.3 | 1.41E-02 |
| LOC129294314 | 129294314 | uncharacterized LOC129294314 | -1.3 | 2.68E-04 |
| LOC129294686 | 129294686 | uncharacterized LOC129294686 | -1.3 | 3.13E-03 |
| LOC129304625 | 129304625 | uncharacterized LOC129304625 | -1.3 | 4.14E-03 |
| LOC129310377 | 129310377 | BTB/POZ domain-containing protein SR1IP1 | -1.3 | 1.63E-03 |
| LOC129310921 | 129310921 | serine/threonine-protein kinase STY17-like | -1.3 | 4.20E-02 |
| LOC129291371 | 129291371 | (R)-mandelonitrile lyase-like | -1.3 | 4.92E-02 |
| LOC129312711 | 129312711 | E3 ubiquitin-protein ligase SDIR1-like | -1.3 | 1.51E-02 |
| LOC129308621 | 129308621 | phytochrome B-2-like | -1.3 | 2.17E-02 |
| LOC129293363 | 129293363 | protein DETOXIFICATION 46, chloroplastic | -1.3 | 2.88E-03 |
| LOC129319198 | 129319198 | probable inactive purple acid phosphatase 27 | -1.3 | 1.70E-02 |
| LOC129321396 | 129321396 | embryogenesis-associated protein EMB8-like | -1.3 | 2.68E-03 |
| LOC129318104 | 129318104 | uncharacterized LOC129318104 | -1.3 | 2.87E-02 |
| LOC129319950 | 129319950 | cellulose synthase A catalytic subunit 6 [UDP-forming]-like | -1.3 | 2.99E-02 |
| LOC129322201 | 129322201 | probable alpha,alpha-trehalose-phosphate synthase [UDP-formir | -1.3 | 2.84E-03 |
| LOC129291135 | 129291135 | 2-succinylbenzoate--CoA ligase, chloroplastic/peroxisomal | -1.3 | 6.61E-03 |
| LOC129318547 | 129318547 | transcription termination factor MTERF6, chloroplastic/mitocho | -1.3 | 3.07E-02 |
| LOC129304069 | 129304069 | U-box domain-containing protein 15 | -1.3 | 2.61E-02 |
| LOC129315285 | 129315285 | phototropin-1 | -1.4 | 2.21E-02 |
| LOC129290962 | 129290962 | transcription termination factor MTERF5, chloroplastic | -1.4 | 1.29E-02 |
| LOC129305874 | 129305874 | probable metal-nicotianamine transporter YSL6 | -1.4 | 2.02E-02 |
| LOC129306278 | 129306278 | phosphoglycerate kinase, cytosolic-like | -1.4 | 1.96E-02 |
| LOC129292475 | 129292475 | probable protein arginine N-methyltransferase 1 | -1.4 | 1.89E-02 |
| LOC129314272 | 129314272 | arogenate dehydratase/prephenate dehydratase 1, chloroplastic-li | -1.4 | 3.08E-03 |
| LOC129285236 | 129285236 | probable protein phosphatase 2C 62 | -1.4 | 1.62E-02 |
| LOC129320242 | 129320242 | pentatricopeptide repeat-containing protein At1g74850, chloropl | -1.4 | 2.06E-02 |
| LOC129288768 | 129288768 | uncharacterized LOC129288768 | -1.4 | 4.71E-02 |
| LOC129319857 | 129319857 | uncharacterized LOC129319857 | -1.4 | 3.61E-02 |

| | | | | |
|---|---|---|---|---|
| LOC129309206 | 129309206 | divinyl chlorophyllide a 8-vinyl-reductase, chloroplastic | -1.4 | 2.37E-02 |
| LOC129309373 | 129309373 | calnexin homolog | -1.4 | 4.61E-02 |
| LOC129309153 | 129309153 | receptor-like kinase TMK4 | -1.4 | 4.73E-02 |
| LOC129321088 | 129321088 | photosynthetic NDH subunit of lumenal location 4, chloroplastic | -1.4 | 9.08E-03 |
| LOC129322666 | 129322666 | uncharacterized LOC129322666 | -1.4 | 3.44E-02 |
| LOC129301984 | 129301984 | UDP-galactose/UDP-glucose transporter 7-like | -1.4 | 2.14E-02 |
| LOC129305257 | 129305257 | organelle RRM domain-containing protein 1, chloroplastic | -1.4 | 2.25E-02 |
| LOC129305749 | 129305749 | autophagy-related protein 9-like | -1.4 | 8.29E-03 |
| LOC129306133 | 129306133 | protein CHUP1, chloroplastic-like | -1.4 | 1.29E-02 |
| LOC129295766 | 129295766 | bifunctional riboflavin kinase/FMN phosphatase-like | -1.4 | 1.46E-02 |
| LOC129295286 | 129295286 | transcription factor TCP4-like | -1.4 | 5.08E-03 |
| LOC129313660 | 129313660 | protein SPA1-RELATED 3-like | -1.4 | 4.91E-02 |
| LOC129312816 | 129312816 | uncharacterized LOC129312816 | -1.4 | 1.20E-02 |
| LOC129284897 | 129284897 | uncharacterized LOC129284897 | -1.4 | 1.83E-02 |
| LOC129300720 | 129300720 | glutamate-1-semialdehyde 2,1-aminomutase, chloroplastic | -1.4 | 1.54E-02 |
| LOC129305837 | 129305837 | potassium channel AKT2/3-like | -1.4 | 4.22E-03 |
| LOC129313343 | 129313343 | probable leucine-rich repeat receptor-like serine/threonine-protei | -1.4 | 1.21E-02 |
| LOC129288945 | 129288945 | uncharacterized LOC129288945 | -1.4 | 5.22E-03 |
| LOC129302501 | 129302501 | pleiotropic drug resistance protein 1-like | -1.4 | 6.49E-03 |
| LOC129321714 | 129321714 | dihydrolipoyl dehydrogenase, mitochondrial-like | -1.4 | 1.01E-03 |
| LOC129321401 | 129321401 | isopentenyl phosphate kinase | -1.4 | 4.74E-02 |
| LOC129309027 | 129309027 | rubisco accumulation factor 1.1, chloroplastic-like | -1.4 | 2.81E-03 |
| LOC129285932 | 129285932 | nodulin-26-like | -1.4 | 2.14E-02 |
| LOC129307636 | 129307636 | filament-like plant protein | -1.4 | 9.61E-03 |
| LOC129314434 | 129314434 | TPR repeat-containing thioredoxin TTL1-like | -1.4 | 3.45E-02 |
| LOC129286080 | 129286080 | uncharacterized LOC129286080 | -1.4 | 1.07E-02 |
| LOC129308030 | 129308030 | uncharacterized LOC129308030 | -1.4 | 3.10E-02 |
| LOC129293221 | 129293221 | sorbitol dehydrogenase | -1.4 | 1.22E-03 |
| LOC129292958 | 129292958 | transcription factor bHLH7-like | -1.4 | 9.81E-03 |
| LOC129303328 | 129303328 | RING-H2 finger protein ATL66-like | -1.4 | 3.82E-02 |
| LOC129312769 | 129312769 | dihydroorotase, mitochondrial-like | -1.4 | 8.33E-03 |
| LOC129303219 | 129303219 | xylulose kinase 2-like | -1.4 | 3.89E-03 |
| LOC129299639 | 129299639 | cysteine proteinase mucunain-like | -1.4 | 2.20E-03 |
| LOC129305297 | 129305297 | transcription factor EMB1444-like | -1.4 | 9.87E-04 |
| LOC129292930 | 129292930 | kinesin-like protein KIN-7E | -1.4 | 1.69E-05 |
| LOC129321082 | 129321082 | uncharacterized LOC129321082 | -1.4 | 6.70E-03 |
| LOC129305825 | 129305825 | ATP-dependent DNA helicase Q-like SIM | -1.4 | 2.30E-03 |
| LOC129285523 | 129285523 | uncharacterized LOC129285523 | -1.4 | 4.18E-02 |
| LOC129322953 | 129322953 | uncharacterized LOC129322953 | -1.4 | 3.08E-02 |
| LOC129313571 | 129313571 | WEB family protein At3g02930, chloroplastic-like | -1.4 | 1.36E-02 |
| LOC129322965 | 129322965 | protein indeterminate-domain 4, chloroplastic-like | -1.4 | 2.08E-02 |
| LOC129311150 | 129311150 | protein CHUP1, chloroplastic-like | -1.4 | 9.08E-06 |
| LOC129306801 | 129306801 | cyclic dof factor 2 | -1.4 | 1.52E-03 |
| LOC129298366 | 129298366 | glyoxylate/hydroxypyruvate/pyruvate reductase 2KGR-like | -1.4 | 1.76E-04 |
| LOC129322811 | 129322811 | SPX domain-containing membrane protein At4g22990-like | -1.4 | 4.50E-03 |
| LOC129305209 | 129305209 | probable ribose-5-phosphate isomerase 3, chloroplastic | -1.4 | 4.82E-02 |
| LOC129293751 | 129293751 | ATP synthase gamma chain, chloroplastic | -1.4 | 1.49E-02 |
| LOC129292492 | 129292492 | very-long-chain 3-oxoacyl-CoA reductase 1-like | -1.4 | 5.97E-03 |

| | | | | |
|---|---|---|---|---|
| LOC129308801 | 129308801 | uncharacterized LOC129308801 | -1.4 | 4.99E-03 |
| LOC129293368 | 129293368 | pentatricopeptide repeat-containing protein At4g21065-like | -1.4 | 3.01E-02 |
| LOC129321945 | 129321945 | protein PSK SIMULATOR 1-like | -1.4 | 3.23E-02 |
| LOC129316328 | 129316328 | protein trichome birefringence-like 39 | -1.4 | 6.19E-03 |
| LOC129302380 | 129302380 | pentatricopeptide repeat-containing protein At5g27460 | -1.4 | 5.81E-03 |
| LOC129295527 | 129295527 | rhodanese-like domain-containing protein 4, chloroplastic | -1.4 | 1.11E-02 |
| LOC129316796 | 129316796 | E3 ubiquitin-protein ligase RMA1H1-like | -1.4 | 6.79E-03 |
| LOC129302881 | 129302881 | pyridoxal reductase, chloroplastic | -1.4 | 9.91E-05 |
| LOC129294446 | 129294446 | transcription factor bHLH143-like | -1.4 | 3.36E-04 |
| LOC129322933 | 129322933 | uncharacterized LOC129322933 | -1.4 | 2.83E-02 |
| LOC129320881 | 129320881 | pyrophosphate-energized vacuolar membrane proton pump | -1.4 | 1.16E-04 |
| LOC129303256 | 129303256 | uncharacterized LOC129303256 | -1.4 | 8.50E-03 |
| LOC129309298 | 129309298 | low affinity inorganic phosphate transporter 1-like | -1.4 | 2.70E-05 |
| LOC129297670 | 129297670 | 8-hydroxygeraniol oxidoreductase-like | -1.4 | 1.94E-03 |
| LOC129291200 | 129291200 | F-box/kelch-repeat protein At1g51550 | -1.4 | 3.52E-02 |
| LOC129312835 | 129312835 | uncharacterized LOC129312835 | -1.4 | 3.70E-02 |
| LOC129320180 | 129320180 | uncharacterized LOC129320180 | -1.4 | 3.31E-03 |
| LOC129311115 | 129311115 | putative pentatricopeptide repeat-containing protein At5g65820 | -1.5 | 2.84E-02 |
| LOC129291749 | 129291749 | uncharacterized LOC129291749 | -1.5 | 1.59E-02 |
| LOC129309098 | 129309098 | probable methyltransferase PMT2 | -1.5 | 4.75E-02 |
| LOC129293108 | 129293108 | probable RNA-dependent RNA polymerase 3 | -1.5 | 2.79E-02 |
| LOC129308591 | 129308591 | RNA polymerase II C-terminal domain phosphatase-like 3 | -1.5 | 7.41E-04 |
| LOC129317123 | 129317123 | uncharacterized LOC129317123 | -1.5 | 1.74E-03 |
| LOC129307426 | 129307426 | LEC14B protein | -1.5 | 6.15E-03 |
| LOC129297095 | 129297095 | CDP-diacylglycerol--serine O-phosphatidyltransferase 1-like | -1.5 | 2.75E-02 |
| LOC129313403 | 129313403 | thiosulfate/3-mercaptopyruvate sulfurtransferase 1, mitochondri | -1.5 | 1.98E-02 |
| LOC129308426 | 129308426 | protein TIME FOR COFFEE-like | -1.5 | 1.08E-03 |
| LOC129320823 | 129320823 | transcription termination factor MTEF1, chloroplastic | -1.5 | 3.52E-02 |
| LOC129303268 | 129303268 | NAD(P)H-quinone oxidoreductase subunit N, chloroplastic | -1.5 | 4.23E-03 |
| LOC129308270 | 129308270 | mediator of RNA polymerase II transcription subunit 33A-like | -1.5 | 8.31E-03 |
| LOC129315151 | 129315151 | sterol 24-C-methyltransferase-like | -1.5 | 2.48E-02 |
| LOC129301947 | 129301947 | methyl-CpG-binding domain-containing protein 8-like | -1.5 | 6.67E-03 |
| LOC129303421 | 129303421 | bZIP transcription factor 44-like | -1.5 | 2.40E-02 |
| LOC129322011 | 129322011 | protein ABCI7, chloroplastic | -1.5 | 3.32E-04 |
| LOC129292165 | 129292165 | rhicadhesin receptor-like | -1.5 | 1.75E-02 |
| LOC129306433 | 129306433 | uncharacterized LOC129306433 | -1.5 | 5.08E-03 |
| LOC129295448 | 129295448 | uncharacterized LOC129295448 | -1.5 | 1.71E-03 |
| LOC129314786 | 129314786 | zinc finger protein BALDIBIS-like | -1.5 | 1.46E-02 |
| LOC129299266 | 129299266 | zinc finger protein CONSTANS-LIKE 5-like | -1.5 | 4.30E-04 |
| LOC129316101 | 129316101 | uncharacterized LOC129316101 | -1.5 | 2.74E-02 |
| LOC129311323 | 129311323 | flavonoid 3'-monooxygenase CYP75B137-like | -1.5 | 7.41E-03 |
| LOC129304852 | 129304852 | 4-alpha-glucanotransferase DPE2 | -1.5 | 1.38E-03 |
| LOC129315551 | 129315551 | putative disease resistance RPP13-like protein 1 | -1.5 | 3.08E-02 |
| LOC129311217 | 129311217 | protein NLP5-like | -1.5 | 1.24E-02 |
| LOC129285628 | 129285628 | magnesium transporter MRS2-I-like | -1.5 | 6.36E-06 |
| LOC129314151 | 129314151 | nudix hydrolase 20, chloroplastic-like | -1.5 | 2.66E-02 |
| LOC129299649 | 129299649 | desmethyl-deoxy-podophyllotoxin synthase-like | -1.5 | 4.93E-02 |
| LOC129304454 | 129304454 | transcription factor bHLH130-like | -1.5 | 8.67E-03 |

| | | | | |
|---|---|---|---|---|
| LOC129288962 | 129288962 | BTB/POZ domain and ankyrin repeat-containing protein NOOT | -1.5 | 5.81E-03 |
| LOC129312799 | 129312799 | uncharacterized LOC129312799 | -1.5 | 9.09E-03 |
| LOC129308474 | 129308474 | purple acid phosphatase 23 | -1.5 | 1.11E-03 |
| LOC129295749 | 129295749 | E3 ubiquitin-protein ligase MBR2-like | -1.5 | 1.37E-03 |
| LOC129285310 | 129285310 | cysteine-rich receptor-like protein kinase 25 | -1.5 | 4.94E-03 |
| LOC129319237 | 129319237 | calmodulin calcium-dependent NAD kinase-like | -1.5 | 2.94E-02 |
| LOC129293471 | 129293471 | zinc finger protein 7-like | -1.5 | 4.18E-02 |
| LOC129299151 | 129299151 | uncharacterized LOC129299151 | -1.5 | 5.14E-04 |
| LOC129304680 | 129304680 | double-stranded RNA-binding protein 1-like | -1.5 | 1.55E-02 |
| LOC129304049 | 129304049 | uncharacterized LOC129304049 | -1.5 | 4.64E-02 |
| LOC129295914 | 129295914 | probable galacturonosyltransferase-like 3 | -1.5 | 2.47E-02 |
| LOC129307382 | 129307382 | beta carbonic anhydrase 5, chloroplastic-like | -1.5 | 4.03E-03 |
| LOC129294812 | 129294812 | phosphoglucan phosphatase LSF2, chloroplastic | -1.5 | 1.30E-02 |
| LOC129291346 | 129291346 | squamosa promoter-binding-like protein 8 | -1.5 | 3.84E-02 |
| LOC129293439 | 129293439 | sugar transport protein 14-like | -1.5 | 7.94E-04 |
| LOC129309102 | 129309102 | chalcone synthase 1-like | -1.5 | 1.54E-03 |
| LOC129311109 | 129311109 | G-type lectin S-receptor-like serine/threonine-protein kinase At1 | -1.5 | 1.84E-05 |
| LOC129288539 | 129288539 | pentatricopeptide repeat-containing protein At3g26630, chloropl | -1.5 | 7.07E-03 |
| LOC129294633 | 129294633 | uncharacterized LOC129294633 | -1.5 | 4.60E-02 |
| LOC129302320 | 129302320 | low affinity sulfate transporter 3-like | -1.5 | 2.62E-05 |
| LOC129290120 | 129290120 | uncharacterized LOC129290120 | -1.5 | 1.75E-02 |
| LOC129321748 | 129321748 | probable methyltransferase PMT14 | -1.5 | 1.12E-02 |
| LOC129306693 | 129306693 | uncharacterized LOC129306693 | -1.5 | 1.06E-02 |
| LOC129310916 | 129310916 | alpha-1,4 glucan phosphorylase L-2 isozyme, chloroplastic/amyl | -1.5 | 1.07E-03 |
| LOC129291133 | 129291133 | sanguinarine reductase-like | -1.5 | 6.89E-04 |
| LOC129310984 | 129310984 | light-harvesting complex-like protein OHP2, chloroplastic | -1.5 | 4.43E-02 |
| LOC129308644 | 129308644 | SNF1-related protein kinase regulatory subunit gamma-1 | -1.5 | 3.65E-02 |
| LOC129307365 | 129307365 | cyclic nucleotide-gated ion channel 2-like | -1.5 | 2.25E-02 |
| LOC129294145 | 129294145 | sulfite exporter TauE/SafE family protein 3-like | -1.5 | 1.17E-02 |
| LOC129316244 | 129316244 | UDP-xylose transporter 1-like | -1.5 | 3.41E-02 |
| LOC129293459 | 129293459 | uncharacterized LOC129293459 | -1.5 | 1.22E-02 |
| LOC129302352 | 129302352 | F-box protein At5g07610-like | -1.5 | 3.30E-02 |
| LOC129322239 | 129322239 | LRR receptor-like serine/threonine-protein kinase FEI 1 | -1.5 | 1.89E-02 |
| LOC129323183 | 129323183 | uncharacterized LOC129323183 | -1.5 | 1.67E-03 |
| LOC129321574 | 129321574 | clp protease adapter protein ClpF, chloroplastic-like | -1.5 | 1.36E-02 |
| LOC129306647 | 129306647 | endonuclease 1-like | -1.5 | 2.49E-02 |
| LOC129315387 | 129315387 | protein NARROW LEAF 1 | -1.5 | 2.38E-03 |
| LOC129290407 | 129290407 | threonine synthase, chloroplastic-like | -1.5 | 1.29E-03 |
| LOC129318084 | 129318084 | nicotinamide/nicotinic acid mononucleotide adenylyltransferase- | -1.5 | 8.06E-03 |
| LOC129303554 | 129303554 | synaptonemal complex protein 1-like | -1.5 | 3.56E-03 |
| LOC129309637 | 129309637 | probable serine/threonine-protein kinase WNK9 | -1.5 | 2.78E-02 |
| LOC129320680 | 129320680 | bidirectional sugar transporter N3-like | -1.5 | 4.25E-02 |
| LOC129294271 | 129294271 | aspartic proteinase Asp1-like | -1.5 | 3.28E-02 |
| LOC129318485 | 129318485 | uncharacterized LOC129318485 | -1.5 | 1.03E-02 |
| LOC129309166 | 129309166 | HIPL1 protein | -1.5 | 3.50E-04 |
| LOC129291288 | 129291288 | serine carboxypeptidase-like 20 | -1.5 | 1.10E-03 |
| LOC129307711 | 129307711 | homeobox-leucine zipper protein ATHB-6-like | -1.5 | 2.67E-02 |
| LOC129313573 | 129313573 | 3-phosphoinositide-dependent protein kinase 2-like | -1.5 | 2.38E-02 |

| | | | | |
|---|---|---|---|---|
| LOC129299594 | 129299594 | transcription termination factor MTERF4, chloroplastic-like | -1.5 | 9.24E-03 |
| LOC129317102 | 129317102 | KH domain-containing protein At3g08620-like | -1.5 | 7.08E-05 |
| LOC129288535 | 129288535 | scarecrow-like protein 27 | -1.5 | 1.83E-05 |
| LOC129306988 | 129306988 | uncharacterized LOC129306988 | -1.5 | 1.15E-02 |
| LOC129291226 | 129291226 | auxin transporter-like protein 4 | -1.5 | 4.39E-03 |
| LOC129322877 | 129322877 | homeobox-leucine zipper protein ATHB-13-like | -1.6 | 2.04E-02 |
| LOC129322212 | 129322212 | probable purine permease 11 | -1.6 | 3.27E-02 |
| LOC129322373 | 129322373 | pheophytinase, chloroplastic | -1.6 | 3.99E-03 |
| LOC129311858 | 129311858 | uncharacterized LOC129311858 | -1.6 | 8.06E-03 |
| LOC129322644 | 129322644 | floral homeotic protein APETALA 2-like | -1.6 | 7.29E-03 |
| LOC129308600 | 129308600 | glycosyl hydrolase 5 family protein | -1.6 | 1.99E-03 |
| LOC129284595 | 129284595 | NAD(P)H-quinone oxidoreductase subunit S, chloroplastic | -1.6 | 1.38E-02 |
| LOC129322843 | 129322843 | receptor protein-tyrosine kinase CEPR2 | -1.6 | 3.02E-02 |
| LOC129304604 | 129304604 | tyrosine decarboxylase 2 | -1.6 | 7.61E-04 |
| LOC129308420 | 129308420 | auxin response factor 19 | -1.6 | 2.48E-03 |
| LOC129295984 | 129295984 | uncharacterized LOC129295984 | -1.6 | 1.95E-02 |
| LOC129320144 | 129320144 | probable pectinesterase/pectinesterase inhibitor 51 | -1.6 | 3.44E-03 |
| LOC129314238 | 129314238 | homeobox-leucine zipper protein HDG2-like | -1.6 | 1.72E-02 |
| LOC129290635 | 129290635 | hydroquinone glucosyltransferase-like | -1.6 | 3.72E-02 |
| LOC129287142 | 129287142 | uncharacterized LOC129287142 | -1.6 | 2.51E-02 |
| LOC129322836 | 129322836 | kinesin-like protein KIN-4A | -1.6 | 5.30E-03 |
| LOC129309728 | 129309728 | carboxyl-terminal-processing peptidase 2, chloroplastic | -1.6 | 4.08E-03 |
| LOC129291277 | 129291277 | beta-glucuronosyltransferase GlcAT14C-like | -1.6 | 4.63E-02 |
| LOC129322243 | 129322243 | probable methyltransferase PMT2 | -1.6 | 7.26E-04 |
| LOC129287718 | 129287718 | protein trichome birefringence-like 35 | -1.6 | 2.65E-03 |
| LOC129313140 | 129313140 | IRK-interacting protein | -1.6 | 1.67E-02 |
| LOC129300239 | 129300239 | uncharacterized LOC129300239 | -1.6 | 7.77E-03 |
| LOC129309383 | 129309383 | squamosa promoter-binding-like protein 2 | -1.6 | 2.25E-02 |
| LOC129291873 | 129291873 | zinc finger protein 8-like | -1.6 | 3.50E-02 |
| LOC129305242 | 129305242 | thioredoxin reductase NTRC-like | -1.6 | 1.21E-05 |
| LOC129317717 | 129317717 | GDSL esterase/lipase EXL3-like | -1.6 | 2.98E-02 |
| LOC129311418 | 129311418 | cellulose synthase-like protein D3 | -1.6 | 2.45E-03 |
| LOC129295719 | 129295719 | protein STRICTOSIDINE SYNTHASE-LIKE 4-like | -1.6 | 4.18E-03 |
| LOC129313488 | 129313488 | beta-amylase-like | -1.6 | 1.83E-04 |
| LOC129319487 | 129319487 | 2-deoxy-glucose resistant protein 2-like | -1.6 | 3.91E-03 |
| LOC129298244 | 129298244 | receptor-like protein kinase FERONIA | -1.6 | 4.96E-02 |
| LOC129302850 | 129302850 | 4-diphosphocytidyl-2-C-methyl-D-erythritol kinase, chloroplasti | -1.6 | 1.06E-02 |
| LOC129285726 | 129285726 | dicarboxylate transporter 2.1, chloroplastic-like | -1.6 | 8.76E-04 |
| LOC129319222 | 129319222 | protein PLASTID MOVEMENT IMPAIRED 1 | -1.6 | 3.04E-03 |
| LOC129317169 | 129317169 | uncharacterized LOC129317169 | -1.6 | 1.05E-02 |
| LOC129311656 | 129311656 | uncharacterized LOC129311656 | -1.6 | 4.10E-02 |
| LOC129291614 | 129291614 | protein SUPPRESSOR OF PHYA-105 1 | -1.6 | 1.43E-02 |
| LOC129319778 | 129319778 | phytyl ester synthase 2, chloroplastic-like | -1.6 | 9.70E-03 |
| LOC129288677 | 129288677 | putative DEAD-box ATP-dependent RNA helicase 33 | -1.6 | 1.02E-04 |
| LOC129303566 | 129303566 | geranylgeranyl pyrophosphate synthase 7, chloroplastic-like | -1.6 | 2.59E-02 |
| LOC129286700 | 129286700 | ferredoxin--NADP reductase, root isozyme, chloroplastic | -1.6 | 4.35E-02 |
| LOC129322990 | 129322990 | protein ACCUMULATION AND REPLICATION OF CHLORO | -1.6 | 1.16E-02 |
| LOC129305271 | 129305271 | transcription factor MYB60-like | -1.6 | 7.03E-03 |

| | | | | |
|---|---|---|---|---|
| LOC129302276 | 129302276 | BTB/POZ domain-containing protein NPY1-like | -1.6 | 3.45E-03 |
| LOC129313541 | 129313541 | monofunctional riboflavin biosynthesis protein RIBA 3, chlorop | -1.6 | 1.06E-03 |
| LOC129315992 | 129315992 | conserved oligomeric Golgi complex subunit 7 | -1.6 | 1.62E-02 |
| LOC129302999 | 129302999 | chitinase 2-like | -1.6 | 3.43E-03 |
| LOC129306327 | 129306327 | AUGMIN subunit 6-like | -1.6 | 7.77E-03 |
| LOC129296043 | 129296043 | enoyl-[acyl-carrier-protein] reductase [NADH], chloroplastic-lik | -1.6 | 4.35E-02 |
| LOC129321087 | 129321087 | uncharacterized LOC129321087 | -1.6 | 4.15E-02 |
| LOC129314522 | 129314522 | glucosamine inositolphosphorylceramide transferase 1 | -1.6 | 2.03E-03 |
| LOC129304136 | 129304136 | protein NPGR1 | -1.6 | 1.55E-02 |
| LOC129322788 | 129322788 | protein ESMERALDA 1-like | -1.6 | 1.78E-03 |
| LOC129309711 | 129309711 | transcription factor IBH1 | -1.6 | 1.62E-02 |
| LOC129290156 | 129290156 | RING-H2 finger protein ATL39-like | -1.6 | 2.05E-03 |
| LOC129291605 | 129291605 | probable glycerol-3-phosphate acyltransferase 8 | -1.6 | 1.03E-02 |
| LOC129290169 | 129290169 | D-glycerate 3-kinase, chloroplastic | -1.6 | 1.70E-04 |
| LOC129300575 | 129300575 | probable alpha-mannosidase At5g13980 | -1.6 | 2.19E-02 |
| LOC129307919 | 129307919 | uncharacterized LOC129307919 | -1.6 | 1.50E-02 |
| LOC129307752 | 129307752 | alkaline/neutral invertase A, mitochondrial-like | -1.6 | 3.06E-03 |
| LOC129315260 | 129315260 | putative protein FAR1-RELATED SEQUENCE 10 | -1.6 | 3.80E-02 |
| LOC129286861 | 129286861 | uncharacterized LOC129286861 | -1.6 | 1.19E-02 |
| LOC129294390 | 129294390 | pentatricopeptide repeat-containing protein At3g24000, mitocho | -1.6 | 3.47E-02 |
| LOC129288967 | 129288967 | UDP-glycosyltransferase 74G1-like | -1.6 | 4.50E-03 |
| LOC129322020 | 129322020 | monodehydroascorbate reductase, chloroplastic/mitochondrial | -1.6 | 3.06E-03 |
| LOC129322014 | 129322014 | RING-H2 finger protein ATL52-like | -1.6 | 2.01E-03 |
| LOC129309399 | 129309399 | agmatine deiminase | -1.6 | 1.58E-04 |
| LOC129294634 | 129294634 | glycerol-3-phosphate dehydrogenase [NAD(+)] 2, chloroplastic | -1.6 | 1.54E-02 |
| LOC129307012 | 129307012 | beta-carotene isomerase D27, chloroplastic | -1.6 | 7.80E-06 |
| LOC129287400 | 129287400 | uncharacterized LOC129287400 | -1.6 | 2.94E-02 |
| LOC129320170 | 129320170 | brassinosteroid LRR receptor kinase-like | -1.6 | 5.78E-03 |
| LOC129322700 | 129322700 | amino acid transporter ANT1 | -1.6 | 1.06E-03 |
| LOC129321994 | 129321994 | serine carboxypeptidase-like 45 | -1.6 | 2.81E-03 |
| LOC129322333 | 129322333 | protein indeterminate-domain 5, chloroplastic-like | -1.6 | 1.47E-06 |
| LOC129290683 | 129290683 | F-box protein At2g39490 | -1.6 | 2.84E-03 |
| LOC129313126 | 129313126 | uncharacterized LOC129313126 | -1.6 | 3.62E-03 |
| LOC129291098 | 129291098 | uncharacterized LOC129291098 | -1.6 | 1.48E-03 |
| LOC129286712 | 129286712 | guanine nucleotide-binding protein subunit beta-2 | -1.6 | 2.86E-04 |
| LOC129286616 | 129286616 | tyrosine--tRNA ligase, chloroplastic/mitochondrial-like | -1.6 | 2.54E-03 |
| LOC129315802 | 129315802 | glutamine synthetase nodule isozyme-like | -1.6 | 1.76E-02 |
| LOC129322260 | 129322260 | protein COFACTOR ASSEMBLY OF COMPLEX C SUBUNI | -1.6 | 4.36E-02 |
| LOC129320852 | 129320852 | ras-related protein Rab7-like | -1.6 | 4.34E-02 |
| LOC129297038 | 129297038 | hydroxymethylglutaryl-CoA lyase, mitochondrial-like | -1.6 | 1.25E-02 |
| LOC129299842 | 129299842 | nudix hydrolase 18, mitochondrial-like | -1.6 | 2.71E-05 |
| LOC129291472 | 129291472 | photosynthetic NDH subunit of subcomplex B 1, chloroplastic-l | -1.6 | 2.57E-02 |
| LOC129299540 | 129299540 | heavy metal-associated isoprenylated plant protein 7-like | -1.7 | 1.58E-02 |
| LOC129285307 | 129285307 | cysteine-rich receptor-like protein kinase 10 | -1.7 | 2.19E-03 |
| LOC129304186 | 129304186 | uncharacterized LOC129304186 | -1.7 | 4.90E-02 |
| LOC129320485 | 129320485 | probable aspartic proteinase GIP2 | -1.7 | 4.77E-02 |
| LOC129300335 | 129300335 | ribosome-binding factor PSRP1, chloroplastic-like | -1.7 | 1.29E-02 |
| LOC129318899 | 129318899 | uncharacterized LOC129318899 | -1.7 | 2.14E-02 |

| | | | | |
|---|---|---|---|---|
| LOC129290775 | 129290775 | uncharacterized LOC129290775 | -1.7 | 2.57E-02 |
| LOC129290792 | 129290792 | uncharacterized LOC129290792 | -1.7 | 4.23E-03 |
| LOC129315272 | 129315272 | probable BOI-related E3 ubiquitin-protein ligase 2 | -1.7 | 1.04E-03 |
| LOC129320496 | 129320496 | pentatricopeptide repeat-containing protein At4g30825, chloropl | -1.7 | 4.94E-03 |
| LOC129285057 | 129285057 | berberine bridge enzyme-like 13 | -1.7 | 2.12E-02 |
| LOC129307082 | 129307082 | uncharacterized protein At5g39570 | -1.7 | 1.33E-02 |
| LOC129323154 | 129323154 | polygalacturonase 1 beta-like protein 3 | -1.7 | 4.33E-02 |
| LOC129293576 | 129293576 | non-specific lipid-transfer protein 2-like | -1.7 | 7.16E-03 |
| LOC129292063 | 129292063 | uncharacterized LOC129292063 | -1.7 | 3.32E-04 |
| LOC129321898 | 129321898 | UDP-galactose/UDP-glucose transporter 3 | -1.7 | 2.20E-02 |
| LOC129303044 | 129303044 | receptor protein-tyrosine kinase CEPR1 | -1.7 | 4.47E-04 |
| LOC129313672 | 129313672 | actin | -1.7 | 1.22E-02 |
| LOC129312085 | 129312085 | uncharacterized LOC129312085 | -1.7 | 1.48E-02 |
| LOC129284460 | 129284460 | protein root UVB sensitive 1, chloroplastic | -1.7 | 2.20E-05 |
| LOC129287850 | 129287850 | NDR1/HIN1-like protein 13 | -1.7 | 8.50E-03 |
| LOC129303005 | 129303005 | BTB/POZ domain-containing protein At2g30600 | -1.7 | 7.85E-03 |
| LOC129288506 | 129288506 | mogroside IE synthase-like | -1.7 | 5.45E-05 |
| LOC129306710 | 129306710 | nudix hydrolase 12, mitochondrial-like | -1.7 | 8.00E-03 |
| LOC129311310 | 129311310 | legumain-like | -1.7 | 3.44E-02 |
| LOC129294162 | 129294162 | CLP protease regulatory subunit CLPX1, mitochondrial-like | -1.7 | 1.26E-02 |
| LOC129305811 | 129305811 | probable ubiquitin-conjugating enzyme E2 24 | -1.7 | 6.80E-06 |
| LOC129312819 | 129312819 | uncharacterized LOC129312819 | -1.7 | 4.29E-06 |
| LOC129310592 | 129310592 | uncharacterized LOC129310592 | -1.7 | 5.03E-04 |
| LOC129293328 | 129293328 | cysteine-rich receptor-like protein kinase 29 | -1.7 | 2.62E-02 |
| LOC129311831 | 129311831 | ethylene-responsive transcription factor 5-like | -1.7 | 1.09E-03 |
| LOC129288096 | 129288096 | U-box domain-containing protein 44-like | -1.7 | 2.26E-03 |
| LOC129304117 | 129304117 | uncharacterized LOC129304117 | -1.7 | 3.03E-02 |
| LOC129289553 | 129289553 | glycerol-3-phosphate 2-O-acyltransferase 6-like | -1.7 | 3.97E-03 |
| LOC129317122 | 129317122 | uncharacterized LOC129317122 | -1.7 | 4.03E-03 |
| LOC129306340 | 129306340 | pentatricopeptide repeat-containing protein At4g11690-like | -1.7 | 4.39E-03 |
| LOC129301674 | 129301674 | peptide-N4-(N-acetyl-beta-glucosaminyl)asparagine amidase A | -1.7 | 2.43E-04 |
| LOC129291331 | 129291331 | protein trichome birefringence-like 3 | -1.7 | 1.57E-02 |
| LOC129303081 | 129303081 | probable inorganic phosphate transporter 1-5 | -1.7 | 2.61E-02 |
| LOC129315764 | 129315764 | zinc finger CCCH domain-containing protein 53-like | -1.7 | 1.78E-02 |
| LOC129285865 | 129285865 | probable carboxylesterase 12 | -1.7 | 7.33E-04 |
| LOC129322647 | 129322647 | mediator of RNA polymerase II transcription subunit 25-like | -1.7 | 3.12E-02 |
| LOC129308673 | 129308673 | peroxidase A2-like | -1.7 | 4.37E-02 |
| LOC129284449 | 129284449 | metalloendoproteinase 2-MMP-like | -1.7 | 4.56E-02 |
| LOC129287875 | 129287875 | monocopper oxidase-like protein SKU5 | -1.7 | 1.37E-03 |
| LOC129298423 | 129298423 | cysteine-rich receptor-like protein kinase 10 | -1.7 | 2.08E-02 |
| LOC129305849 | 129305849 | fasciclin-like arabinogalactan protein 15 | -1.7 | 1.65E-02 |
| LOC129315118 | 129315118 | uncharacterized LOC129315118 | -1.7 | 1.05E-02 |
| LOC129292712 | 129292712 | transcription factor DICHOTOMA-like | -1.7 | 4.98E-02 |
| LOC129287547 | 129287547 | uncharacterized LOC129287547 | -1.7 | 4.01E-03 |
| LOC129303055 | 129303055 | uncharacterized LOC129303055 | -1.7 | 1.65E-04 |
| LOC129306904 | 129306904 | splicing factor-like protein 1 | -1.7 | 5.91E-03 |
| LOC129315395 | 129315395 | protein ZINC INDUCED FACILITATOR-LIKE 1-like | -1.7 | 1.66E-02 |
| LOC129302473 | 129302473 | pentatricopeptide repeat-containing protein At5g18475 | -1.7 | 2.86E-03 |

| | | | | |
|---|---|---|---|---|
| LOC129320879 | 129320879 | uncharacterized LOC129320879 | -1.7 | 1.03E-04 |
| LOC129293452 | 129293452 | uncharacterized LOC129293452 | -1.7 | 2.97E-02 |
| LOC129322115 | 129322115 | uncharacterized LOC129322115 | -1.7 | 4.60E-03 |
| LOC129311934 | 129311934 | probable LRR receptor-like serine/threonine-protein kinase At1g | -1.7 | 4.43E-02 |
| LOC129290544 | 129290544 | UDP-glycosyltransferase 74F2-like | -1.7 | 3.05E-03 |
| LOC129317887 | 129317887 | 6,7,8-trihydroxycoumarin synthase-like | -1.7 | 1.28E-02 |
| LOC129292785 | 129292785 | probable CoA ligase CCL6 | -1.7 | 1.20E-02 |
| LOC129290673 | 129290673 | BEACH domain-containing protein C2 | -1.7 | 2.16E-02 |
| LOC129292560 | 129292560 | probable pinoresinol-lariciresinol reductase 3 | -1.7 | 4.18E-03 |
| LOC129318766 | 129318766 | J domain-containing protein required for chloroplast accumulati | -1.7 | 1.07E-02 |
| LOC129313203 | 129313203 | uncharacterized LOC129313203 | -1.7 | 4.11E-02 |
| LOC129306663 | 129306663 | callose synthase 11 | -1.7 | 2.92E-02 |
| LOC129295562 | 129295562 | probable glucan 1,3-alpha-glucosidase | -1.7 | 2.65E-02 |
| LOC129290145 | 129290145 | disease resistance protein RPV1-like | -1.8 | 5.67E-04 |
| LOC129315983 | 129315983 | 3-ketoacyl-CoA synthase 3-like | -1.8 | 1.36E-03 |
| LOC129301969 | 129301969 | 3beta,22alpha-dihydroxysteroid 3-dehydrogenase | -1.8 | 4.03E-02 |
| LOC129285398 | 129285398 | NAC domain-containing protein 6-like | -1.8 | 8.31E-03 |
| LOC129308609 | 129308609 | glucose-1-phosphate adenylyltransferase large subunit 1, chlorop | -1.8 | 6.62E-07 |
| LOC129294972 | 129294972 | protein WVD2-like 7 | -1.8 | 1.45E-07 |
| LOC129312642 | 129312642 | O-fucosyltransferase 20-like | -1.8 | 3.16E-03 |
| LOC129312435 | 129312435 | transcription factor TCP4-like | -1.8 | 7.03E-03 |
| LOC129284610 | 129284610 | CDK5RAP1-like protein | -1.8 | 5.49E-04 |
| LOC129308111 | 129308111 | uncharacterized LOC129308111 | -1.8 | 3.26E-02 |
| LOC129288778 | 129288778 | uncharacterized LOC129288778 | -1.8 | 1.29E-07 |
| LOC129287848 | 129287848 | E3 ubiquitin-protein ligase AIRP2 | -1.8 | 2.97E-02 |
| LOC129297187 | 129297187 | uncharacterized GPI-anchored protein At1g61900-like | -1.8 | 2.53E-02 |
| LOC129311177 | 129311177 | uncharacterized LOC129311177 | -1.8 | 3.62E-02 |
| LOC129288568 | 129288568 | vacuolar cation/proton exchanger 3-like | -1.8 | 5.40E-03 |
| LOC129291417 | 129291417 | 4-hydroxybenzoate polyprenyltransferase, mitochondrial-like | -1.8 | 4.37E-02 |
| LOC129320297 | 129320297 | uncharacterized LOC129320297 | -1.8 | 1.88E-02 |
| LOC129287596 | 129287596 | probable serine/threonine-protein kinase At1g01540 | -1.8 | 1.32E-03 |
| LOC129304657 | 129304657 | phosphatidylinositol N-acetylglucosaminyltransferase subunit P- | -1.8 | 1.70E-02 |
| LOC129288650 | 129288650 | uncharacterized LOC129288650 | -1.8 | 3.04E-03 |
| LOC129308513 | 129308513 | cadmium/zinc-transporting ATPase HMA2-like | -1.8 | 8.70E-03 |
| LOC129320072 | 129320072 | CASP-like protein 5B3 | -1.8 | 4.39E-03 |
| LOC129288749 | 129288749 | transcription factor MYBS3-like | -1.8 | 4.23E-02 |
| LOC129291131 | 129291131 | universal stress protein PHOS32-like | -1.8 | 5.81E-04 |
| LOC129303350 | 129303350 | uncharacterized LOC129303350 | -1.8 | 5.65E-03 |
| LOC129307733 | 129307733 | phosphoglucan, water dikinase, chloroplastic | -1.8 | 2.35E-02 |
| LOC129311883 | 129311883 | probable beta-D-xylosidase 6 | -1.8 | 1.69E-02 |
| LOC129312153 | 129312153 | auxin efflux carrier component 3-like | -1.8 | 5.20E-04 |
| LOC129286247 | 129286247 | protein SCAR3 | -1.8 | 1.50E-02 |
| LOC129314444 | 129314444 | pentatricopeptide repeat-containing protein At2g27800, mitocho | -1.8 | 4.06E-02 |
| LOC129298329 | 129298329 | uncharacterized LOC129298329 | -1.8 | 1.70E-03 |
| LOC129315180 | 129315180 | uncharacterized LOC129315180 | -1.8 | 3.77E-03 |
| LOC129284318 | 129284318 | MLP-like protein 28 | -1.8 | 1.33E-03 |
| LOC129315883 | 129315883 | putative disease resistance protein RGA4 | -1.8 | 3.35E-02 |
| LOC129304095 | 129304095 | ent-kaurene oxidase-like | -1.8 | 2.26E-03 |

| | | | | |
|---|---|---|---|---|
| LOC129288987 | 129288987 | protein trichome birefringence-like 3 | -1.8 | 4.33E-02 |
| LOC129306120 | 129306120 | DEAD-box ATP-dependent RNA helicase 39-like | -1.8 | 6.96E-04 |
| LOC129294638 | 129294638 | beta-galactosidase 16 | -1.8 | 3.41E-02 |
| LOC129316804 | 129316804 | probable F-box protein At2g36090 | -1.8 | 3.90E-02 |
| LOC129304040 | 129304040 | methylesterase 17-like | -1.8 | 6.94E-03 |
| LOC129291651 | 129291651 | uncharacterized LOC129291651 | -1.8 | 2.12E-03 |
| LOC129319386 | 129319386 | uncharacterized LOC129319386 | -1.8 | 3.94E-03 |
| LOC129320544 | 129320544 | superoxide dismutase [Fe], chloroplastic-like | -1.8 | 5.65E-03 |
| LOC129311978 | 129311978 | metal transporter Nramp3-like | -1.8 | 4.03E-04 |
| LOC129286977 | 129286977 | catalase isozyme 1-like | -1.8 | 1.26E-07 |
| LOC129305926 | 129305926 | UDP-glycosyltransferase 73C4-like | -1.8 | 3.15E-03 |
| LOC129307262 | 129307262 | protein ASPARTIC PROTEASE IN GUARD CELL 1-like | -1.8 | 1.16E-02 |
| LOC129285565 | 129285565 | pentatricopeptide repeat-containing protein At5g55840 | -1.8 | 2.21E-02 |
| LOC129302032 | 129302032 | transcription repressor KAN1 | -1.8 | 2.13E-02 |
| LOC129300444 | 129300444 | VQ motif-containing protein 9-like | -1.8 | 1.47E-02 |
| LOC129301220 | 129301220 | probable LRR receptor-like serine/threonine-protein kinase At2g | -1.8 | 3.15E-02 |
| LOC129308207 | 129308207 | serine/threonine/tyrosine-protein kinase HT1-like | -1.8 | 3.00E-03 |
| LOC129307455 | 129307455 | probable inactive leucine-rich repeat receptor-like protein kinase | -1.8 | 2.35E-03 |
| LOC129322094 | 129322094 | tRNA(adenine(34)) deaminase, chloroplastic | -1.8 | 6.12E-03 |
| LOC129294478 | 129294478 | uncharacterized LOC129294478 | -1.8 | 2.92E-05 |
| LOC129310395 | 129310395 | subtilisin-like serine-protease S | -1.8 | 1.22E-03 |
| LOC129295371 | 129295371 | serine/threonine/tyrosine-protein kinase HT1-like | -1.8 | 6.59E-03 |
| LOC129317876 | 129317876 | zinc transporter 11-like | -1.8 | 1.81E-02 |
| LOC129303881 | 129303881 | transcription factor bHLH74-like | -1.8 | 1.11E-03 |
| LOC129291591 | 129291591 | uncharacterized LOC129291591 | -1.8 | 2.41E-02 |
| LOC129313809 | 129313809 | aquaporin SIP1-1-like | -1.8 | 5.20E-03 |
| LOC129320426 | 129320426 | transcription factor MYBS1 | -1.8 | 5.59E-04 |
| LOC129304976 | 129304976 | ABC transporter G family member 1-like | -1.8 | 2.30E-05 |
| LOC129297364 | 129297364 | uncharacterized LOC129297364 | -1.8 | 6.60E-03 |
| LOC129303660 | 129303660 | probable plastidic glucose transporter 1 | -1.8 | 2.38E-03 |
| LOC129315919 | 129315919 | diacylglycerol kinase 5-like | -1.8 | 2.04E-07 |
| LOC129315498 | 129315498 | protein SIEL-like | -1.8 | 3.15E-03 |
| LOC129295491 | 129295491 | probable serine/threonine-protein kinase At1g01540 | -1.8 | 2.17E-03 |
| LOC129284493 | 129284493 | uncharacterized LOC129284493 | -1.8 | 1.56E-02 |
| LOC129291556 | 129291556 | scarecrow-like protein 6 | -1.8 | 1.54E-02 |
| LOC129321039 | 129321039 | uncharacterized LOC129321039 | -1.8 | 1.11E-03 |
| LOC129308571 | 129308571 | plasma membrane ATPase 4 | -1.8 | 1.93E-06 |
| LOC129308243 | 129308243 | protein CELLULOSE SYNTHASE INTERACTIVE 1-like | -1.8 | 3.73E-04 |
| LOC129285977 | 129285977 | leucine-rich repeat extensin-like protein 3 | -1.8 | 1.40E-02 |
| LOC129303596 | 129303596 | protein DOWNY MILDEW RESISTANCE 6-like | -1.8 | 2.79E-02 |
| LOC129308713 | 129308713 | ATPase 11, plasma membrane-type | -1.9 | 4.42E-07 |
| LOC129307018 | 129307018 | uncharacterized LOC129307018 | -1.9 | 2.88E-04 |
| LOC129320339 | 129320339 | uncharacterized LOC129320339 | -1.9 | 1.01E-03 |
| LOC129290749 | 129290749 | protein REDUCED CHLOROPLAST COVERAGE 1 | -1.9 | 4.18E-03 |
| LOC129293359 | 129293359 | probable ADP-ribosylation factor GTPase-activating protein AG | -1.9 | 1.05E-02 |
| LOC129309619 | 129309619 | G-type lectin S-receptor-like serine/threonine-protein kinase At4 | -1.9 | 5.31E-03 |
| LOC129303385 | 129303385 | uncharacterized LOC129303385 | -1.9 | 1.07E-03 |
| LOC129291699 | 129291699 | molybdate transporter 2 | -1.9 | 1.94E-06 |

| | | | | |
|---|---|---|---|---|
| LOC129290864 | 129290864 | nitrate regulatory gene2 protein-like | -1.9 | 2.28E-03 |
| LOC129313778 | 129313778 | endochitinase PR4-like | -1.9 | 6.60E-03 |
| LOC129293144 | 129293144 | uncharacterized LOC129293144 | -1.9 | 1.07E-03 |
| LOC129287755 | 129287755 | cyclic nucleotide-gated ion channel 1-like | -1.9 | 4.76E-02 |
| LOC129302302 | 129302302 | plasma membrane ATPase 4 | -1.9 | 7.92E-06 |
| LOC129319814 | 129319814 | probable WRKY transcription factor 31 | -1.9 | 6.89E-03 |
| LOC129292704 | 129292704 | zinc finger protein JACKDAW-like | -1.9 | 3.13E-04 |
| LOC129298974 | 129298974 | dicarboxylate transporter 2.1, chloroplastic-like | -1.9 | 1.27E-02 |
| LOC129306291 | 129306291 | uncharacterized LOC129306291 | -1.9 | 4.85E-04 |
| LOC129298627 | 129298627 | BTB/POZ domain-containing protein NPY5-like | -1.9 | 2.51E-02 |
| LOC129297224 | 129297224 | uncharacterized aarF domain-containing protein kinase At1g718 | -1.9 | 2.96E-02 |
| LOC129303502 | 129303502 | probable transcription factor KAN4 | -1.9 | 1.02E-02 |
| LOC129304754 | 129304754 | heavy metal-associated isoprenylated plant protein 6-like | -1.9 | 2.54E-05 |
| LOC129303628 | 129303628 | non-specific lipid transfer protein GPI-anchored 2-like | -1.9 | 1.42E-02 |
| LOC129292650 | 129292650 | strigolactone esterase D14-like | -1.9 | 3.27E-08 |
| LOC129310679 | 129310679 | formyltetrahydrofolate deformylase 1, mitochondrial-like | -1.9 | 4.43E-02 |
| LOC129287521 | 129287521 | uncharacterized LOC129287521 | -1.9 | 1.93E-04 |
| LOC129305603 | 129305603 | 2-Cys peroxiredoxin BAS1, chloroplastic-like | -1.9 | 3.93E-02 |
| LOC129301821 | 129301821 | membrane protein of ER body-like protein | -1.9 | 3.17E-02 |
| LOC129307756 | 129307756 | uncharacterized LOC129307756 | -1.9 | 2.32E-03 |
| LOC129309539 | 129309539 | uncharacterized LOC129309539 | -1.9 | 1.74E-03 |
| LOC129314427 | 129314427 | uncharacterized LOC129314427 | -1.9 | 3.70E-02 |
| LOC129305246 | 129305246 | uncharacterized LOC129305246 | -1.9 | 2.69E-02 |
| LOC129285193 | 129285193 | transcription factor TCP8 | -1.9 | 3.00E-03 |
| LOC129307314 | 129307314 | uncharacterized LOC129307314 | -1.9 | 3.98E-02 |
| LOC129320231 | 129320231 | probable serine/threonine-protein kinase SIS8 | -1.9 | 2.25E-06 |
| LOC129288814 | 129288814 | D-3-phosphoglycerate dehydrogenase 2, chloroplastic-like | -1.9 | 2.85E-03 |
| LOC129311513 | 129311513 | LOB domain-containing protein 38-like | -1.9 | 1.30E-03 |
| LOC129290370 | 129290370 | uncharacterized LOC129290370 | -1.9 | 7.17E-05 |
| LOC129318848 | 129318848 | uncharacterized LOC129318848 | -1.9 | 8.79E-04 |
| LOC129302472 | 129302472 | probable LRR receptor-like serine/threonine-protein kinase At1g | -1.9 | 3.80E-03 |
| LOC129317203 | 129317203 | uncharacterized LOC129317203 | -1.9 | 2.36E-03 |
| LOC129309726 | 129309726 | pentatricopeptide repeat-containing protein At2g18940, chloropl | -1.9 | 2.04E-04 |
| LOC129302163 | 129302163 | BRI1 kinase inhibitor 1-like | -1.9 | 1.01E-04 |
| LOC129301494 | 129301494 | 2-oxoglutarate-dependent dioxygenase 19-like | -1.9 | 2.47E-02 |
| LOC129293307 | 129293307 | 2-hydroxyisoflavanone dehydratase-like | -1.9 | 1.34E-02 |
| LOC129291344 | 129291344 | protein SUPPRESSOR OF K(+) TRANSPORT GROWTH DEI | -1.9 | 4.74E-06 |
| LOC129293735 | 129293735 | trihelix transcription factor DF1-like | -1.9 | 7.80E-06 |
| LOC129314595 | 129314595 | uncharacterized LOC129314595 | -1.9 | 5.70E-03 |
| LOC129288295 | 129288295 | alcohol dehydrogenase-like 4 | -1.9 | 1.42E-04 |
| LOC129319446 | 129319446 | calmodulin-binding protein 60 A-like | -1.9 | 2.08E-04 |
| LOC129317584 | 129317584 | CASP-like protein 2A2 | -1.9 | 4.75E-05 |
| LOC129294479 | 129294479 | syntaxin-31 | -1.9 | 1.35E-02 |
| LOC129308560 | 129308560 | serine/threonine-protein kinase STY13 | -1.9 | 3.21E-02 |
| LOC129292215 | 129292215 | alpha-galactosidase-like | -1.9 | 2.53E-04 |
| LOC129300226 | 129300226 | SCARECROW-LIKE protein 7-like | -2.0 | 1.29E-02 |
| LOC129301353 | 129301353 | fatty acid amide hydrolase-like | -2.0 | 3.02E-02 |
| LOC129296436 | 129296436 | uncharacterized LOC129296436 | -2.0 | 2.03E-03 |

| | | | | |
|---|---|---|---|---|
| LOC129320585 | 129320585 | thylakoid lumenal 16.5 kDa protein, chloroplastic-like | -2.0 | 4.41E-02 |
| LOC129311959 | 129311959 | probable xyloglucan endotransglucosylase/hydrolase protein 28 | -2.0 | 4.50E-04 |
| LOC129310841 | 129310841 | clp protease adapter protein ClpF, chloroplastic-like | -2.0 | 1.98E-02 |
| LOC129296175 | 129296175 | increased DNA methylation 3-like | -2.0 | 4.67E-02 |
| LOC129306585 | 129306585 | uncharacterized LOC129306585 | -2.0 | 2.66E-03 |
| LOC129318509 | 129318509 | uncharacterized LOC129318509 | -2.0 | 4.30E-10 |
| LOC129313332 | 129313332 | photosynthetic NDH subunit of lumenal location 3, chloroplastic | -2.0 | 1.63E-03 |
| LOC129286496 | 129286496 | histone H1-like | -2.0 | 2.23E-02 |
| LOC129320484 | 129320484 | transcription factor BIM1 | -2.0 | 9.77E-04 |
| LOC129288475 | 129288475 | MLO-like protein 1 | -2.0 | 8.90E-06 |
| LOC129301937 | 129301937 | uncharacterized LOC129301937 | -2.0 | 1.02E-02 |
| LOC129293672 | 129293672 | ferredoxin--nitrite reductase, chloroplastic-like | -2.0 | 3.94E-04 |
| LOC129286882 | 129286882 | uncharacterized LOC129286882 | -2.0 | 9.61E-03 |
| LOC129304928 | 129304928 | uncharacterized LOC129304928 | -2.0 | 3.00E-02 |
| LOC129313786 | 129313786 | probable carboxylesterase 15 | -2.0 | 2.20E-02 |
| LOC129321594 | 129321594 | peroxidase 35-like | -2.0 | 8.00E-06 |
| LOC129294124 | 129294124 | uncharacterized LOC129294124 | -2.0 | 4.85E-04 |
| LOC129308682 | 129308682 | uncharacterized LOC129308682 | -2.0 | 4.36E-02 |
| LOC129288522 | 129288522 | uncharacterized LOC129288522 | -2.0 | 5.58E-08 |
| LOC129290525 | 129290525 | uncharacterized LOC129290525 | -2.0 | 3.23E-02 |
| LOC129311234 | 129311234 | solanesyl diphosphate synthase 3, chloroplastic/mitochondrial-lil | -2.0 | 1.23E-04 |
| LOC129318019 | 129318019 | pentatricopeptide repeat-containing protein At3g22150, chloropl | -2.0 | 4.36E-02 |
| LOC129294086 | 129294086 | uncharacterized LOC129294086 | -2.0 | 1.86E-03 |
| LOC129288559 | 129288559 | PTI1-like tyrosine-protein kinase At3g15890 | -2.0 | 4.81E-03 |
| LOC129295467 | 129295467 | strigolactone esterase D14-like | -2.0 | 2.94E-04 |
| LOC129289259 | 129289259 | protein HIGH CHLOROPHYLL FLUORESCENCE PHENOTY | -2.0 | 6.06E-04 |
| LOC129312965 | 129312965 | probable LRR receptor-like serine/threonine-protein kinase At1g | -2.0 | 8.78E-03 |
| LOC129307025 | 129307025 | uncharacterized protein At2g23090-like | -2.0 | 3.16E-03 |
| LOC129317594 | 129317594 | fasciclin-like arabinogalactan protein 2 | -2.0 | 6.88E-04 |
| LOC129291274 | 129291274 | lysM domain receptor-like kinase 3 | -2.0 | 1.41E-03 |
| LOC129292141 | 129292141 | uncharacterized LOC129292141 | -2.0 | 3.26E-04 |
| LOC129290314 | 129290314 | BTB/POZ domain-containing protein At5g48800-like | -2.0 | 1.37E-04 |
| LOC129322722 | 129322722 | BTB/POZ domain-containing protein At1g67900-like | -2.0 | 1.23E-02 |
| LOC129318340 | 129318340 | pathogenesis-related thaumatin-like protein 3.5 | -2.0 | 5.51E-03 |
| LOC129315672 | 129315672 | auxin response factor 18-like | -2.0 | 1.27E-02 |
| LOC129288177 | 129288177 | G-type lectin S-receptor-like serine/threonine-protein kinase SD | -2.0 | 4.54E-03 |
| LOC129306742 | 129306742 | alkane hydroxylase MAH1-like | -2.0 | 5.48E-03 |
| LOC129315317 | 129315317 | putative disease resistance RPP13-like protein 1 | -2.0 | 2.00E-02 |
| LOC129295838 | 129295838 | basic leucine zipper 24-like | -2.0 | 1.81E-03 |
| LOC129317785 | 129317785 | probable galactinol--sucrose galactosyltransferase 1 | -2.0 | 3.83E-03 |
| LOC129301917 | 129301917 | cucumisin-like | -2.0 | 4.31E-07 |
| LOC129291702 | 129291702 | 2-methylene-furan-3-one reductase-like | -2.0 | 7.13E-04 |
| LOC129288195 | 129288195 | putative pentatricopeptide repeat-containing protein At3g23330 | -2.0 | 1.27E-02 |
| LOC129293426 | 129293426 | probable carboxylesterase 12 | -2.0 | 1.05E-04 |
| LOC129310375 | 129310375 | plastidal glycolate/glycerate translocator 1, chloroplastic-like | -2.0 | 8.47E-08 |
| LOC129315780 | 129315780 | protein FAR-RED IMPAIRED RESPONSE 1-like | -2.0 | 1.84E-02 |
| LOC129303663 | 129303663 | uncharacterized LOC129303663 | -2.0 | 7.69E-03 |
| LOC129291367 | 129291367 | protein ROOT HAIR DEFECTIVE 3 homolog 1-like | -2.0 | 4.65E-06 |

| | | | | |
|---|---|---|---|---|
| LOC129322567 | 129322567 | protein ENHANCED PSEUDOMONAS SUSCEPTIBILITY 1- | -2.0 | 1.96E-03 |
| LOC129308490 | 129308490 | protein NSP-INTERACTING KINASE 1-like | -2.0 | 9.00E-03 |
| LOC129315282 | 129315282 | probable WRKY transcription factor 14 | -2.1 | 1.04E-02 |
| LOC129288494 | 129288494 | uncharacterized LOC129288494 | -2.1 | 1.83E-03 |
| LOC129322995 | 129322995 | vacuolar iron transporter 1 | -2.1 | 4.27E-03 |
| LOC129291413 | 129291413 | zinc finger protein VAR3, chloroplastic-like | -2.1 | 5.80E-03 |
| LOC129296466 | 129296466 | uncharacterized LOC129296466 | -2.1 | 7.77E-03 |
| LOC129287140 | 129287140 | NAC domain-containing protein 35-like | -2.1 | 2.17E-02 |
| LOC129320042 | 129320042 | uncharacterized LOC129320042 | -2.1 | 7.59E-03 |
| LOC129304665 | 129304665 | uncharacterized LOC129304665 | -2.1 | 6.09E-05 |
| LOC129310915 | 129310915 | BTB/POZ domain-containing protein At1g50280-like | -2.1 | 1.46E-02 |
| LOC129288105 | 129288105 | methylsterol monooxygenase 1-1-like | -2.1 | 2.30E-05 |
| LOC129288812 | 129288812 | uncharacterized LOC129288812 | -2.1 | 3.26E-05 |
| LOC129299895 | 129299895 | protein trichome birefringence-like 39 | -2.1 | 2.20E-02 |
| LOC129291157 | 129291157 | putative lipid-binding protein At4g00165 | -2.1 | 2.25E-02 |
| LOC129293535 | 129293535 | U-box domain-containing protein 26-like | -2.1 | 4.74E-02 |
| LOC129316772 | 129316772 | uncharacterized LOC129316772 | -2.1 | 1.93E-04 |
| LOC129289381 | 129289381 | ethylene-responsive transcription factor CRF5-like | -2.1 | 3.02E-02 |
| LOC129306267 | 129306267 | protein ILITYHIA | -2.1 | 4.15E-02 |
| LOC129307674 | 129307674 | uncharacterized LOC129307674 | -2.1 | 1.98E-03 |
| LOC129292373 | 129292373 | receptor-like cytosolic serine/threonine-protein kinase RBK2 | -2.1 | 2.37E-02 |
| LOC129284384 | 129284384 | MLP-like protein 28 | -2.1 | 2.03E-02 |
| LOC129305071 | 129305071 | nudix hydrolase 23, chloroplastic | -2.1 | 1.75E-03 |
| LOC129306941 | 129306941 | root phototropism protein 3 | -2.1 | 4.87E-04 |
| LOC129316095 | 129316095 | probable serine/threonine-protein kinase PBL9 | -2.1 | 1.37E-03 |
| LOC129301903 | 129301903 | cysteine-rich receptor-like protein kinase 42 | -2.1 | 2.49E-02 |
| LOC129320935 | 129320935 | heavy metal-associated isoprenylated plant protein 7-like | -2.1 | 2.63E-02 |
| LOC129311769 | 129311769 | E3 ubiquitin-protein ligase Os04g0590900-like | -2.1 | 2.69E-03 |
| LOC129302028 | 129302028 | uncharacterized protein At4g06744 | -2.1 | 2.62E-02 |
| LOC129301048 | 129301048 | non-symbiotic hemoglobin-like | -2.1 | 3.50E-02 |
| LOC129288809 | 129288809 | probable CoA ligase CCL8 | -2.1 | 4.75E-04 |
| LOC129308048 | 129308048 | glutathione S-transferase U9 | -2.1 | 6.15E-04 |
| LOC129295840 | 129295840 | putative disease resistance protein At3g14460 | -2.1 | 4.13E-02 |
| LOC129305241 | 129305241 | ABC transporter G family member 15-like | -2.1 | 6.68E-03 |
| LOC129291823 | 129291823 | uncharacterized LOC129291823 | -2.1 | 4.51E-03 |
| LOC129285324 | 129285324 | probable WRKY transcription factor 27 | -2.1 | 4.58E-02 |
| LOC129315715 | 129315715 | ankyrin repeat-containing protein At5g02620 | -2.1 | 1.23E-04 |
| LOC129317184 | 129317184 | pyruvate decarboxylase 1-like | -2.1 | 1.59E-02 |
| LOC129284989 | 129284989 | U-box domain-containing protein 26-like | -2.1 | 1.25E-04 |
| LOC129322930 | 129322930 | metal transporter Nramp3-like | -2.1 | 5.84E-04 |
| LOC129303335 | 129303335 | linoleate 13S-lipoxygenase 2-1, chloroplastic-like | -2.1 | 3.22E-04 |
| LOC129318971 | 129318971 | probable inorganic phosphate transporter 1-5 | -2.1 | 4.91E-02 |
| LOC129319462 | 129319462 | leucine-rich repeat receptor-like serine/threonine/tyrosine-protei | -2.1 | 3.74E-02 |
| LOC129291321 | 129291321 | NAC domain-containing protein 43-like | -2.1 | 5.35E-06 |
| LOC129315295 | 129315295 | protein PNS1 | -2.1 | 1.43E-03 |
| LOC129311860 | 129311860 | cytochrome P450 CYP82D47-like | -2.1 | 1.42E-02 |
| LOC129303445 | 129303445 | calcineurin B-like protein 4 | -2.1 | 2.52E-02 |
| LOC129291106 | 129291106 | U-box domain-containing protein 33-like | -2.1 | 2.38E-08 |

| | | | | |
|---|---|---|---|---|
| LOC129309329 | 129309329 | adenylate isopentenyltransferase 5, chloroplastic-like | -2.1 | 1.11E-03 |
| LOC129308782 | 129308782 | calcium uptake protein, mitochondrial-like | -2.1 | 1.34E-03 |
| LOC129308791 | 129308791 | endo-1,3;1,4-beta-D-glucanase-like | -2.1 | 1.26E-02 |
| LOC129309087 | 129309087 | AT-rich interactive domain-containing protein 4-like | -2.1 | 5.65E-03 |
| LOC129302536 | 129302536 | uncharacterized LOC129302536 | -2.1 | 7.23E-03 |
| LOC129314964 | 129314964 | gamma-glutamyl hydrolase 2-like | -2.1 | 1.38E-05 |
| LOC129295949 | 129295949 | class 10 plant pathogenesis-related protein 2A-like | -2.1 | 1.47E-02 |
| LOC129309476 | 129309476 | uncharacterized LOC129309476 | -2.1 | 3.33E-02 |
| LOC129311753 | 129311753 | WPP domain-interacting tail-anchored protein 2-like | -2.2 | 1.91E-02 |
| LOC129291620 | 129291620 | FCS-Like Zinc finger 10-like | -2.2 | 4.86E-03 |
| LOC129306572 | 129306572 | uncharacterized LOC129306572 | -2.2 | 3.55E-02 |
| LOC129315606 | 129315606 | S-adenosyl-L-methionine-dependent uroporphyrinogen III methy | -2.2 | 1.45E-02 |
| LOC129312109 | 129312109 | lipid phosphate phosphatase 2-like | -2.2 | 1.43E-02 |
| LOC129291183 | 129291183 | UDP-glycosyltransferase 91C1-like | -2.2 | 5.51E-03 |
| LOC129316015 | 129316015 | RNA polymerase sigma factor sigD, chloroplastic | -2.2 | 9.51E-04 |
| LOC129289468 | 129289468 | dehydrogenase/reductase SDR family member FEY-like | -2.2 | 1.74E-02 |
| LOC129291006 | 129291006 | G-type lectin S-receptor-like serine/threonine-protein kinase At4 | -2.2 | 4.19E-02 |
| LOC129318855 | 129318855 | class V chitinase CHIT5-like | -2.2 | 7.94E-04 |
| LOC129295669 | 129295669 | phosphatidylglycerophosphate phosphatase 1, chloroplastic/mito | -2.2 | 3.86E-02 |
| LOC129322224 | 129322224 | probable aspartyl protease At4g16563 | -2.2 | 1.10E-04 |
| LOC129298284 | 129298284 | protein NRT1/ PTR FAMILY 5.10-like | -2.2 | 1.88E-02 |
| LOC129284893 | 129284893 | G-type lectin S-receptor-like serine/threonine-protein kinase RK | -2.2 | 2.24E-03 |
| LOC129314355 | 129314355 | uncharacterized LOC129314355 | -2.2 | 5.58E-04 |
| LOC129294701 | 129294701 | 14 kDa proline-rich protein DC2.15-like | -2.2 | 1.54E-02 |
| LOC129312031 | 129312031 | uncharacterized LOC129312031 | -2.2 | 2.99E-02 |
| LOC129302880 | 129302880 | filament-like plant protein 4 | -2.2 | 2.69E-03 |
| LOC129297613 | 129297613 | zinc finger CCCH domain-containing protein 18-like | -2.2 | 6.38E-05 |
| LOC129290326 | 129290326 | phospholipase A1 PLIP1, chloroplastic | -2.2 | 1.66E-03 |
| LOC129303823 | 129303823 | auxin-responsive protein IAA31 | -2.2 | 2.38E-02 |
| LOC129319812 | 129319812 | cyprosin-like | -2.2 | 1.09E-04 |
| LOC129316132 | 129316132 | U-box domain-containing protein 1-like | -2.2 | 8.02E-03 |
| LOC129322369 | 129322369 | glutamate--glyoxylate aminotransferase 2 | -2.2 | 7.48E-07 |
| LOC129288752 | 129288752 | probable protein phosphatase 2C 65 | -2.2 | 2.94E-02 |
| LOC129312149 | 129312149 | NAC domain-containing protein 37-like | -2.2 | 4.41E-02 |
| LOC129315941 | 129315941 | BEL1-like homeodomain protein 1 | -2.2 | 4.38E-05 |
| LOC129318522 | 129318522 | glycerophosphodiester phosphodiesterase GDPDL3-like | -2.2 | 1.02E-03 |
| LOC129285857 | 129285857 | uncharacterized LOC129285857 | -2.2 | 7.13E-04 |
| LOC129307789 | 129307789 | ferric reduction oxidase 8, mitochondrial | -2.2 | 1.14E-03 |
| LOC129304384 | 129304384 | beta-xylosidase/alpha-L-arabinofuranosidase 2-like | -2.2 | 2.82E-02 |
| LOC129299678 | 129299678 | F-box/LRR-repeat protein 3-like | -2.2 | 9.77E-04 |
| LOC129321431 | 129321431 | 3-ketoacyl-CoA synthase 6 | -2.2 | 9.75E-06 |
| LOC129290289 | 129290289 | uncharacterized LOC129290289 | -2.2 | 1.30E-02 |
| LOC129322713 | 129322713 | plastidal glycolate/glycerate translocator 1, chloroplastic-like | -2.2 | 6.84E-04 |
| LOC129307349 | 129307349 | polygalacturonase inhibitor-like | -2.2 | 2.64E-02 |
| LOC129314329 | 129314329 | 3-ketoacyl-CoA synthase 12-like | -2.2 | 2.60E-02 |
| LOC129306274 | 129306274 | EID1-like F-box protein 2 | -2.2 | 5.08E-03 |
| LOC129319524 | 129319524 | G-type lectin S-receptor-like serine/threonine-protein kinase LE( | -2.2 | 6.50E-03 |
| LOC129293431 | 129293431 | uncharacterized LOC129293431 | -2.2 | 2.22E-02 |

| | | | | |
|---|---|---|---|---|
| LOC129310477 | 129310477 | homeobox-leucine zipper protein ATHB-13 | -2.2 | 6.88E-08 |
| LOC129319896 | 129319896 | transcription factor TGA4 | -2.2 | 5.22E-06 |
| LOC129319082 | 129319082 | 2-oxoglutarate-dependent dioxygenase 19-like | -2.2 | 4.08E-03 |
| LOC129310997 | 129310997 | glycine-rich domain-containing protein 1-like | -2.2 | 2.46E-03 |
| LOC129305562 | 129305562 | O-fucosyltransferase 27 | -2.2 | 2.10E-02 |
| LOC129287471 | 129287471 | uncharacterized LOC129287471 | -2.2 | 2.27E-02 |
| LOC129306389 | 129306389 | COP1-interacting protein 7 | -2.2 | 1.69E-04 |
| LOC129312956 | 129312956 | uncharacterized LOC129312956 | -2.2 | 1.62E-03 |
| LOC129294346 | 129294346 | thioredoxin M-type, chloroplastic-like | -2.2 | 5.22E-06 |
| LOC129297598 | 129297598 | receptor-like protein 7 | -2.2 | 4.30E-04 |
| LOC129302174 | 129302174 | subtilisin-like protease SBT3 | -2.2 | 7.12E-03 |
| LOC129305231 | 129305231 | cation/H(+) antiporter 20-like | -2.2 | 2.87E-02 |
| LOC129302071 | 129302071 | 2-oxoglutarate-dependent dioxygenase 19-like | -2.2 | 1.10E-03 |
| LOC129320773 | 129320773 | uncharacterized LOC129320773 | -2.3 | 5.40E-04 |
| LOC129309218 | 129309218 | protein CHUP1, chloroplastic | -2.3 | 1.45E-10 |
| LOC129288127 | 129288127 | probable polygalacturonase | -2.3 | 1.89E-03 |
| LOC129314389 | 129314389 | putative disease resistance protein At3g14460 | -2.3 | 4.28E-02 |
| LOC129291558 | 129291558 | MAPK kinase substrate protein At1g80180-like | -2.3 | 6.87E-04 |
| LOC129322963 | 129322963 | probable trehalose-phosphate phosphatase F | -2.3 | 4.82E-03 |
| LOC129306749 | 129306749 | uncharacterized LOC129306749 | -2.3 | 1.03E-03 |
| LOC129302184 | 129302184 | metal-nicotianamine transporter YSL3-like | -2.3 | 9.69E-03 |
| LOC129285945 | 129285945 | subtilisin-like protease SBT1.6 | -2.3 | 8.29E-03 |
| LOC129296945 | 129296945 | uncharacterized LOC129296945 | -2.3 | 4.70E-02 |
| LOC129293833 | 129293833 | probable membrane-associated kinase regulator 6 | -2.3 | 2.86E-03 |
| LOC129293419 | 129293419 | probable carboxylesterase 12 | -2.3 | 4.93E-06 |
| LOC129321592 | 129321592 | uncharacterized LOC129321592 | -2.3 | 4.37E-02 |
| LOC129319826 | 129319826 | zinc finger protein ZAT9-like | -2.3 | 1.42E-02 |
| LOC129296955 | 129296955 | probable ribose-5-phosphate isomerase 4, chloroplastic | -2.3 | 2.37E-04 |
| LOC129321841 | 129321841 | uncharacterized LOC129321841 | -2.3 | 3.55E-04 |
| LOC129294223 | 129294223 | sugar transport protein 5-like | -2.3 | 3.76E-05 |
| LOC129305985 | 129305985 | uncharacterized LOC129305985 | -2.3 | 2.30E-07 |
| LOC129294413 | 129294413 | RNA polymerase sigma factor sigC | -2.3 | 5.15E-06 |
| LOC129323217 | 129323217 | cinnamoyl-CoA reductase-like SNL6 | -2.3 | 2.40E-04 |
| LOC129305114 | 129305114 | transcription factor MYB8-like | -2.3 | 4.09E-05 |
| LOC129293711 | 129293711 | cytochrome P450 CYP736A12-like | -2.3 | 1.87E-02 |
| LOC129306584 | 129306584 | uncharacterized LOC129306584 | -2.3 | 1.18E-02 |
| LOC129319789 | 129319789 | receptor-like protein kinase FERONIA | -2.3 | 1.25E-02 |
| LOC129315255 | 129315255 | basic leucine zipper 61-like | -2.3 | 1.64E-02 |
| LOC129307238 | 129307238 | uncharacterized LOC129307238 | -2.3 | 3.69E-02 |
| LOC129307825 | 129307825 | probable polyamine transporter At3g13620 | -2.3 | 1.37E-03 |
| LOC129295017 | 129295017 | cytosolic sulfotransferase 15-like | -2.3 | 7.24E-03 |
| LOC129309088 | 129309088 | probable mannitol dehydrogenase | -2.3 | 1.07E-04 |
| LOC129322275 | 129322275 | uncharacterized LOC129322275 | -2.3 | 5.33E-04 |
| LOC129302964 | 129302964 | anthocyanidin 3-O-glucosyltransferase 5-like | -2.3 | 1.42E-02 |
| LOC129322809 | 129322809 | long-chain-alcohol oxidase FAO4A-like | -2.3 | 1.13E-06 |
| LOC129302206 | 129302206 | putative GATA transcription factor 22 | -2.3 | 2.71E-02 |
| LOC129307689 | 129307689 | 3-ketoacyl-CoA synthase 11-like | -2.3 | 1.77E-04 |
| LOC129297653 | 129297653 | hevamine-A-like | -2.3 | 1.66E-02 |

| | | | | |
|---|---|---|---|---|
| LOC129318941 | 129318941 | mannan endo-1,4-beta-mannosidase 7-like | -2.3 | 6.61E-06 |
| LOC129315924 | 129315924 | WAT1-related protein At5g07050-like | -2.3 | 4.66E-05 |
| LOC129322773 | 129322773 | uncharacterized LOC129322773 | -2.3 | 1.30E-02 |
| LOC129309501 | 129309501 | uncharacterized LOC129309501 | -2.3 | 1.70E-06 |
| LOC129288631 | 129288631 | subtilisin-like protease SBT1.1 | -2.3 | 3.04E-03 |
| LOC129320542 | 129320542 | serine carboxypeptidase-like 26 | -2.3 | 2.24E-03 |
| LOC129311526 | 129311526 | uncharacterized LOC129311526 | -2.3 | 1.62E-02 |
| LOC129284414 | 129284414 | glutamate receptor 3.6-like | -2.3 | 8.47E-03 |
| LOC129307051 | 129307051 | uncharacterized LOC129307051 | -2.3 | 2.78E-07 |
| LOC129304856 | 129304856 | protein NRT1/ PTR FAMILY 5.8-like | -2.3 | 6.19E-08 |
| LOC129289529 | 129289529 | uncharacterized LOC129289529 | -2.3 | 9.70E-04 |
| LOC129288370 | 129288370 | disease resistance protein At4g27190-like | -2.3 | 3.06E-03 |
| LOC129305151 | 129305151 | MLO-like protein 2 | -2.3 | 2.19E-04 |
| LOC129311564 | 129311564 | hevamine-A-like | -2.3 | 2.53E-02 |
| LOC129286736 | 129286736 | thiamine thiazole synthase 2, chloroplastic-like | -2.3 | 1.31E-05 |
| LOC129300849 | 129300849 | uncharacterized LOC129300849 | -2.3 | 3.86E-03 |
| LOC129291383 | 129291383 | protein indeterminate-domain 7 | -2.4 | 8.99E-04 |
| LOC129286942 | 129286942 | uncharacterized protein At4g06744-like | -2.4 | 1.24E-04 |
| LOC129321819 | 129321819 | polygalacturonase At1g48100-like | -2.4 | 2.99E-02 |
| LOC129304171 | 129304171 | uncharacterized LOC129304171 | -2.4 | 2.81E-03 |
| LOC129285760 | 129285760 | K(+) efflux antiporter 3, chloroplastic-like | -2.4 | 5.65E-07 |
| LOC129301559 | 129301559 | cysteine-rich receptor-like protein kinase 43 | -2.4 | 3.68E-02 |
| LOC129298455 | 129298455 | cytochrome P450 76C4-like | -2.4 | 3.45E-02 |
| LOC129307934 | 129307934 | probable pectate lyase 18 | -2.4 | 5.97E-03 |
| LOC129316235 | 129316235 | serine carboxypeptidase-like 51 | -2.4 | 1.24E-04 |
| LOC129292599 | 129292599 | uncharacterized oxidoreductase At4g09670-like | -2.4 | 2.35E-02 |
| LOC129320321 | 129320321 | repetitive proline-rich cell wall protein 2-like | -2.4 | 1.61E-02 |
| LOC129287217 | 129287217 | uncharacterized LOC129287217 | -2.4 | 1.29E-02 |
| LOC129317209 | 129317209 | BTB/POZ domain-containing protein NPY1 | -2.4 | 1.83E-05 |
| LOC129294808 | 129294808 | ethylene-responsive transcription factor SHINE 3-like | -2.4 | 1.18E-04 |
| LOC129288035 | 129288035 | transcription factor MYB60-like | -2.4 | 9.73E-07 |
| LOC129306403 | 129306403 | berberine bridge enzyme-like 26 | -2.4 | 3.23E-04 |
| LOC129287836 | 129287836 | laccase-17-like | -2.4 | 1.03E-02 |
| LOC129288941 | 129288941 | RING-H2 finger protein ATL2-like | -2.4 | 2.14E-02 |
| LOC129315792 | 129315792 | expansin-A4-like | -2.4 | 8.13E-03 |
| LOC129302416 | 129302416 | U-box domain-containing protein 44-like | -2.4 | 1.06E-04 |
| LOC129288957 | 129288957 | hydroquinone glucosyltransferase-like | -2.4 | 5.13E-03 |
| LOC129312208 | 129312208 | uncharacterized LOC129312208 | -2.4 | 1.67E-04 |
| LOC129318632 | 129318632 | E3 ubiquitin-protein ligase At1g12760-like | -2.4 | 4.88E-04 |
| LOC129292711 | 129292711 | probable carboxylesterase 1 | -2.4 | 3.14E-03 |
| LOC129288408 | 129288408 | auxin transporter-like protein 2 | -2.4 | 1.05E-06 |
| LOC129313522 | 129313522 | peroxidase P7-like | -2.4 | 2.29E-06 |
| LOC129312113 | 129312113 | uncharacterized LOC129312113 | -2.4 | 1.10E-03 |
| LOC129318510 | 129318510 | putative wall-associated receptor kinase-like 16 | -2.4 | 1.81E-03 |
| LOC129323220 | 129323220 | uncharacterized acetyltransferase At3g50280-like | -2.4 | 2.34E-02 |
| LOC129303565 | 129303565 | cation/calcium exchanger 1-like | -2.4 | 8.60E-04 |
| LOC129314625 | 129314625 | probable glucuronoxylan glucuronosyltransferase IRX7 | -2.4 | 1.46E-04 |
| LOC129293551 | 129293551 | photosynthetic NDH subunit of subcomplex B 2, chloroplastic | -2.4 | 1.46E-03 |

| | | | | |
|---|---|---|---|---|
| LOC129316787 | 129316787 | uncharacterized LOC129316787 | -2.4 | 4.82E-02 |
| LOC129316852 | 129316852 | dirigent protein 22-like | -2.4 | 3.58E-04 |
| LOC129320100 | 129320100 | uncharacterized LOC129320100 | -2.4 | 2.32E-04 |
| LOC129307038 | 129307038 | probable xyloglucan endotransglucosylase/hydrolase protein 32 | -2.4 | 3.80E-03 |
| LOC129306755 | 129306755 | uncharacterized LOC129306755 | -2.4 | 7.55E-05 |
| LOC129293306 | 129293306 | 2-hydroxyisoflavanone dehydratase-like | -2.4 | 5.63E-07 |
| LOC129303808 | 129303808 | root phototropism protein 2 | -2.5 | 2.74E-10 |
| LOC129284646 | 129284646 | RING-H2 finger protein ATL52-like | -2.5 | 2.28E-03 |
| LOC129296710 | 129296710 | hypothetical protein At1g04090-like | -2.5 | 7.84E-04 |
| LOC129296320 | 129296320 | zinc finger protein VAR3, chloroplastic-like | -2.5 | 1.62E-03 |
| LOC129297192 | 129297192 | uncharacterized LOC129297192 | -2.5 | 2.16E-02 |
| LOC129305911 | 129305911 | pectinesterase inhibitor 6-like | -2.5 | 2.67E-02 |
| LOC129294688 | 129294688 | uncharacterized LOC129294688 | -2.5 | 7.59E-03 |
| LOC129306296 | 129306296 | probable aspartic protease At2g35615 | -2.5 | 1.43E-03 |
| LOC129306960 | 129306960 | GDSL esterase/lipase At5g14450-like | -2.5 | 1.55E-04 |
| LOC129320216 | 129320216 | zinc finger protein BRUTUS-like At1g18910 | -2.5 | 1.81E-02 |
| LOC129305175 | 129305175 | 7-deoxyloganetin glucosyltransferase-like | -2.5 | 3.15E-02 |
| LOC129323037 | 129323037 | xylan glycosyltransferase MUCI21-like | -2.5 | 4.18E-02 |
| LOC129315316 | 129315316 | putative disease resistance RPP13-like protein 1 | -2.5 | 3.66E-03 |
| LOC129313564 | 129313564 | protein SOSEKI 3 | -2.5 | 3.49E-07 |
| LOC129290615 | 129290615 | putative receptor-like protein kinase At3g47110 | -2.5 | 1.89E-07 |
| LOC129299111 | 129299111 | pentatricopeptide repeat-containing protein At5g10690 | -2.5 | 1.30E-03 |
| LOC129287548 | 129287548 | aspartyl protease family protein At5g10770 | -2.5 | 6.33E-03 |
| LOC129312458 | 129312458 | cytosolic sulfotransferase 15-like | -2.5 | 1.99E-02 |
| LOC129306840 | 129306840 | potassium channel KAT1-like | -2.5 | 5.69E-06 |
| LOC129289454 | 129289454 | ethylene-responsive transcription factor WIN1 | -2.5 | 3.15E-02 |
| LOC129302470 | 129302470 | serine/threonine-protein kinase STY17-like | -2.5 | 1.44E-02 |
| LOC129322819 | 129322819 | mitochondrial uncoupling protein 5-like | -2.5 | 6.06E-04 |
| LOC129292701 | 129292701 | silicon efflux transporter LSI2-like | -2.5 | 8.78E-05 |
| LOC129308767 | 129308767 | beta-D-xylosidase 1 | -2.5 | 5.34E-05 |
| LOC129295970 | 129295970 | DNA mismatch repair protein MSH5-like | -2.5 | 1.24E-02 |
| LOC129307916 | 129307916 | fatty acid amide hydrolase-like | -2.5 | 1.50E-06 |
| LOC129305077 | 129305077 | chloride channel protein CLC-b | -2.5 | 4.89E-06 |
| LOC129323031 | 129323031 | probable galacturonosyltransferase-like 3 | -2.5 | 1.01E-02 |
| LOC129296039 | 129296039 | uncharacterized LOC129296039 | -2.6 | 2.81E-03 |
| LOC129300913 | 129300913 | uncharacterized LOC129300913 | -2.6 | 2.25E-05 |
| LOC129303672 | 129303672 | anthocyanidin 3-O-glucosyltransferase 6-like | -2.6 | 1.66E-04 |
| LOC129302851 | 129302851 | uncharacterized LOC129302851 | -2.6 | 1.34E-02 |
| LOC129305702 | 129305702 | long-chain-alcohol oxidase FAO2 | -2.6 | 6.46E-04 |
| LOC129288049 | 129288049 | probable serine/threonine-protein kinase SIS8 | -2.6 | 6.18E-08 |
| LOC129293801 | 129293801 | protein NLP6-like | -2.6 | 8.07E-08 |
| LOC129322153 | 129322153 | beta-amylase 3, chloroplastic-like | -2.6 | 1.87E-13 |
| LOC129291741 | 129291741 | copper transporter 2-like | -2.6 | 8.11E-03 |
| LOC129288653 | 129288653 | NAC domain-containing protein 43 | -2.6 | 1.80E-03 |
| LOC129304877 | 129304877 | 7-deoxyloganetin glucosyltransferase-like | -2.6 | 3.73E-03 |
| LOC129315031 | 129315031 | G-type lectin S-receptor-like serine/threonine-protein kinase At4 | -2.6 | 2.58E-02 |
| LOC129284598 | 129284598 | cytochrome P450 86A22 | -2.6 | 4.26E-04 |
| LOC129312923 | 129312923 | uncharacterized LOC129312923 | -2.6 | 1.25E-02 |

| | | | | |
|---|---|---|---|---|
| LOC129293519 | 129293519 | inositol oxygenase 4-like | -2.6 | 2.49E-03 |
| LOC129305111 | 129305111 | aspartyl protease family protein At5g10770-like | -2.6 | 2.81E-04 |
| LOC129295672 | 129295672 | 2-oxoglutarate-dependent dioxygenase 19-like | -2.6 | 4.72E-03 |
| LOC129303788 | 129303788 | WUSCHEL-related homeobox 4-like | -2.6 | 4.14E-03 |
| LOC129318482 | 129318482 | subtilisin-like protease SBT1.7 | -2.6 | 4.02E-05 |
| LOC129311190 | 129311190 | homeobox protein BEL1 homolog | -2.6 | 4.18E-08 |
| LOC129311780 | 129311780 | zinc finger CCCH domain-containing protein 6-like | -2.6 | 5.13E-04 |
| LOC129306916 | 129306916 | auxin-responsive protein SAUR32-like | -2.6 | 7.93E-04 |
| LOC129320735 | 129320735 | FCS-Like Zinc finger 8 | -2.6 | 2.43E-04 |
| LOC129295308 | 129295308 | carboxylesterase 1-like | -2.6 | 1.01E-03 |
| LOC129287828 | 129287828 | aldehyde oxidase GLOX | -2.6 | 2.31E-03 |
| LOC129304481 | 129304481 | calcium-binding protein KRP1-like | -2.6 | 3.87E-02 |
| LOC129314285 | 129314285 | 21 kDa protein | -2.6 | 4.08E-03 |
| LOC129285671 | 129285671 | epimerase family protein SDR39U1 homolog, chloroplastic | -2.6 | 2.39E-10 |
| LOC129300001 | 129300001 | cytochrome P450 77A3-like | -2.6 | 7.32E-03 |
| LOC129309492 | 129309492 | cytochrome P450 71AP13-like | -2.7 | 1.44E-02 |
| LOC129292891 | 129292891 | non-symbiotic hemoglobin-like | -2.7 | 3.75E-02 |
| LOC129298141 | 129298141 | chlorophyllase-2-like | -2.7 | 4.52E-03 |
| LOC129296928 | 129296928 | chaperone protein dnaJ C76, chloroplastic-like | -2.7 | 5.46E-06 |
| LOC129304987 | 129304987 | type I inositol polyphosphate 5-phosphatase 4-like | -2.7 | 5.31E-03 |
| LOC129295134 | 129295134 | proline dehydrogenase 2, mitochondrial-like | -2.7 | 1.76E-04 |
| LOC129301994 | 129301994 | laccase-5-like | -2.7 | 1.84E-02 |
| LOC129287957 | 129287957 | uncharacterized LOC129287957 | -2.7 | 9.65E-03 |
| LOC129322694 | 129322694 | uncharacterized acetyltransferase At3g50280-like | -2.7 | 6.65E-07 |
| LOC129291103 | 129291103 | 3-ketoacyl-CoA synthase 1-like | -2.7 | 1.22E-07 |
| LOC129304242 | 129304242 | transcription factor PIF4-like | -2.7 | 1.42E-09 |
| LOC129285319 | 129285319 | putative F-box protein At3g23960 | -2.7 | 3.92E-03 |
| LOC129301918 | 129301918 | cucumisin-like | -2.7 | 7.22E-05 |
| LOC129293555 | 129293555 | receptor-like protein kinase FERONIA | -2.7 | 2.25E-07 |
| LOC129303883 | 129303883 | zinc transporter 8-like | -2.7 | 1.30E-02 |
| LOC129288037 | 129288037 | uncharacterized LOC129288037 | -2.7 | 2.47E-02 |
| LOC129311929 | 129311929 | rhamnogalacturonan I rhamnosyltransferase 1-like | -2.7 | 1.97E-02 |
| LOC129300216 | 129300216 | phenylacetaldehyde oxime monooxygenase CYP71AN24-like | -2.7 | 5.74E-15 |
| LOC129284440 | 129284440 | uncharacterized LOC129284440 | -2.7 | 1.37E-03 |
| LOC129286752 | 129286752 | uncharacterized LOC129286752 | -2.7 | 3.95E-03 |
| LOC129320864 | 129320864 | protein NDL1-like | -2.7 | 7.08E-04 |
| LOC129289162 | 129289162 | MDIS1-interacting receptor like kinase 2-like | -2.7 | 2.89E-02 |
| LOC129303406 | 129303406 | glycerol-3-phosphate acyltransferase 5-like | -2.7 | 6.87E-03 |
| LOC129318737 | 129318737 | aspartyl protease family protein At5g10770-like | -2.7 | 3.58E-02 |
| LOC129296490 | 129296490 | probable inactive purple acid phosphatase 29 | -2.7 | 6.21E-06 |
| LOC129309278 | 129309278 | uncharacterized LOC129309278 | -2.7 | 1.86E-11 |
| LOC129288690 | 129288690 | uncharacterized LOC129288690 | -2.7 | 1.25E-02 |
| LOC129322192 | 129322192 | putative pentatricopeptide repeat-containing protein At3g01580 | -2.7 | 6.29E-03 |
| LOC129314446 | 129314446 | kiwellin-like | -2.7 | 8.24E-06 |
| LOC129310945 | 129310945 | protein argonaute 7 | -2.7 | 5.69E-05 |
| LOC129293461 | 129293461 | polyol transporter 5-like | -2.7 | 1.64E-05 |
| LOC129321922 | 129321922 | mechanosensitive ion channel protein 10-like | -2.7 | 2.49E-03 |
| LOC129320396 | 129320396 | leucine-rich repeat receptor-like serine/threonine-protein kinase | -2.8 | 7.08E-05 |

| | | | | |
|---|---|---|---|---|
| LOC129304369 | 129304369 | cation/calcium exchanger 1-like | -2.8 | 1.52E-05 |
| LOC129318599 | 129318599 | aspartyl protease family protein At5g10770-like | -2.8 | 1.13E-06 |
| LOC129287582 | 129287582 | uncharacterized LOC129287582 | -2.8 | 2.85E-02 |
| LOC129317971 | 129317971 | aspartyl protease family protein At5g10770-like | -2.8 | 1.88E-03 |
| LOC129320988 | 129320988 | uncharacterized LOC129320988 | -2.8 | 1.76E-05 |
| LOC129294652 | 129294652 | leucine-rich repeat receptor-like serine/threonine-protein kinase | -2.8 | 2.08E-03 |
| LOC129309973 | 129309973 | transcription factor IBH1-like 1 | -2.8 | 7.77E-04 |
| LOC129290692 | 129290692 | cytokinin hydroxylase | -2.8 | 2.90E-02 |
| LOC129321773 | 129321773 | uncharacterized LOC129321773 | -2.8 | 3.43E-03 |
| LOC129304371 | 129304371 | protein PNS1-like | -2.8 | 3.21E-03 |
| LOC129322932 | 129322932 | serine carboxypeptidase-like 50 | -2.8 | 3.39E-04 |
| LOC129306690 | 129306690 | alkane hydroxylase MAH1-like | -2.8 | 1.93E-06 |
| LOC129315366 | 129315366 | UPF0481 protein At3g47200-like | -2.8 | 4.77E-04 |
| LOC129303440 | 129303440 | LRR receptor-like serine/threonine-protein kinase RGI2 | -2.8 | 2.28E-03 |
| LOC129284394 | 129284394 | uncharacterized acetyltransferase At3g50280-like | -2.8 | 5.08E-03 |
| LOC129318385 | 129318385 | glucan endo-1,3-beta-glucosidase, acidic-like | -2.8 | 7.51E-03 |
| LOC129317564 | 129317564 | acyl-lipid (9-3)-desaturase-like | -2.8 | 2.43E-03 |
| LOC129293639 | 129293639 | transcription factor HBI1-like | -2.8 | 1.69E-05 |
| LOC129290499 | 129290499 | probable disease resistance protein At5g66900 | -2.8 | 4.27E-03 |
| LOC129293495 | 129293495 | probable caffeine synthase MTL2 | -2.8 | 2.57E-04 |
| LOC129288903 | 129288903 | uncharacterized LOC129288903 | -2.8 | 6.75E-04 |
| LOC129288120 | 129288120 | beta-galactosidase 1-like | -2.8 | 6.51E-08 |
| LOC129308185 | 129308185 | uncharacterized LOC129308185 | -2.8 | 9.11E-07 |
| LOC129298269 | 129298269 | uncharacterized protein At4g06744-like | -2.9 | 1.45E-02 |
| LOC129318922 | 129318922 | receptor like protein 21-like | -2.9 | 4.38E-03 |
| LOC129292415 | 129292415 | disease resistance protein RPV1-like | -2.9 | 3.04E-02 |
| LOC129285338 | 129285338 | mannose/glucose-specific lectin-like | -2.9 | 6.06E-03 |
| LOC129322252 | 129322252 | auxin efflux carrier component 3-like | -2.9 | 1.16E-05 |
| LOC129313766 | 129313766 | transcription factor TCP4-like | -2.9 | 2.72E-06 |
| LOC129307816 | 129307816 | xyloglucan endotransglucosylase/hydrolase 2 | -2.9 | 3.38E-03 |
| LOC129292053 | 129292053 | acetate--CoA ligase CCL3-like | -2.9 | 3.87E-19 |
| LOC129317186 | 129317186 | aspartyl protease family protein At5g10770-like | -2.9 | 2.62E-05 |
| LOC129292123 | 129292123 | S-type anion channel SLAH2-like | -2.9 | 1.48E-03 |
| LOC129298790 | 129298790 | coniferyl alcohol acyltransferase-like | -2.9 | 1.06E-02 |
| LOC129300381 | 129300381 | aspartyl protease family protein At5g10770-like | -2.9 | 1.74E-08 |
| LOC129322798 | 129322798 | uncharacterized LOC129322798 | -2.9 | 2.10E-05 |
| LOC129303652 | 129303652 | trihelix transcription factor DF1-like | -2.9 | 5.11E-08 |
| LOC129311774 | 129311774 | uncharacterized LOC129311774 | -2.9 | 1.77E-04 |
| LOC129322973 | 129322973 | protein SMALL AUXIN UP-REGULATED RNA 12 | -2.9 | 4.24E-03 |
| LOC129311672 | 129311672 | probable leucine-rich repeat receptor-like protein kinase At5g63 | -2.9 | 1.05E-03 |
| LOC129310073 | 129310073 | uncharacterized protein At4g14100-like | -2.9 | 8.62E-04 |
| LOC129320518 | 129320518 | BTB/POZ and TAZ domain-containing protein 1-like | -2.9 | 3.74E-08 |
| LOC129306719 | 129306719 | polygalacturonase QRT3 | -2.9 | 2.10E-03 |
| LOC129284395 | 129284395 | 1-aminocyclopropane-1-carboxylate synthase 6 | -2.9 | 4.99E-08 |
| LOC129306131 | 129306131 | GDSL esterase/lipase At1g31550-like | -2.9 | 1.04E-03 |
| LOC129320618 | 129320618 | amine oxidase [copper-containing] zeta, peroxisomal-like | -2.9 | 5.40E-03 |
| LOC129291486 | 129291486 | sulfite exporter TauE/SafE family protein 3-like | -2.9 | 7.48E-03 |
| LOC129287531 | 129287531 | factor Xa inhibitor BuXI-like | -2.9 | 2.87E-03 |

| | | | | |
|---|---|---|---|---|
| LOC129319906 | 129319906 | ras-related protein RABE1c-like | -2.9 | 2.39E-04 |
| LOC129294625 | 129294625 | agamous-like MADS-box protein MADS9 | -2.9 | 4.87E-03 |
| LOC129302211 | 129302211 | topless-related protein 4 | -3.0 | 2.00E-05 |
| LOC129306821 | 129306821 | uncharacterized LOC129306821 | -3.0 | 2.24E-02 |
| LOC129315958 | 129315958 | phospholipase A1-IIdelta-like | -3.0 | 4.34E-05 |
| LOC129317844 | 129317844 | potassium transporter 1-like | -3.0 | 5.65E-07 |
| LOC129301292 | 129301292 | probable carotenoid cleavage dioxygenase 4, chloroplastic | -3.0 | 3.87E-03 |
| LOC129286989 | 129286989 | uncharacterized LOC129286989 | -3.0 | 2.08E-11 |
| LOC129286982 | 129286982 | uncharacterized LOC129286982 | -3.0 | 1.27E-09 |
| LOC129301894 | 129301894 | uncharacterized LOC129301894 | -3.0 | 3.61E-02 |
| LOC129316349 | 129316349 | putative disease resistance protein At3g14460 | -3.0 | 2.80E-05 |
| LOC129310673 | 129310673 | rust resistance kinase Lr10-like | -3.0 | 4.11E-02 |
| LOC129321590 | 129321590 | uncharacterized LOC129321590 | -3.0 | 9.86E-04 |
| LOC129294253 | 129294253 | G-type lectin S-receptor-like serine/threonine-protein kinase SD[illegible] | -3.0 | 4.82E-04 |
| LOC129285926 | 129285926 | oligopeptide transporter 3 | -3.0 | 4.16E-18 |
| LOC129287312 | 129287312 | leucine-rich repeat receptor protein kinase HPCA1-like | -3.0 | 3.48E-04 |
| LOC129292072 | 129292072 | 3-ketoacyl-CoA synthase 19 | -3.0 | 5.39E-14 |
| LOC129301113 | 129301113 | aspartyl protease AED3-like | -3.0 | 6.41E-03 |
| LOC129320533 | 129320533 | U-box domain-containing protein 26-like | -3.0 | 9.70E-03 |
| LOC129284422 | 129284422 | uncharacterized LOC129284422 | -3.0 | 3.09E-05 |
| LOC129298851 | 129298851 | probable xyloglucan endotransglucosylase/hydrolase protein 8 | -3.0 | 3.49E-02 |
| LOC129284455 | 129284455 | protein PYRICULARIA ORYZAE RESISTANCE 21-like | -3.0 | 2.41E-02 |
| LOC129284674 | 129284674 | transcription repressor OFP15-like | -3.1 | 2.45E-02 |
| LOC129320632 | 129320632 | uncharacterized LOC129320632 | -3.1 | 3.42E-03 |
| LOC129290656 | 129290656 | dynein light chain 1, cytoplasmic-like | -3.1 | 1.14E-03 |
| LOC129310719 | 129310719 | probable carotenoid cleavage dioxygenase 4, chloroplastic | -3.1 | 1.91E-08 |
| LOC129315171 | 129315171 | IRK-interacting protein-like | -3.1 | 2.03E-05 |
| LOC129316329 | 129316329 | vascular-related unknown protein 1-like | -3.1 | 9.10E-05 |
| LOC129309536 | 129309536 | uncharacterized LOC129309536 | -3.1 | 3.78E-02 |
| LOC129292502 | 129292502 | cationic amino acid transporter 7, chloroplastic | -3.1 | 8.29E-04 |
| LOC129306227 | 129306227 | uncharacterized LOC129306227 | -3.1 | 1.01E-04 |
| LOC129307291 | 129307291 | boron transporter 1-like | -3.1 | 8.34E-10 |
| LOC129306568 | 129306568 | RNA demethylase ALKBH10B-like | -3.1 | 2.53E-16 |
| LOC129309406 | 129309406 | RING-H2 finger protein ATL13-like | -3.1 | 9.63E-03 |
| LOC129291563 | 129291563 | GDSL esterase/lipase APG-like | -3.1 | 4.98E-09 |
| LOC129311798 | 129311798 | probable methyltransferase TCM_000336 | -3.1 | 2.01E-03 |
| LOC129313756 | 129313756 | probable serine/threonine-protein kinase PBL7 | -3.1 | 4.39E-03 |
| LOC129322923 | 129322923 | sulfite exporter TauE/SafE family protein 3-like | -3.1 | 2.03E-03 |
| LOC129298799 | 129298799 | superoxide dismutase [Cu-Zn] 2-like | -3.1 | 1.30E-02 |
| LOC129286795 | 129286795 | protein EXORDIUM-like 2 | -3.1 | 1.35E-03 |
| LOC129313524 | 129313524 | E3 ubiquitin-protein ligase RZFP34-like | -3.1 | 3.11E-04 |
| LOC129312709 | 129312709 | transcription factor MYB59 | -3.2 | 4.26E-04 |
| LOC129312114 | 129312114 | putative invertase inhibitor | -3.2 | 1.96E-04 |
| LOC129315185 | 129315185 | disease resistance protein RUN1-like | -3.2 | 4.11E-02 |
| LOC129299396 | 129299396 | squalene monooxygenase SE1-like | -3.2 | 6.06E-05 |
| LOC129298560 | 129298560 | peroxidase P7-like | -3.2 | 6.54E-07 |
| LOC129314282 | 129314282 | probable serine/threonine-protein kinase PBL10 | -3.2 | 4.00E-04 |
| LOC129302195 | 129302195 | coniferyl alcohol acyltransferase-like | -3.2 | 7.30E-06 |

| | | | | |
|---|---|---|---|---|
| LOC129308003 | 129308003 | glutathione hydrolase 3 | -3.2 | 3.92E-10 |
| LOC129296228 | 129296228 | acidic endochitinase SE2-like | -3.2 | 1.20E-02 |
| LOC129284446 | 129284446 | uncharacterized LOC129284446 | -3.2 | 2.95E-03 |
| LOC129301916 | 129301916 | cucumisin-like | -3.2 | 6.60E-14 |
| LOC129321692 | 129321692 | anthocyanidin 3-O-glucosyltransferase 5-like | -3.2 | 2.89E-04 |
| LOC129290153 | 129290153 | uncharacterized LOC129290153 | -3.2 | 5.07E-03 |
| LOC129291052 | 129291052 | aspartic proteinase PCS1 | -3.2 | 1.18E-09 |
| LOC129302242 | 129302242 | pentatricopeptide repeat-containing protein At4g33990-like | -3.2 | 4.40E-02 |
| LOC129310046 | 129310046 | 4-coumarate--CoA ligase CCL1-like | -3.2 | 2.06E-06 |
| LOC129314577 | 129314577 | cytosolic sulfotransferase 15-like | -3.2 | 2.30E-02 |
| LOC129322230 | 129322230 | adenine phosphoribosyltransferase 3-like | -3.3 | 1.39E-06 |
| LOC129296256 | 129296256 | class 10 plant pathogenesis-related protein 2A-like | -3.3 | 5.13E-09 |
| LOC129308800 | 129308800 | LRR receptor-like serine/threonine-protein kinase ERL1 | -3.3 | 1.01E-04 |
| LOC129306649 | 129306649 | UDP-glycosyltransferase 83A1-like | -3.3 | 1.06E-02 |
| LOC129317146 | 129317146 | 3-ketoacyl-CoA synthase 10 | -3.3 | 1.54E-07 |
| LOC129314946 | 129314946 | serine carboxypeptidase-like | -3.3 | 1.22E-05 |
| LOC129292686 | 129292686 | protein trichome birefringence-like 19 | -3.3 | 1.35E-18 |
| LOC129292677 | 129292677 | receptor-like protein EIX1 | -3.3 | 8.92E-03 |
| LOC129294757 | 129294757 | protein MKS1-like | -3.3 | 3.33E-02 |
| LOC129303249 | 129303249 | protein WALLS ARE THIN 1-like | -3.3 | 1.26E-07 |
| LOC129300167 | 129300167 | uncharacterized LOC129300167 | -3.3 | 1.66E-08 |
| LOC129321220 | 129321220 | subtilisin-like protease SBT1.7 | -3.3 | 1.88E-10 |
| LOC129308368 | 129308368 | probable glycosyltransferase At5g03795 | -3.3 | 2.06E-07 |
| LOC129288892 | 129288892 | F-box/LRR-repeat protein 3-like | -3.3 | 1.92E-03 |
| LOC129286842 | 129286842 | ervatamin-B-like | -3.3 | 7.23E-04 |
| LOC129321845 | 129321845 | serine carboxypeptidase-like 34 | -3.3 | 1.30E-02 |
| LOC129316079 | 129316079 | disease resistance protein RPV1-like | -3.4 | 1.12E-03 |
| LOC129284463 | 129284463 | uncharacterized LOC129284463 | -3.4 | 3.31E-04 |
| LOC129292729 | 129292729 | S-adenosylmethionine decarboxylase proenzyme 4-like | -3.4 | 1.56E-02 |
| LOC129315460 | 129315460 | ABC transporter B family member 15-like | -3.4 | 1.43E-10 |
| LOC129304030 | 129304030 | protein SOB FIVE-LIKE 6-like | -3.4 | 7.04E-03 |
| LOC129304525 | 129304525 | uncharacterized LOC129304525 | -3.4 | 1.45E-02 |
| LOC129318363 | 129318363 | probable amino acid permease 7 | -3.4 | 3.81E-02 |
| LOC129306247 | 129306247 | protein CANDIDATE G-PROTEIN COUPLED RECEPTOR 7- | -3.4 | 4.25E-02 |
| LOC129311767 | 129311767 | ABC transporter B family member 2-like | -3.4 | 3.18E-09 |
| LOC129293446 | 129293446 | gibberellin 20 oxidase 2-like | -3.4 | 2.76E-04 |
| LOC129306652 | 129306652 | fructose-1,6-bisphosphatase, cytosolic-like | -3.4 | 1.83E-02 |
| LOC129285143 | 129285143 | UDP-glycosyltransferase 88A1-like | -3.4 | 7.15E-03 |
| LOC129295198 | 129295198 | uncharacterized LOC129295198 | -3.4 | 1.11E-06 |
| LOC129321413 | 129321413 | protein DETOXIFICATION 27-like | -3.4 | 1.83E-07 |
| LOC129321247 | 129321247 | uncharacterized LOC129321247 | -3.4 | 1.06E-02 |
| LOC129298582 | 129298582 | gibberellin 20 oxidase 1-D-like | -3.4 | 2.01E-06 |
| LOC129321450 | 129321450 | receptor-like protein kinase FERONIA | -3.4 | 9.48E-08 |
| LOC129310667 | 129310667 | protein HOTHEAD-like | -3.4 | 9.74E-08 |
| LOC129311944 | 129311944 | glycosyltransferase BC10-like | -3.5 | 1.70E-06 |
| LOC129313390 | 129313390 | receptor-like protein kinase THESEUS 1 | -3.5 | 3.88E-06 |
| LOC129299723 | 129299723 | aspartyl protease family protein At5g10770-like | -3.5 | 8.66E-16 |
| LOC129314608 | 129314608 | ankyrin repeat-containing protein At5g02620-like | -3.5 | 3.29E-02 |

| | | | | |
|---|---|---|---|---|
| LOC129316191 | 129316191 | GDSL esterase/lipase At5g14450-like | -3.5 | 4.13E-03 |
| LOC129310939 | 129310939 | palmitoyl-acyl carrier protein thioesterase, chloroplastic-like | -3.5 | 2.33E-05 |
| LOC129288245 | 129288245 | uncharacterized LOC129288245 | -3.5 | 1.86E-02 |
| LOC129293248 | 129293248 | protein trichome birefringence-like 19 | -3.5 | 4.47E-14 |
| LOC129322186 | 129322186 | U-box domain-containing protein 5-like | -3.5 | 1.95E-05 |
| LOC129291452 | 129291452 | germin-like protein subfamily 3 member 1 | -3.5 | 4.95E-03 |
| LOC129315997 | 129315997 | vicianin hydrolase-like | -3.5 | 6.50E-03 |
| LOC129297135 | 129297135 | BURP domain protein RD22-like | -3.5 | 2.02E-02 |
| LOC129311781 | 129311781 | cationic peroxidase 1-like | -3.5 | 2.89E-02 |
| LOC129288178 | 129288178 | E3 ubiquitin-protein ligase ATL42 | -3.5 | 1.76E-03 |
| LOC129310118 | 129310118 | ABC transporter B family member 25-like | -3.5 | 2.44E-05 |
| LOC129285271 | 129285271 | anthocyanidin 3-O-glucosyltransferase 7-like | -3.5 | 1.28E-02 |
| LOC129320820 | 129320820 | RING-H2 finger protein ATL78-like | -3.5 | 2.84E-06 |
| LOC129300717 | 129300717 | 7-deoxyloganetin glucosyltransferase-like | -3.5 | 1.58E-02 |
| LOC129320507 | 129320507 | probable xyloglucan endotransglucosylase/hydrolase protein 23 | -3.5 | 3.52E-03 |
| LOC129318897 | 129318897 | protein NUCLEAR FUSION DEFECTIVE 4 | -3.6 | 1.07E-06 |
| LOC129284820 | 129284820 | uncharacterized LOC129284820 | -3.6 | 1.39E-03 |
| LOC129303407 | 129303407 | patatin-like protein 2 | -3.6 | 1.06E-07 |
| LOC129286750 | 129286750 | protein NUCLEAR FUSION DEFECTIVE 4 | -3.6 | 2.00E-10 |
| LOC129308873 | 129308873 | uncharacterized LOC129308873 | -3.6 | 3.04E-11 |
| LOC129302025 | 129302025 | transcription factor AS1-like | -3.6 | 1.44E-02 |
| LOC129311565 | 129311565 | protein JINGUBANG | -3.6 | 8.62E-09 |
| LOC129321993 | 129321993 | uncharacterized LOC129321993 | -3.6 | 1.06E-21 |
| LOC129286899 | 129286899 | probable hexosyltransferase MUCI70 | -3.6 | 2.87E-03 |
| LOC129318920 | 129318920 | uncharacterized LOC129318920 | -3.7 | 2.83E-02 |
| LOC129297933 | 129297933 | uncharacterized LOC129297933 | -3.7 | 3.41E-02 |
| LOC129286066 | 129286066 | auxin-responsive protein SAUR21-like | -3.7 | 3.18E-05 |
| LOC129298942 | 129298942 | uncharacterized LOC129298942 | -3.7 | 3.19E-02 |
| LOC129295001 | 129295001 | receptor-like protein kinase | -3.7 | 1.96E-02 |
| LOC129310926 | 129310926 | protein FANTASTIC FOUR 1-like | -3.7 | 1.41E-06 |
| LOC129312095 | 129312095 | glyoxylase I 4-like | -3.7 | 3.84E-05 |
| LOC129307422 | 129307422 | elongation of fatty acids protein 3-like | -3.8 | 7.58E-03 |
| LOC129302892 | 129302892 | inducible nitrate reductase [NADH] 2-like | -3.8 | 1.05E-03 |
| LOC129309086 | 129309086 | LRR receptor-like serine/threonine-protein kinase RGI3 | -3.8 | 6.79E-06 |
| LOC129305023 | 129305023 | alkane hydroxylase MAH1-like | -3.8 | 2.55E-09 |
| LOC129314851 | 129314851 | uncharacterized LOC129314851 | -3.8 | 3.60E-04 |
| LOC129295736 | 129295736 | beta-galactosidase 1-like | -3.8 | 2.08E-14 |
| LOC129303583 | 129303583 | protein PHOSPHATE-INDUCED 1-like | -3.9 | 2.32E-04 |
| LOC129301160 | 129301160 | photosystem II reaction center W protein, chloroplastic-like | -3.9 | 1.35E-03 |
| LOC129314919 | 129314919 | transcription factor MYB46-like | -3.9 | 5.18E-04 |
| LOC129290873 | 129290873 | disease resistance-like protein DSC1 | -3.9 | 6.69E-04 |
| LOC129304074 | 129304074 | uncharacterized LOC129304074 | -3.9 | 1.42E-03 |
| LOC129284445 | 129284445 | adenylate isopentenyltransferase 7, mitochondrial-like | -3.9 | 6.97E-05 |
| LOC129290567 | 129290567 | F-box/LRR-repeat protein 3-like | -3.9 | 1.39E-08 |
| LOC129303446 | 129303446 | PAN domain-containing protein At5g03700 | -3.9 | 3.37E-11 |
| LOC129298195 | 129298195 | ethylene-responsive transcription factor SHINE 2-like | -3.9 | 3.23E-04 |
| LOC129301804 | 129301804 | transcription repressor OFP7-like | -3.9 | 1.10E-02 |
| LOC129320609 | 129320609 | cytochrome P450 734A1-like | -3.9 | 1.03E-03 |

| | | | | |
|---|---|---|---|---|
| LOC129301148 | 129301148 | expansin-A8-like | -3.9 | 3.88E-06 |
| LOC129306485 | 129306485 | GDSL esterase/lipase At5g33370-like | -3.9 | 9.70E-06 |
| LOC129316417 | 129316417 | uncharacterized LOC129316417 | -4.0 | 3.35E-02 |
| LOC129291783 | 129291783 | uncharacterized LOC129291783 | -4.0 | 4.51E-06 |
| LOC129320267 | 129320267 | uncharacterized LOC129320267 | -4.0 | 3.40E-06 |
| LOC129293707 | 129293707 | uncharacterized LOC129293707 | -4.0 | 9.89E-03 |
| LOC129311163 | 129311163 | peroxidase 64-like | -4.0 | 1.15E-10 |
| LOC129300986 | 129300986 | uncharacterized LOC129300986 | -4.0 | 2.51E-02 |
| LOC129321612 | 129321612 | mavicyanin-like | -4.0 | 2.79E-04 |
| LOC129309350 | 129309350 | sugar transporter ERD6-like 16 | -4.0 | 1.47E-03 |
| LOC129311484 | 129311484 | probable mannitol dehydrogenase | -4.0 | 2.87E-13 |
| LOC129304076 | 129304076 | protein NRT1/ PTR FAMILY 5.1-like | -4.0 | 2.59E-10 |
| LOC129288095 | 129288095 | protein CNGC15a-like | -4.0 | 1.75E-13 |
| LOC129288855 | 129288855 | uncharacterized LOC129288855 | -4.1 | 2.45E-07 |
| LOC129314674 | 129314674 | receptor-like protein kinase THESEUS 1 | -4.1 | 3.60E-02 |
| LOC129297754 | 129297754 | disease resistance protein RPM1-like | -4.1 | 1.09E-02 |
| LOC129285521 | 129285521 | uncharacterized LOC129285521 | -4.1 | 3.92E-02 |
| LOC129320953 | 129320953 | uncharacterized LOC129320953 | -4.1 | 4.90E-09 |
| LOC129288548 | 129288548 | probable LRR receptor-like serine/threonine-protein kinase At4g | -4.1 | 8.21E-05 |
| LOC129309775 | 129309775 | peroxidase A2-like | -4.1 | 1.08E-11 |
| LOC129316285 | 129316285 | repetitive proline-rich cell wall protein 1-like | -4.1 | 1.98E-15 |
| LOC129293231 | 129293231 | uncharacterized LOC129293231 | -4.2 | 1.01E-02 |
| LOC129322924 | 129322924 | probable xyloglucan endotransglucosylase/hydrolase protein 6 | -4.2 | 7.25E-12 |
| LOC129293797 | 129293797 | 3-hydroxyisobutyryl-CoA hydrolase-like protein 3, mitochondria | -4.2 | 6.69E-03 |
| LOC129285252 | 129285252 | cationic peroxidase 2-like | -4.2 | 1.68E-04 |
| LOC129295183 | 129295183 | glycosyltransferase family 92 protein RCOM_0530710-like | -4.2 | 1.85E-03 |
| LOC129301274 | 129301274 | UDP-glycosyltransferase 708G1-like | -4.2 | 3.39E-02 |
| LOC129306451 | 129306451 | ABC transporter B family member 15-like | -4.2 | 4.73E-07 |
| LOC129298763 | 129298763 | protein SMALL AUXIN UP-REGULATED RNA 12 | -4.2 | 3.85E-02 |
| LOC129303682 | 129303682 | WAT1-related protein At4g08290-like | -4.2 | 2.27E-04 |
| LOC129290579 | 129290579 | protein CURVATURE THYLAKOID 1B, chloroplastic-like | -4.2 | 1.93E-06 |
| LOC129313606 | 129313606 | zinc transporter 1 | -4.3 | 1.17E-04 |
| LOC129286996 | 129286996 | uncharacterized LOC129286996 | -4.3 | 3.86E-05 |
| LOC129288934 | 129288934 | uncharacterized protein At4g19900-like | -4.3 | 3.07E-09 |
| LOC129323056 | 129323056 | probable xyloglucan endotransglucosylase/hydrolase protein 33 | -4.3 | 2.57E-02 |
| LOC129287025 | 129287025 | beta-xylosidase/alpha-L-arabinofuranosidase 2-like | -4.3 | 1.21E-04 |
| LOC129294684 | 129294684 | probable WRKY transcription factor 70 | -4.3 | 3.31E-03 |
| LOC129293184 | 129293184 | uncharacterized LOC129293184 | -4.3 | 1.02E-07 |
| LOC129301838 | 129301838 | protein PHOSPHATE-INDUCED 1-like | -4.3 | 3.51E-06 |
| LOC129293678 | 129293678 | uncharacterized LOC129293678 | -4.3 | 1.52E-03 |
| LOC129288715 | 129288715 | protein trichome birefringence-like 25 | -4.3 | 1.12E-05 |
| LOC129314878 | 129314878 | uncharacterized LOC129314878 | -4.4 | 8.55E-05 |
| LOC129288979 | 129288979 | lectin-domain containing receptor kinase VI.4-like | -4.4 | 3.04E-07 |
| LOC129304791 | 129304791 | uncharacterized LOC129304791 | -4.4 | 1.04E-03 |
| LOC129320511 | 129320511 | xyloglucan endotransglucosylase/hydrolase protein 22-like | -4.4 | 1.57E-06 |
| LOC129318529 | 129318529 | GDSL esterase/lipase EXL3-like | -4.5 | 3.18E-09 |
| LOC129305457 | 129305457 | extensin-like | -4.5 | 1.73E-06 |
| LOC129285430 | 129285430 | zingipain-1-like | -4.5 | 8.57E-04 |

| | | | | |
|---|---|---|---|---|
| LOC129322133 | 129322133 | meiotic recombination protein SPO11-2 | -4.5 | 2.70E-03 |
| LOC129296903 | 129296903 | mediator of RNA polymerase II transcription subunit 20a-like | -4.5 | 9.00E-03 |
| LOC129293485 | 129293485 | sugar transporter ERD6-like 16 | -4.5 | 1.59E-04 |
| LOC129288634 | 129288634 | cytochrome b561 and DOMON domain-containing protein At3g | -4.5 | 1.66E-06 |
| LOC129308140 | 129308140 | probable glycosyltransferase At3g07620 | -4.5 | 3.87E-16 |
| LOC129290515 | 129290515 | pentatricopeptide repeat-containing protein At5g48730, chloropl | -4.5 | 6.13E-03 |
| LOC129289530 | 129289530 | indole-3-pyruvate monooxygenase YUCCA2-like | -4.6 | 1.15E-05 |
| LOC129320949 | 129320949 | uncharacterized LOC129320949 | -4.6 | 5.19E-03 |
| LOC129303452 | 129303452 | carboxylesterase 1-like | -4.6 | 3.81E-03 |
| LOC129297432 | 129297432 | probable sarcosine oxidase | -4.6 | 1.45E-08 |
| LOC129284909 | 129284909 | uncharacterized LOC129284909 | -4.6 | 6.12E-03 |
| LOC129311729 | 129311729 | peroxidase 10 | -4.7 | 9.77E-03 |
| LOC129321353 | 129321353 | protein STRICTOSIDINE SYNTHASE-LIKE 10-like | -4.7 | 2.01E-05 |
| LOC129285234 | 129285234 | leucine-rich repeat receptor protein kinase HPCA1-like | -4.7 | 2.33E-13 |
| LOC129286864 | 129286864 | uncharacterized LOC129286864 | -4.7 | 5.96E-09 |
| LOC129314278 | 129314278 | fasciclin-like arabinogalactan protein 9 | -4.8 | 6.12E-04 |
| LOC129287438 | 129287438 | V-type proton ATPase catalytic subunit A-like | -4.8 | 2.88E-03 |
| LOC129289616 | 129289616 | uncharacterized LOC129289616 | -4.8 | 1.15E-05 |
| LOC129315246 | 129315246 | uncharacterized LOC129315246 | -4.8 | 3.80E-02 |
| LOC129285854 | 129285854 | uncharacterized LOC129285854 | -4.8 | 2.12E-04 |
| LOC129291555 | 129291555 | transcription factor MYC2-like | -4.8 | 1.53E-10 |
| LOC129310345 | 129310345 | probable membrane-associated kinase regulator 6 | -4.9 | 4.55E-03 |
| LOC129304350 | 129304350 | alkane hydroxylase MAH1-like | -4.9 | 3.43E-11 |
| LOC129315202 | 129315202 | uncharacterized LOC129315202 | -4.9 | 3.55E-02 |
| LOC129284550 | 129284550 | protein PHYTOCHROME KINASE SUBSTRATE 1-like | -4.9 | 3.66E-06 |
| LOC129300210 | 129300210 | ETHYLENE INSENSITIVE 3-like 1 protein | -4.9 | 3.37E-04 |
| LOC129297611 | 129297611 | mechanosensitive ion channel protein 10-like | -5.0 | 1.09E-02 |
| LOC129289665 | 129289665 | kunitz-type elastase inhibitor BrEI-like | -5.0 | 6.09E-05 |
| LOC129287456 | 129287456 | putative invertase inhibitor | -5.1 | 4.56E-03 |
| LOC129294734 | 129294734 | RING-H2 finger protein ATL54-like | -5.1 | 1.64E-05 |
| LOC129287209 | 129287209 | GDSL esterase/lipase At5g45910 | -5.2 | 4.84E-07 |
| LOC129285353 | 129285353 | G-type lectin S-receptor-like serine/threonine-protein kinase At1 | -5.3 | 1.02E-06 |
| LOC129286997 | 129286997 | uncharacterized LOC129286997 | -5.3 | 3.25E-03 |
| LOC129309979 | 129309979 | UPF0481 protein At3g47200-like | -5.4 | 3.46E-02 |
| LOC129299826 | 129299826 | xyloglucan endotransglucosylase/hydrolase protein 22-like | -5.5 | 2.49E-13 |
| LOC129299655 | 129299655 | glycerol-3-phosphate 2-O-acyltransferase 6-like | -5.5 | 1.49E-02 |
| LOC129306104 | 129306104 | rac-like GTP-binding protein RAC13 | -5.5 | 6.15E-03 |
| LOC129286970 | 129286970 | cysteine-rich receptor-like protein kinase 25 | -5.6 | 4.27E-03 |
| LOC129296593 | 129296593 | protein CURVATURE THYLAKOID 1B, chloroplastic-like | -5.6 | 5.97E-06 |
| LOC129297807 | 129297807 | shikimate O-hydroxycinnamoyltransferase-like | -5.6 | 6.62E-07 |
| LOC129300186 | 129300186 | probable disease resistance protein At4g27220 | -5.7 | 2.45E-02 |
| LOC129315004 | 129315004 | UPF0481 protein At3g47200-like | -5.7 | 2.77E-03 |
| LOC129305594 | 129305594 | beta-amyrin 24-hydroxylase-like | -5.8 | 3.30E-08 |
| LOC129312633 | 129312633 | uncharacterized LOC129312633 | -5.8 | 1.85E-02 |
| LOC129292664 | 129292664 | hevamine-A-like | -5.8 | 3.45E-03 |
| LOC129315597 | 129315597 | purple acid phosphatase-like | -5.8 | 2.87E-02 |
| LOC129297990 | 129297990 | BURP domain-containing protein 1-like | -5.8 | 7.66E-03 |
| LOC129309822 | 129309822 | protein JINGUBANG | -5.9 | 1.36E-02 |

| | | | | |
|---|---|---|---|---|
| LOC129289528 | 129289528 | uncharacterized LOC129289528 | -5.9 | 1.72E-02 |
| LOC129296417 | 129296417 | uncharacterized LOC129296417 | -5.9 | 1.38E-03 |
| LOC129305832 | 129305832 | pectate lyase-like | -5.9 | 5.40E-03 |
| LOC129287587 | 129287587 | plant UBX domain-containing protein 8-like | -6.2 | 9.60E-03 |
| LOC129308339 | 129308339 | uncharacterized LOC129308339 | -6.2 | 4.50E-02 |
| LOC129293871 | 129293871 | protein SMALL AUXIN UP-REGULATED RNA 12-like | -6.4 | 9.49E-09 |
| LOC129312327 | 129312327 | cytosolic sulfotransferase 15-like | -6.4 | 2.29E-05 |
| LOC129310637 | 129310637 | uncharacterized LOC129310637 | -6.5 | 2.88E-02 |
| LOC129299197 | 129299197 | linoleate 13S-lipoxygenase 2-1, chloroplastic-like | -6.8 | 1.41E-04 |
| LOC129312007 | 129312007 | F-box/FBD/LRR-repeat protein At3g26920-like | -6.9 | 1.44E-02 |
| LOC129297712 | 129297712 | uncharacterized LOC129297712 | -6.9 | 1.43E-02 |
| LOC129295047 | 129295047 | uncharacterized LOC129295047 | -6.9 | 6.72E-04 |
| LOC129317179 | 129317179 | uncharacterized LOC129317179 | -6.9 | 1.48E-02 |
| LOC129296287 | 129296287 | hevamine-A-like | -7.0 | 2.04E-05 |
| LOC129284431 | 129284431 | uncharacterized LOC129284431 | -7.0 | 2.75E-03 |
| LOC129287459 | 129287459 | geraniol 8-hydroxylase-like | -7.2 | 2.34E-05 |
| LOC129322564 | 129322564 | protein STRICTOSIDINE SYNTHASE-LIKE 10-like | -7.2 | 3.74E-02 |
| LOC129306851 | 129306851 | receptor-like protein kinase FERONIA | -7.5 | 2.71E-02 |
| LOC129288056 | 129288056 | receptor-like protein kinase | -7.5 | 9.97E-06 |
| LOC129311588 | 129311588 | probable xyloglucan endotransglucosylase/hydrolase protein 6 | -8.2 | 1.69E-33 |
| LOC129289527 | 129289527 | uncharacterized LOC129289527 | -8.4 | 2.59E-07 |
| LOC129310240 | 129310240 | receptor-like protein kinase FERONIA | -8.6 | 7.82E-06 |
| LOC129307109 | 129307109 | short chain aldehyde dehydrogenase 1-like | -10.4 | 4.75E-02 |

Genes differentially expressed in the fifth leaf of one-month-old *Prosopis cineraria* (Indian cultivar) seedlings treated with 5% PEG-6000 versus control are shown. The genes were filtered based on Benjamini Hochberg False Discovery Rate adjusted *P* -value < 0.05 and -1 < $\log_2$(fold-change) > 1. The NCBI gene loci and gene IDs corresponding to the genome of the Arabian cultivar of *P. cineraria* (GCF_029017545.1) are shown.

**Supplementary Table S2. Functional enrichment analysis of the gene networks of *Prosopis cineraria* under drought**

| Number of bac | Number of net | Category | Description | FDR value | *P* -value |
|---|---|---|---|---|---|
| **Up-regulated gene network** | | | | | |
| 2020 | 96 | GO Biological Process | Response to abiotic stimulus | 6.00E-19 | 1.04E-22 |
| 11632 | 277 | GO Molecular Function | Binding | 5.50E-18 | 1.72E-21 |
| 3346 | 120 | GO Biological Process | Regulation of primary metabolic process | 2.42E-15 | 8.41E-19 |
| 627 | 47 | GO Biological Process | Response to temperature stimulus | 6.54E-15 | 3.41E-18 |
| 3384 | 119 | GO Biological Process | Regulation of cellular metabolic process | 6.54E-15 | 5.51E-18 |
| 5300 | 159 | GO Biological Process | Regulation of cellular process | 6.54E-15 | 3.59E-18 |
| 3272 | 116 | GO Biological Process | Regulation of nitrogen compound metabolic process | 9.07E-15 | 9.45E-18 |
| 3735 | 125 | GO Biological Process | Regulation of metabolic process | 2.17E-14 | 2.64E-17 |
| 6689 | 183 | GO Biological Process | Biological regulation | 2.17E-14 | 2.64E-17 |
| 5971 | 169 | GO Biological Process | Regulation of biological process | 2.97E-14 | 4.65E-17 |
| 2909 | 106 | GO Biological Process | Regulation of biosynthetic process | 3.72E-14 | 6.46E-17 |
| 3486 | 117 | GO Biological Process | Regulation of macromolecule metabolic process | 1.82E-13 | 3.48E-16 |
| 2880 | 103 | GO Biological Process | Regulation of cellular biosynthetic process | 3.33E-13 | 6.93E-16 |
| 13552 | 290 | GO Biological Process | Cellular process | 7.30E-13 | 1.65E-15 |
| 2622 | 96 | GO Biological Process | Regulation of nucleobase-containing compound metabolic process | 9.11E-13 | 2.22E-15 |
| 2796 | 99 | GO Biological Process | Regulation of macromolecule biosynthetic process | 2.16E-12 | 5.63E-15 |
| 2508 | 91 | GO Biological Process | Regulation of RNA metabolic process | 8.78E-12 | 2.44E-14 |
| 2386 | 88 | GO Biological Process | Regulation of transcription, DNA-templated | 1.01E-11 | 2.97E-14 |
| 2517 | 92 | GO Molecular Function | DNA binding | 1.77E-11 | 1.11E-14 |
| 1582 | 67 | GO Biological Process | Response to oxygen-containing compound | 4.72E-11 | 1.64E-13 |
| 2960 | 99 | GO Biological Process | Response to chemical | 4.80E-11 | 1.75E-13 |
| 3061 | 101 | GO Biological Process | Regulation of gene expression | 5.44E-11 | 2.08E-13 |
| 4469 | 130 | GO Molecular Function | Nucleic acid binding | 1.99E-10 | 2.07E-13 |
| 2556 | 90 | GO Molecular Function | Protein binding | 1.99E-10 | 1.87E-13 |
| 1797 | 71 | GO Molecular Function | Transcription regulator activity | 4.20E-10 | 6.87E-13 |
| 7692 | 188 | GO Molecular Function | Heterocyclic compound binding | 4.20E-10 | 6.58E-13 |
| 7733 | 188 | GO Molecular Function | Organic cyclic compound binding | 5.00E-10 | 1.09E-12 |
| 1642 | 64 | GO Molecular Function | DNA-binding transcription factor activity | 7.83E-09 | 1.96E-11 |
| 6206 | 156 | GO Biological Process | Response to stimulus | 8.55E-09 | 3.41E-11 |
| 123 | 17 | GO Molecular Function | Protein heterodimerization activity | 1.74E-08 | 4.91E-11 |
| 1335 | 55 | GO Biological Process | Positive regulation of biological process | 2.09E-08 | 8.69E-11 |
| 538 | 33 | GO Molecular Function | Structural molecule activity | 2.16E-08 | 6.75E-11 |
| 379 | 27 | GO Biological Process | Response to water | 4.14E-08 | 1.80E-10 |
| 598 | 34 | GO Biological Process | Cell cycle | 4.98E-08 | 2.29E-10 |
| 909 | 43 | GO Biological Process | Positive regulation of metabolic process | 4.98E-08 | 2.25E-10 |
| 3820 | 108 | GO Biological Process | Response to stress | 5.58E-08 | 2.71E-10 |
| 368 | 26 | GO Biological Process | Response to water deprivation | 9.23E-08 | 4.65E-10 |

| | | | | | |
|---|---|---|---|---|---|
| 827 | 40 | GO Biological Process | Positive regulation of macromolecule metabolic process | 1.04E-07 | 5.41E-10 |
| 727 | 37 | GO Biological Process | Response to inorganic substance | 1.24E-07 | 6.69E-10 |
| 43 | 11 | GO Molecular Function | Structural constituent of chromatin | 1.33E-07 | 4.58E-10 |
| 2952 | 89 | GO Biological Process | Organic substance biosynthetic process | 1.44E-07 | 7.98E-10 |
| 8044 | 183 | GO Biological Process | Cellular metabolic process | 1.73E-07 | 1.02E-09 |
| 614 | 33 | GO Biological Process | Positive regulation of biosynthetic process | 2.68E-07 | 1.63E-09 |
| 254 | 21 | GO Biological Process | Response to heat | 2.73E-07 | 1.71E-09 |
| 2858 | 86 | GO Biological Process | Cellular biosynthetic process | 2.84E-07 | 1.82E-09 |
| 562 | 31 | GO Biological Process | Positive regulation of macromolecule biosynthetic process | 4.40E-07 | 2.90E-09 |
| 921 | 41 | GO Biological Process | Response to lipid | 4.84E-07 | 3.28E-09 |
| 3103 | 90 | GO Biological Process | Biosynthetic process | 5.83E-07 | 4.05E-09 |
| 411 | 26 | GO Biological Process | Cell cycle process | 5.83E-07 | 4.12E-09 |
| 1543 | 56 | GO Biological Process | Response to hormone | 6.82E-07 | 4.97E-09 |
| 2285 | 72 | KEGG Pathways | Metabolic pathways | 8.37E-07 | 8.34E-09 |
| 1219 | 48 | KEGG Pathways | Biosynthesis of secondary metabolites | 8.37E-07 | 6.20E-09 |
| 1959 | 65 | GO Biological Process | Response to organic substance | 1.02E-06 | 7.58E-09 |
| 1661 | 58 | GO Biological Process | Cellular response to chemical stimulus | 1.25E-06 | 9.76E-09 |
| 1257 | 49 | GO Molecular Function | Sequence-specific DNA binding | 1.54E-06 | 5.80E-09 |
| 233 | 19 | GO Biological Process | Cellular response to decreased oxygen levels | 1.58E-06 | 1.26E-08 |
| 540 | 29 | GO Biological Process | Response to abscisic acid | 2.00E-06 | 1.67E-08 |
| 388 | 24 | GO Biological Process | Cell division | 2.83E-06 | 2.46E-08 |
| 805 | 36 | GO Biological Process | Positive regulation of nitrogen compound metabolic process | 3.17E-06 | 2.81E-08 |
| 596 | 30 | GO Biological Process | Positive regulation of cellular biosynthetic process | 4.10E-06 | 3.70E-08 |
| 529 | 28 | GO Biological Process | Positive regulation of transcription, DNA-templated | 4.30E-06 | 3.95E-08 |
| 605 | 30 | GO Biological Process | Positive regulation of RNA metabolic process | 5.21E-06 | 5.07E-08 |
| 231 | 18 | GO Biological Process | Cellular response to hypoxia | 5.86E-06 | 5.80E-08 |
| 3214 | 88 | GO Biological Process | Cellular response to stimulus | 8.18E-06 | 8.52E-08 |
| 391 | 23 | GO Biological Process | Response to cold | 1.07E-05 | 1.13E-07 |
| 318 | 20 | KEGG Pathways | Ribosome | 1.28E-05 | 2.84E-07 |
| 1630 | 54 | GO Biological Process | Reproductive process | 1.57E-05 | 1.74E-07 |
| 830 | 35 | GO Biological Process | Positive regulation of cellular metabolic process | 1.57E-05 | 1.71E-07 |
| 1376 | 48 | GO Biological Process | Post-embryonic development | 1.91E-05 | 2.15E-07 |
| 200 | 16 | GO Biological Process | Mitotic cell cycle process | 1.98E-05 | 2.27E-07 |
| 767 | 33 | GO Biological Process | Negative regulation of cellular process | 2.14E-05 | 2.53E-07 |
| 10167 | 208 | GO Biological Process | Metabolic process | 2.32E-05 | 2.82E-07 |
| 1072 | 40 | GO Biological Process | Positive regulation of cellular process | 3.84E-05 | 4.73E-07 |
| 954 | 38 | GO Molecular Function | Transcription cis-regulatory region binding | 4.88E-05 | 1.99E-07 |
| 341 | 20 | GO Biological Process | Regulation of post-embryonic development | 6.39E-05 | 7.99E-07 |
| 250 | 17 | GO Biological Process | Mitotic cell cycle | 6.40E-05 | 8.12E-07 |
| 703 | 30 | GO Biological Process | Response to radiation | 8.24E-05 | 1.06E-06 |
| 1242 | 43 | GO Biological Process | Cellular response to stress | 8.91E-05 | 1.17E-06 |
| 257 | 17 | GO Biological Process | Regulation of cell cycle | 8.91E-05 | 1.16E-06 |
| 30 | 7 | GO Biological Process | Regulation of ethylene-activated signaling pathway | 9.03E-05 | 1.22E-06 |
| 486 | 24 | GO Biological Process | Carboxylic acid biosynthetic process | 9.03E-05 | 1.21E-06 |
| 524 | 25 | GO Biological Process | Response to osmotic stress | 9.43E-05 | 1.31E-06 |
| 1131 | 40 | GO Biological Process | Reproductive structure development | 1.20E-04 | 1.71E-06 |

| | | | | | |
|---|---|---|---|---|---|
| 1140 | 40 | GO Biological Process | Cellular response to organic substance | 1.40E-04 | 2.07E-06 |
| 1363 | 45 | GO Biological Process | Developmental process involved in reproduction | 1.50E-04 | 2.21E-06 |
| 120 | 11 | KEGG Pathways | Cysteine and methionine metabolism | 1.70E-04 | 5.35E-06 |
| 289 | 17 | KEGG Pathways | Plant hormone signal transduction | 1.70E-04 | 5.15E-06 |
| 1100 | 40 | GO Molecular Function | Double-stranded DNA binding | 1.80E-04 | 8.84E-07 |
| 659 | 29 | GO Molecular Function | Protein dimerization activity | 1.80E-04 | 8.94E-07 |
| 902 | 34 | GO Biological Process | Hormone-mediated signaling pathway | 1.90E-04 | 2.88E-06 |
| 137 | 12 | GO Biological Process | Regulation of cell cycle process | 2.00E-04 | 3.11E-06 |
| 680 | 28 | GO Biological Process | Response to light stimulus | 3.00E-04 | 4.66E-06 |
| 107 | 11 | GO Molecular Function | Protein kinase regulator activity | 3.40E-04 | 1.92E-06 |
| 1233 | 41 | GO Biological Process | Negative regulation of biological process | 3.50E-04 | 5.45E-06 |
| 136 | 11 | KEGG Pathways | MAPK signaling pathway - plant | 3.60E-04 | 1.61E-05 |
| 2594 | 69 | GO Biological Process | Multicellular organismal process | 4.70E-04 | 7.42E-06 |
| 829 | 31 | GO Biological Process | Shoot system development | 5.80E-04 | 9.28E-06 |
| 788 | 30 | GO Biological Process | Regulation of developmental process | 5.80E-04 | 9.32E-06 |
| 671 | 27 | GO Biological Process | Small molecule biosynthetic process | 6.30E-04 | 1.03E-05 |
| 966 | 34 | GO Biological Process | Carboxylic acid metabolic process | 7.00E-04 | 1.17E-05 |
| 384 | 20 | GO Molecular Function | Structural constituent of ribosome | 7.10E-04 | 4.43E-06 |
| 246 | 15 | GO Biological Process | Phosphorelay signal transduction system | 7.20E-04 | 1.23E-05 |
| 84 | 9 | GO Biological Process | Regulation of mitotic cell cycle | 7.20E-04 | 1.22E-05 |
| 159 | 12 | GO Biological Process | Nuclear division | 7.40E-04 | 1.28E-05 |
| 2344 | 63 | GO Biological Process | Multicellular organism development | 8.30E-04 | 1.45E-05 |
| 222 | 14 | GO Biological Process | Monocarboxylic acid biosynthetic process | 9.60E-04 | 1.68E-05 |
| 228 | 14 | GO Biological Process | Response to wounding | 1.20E-03 | 2.23E-05 |
| 1087 | 36 | GO Biological Process | Oxoacid metabolic process | 1.20E-03 | 2.26E-05 |
| 196 | 13 | GO Biological Process | Organelle fission | 1.20E-03 | 2.04E-05 |
| 584 | 24 | GO Biological Process | Intracellular signal transduction | 1.30E-03 | 2.30E-05 |
| 9277 | 184 | GO Biological Process | Organic substance metabolic process | 1.30E-03 | 2.38E-05 |
| 192 | 12 | KEGG Pathways | Spliceosome | 1.40E-03 | 7.25E-05 |
| 444 | 20 | GO Biological Process | Response to salt stress | 1.70E-03 | 3.29E-05 |
| 20 | 5 | GO Biological Process | Negative regulation of ethylene-activated signaling pathway | 1.70E-03 | 3.24E-05 |
| 1475 | 44 | GO Biological Process | Cellular nitrogen compound biosynthetic process | 1.70E-03 | 3.26E-05 |
| 1835 | 51 | GO Biological Process | System development | 2.30E-03 | 4.55E-05 |
| 146 | 10 | KEGG Pathways | Oxidative phosphorylation | 2.40E-03 | 1.40E-04 |
| 241 | 13 | KEGG Pathways | Biosynthesis of amino acids | 2.40E-03 | 1.50E-04 |
| 497 | 21 | GO Biological Process | Negative regulation of nitrogen compound metabolic process | 2.50E-03 | 5.09E-05 |
| 1507 | 44 | GO Biological Process | Organonitrogen compound biosynthetic process | 2.60E-03 | 5.30E-05 |
| 500 | 21 | GO Biological Process | Cellular response to lipid | 2.70E-03 | 5.53E-05 |
| 424 | 20 | GO Molecular Function | Enzyme binding | 2.70E-03 | 1.76E-05 |
| 392 | 18 | GO Biological Process | Negative regulation of biosynthetic process | 3.10E-03 | 6.39E-05 |
| 2728 | 68 | GO Biological Process | Anatomical structure development | 3.10E-03 | 6.48E-05 |
| 254 | 14 | GO Biological Process | Negative regulation of RNA metabolic process | 3.20E-03 | 6.76E-05 |
| 194 | 12 | GO Biological Process | Ethylene-activated signaling pathway | 3.80E-03 | 7.95E-05 |
| 25 | 5 | GO Biological Process | Negative regulation of mitotic nuclear division | 3.80E-03 | 8.18E-05 |
| 54 | 6 | KEGG Pathways | Arginine and proline metabolism | 4.00E-03 | 2.90E-04 |
| 480 | 20 | GO Biological Process | Negative regulation of cellular metabolic process | 4.20E-03 | 9.18E-05 |

| | | | | | |
|---|---|---|---|---|---|
| 483 | 20 | GO Biological Process | Monocarboxylic acid metabolic process | 4.50E-03 | 9.95E-05 |
| 2875 | 70 | GO Biological Process | Developmental process | 4.60E-03 | 1.00E-04 |
| 448 | 19 | GO Biological Process | Flower development | 4.90E-03 | 1.10E-04 |
| 1705 | 47 | GO Biological Process | Small molecule metabolic process | 5.00E-03 | 1.10E-04 |
| 780 | 27 | GO Biological Process | Regulation of response to stimulus | 5.40E-03 | 1.20E-04 |
| 46 | 6 | GO Biological Process | Negative regulation of mitotic cell cycle | 5.80E-03 | 1.30E-04 |
| 147 | 10 | GO Biological Process | Regulation of flower development | 6.50E-03 | 1.50E-04 |
| 241 | 13 | GO Biological Process | Negative regulation of transcription, DNA-templated | 6.50E-03 | 1.50E-04 |
| 386 | 17 | GO Biological Process | Negative regulation of cellular biosynthetic process | 6.80E-03 | 1.60E-04 |
| 179 | 11 | GO Biological Process | Regulation of organelle organization | 6.80E-03 | 1.70E-04 |
| 630 | 23 | GO Biological Process | Translation | 7.60E-03 | 1.90E-04 |
| 1071 | 33 | GO Biological Process | Organic cyclic compound biosynthetic process | 7.60E-03 | 1.80E-04 |
| 632 | 23 | GO Biological Process | Protein-containing complex organization | 7.80E-03 | 1.90E-04 |
| 893 | 29 | GO Biological Process | Cellular component assembly | 7.90E-03 | 2.00E-04 |
| 97 | 8 | GO Biological Process | Cellular response to fatty acid | 8.10E-03 | 2.00E-04 |
| 51 | 6 | GO Biological Process | Sulfur amino acid biosynthetic process | 8.70E-03 | 2.20E-04 |
| 767 | 26 | GO Biological Process | Negative regulation of metabolic process | 8.70E-03 | 2.20E-04 |
| 74 | 7 | GO Biological Process | Negative regulation of cell cycle | 8.90E-03 | 2.30E-04 |
| 32 | 5 | GO Biological Process | Microtubule cytoskeleton organization involved in mitosis | 8.90E-03 | 2.30E-04 |
| 1917 | 50 | GO Biological Process | Signaling | 9.30E-03 | 2.40E-04 |
| 43 | 5 | KEGG Pathways | alpha-Linolenic acid metabolism | 9.60E-03 | 7.90E-04 |
| 8604 | 167 | GO Biological Process | Primary metabolic process | 9.80E-03 | 2.60E-04 |
| 190 | 11 | GO Biological Process | Meiotic cell cycle | 1.00E-02 | 2.70E-04 |
| 1879 | 49 | GO Biological Process | Signal transduction | 1.04E-02 | 2.80E-04 |
| 367 | 16 | GO Biological Process | Negative regulation of macromolecule biosynthetic process | 1.04E-02 | 2.80E-04 |
| 608 | 22 | GO Biological Process | Regulation of transcription by RNA polymerase II | 1.05E-02 | 2.90E-04 |
| 331 | 15 | GO Biological Process | Regulation of signal transduction | 1.06E-02 | 2.90E-04 |
| 132 | 9 | GO Biological Process | Rhythmic process | 1.13E-02 | 3.10E-04 |
| 701 | 24 | GO Biological Process | Amide biosynthetic process | 1.19E-02 | 3.40E-04 |
| 7251 | 144 | GO Biological Process | Nitrogen compound metabolic process | 1.21E-02 | 3.40E-04 |
| 107 | 8 | GO Biological Process | Phenylpropanoid biosynthetic process | 1.32E-02 | 3.80E-04 |
| 2118 | 53 | GO Biological Process | Cell communication | 1.43E-02 | 4.10E-04 |
| 58 | 6 | GO Biological Process | Cold acclimation | 1.44E-02 | 4.20E-04 |
| 58 | 6 | GO Biological Process | Olefinic compound metabolic process | 1.44E-02 | 4.20E-04 |
| 938 | 29 | GO Biological Process | Aromatic compound biosynthetic process | 1.45E-02 | 4.30E-04 |
| 717 | 24 | GO Biological Process | Negative regulation of macromolecule metabolic process | 1.54E-02 | 4.60E-04 |
| 273 | 13 | GO Biological Process | Response to ethylene | 1.57E-02 | 4.70E-04 |
| 111 | 8 | GO Biological Process | Regulation of phosphorylation | 1.58E-02 | 4.80E-04 |
| 205 | 11 | GO Biological Process | Response to fatty acid | 1.64E-02 | 5.00E-04 |
| 27 | 5 | GO Molecular Function | Protein kinase inhibitor activity | 1.64E-02 | 1.10E-04 |
| 1932 | 49 | GO Biological Process | Gene expression | 1.71E-02 | 5.20E-04 |
| 208 | 11 | GO Biological Process | Protein folding | 1.82E-02 | 5.60E-04 |
| 144 | 9 | GO Biological Process | Phenylpropanoid metabolic process | 1.86E-02 | 5.70E-04 |
| 62 | 6 | GO Biological Process | Regulation of cyclin-dependent protein serine/threonine kinase activity | 1.88E-02 | 5.80E-04 |
| 62 | 6 | GO Biological Process | Lignin metabolic process | 1.88E-02 | 5.80E-04 |
| 40 | 5 | GO Biological Process | Regulation of jasmonic acid mediated signaling pathway | 1.88E-02 | 5.80E-04 |

| | | | | | |
|---|---|---|---|---|---|
| 88 | 7 | GO Biological Process | Jasmonic acid mediated signaling pathway | 1.94E-02 | 6.20E-04 |
| 562 | 20 | GO Biological Process | Protein-containing complex assembly | 2.05E-02 | 6.50E-04 |
| 741 | 24 | GO Biological Process | Peptide metabolic process | 2.23E-02 | 7.20E-04 |
| 1456 | 39 | GO Biological Process | Macromolecule biosynthetic process | 2.27E-02 | 7.40E-04 |
| 65 | 6 | GO Biological Process | Negative regulation of cell cycle process | 2.27E-02 | 7.30E-04 |
| 698 | 23 | GO Biological Process | Cellular response to oxygen-containing compound | 2.27E-02 | 7.30E-04 |
| 32 | 4 | KEGG Pathways | Phenylalanine metabolism | 2.36E-02 | 2.10E-03 |
| 151 | 9 | GO Biological Process | Response to reactive oxygen species | 2.40E-02 | 7.90E-04 |
| 151 | 9 | GO Biological Process | Sulfur compound biosynthetic process | 2.40E-02 | 7.90E-04 |
| 5 | 3 | GO Molecular Function | Glycogen (starch) synthase activity | 2.44E-02 | 1.80E-04 |
| 93 | 7 | GO Biological Process | Regulation of protein kinase activity | 2.53E-02 | 8.40E-04 |
| 67 | 6 | GO Biological Process | Spindle organization | 2.54E-02 | 8.50E-04 |
| 10 | 3 | GO Biological Process | Induced systemic resistance, jasmonic acid mediated signaling pathway | 2.63E-02 | 8.90E-04 |
| 45 | 5 | GO Biological Process | Olefinic compound biosynthetic process | 2.77E-02 | 9.50E-04 |
| 5275 | 108 | GO Biological Process | Organonitrogen compound metabolic process | 2.77E-02 | 9.40E-04 |
| 224 | 11 | GO Biological Process | Abscisic acid-activated signaling pathway | 2.89E-02 | 9.90E-04 |
| 32 | 5 | GO Molecular Function | DNA-binding transcription activator activity | 2.93E-02 | 2.30E-04 |
| 52 | 6 | GO Molecular Function | Cyclin-dependent protein serine/threonine kinase regulator activity | 3.00E-02 | 2.40E-04 |
| 26 | 4 | GO Biological Process | Methionine biosynthetic process | 3.03E-02 | 1.00E-03 |
| 1441 | 38 | GO Biological Process | Cellular component biogenesis | 3.27E-02 | 1.10E-03 |
| 161 | 9 | GO Biological Process | Vegetative to reproductive phase transition of meristem | 3.44E-02 | 1.20E-03 |
| 161 | 9 | GO Biological Process | Secondary metabolite biosynthetic process | 3.44E-02 | 1.20E-03 |
| 306 | 13 | GO Biological Process | Alpha-amino acid metabolic process | 3.60E-02 | 1.30E-03 |
| 18 | 4 | GO Molecular Function | RNA polymerase II complex binding | 3.64E-02 | 3.10E-04 |
| 163 | 9 | GO Biological Process | Cellular respiration | 3.66E-02 | 1.30E-03 |
| 12 | 3 | GO Biological Process | Mitotic recombination | 3.83E-02 | 1.40E-03 |
| 132 | 8 | GO Biological Process | Chromosome segregation | 3.83E-02 | 1.40E-03 |
| 3497 | 76 | GO Biological Process | Cellular nitrogen compound metabolic process | 3.83E-02 | 1.40E-03 |
| 471 | 17 | GO Biological Process | Phyllome development | 3.83E-02 | 1.40E-03 |
| 12 | 3 | GO Biological Process | L-methionine biosynthetic process | 3.83E-02 | 1.40E-03 |
| 102 | 7 | GO Biological Process | Meiotic nuclear division | 3.83E-02 | 1.40E-03 |
| 196 | 11 | GO Molecular Function | Chromatin binding | 3.95E-02 | 3.50E-04 |
| 29 | 4 | GO Biological Process | Mitotic spindle organization | 4.08E-02 | 1.50E-03 |
| 201 | 10 | GO Biological Process | Response to jasmonic acid | 4.08E-02 | 1.50E-03 |
| 29 | 4 | GO Biological Process | Oxylipin biosynthetic process | 4.08E-02 | 1.50E-03 |
| 29 | 4 | GO Biological Process | Response to calcium ion | 4.08E-02 | 1.50E-03 |
| 270 | 11 | KEGG Pathways | Carbon metabolism | 4.15E-02 | 4.00E-03 |
| 39 | 4 | KEGG Pathways | Circadian rhythm - plant | 4.15E-02 | 4.00E-03 |
| 51 | 5 | GO Biological Process | Regulation of mitotic cell cycle phase transition | 4.19E-02 | 1.60E-03 |
| 167 | 10 | GO Molecular Function | Transcription coregulator activity | 4.34E-02 | 3.90E-04 |
| 13 | 3 | GO Biological Process | NLS-bearing protein import into nucleus | 4.41E-02 | 1.70E-03 |
| 108 | 7 | GO Biological Process | Circadian rhythm | 4.89E-02 | 1.90E-03 |
| 108 | 7 | GO Biological Process | Negative regulation of signal transduction | 4.89E-02 | 1.90E-03 |
| 79 | 6 | GO Biological Process | Positive regulation of post-embryonic development | 4.89E-02 | 1.90E-03 |

**Down-regulated gene network**

| | | | | | |
|---|---|---|---|---|---|
| 2285 | 32 | KEGG Pathways | Metabolic pathways | 1.87E-13 | 1.38E-15 |

| | | | | | |
|---|---|---|---|---|---|
| 1219 | 24 | KEGG Pathways | Biosynthesis of secondary metabolites | 7.86E-13 | 1.16E-14 |
| 966 | 23 | GO Biological Process | Carboxylic acid metabolic process | 5.33E-12 | 9.26E-16 |
| 1705 | 27 | GO Biological Process | Small molecule metabolic process | 4.16E-11 | 2.89E-14 |
| 483 | 16 | GO Biological Process | Monocarboxylic acid metabolic process | 3.93E-10 | 3.41E-13 |
| 671 | 17 | GO Biological Process | Small molecule biosynthetic process | 3.50E-09 | 3.65E-12 |
| 90 | 9 | GO Biological Process | Response to blue light | 5.88E-09 | 7.14E-12 |
| 13552 | 65 | GO Biological Process | Cellular process | 9.50E-09 | 1.32E-11 |
| 680 | 16 | GO Biological Process | Response to light stimulus | 3.20E-08 | 5.01E-11 |
| 36 | 6 | KEGG Pathways | Fatty acid elongation | 7.27E-08 | 1.62E-09 |
| 254 | 11 | GO Biological Process | Fatty acid metabolic process | 7.93E-08 | 1.51E-10 |
| 149 | 9 | GO Biological Process | Fatty acid biosynthetic process | 2.34E-07 | 4.87E-10 |
| 222 | 10 | GO Biological Process | Monocarboxylic acid biosynthetic process | 3.38E-07 | 7.62E-10 |
| 486 | 13 | GO Biological Process | Carboxylic acid biosynthetic process | 3.64E-07 | 8.86E-10 |
| 2020 | 23 | GO Biological Process | Response to abiotic stimulus | 9.06E-07 | 2.52E-09 |
| 1543 | 20 | GO Biological Process | Response to hormone | 1.44E-06 | 4.25E-09 |
| 1959 | 22 | GO Biological Process | Response to organic substance | 2.41E-06 | 7.96E-09 |
| 3103 | 27 | GO Biological Process | Biosynthetic process | 5.93E-06 | 2.14E-08 |
| 902 | 15 | GO Biological Process | Hormone-mediated signaling pathway | 5.93E-06 | 2.17E-08 |
| 763 | 14 | GO Biological Process | Cellular lipid metabolic process | 5.93E-06 | 2.06E-08 |
| 30 | 5 | GO Biological Process | Chloroplast relocation | 9.98E-06 | 3.99E-08 |
| 10167 | 51 | GO Biological Process | Metabolic process | 1.21E-05 | 5.45E-08 |
| 1879 | 20 | GO Biological Process | Signal transduction | 2.08E-05 | 1.09E-07 |
| 6206 | 38 | GO Biological Process | Response to stimulus | 2.08E-05 | 1.08E-07 |
| 619 | 12 | GO Biological Process | Lipid biosynthetic process | 2.38E-05 | 1.32E-07 |
| 9277 | 47 | GO Biological Process | Organic substance metabolic process | 5.35E-05 | 3.16E-07 |
| 70 | 5 | KEGG Pathways | Glycine, serine and threonine metabolism | 6.57E-05 | 1.95E-06 |
| 70 | 5 | KEGG Pathways | Fatty acid metabolism | 6.57E-05 | 1.95E-06 |
| 441 | 10 | GO Biological Process | Response to auxin | 6.63E-05 | 4.03E-07 |
| 8044 | 43 | GO Biological Process | Cellular metabolic process | 6.63E-05 | 4.13E-07 |
| 2952 | 24 | GO Biological Process | Organic substance biosynthetic process | 9.31E-05 | 6.14E-07 |
| 2960 | 24 | GO Biological Process | Response to chemical | 9.52E-05 | 6.45E-07 |
| 23 | 4 | GO Biological Process | Phototropism | 1.20E-04 | 8.49E-07 |
| 8604 | 44 | GO Biological Process | Primary metabolic process | 1.30E-04 | 9.84E-07 |
| 63 | 5 | GO Biological Process | Auxin metabolic process | 1.60E-04 | 1.19E-06 |
| 64 | 5 | GO Biological Process | Response to red light | 1.70E-04 | 1.28E-06 |
| 284 | 8 | GO Biological Process | Regulation of hormone levels | 1.70E-04 | 1.32E-06 |
| 2858 | 23 | GO Biological Process | Cellular biosynthetic process | 1.70E-04 | 1.37E-06 |
| 270 | 7 | KEGG Pathways | Carbon metabolism | 2.40E-04 | 1.07E-05 |
| 133 | 6 | GO Biological Process | Cellular response to light stimulus | 2.70E-04 | 2.24E-06 |
| 306 | 8 | GO Biological Process | Alpha-amino acid metabolic process | 2.70E-04 | 2.27E-06 |
| 310 | 8 | GO Biological Process | Nucleotide metabolic process | 2.90E-04 | 2.50E-06 |
| 3214 | 24 | GO Biological Process | Cellular response to stimulus | 3.20E-04 | 2.80E-06 |
| 289 | 7 | KEGG Pathways | Plant hormone signal transduction | 3.20E-04 | 1.64E-05 |
| 221 | 7 | GO Biological Process | Purine ribonucleotide metabolic process | 3.30E-04 | 2.99E-06 |
| 80 | 5 | GO Biological Process | Nucleoside diphosphate phosphorylation | 3.80E-04 | 3.62E-06 |
| 9 | 3 | GO Biological Process | Purine ribonucleotide salvage | 4.20E-04 | 4.19E-06 |

| | | | | | |
|---|---|---|---|---|---|
| 89 | 5 | GO Biological Process | Cellular metabolic compound salvage | 5.60E-04 | 5.95E-06 |
| 248 | 7 | GO Biological Process | Auxin-activated signaling pathway | 5.70E-04 | 6.24E-06 |
| 12 | 3 | GO Biological Process | AMP metabolic process | 7.30E-04 | 8.62E-06 |
| 1661 | 16 | GO Biological Process | Cellular response to chemical stimulus | 7.70E-04 | 9.27E-06 |
| 642 | 10 | GO Biological Process | Organophosphate metabolic process | 8.70E-04 | 1.07E-05 |
| 241 | 6 | KEGG Pathways | Biosynthesis of amino acids | 9.90E-04 | 5.88E-05 |
| 391 | 8 | GO Biological Process | Response to cold | 1.00E-03 | 1.31E-05 |
| 77 | 4 | KEGG Pathways | Glyoxylate and dicarboxylate metabolism | 1.10E-03 | 7.20E-05 |
| 335 | 9 | GO Molecular Function | Acyltransferase activity, transferring groups other than amino-acyl grou | 1.30E-03 | 4.06E-07 |
| 23 | 4 | GO Molecular Function | Very-long-chain 3-ketoacyl-CoA synthase activity | 1.40E-03 | 8.49E-07 |
| 54 | 4 | GO Biological Process | Aromatic amino acid family biosynthetic process | 1.50E-03 | 1.93E-05 |
| 1582 | 15 | GO Biological Process | Response to oxygen-containing compound | 1.70E-03 | 2.23E-05 |
| 307 | 7 | GO Biological Process | Plastid organization | 1.80E-03 | 2.40E-05 |
| 18 | 3 | GO Biological Process | Cutin biosynthetic process | 1.90E-03 | 2.49E-05 |
| 313 | 7 | GO Biological Process | Response to organic cyclic compound | 2.00E-03 | 2.70E-05 |
| 125 | 5 | GO Biological Process | Tropism | 2.10E-03 | 2.87E-05 |
| 100 | 4 | KEGG Pathways | Purine metabolism | 2.60E-03 | 1.90E-04 |
| 22 | 3 | GO Biological Process | Wax metabolic process | 2.90E-03 | 4.27E-05 |
| 921 | 11 | GO Biological Process | Response to lipid | 2.90E-03 | 4.26E-05 |
| 22 | 3 | GO Biological Process | Very long-chain fatty acid biosynthetic process | 2.90E-03 | 4.27E-05 |
| 70 | 4 | GO Biological Process | Nucleoside monophosphate metabolic process | 3.20E-03 | 5.05E-05 |
| 233 | 6 | GO Biological Process | Chloroplast organization | 3.20E-03 | 4.90E-05 |
| 23 | 3 | GO Biological Process | Chloroplast accumulation movement | 3.20E-03 | 4.82E-05 |
| 941 | 11 | GO Biological Process | Regulation of biological quality | 3.30E-03 | 5.16E-05 |
| 43 | 3 | KEGG Pathways | Fatty acid biosynthesis | 3.30E-03 | 2.70E-04 |
| 24 | 3 | GO Biological Process | Chloroplast avoidance movement | 3.40E-03 | 5.42E-05 |
| 72 | 4 | GO Biological Process | Glycolytic process | 3.50E-03 | 5.61E-05 |
| 148 | 5 | GO Biological Process | Nucleoside triphosphate metabolic process | 3.80E-03 | 6.26E-05 |
| 118 | 4 | KEGG Pathways | Glycolysis / Gluconeogenesis | 3.90E-03 | 3.50E-04 |
| 1376 | 13 | GO Biological Process | Post-embryonic development | 5.00E-03 | 8.82E-05 |
| 53 | 3 | KEGG Pathways | Phenylalanine, tyrosine and tryptophan biosynthesis | 5.00E-03 | 4.80E-04 |
| 29 | 3 | GO Biological Process | Purine ribonucleoside monophosphate biosynthetic process | 5.10E-03 | 9.09E-05 |
| 29 | 3 | GO Biological Process | Cellular response to blue light | 5.10E-03 | 9.09E-05 |
| 2728 | 19 | GO Biological Process | Anatomical structure development | 5.70E-03 | 1.10E-04 |
| 273 | 6 | GO Biological Process | Response to ethylene | 6.10E-03 | 1.10E-04 |
| 62 | 3 | KEGG Pathways | Tryptophan metabolism | 7.20E-03 | 7.50E-04 |
| 6689 | 33 | GO Biological Process | Biological regulation | 8.00E-03 | 1.60E-04 |
| 16 | 2 | KEGG Pathways | Biotin metabolism | 9.80E-03 | 1.10E-03 |
| 73 | 3 | KEGG Pathways | 2-Oxocarboxylic acid metabolism | 1.00E-02 | 1.20E-03 |
| 40 | 3 | GO Biological Process | Auxin biosynthetic process | 1.11E-02 | 2.20E-04 |
| 938 | 10 | GO Biological Process | Aromatic compound biosynthetic process | 1.21E-02 | 2.40E-04 |
| 7 | 2 | GO Biological Process | AMP salvage | 1.28E-02 | 2.60E-04 |
| 45 | 3 | GO Biological Process | Response to far red light | 1.50E-02 | 3.10E-04 |
| 115 | 4 | GO Biological Process | Response to brassinosteroid | 1.51E-02 | 3.20E-04 |
| 8 | 2 | GO Biological Process | Purine ribonucleoside salvage | 1.53E-02 | 3.20E-04 |
| 50 | 3 | GO Biological Process | Glycosyl compound biosynthetic process | 1.85E-02 | 4.10E-04 |

| | | | | | |
|---|---|---|---|---|---|
| 3820 | 22 | GO Biological Process | Response to stress | 1.86E-02 | 4.20E-04 |
| 25 | 2 | KEGG Pathways | Biosynthesis of unsaturated fatty acids | 1.91E-02 | 2.50E-03 |
| 201 | 4 | KEGG Pathways | Plant-pathogen interaction | 1.91E-02 | 2.40E-03 |
| 53 | 3 | GO Biological Process | Glucose metabolic process | 2.06E-02 | 4.80E-04 |
| 54 | 3 | GO Biological Process | Red, far-red light phototransduction | 2.12E-02 | 5.10E-04 |
| 2344 | 16 | GO Biological Process | Multicellular organism development | 2.14E-02 | 5.20E-04 |
| 2594 | 17 | GO Biological Process | Multicellular organismal process | 2.18E-02 | 5.40E-04 |
| 1071 | 10 | GO Biological Process | Organic cyclic compound biosynthetic process | 2.71E-02 | 6.80E-04 |
| 13 | 2 | GO Biological Process | Response to herbicide | 2.96E-02 | 7.50E-04 |
| 37 | 2 | KEGG Pathways | Cutin, suberine and wax biosynthesis | 3.43E-02 | 5.10E-03 |
| 36 | 2 | KEGG Pathways | Arginine biosynthesis | 3.43E-02 | 4.80E-03 |
| 9473 | 43 | GO Molecular Function | Catalytic activity | 3.57E-02 | 4.47E-05 |
| 39 | 2 | KEGG Pathways | Circadian rhythm - plant | 3.60E-02 | 5.60E-03 |
| 15 | 2 | GO Biological Process | Aromatic amino acid family biosynthetic process, prephenate pathway | 3.75E-02 | 9.70E-04 |
| 15 | 2 | GO Biological Process | Root cap development | 3.75E-02 | 9.70E-04 |
| 420 | 6 | GO Biological Process | Organophosphate biosynthetic process | 4.11E-02 | 1.10E-03 |
| 5275 | 26 | GO Biological Process | Organonitrogen compound metabolic process | 4.50E-02 | 1.20E-03 |
| 3047 | 18 | GO Biological Process | Cellular aromatic compound metabolic process | 4.55E-02 | 1.20E-03 |
| 774 | 8 | GO Biological Process | Carbohydrate derivative metabolic process | 4.75E-02 | 1.30E-03 |
| 27 | 3 | GO Molecular Function | Fatty acid synthase activity | 4.77E-02 | 7.47E-05 |
| 76 | 3 | GO Biological Process | Indole-containing compound metabolic process | 4.85E-02 | 1.30E-03 |

An enrichment analysis of genes in the up-regulated gene network and down-regulated gene networks (see Fig. 2A and 2B) in *P. cineraria* seedlings using Cytoscape is shown. The Gene Ontology (GO) and KEGG PATHWAY databases were used to identify the enriched molecular function, biological pathways, and cellular components of the genes. The frequency of occurrence of a particular pathway in the set of network genes (input number) versus the frequency in the *P. cineraria* genome (background number) determined the enrichment. Enrichment categories with *P*-value < 0.05 are presented (Benjamini-Hochberg FDR). FDR, false discovery rate.

**Table S3. Hub gene identification in *Prosopis cineraria* under drought stress**

MCODE analysis results for Upregulated DEGs

| AGI Code | Gene | Description | Degree |
|---|---|---|---|
| AT3G43980 | RPS29A | 40S ribosomal protein S29 | 33 |
| AT2G09990 | RPS16A | 40S ribosomal protein S16-1 | 30 |
| AT2G07675 | rps12 | Ribosomal protein S12 | 30 |
| AT5G03850 | RPS28A | 40S ribosomal protein S28-1 | 29 |
| AT5G27700 | RPS21C | 40S ribosomal protein S21-2 | 28 |
| AT1G71760 | F14O23.14 | Uncharacterized protein | 26 |
| AT4G31985 | RPL39A | 60S ribosomal protein L39-1 | 26 |
| AT4G00100 | RPS13B | 40S ribosomal protein S13-2 | 26 |
| AT5G04800 | RPS17D | 40S ribosomal protein S17-4 | 24 |
| AT1G70600 | RPL27AC | 60S ribosomal protein L27a-3 | 24 |
| AT3G53740 | RPL36B | 60S ribosomal protein L36-2 | 24 |
| AT5G64620 | C/VIF2 | Cell wall / vacuolar inhibitor of fructosidase 2 | 23 |
| AT5G56710 | RPL31C | 60S ribosomal protein L31-3 | 23 |
| AT3G10950 | RPL37AB | Putative 60S ribosomal protein L37a-1 | 23 |
| AT2G37190 | RPL12A-2 | 60S ribosomal protein L12-1 | 23 |
| AT1G14320 | RPL10A | 60S ribosomal protein L10-1 | 23 |
| AT5G48760 | RPL13AD | 60S ribosomal protein L13a-4 | 22 |
| AT1G07660 | F24B9.25 | Histone H4 | 22 |
| AT1G26880 | RPL34A | 60S ribosomal protein L34-1 | 21 |
| AT3G05560 | RPL22B | 60S ribosomal protein L22-2 | 21 |
| AT2G28290 | SYD | Chromatin structure-remodeling complex protein SYD | 19 |
| AT3G12580 | HSP70-4 | Heat shock 70 kDa protein 4 | 18 |
| AT1G09200 | HTR2 | Histone H3.2 | 18 |
| AT4G27090 | RPL14B | 60S ribosomal protein L14-2 | 18 |
| AT4G30220 | RUXF | Probable small nuclear ribonucleoprotein F | 17 |
| AT1G16210 | F3O9.2 | Coiled-coil protein | 16 |
| AT5G52310 | RD29A | Low-temperature-induced 78 kDa protein | 16 |
| AT3G28730 | SSRP1 | FACT complex subunit SSRP1 | 15 |
| AT1G15120 | QCR6-1 | Cytochrome b-c1 complex subunit 6-1, mitochondrial | 12 |
| AT2G37470 | F3G5.26 | Histone H2B.4 | 12 |
| AT1G54690 | GAMMA-H2AX | Probable histone H2AXb | 12 |
| AT2G28720 | T11P11.3 | Histone H2B.3 | 11 |
| AT3G22590 | CDC73 | Protein CDC73 homolog | 11 |
| AT2G07727 | MT-CYB | Cytochrome b | 11 |
| AT1G61040 | VIP5 | Protein RTF1 homolog | 11 |
| AT1G05010 | ACO4 | 1-aminocyclopropane-1-carboxylate oxidase 4 | 10 |
| AT1G16030 | HSP70-5 | Heat shock 70 kDa protein 5 | 10 |
| AT5G43630 | TZP | Zinc knuckle (CCHC-type) family protein | 10 |
| AT3G26744 | SCRM | Transcription factor ICE1 | 10 |
| AT4G27670 | HSP21 | Heat shock protein 21, chloroplastic | 10 |
| AT1G65700 | LSM8 | Sm-like protein LSM8 | 10 |
| AT2G07785 | ND1 | NADH-ubiquinone oxidoreductase chain 1 | 9 |
| AT5G44500 | MFC16.18 | Small nuclear ribonucleoprotein-associated protein | 9 |
| AT3G19290 | ABF4 | ABSCISIC ACID-INSENSITIVE 5-like protein 7 | 9 |
| AT3G54560 | H2AV | Histone H2A variant 1 | 9 |
| ATMG00580 | ND4 | NADH-ubiquinone oxidoreductase chain 4 | 9 |
| AT1G07170 | F23F1.8 | PHD finger-like domain-containing protein 5A | 8 |
| ATMG00990 | ND3 | NADH-ubiquinone oxidoreductase chain 3 | 8 |
| AT3G28870 | F4J0F5_ARATH | Paired amphipathic helix SIN3-like protein | 8 |
| AT2G07741 | atp6 | ATP synthase subunit a | 8 |
| AT1G51060 | HTA10 | Probable histone H2A.1 | 8 |
| AT5G61850 | LFY | Protein LEAFY | 7 |
| AT5G26910 | TRM8 | GPI-anchored adhesin-like protein | 7 |
| AT4G21320 | HSA32 | Protein HEAT-STRESS-ASSOCIATED 32 | 7 |

MCODE analysis results for Downregulated DEGs

| AGI Code | Gene | Description | Degree |
|---|---|---|---|
| AT1G68530 | CUT1 | 3-ketoacyl-CoA synthase 6 | 8 |
| AT1G25540 | MED25 | Mediator of RNA polymerase II transcription subunit 25 | 4 |
| AT1G75100 | JAC1 | J domain-containing protein required for chloroplast accumulation response 1 | 4 |
| AT4G03020 | T4I9.10 | LEC14B homolog | 3 |
| AT3G25690 | CHUP1 | Protein CHUP1, chloroplastic | 3 |
| AT2G38280 | AMPD | AMP deaminase | 3 |
| AT4G24620 | PGI1 | Glucose-6-phosphate isomerase 1, chloroplastic | 5 |
| AT1G18270 | T10O22.24 | Ketose-bisphosphate aldolase class-II family protein | 5 |
| AT5G34930 | TYRAAT1 | Arogenate dehydrogenase 1, chloroplastic | 2 |
| AT1G15780 | MED15A | Mediator of RNA polymerase II transcription subunit 15a | 3 |
| AT2G05990 | MOD1 | Enoyl-[acyl-carrier-protein] reductase [NADH], chloroplastic | 3 |
| AT1G48030 | LPD1-2 | Dihydrolipoyl dehydrogenase 1, mitochondrial | 3 |
| AT1G01610 | GPAT4 | Glycerol-3-phosphate 2-O-acyltransferase 4 | 6 |
| AT4G00360 | CYP86A2 | Cytochrome P450 86A2 | 8 |
| AT2G46340 | SPA1 | Protein SUPPRESSOR OF PHYA-105 1 | 6 |
| AT1G09780 | PGM1 | 2,3-bisphosphoglycerate-independent phosphoglycerate mutase 1 | 2 |
| AT4G22570 | APT3 | Adenine phosphoribosyltransferase 3 | 3 |
| AT1G11860 | GDCST | Aminomethyltransferase, mitochondrial | 3 |
| AT2G16780 | MSI2 | WD-40 repeat-containing protein MSI2 | 3 |
| AT2G28230 | MED20A | Mediator of RNA polymerase II transcription subunit 20a | 3 |
| AT1G08510 | FATB | Palmitoyl-acyl carrier protein thioesterase, chloroplastic | 3 |
| AT3G45780 | PHOT1 | Phototropin-1 | 7 |
| AT1G15360 | WIN1-2 | Ethylene-responsive transcription factor WIN1 | 6 |
| AT3G12780 | PGK1 | Phosphoglycerate kinase 1, chloroplastic | 3 |
| AT1G70580 | GGAT2 | Glutamate--glyoxylate aminotransferase 2 | 4 |
| AT5G03300 | ADK2-2 | Adenosine kinase 2 | 4 |
| AT4G34460 | GB1 | Guanine nucleotide-binding protein subunit beta | 5 |
| AT1G11790 | ADT1 | Arogenate dehydratase/prephenate dehydratase 1, chloroplastic | 2 |
| AT1G17840 | ABCG11 | ABC transporter G family member 11 | 5 |
| AT1G04640 | LIP2-4 | Octanoyltransferase LIP2, mitochondrial | 2 |
| AT4G31990 | ASP5 | Aspartate aminotransferase, chloroplastic | 5 |
| AT3G23590 | MED33A | Mediator of RNA polymerase II transcription subunit 33A | 3 |
| AT5G47840 | AMK2 | Adenylate kinase 2, chloroplastic | 4 |
| AT1G67730 | KCR1 | Very-long-chain 3-oxoacyl-CoA reductase 1 | 8 |
| AT1G42550 | PMI1 | Protein PLASTID MOVEMENT IMPAIRED 1 | 5 |

Hub analysis for upregulated DEGs

| AGI Code | Gene | Degree | logfc |
|---|---|---|---|
| AT2G28290 | SYD | 19 | 5.716 |
| AT3G24500 | MBF1C | 13 | 4.207 |
| AT1G19180 | TIFY10A | 11 | 4.867 |
| AT1G61040 | VIP5 | 11 | 7.373 |
| AT1G16030 | HSP70-5 | 10 | 5.688 |
| AT1G17420 | LOX3 | 10 | 7.708 |
| AT3G26744 | SCRM | 10 | 4.517 |
| AT4G33950 | SRK2E | 7 | 7.915 |
| AT5G54160 | OMT1 | 7 | 5.136 |
| AT4G25480 | DREB1A | 6 | 10.659 |
| AT3G23240 | ERF1B | 6 | 4.212 |
| AT5G61850 | LFY | 7 | 6.484 |

Hub analysis for downregulated DEGs

| AGI Code | Gene | Degree | logfc |
|---|---|---|---|
| AT2G43010 | PIF4 | 8 | -2.681 |
| AT1G01120 | KCS1 | 5 | -2.680 |

| | | | |
|---|---|---|---|
| AT2G03870 | LSM7 | Sm-like protein LSM7 | 7 |
| AT3G09640 | APX2 | L-ascorbate peroxidase 2, cytosolic | 7 |
| AT5G40650 | SDH2-2 | Succinate dehydrogenase [ubiquinone] iron-sulfur subunit 2, mit | 7 |
| AT5G08290 | YLS8 | Thioredoxin-like protein YLS8 | 7 |
| AT1G23230 | MED23 | Mediator of RNA polymerase II transcription subunit 23 | 7 |
| AT3G25230 | FKBP62 | Peptidyl-prolyl cis-trans isomerase FKBP62 | 7 |
| AT4G17615 | CBL1 | Calcineurin B-like protein 1 | 6 |
| AT1G64670 | BDG1 | Probable lysophospholipase BODYGUARD 1 | 6 |
| AT5G24470 | APRR5 | Two-component response regulator-like APRR5 | 6 |
| AT1G66410 | CAM1 | Calmodulin-1 | 6 |
| AT2G25490 | EBF1 | EIN3-binding F-box protein 1 | 6 |
| AT5G02810 | APRR7 | Two-component response regulator-like APRR7 | 6 |
| AT2G16510 | VHA-c1 | V-type proton ATPase subunit c1 | 6 |
| AT4G25480 | DREB1A | Dehydration-responsive element-binding protein 1A | 6 |
| AT3G23240 | ERF1B | Ethylene-responsive transcription factor 1B | 6 |
| AT1G03780 | TPX2 | Protein TPX2 | 6 |
| AT1G16430 | MED22A | Mediator of RNA polymerase II transcription subunit 22a | 6 |
| AT4G10250 | HSP22.0 | 22.0 kDa heat shock protein | 6 |
| AT1G66340 | ETR1 | Ethylene receptor 1 | 6 |
| AT2G47190 | Atmyb2 | MYB transcription factor (Atmyb2) | 5 |
| AT2G07689 | T18C6.13 | NADH-Ubiquinone/plastoquinone (Complex I) protein | 5 |
| AT5G13220 | TIFY9 | Protein TIFY 9 | 5 |
| AT2G40940 | ERS1 | Ethylene response sensor 1 | 5 |
| AT1G01060 | LHY | Protein LHY | 5 |
| AT5G12840 | NFYA1 | Nuclear transcription factor Y subunit A-1 | 5 |
| AT4G39080 | VHA-a3 | V-type proton ATPase subunit a3 | 5 |
| AT1G32900 | GBSS1 | Granule-bound starch synthase 1, chloroplastic/amyloplastic | 5 |
| AT1G56170 | NFYC2 | Nuclear transcription factor Y subunit C-2 | 5 |
| AT1G23800 | ALDH2B7 | Aldehyde dehydrogenase family 2 member B7, mitochondrial | 5 |
| AT4G14540 | NFYB3 | Nuclear transcription factor Y subunit B-3 | 5 |
| AT4G27410 | NAC072 | NAC domain-containing protein 72 | 5 |
| AT3G48590 | NFYC1 | Nuclear transcription factor Y subunit C-1 | 4 |
| AT1G08980 | AMI1 | Amidase 1 | 4 |
| AT5G55160 | SUMO2 | Small ubiquitin-related modifier 2 | 4 |
| AT1G54410 | HIRD11 | Dehydrin HIRD11 | 4 |
| AT1G09530 | PIF3 | Transcription factor PIF3 | 4 |
| AT4G34880 | F11I11.120 | Probable amidase At4g34880 | 4 |
| AT2G13570 | NFYB7 | Nuclear transcription factor Y subunit B-7 | 4 |
| AT1G68480 | JAG | Zinc finger protein JAGGED | 4 |
| AT1G31360 | RECQL2 | ATP-dependent DNA helicase Q-like 2 | 4 |
| AT1G30135 | TIFY5A | Protein TIFY 5A | 4 |
| AT2G43160 | EPSIN2 | Clathrin interactor EPSIN 2 | 4 |
| ATMG00090 | RPS3 | Ribosomal protein S3, mitochondrial | 4 |
| AT5G53210 | SPCH | Transcription factor SPEECHLESS | 4 |
| AT2G38250 | GT-3B | Trihelix transcription factor GT-3b | 4 |
| AT4G32190 | F10M6.170 | Myosin heavy chain-related protein | 4 |
| AT5G66400 | RAB18 | Dehydrin Rab18 | 4 |
| AT5G12230 | MED19A | Mediator of RNA polymerase II transcription subunit 19a | 4 |
| AT4G25000 | AMY1 | Alpha-amylase 1 | 3 |
| AT2G45190 | YAB1 | Axial regulator YABBY 1 | 3 |
| AT2G42660 | F14N22.7 | Homeodomain-like superfamily protein | 3 |
| AT4G24940 | SAE1A | SUMO-activating enzyme subunit 1A | 3 |
| AT4G39130 | T22F8.30 | Dehydrin family protein | 3 |
| AT3G15680 | Q9LW11_ARATH | Zinc finger protein-like Ser/Thr protein kinase-like protein | 3 |
| AT1G64520 | RPN12A | 26S proteasome non-ATPase regulatory subunit 8 homolog A | 3 |

| | | | |
|---|---|---|---|
| AT4G00360 | CYP86A2 | 8 | -2.591 |
| AT1G15360 | WIN1-2 | 6 | -2.518 |
| AT2G38120 | AUX1 | 11 | -2.406 |
| AT1G68530 | CUT1 | 8 | -2.220 |
| AT4G31990 | ASP5 | 5 | -2.162 |
| AT1G17840 | ABCG11 | 5 | -1.838 |
| AT4G34460 | GB1 | 5 | -1.640 |
| AT2G46340 | SPA1 | 6 | -1.591 |
| AT1G42550 | PMI1 | 5 | -1.587 |

| | | | |
|---|---|---|---|
| AT2G28550 | RAP2-7 | Ethylene-responsive transcription factor RAP2-7 | 3 |
| AT2G32720 | CYTB5-B | Cytochrome b5 isoform B | 3 |
| AT1G70530 | CRK3 | Cysteine-rich receptor-like protein kinase 3 | 3 |
| AT1G14130 | F7A19.21 | 2-oxoglutarate (2OG) and Fe(II)-dependent oxygenase superfami | 3 |
| AT1G04110 | SBT1.2 | Subtilisin-like protease SBT1.2 | 3 |
| AT2G38600 | T6A23.20 | HAD superfamily, subfamily IIIB acid phosphatase | 3 |
| AT4G30960 | CIPK6 | CBL-interacting serine/threonine-protein kinase 6 | 3 |
| AT4G18240 | SS4 | Probable starch synthase 4, chloroplastic/amyloplastic | 3 |
| AT4G05400 | C6L9.80 | Copper ion binding protein | 3 |
| AT1G17730 | CHMP1A | ESCRT-related protein CHMP1A | 3 |
| AT5G55290 | VHA-e1 | V-type proton ATPase subunit e1 | 3 |
| AT5G45010 | DSS1(V) | Protein DSS1 HOMOLOG ON CHROMOSOME V | 3 |
| AT5G17770 | CBR1 | NADH--cytochrome b5 reductase 1 | 3 |
| AT1G12910 | LWD1 | WD repeat-containing protein LWD1 | 3 |
| AT5G01370 | ACI1 | ALC-interacting protein 1 | 3 |
| AT1G01260 | BHLH13 | Transcription factor bHLH13 | 3 |
| AT2G07734 | RPS4 | Ribosomal protein S4, mitochondrial | 3 |
| AT3G06120 | MUTE | Transcription factor MUTE | 3 |
| AT1G76180 | ERD14 | Dehydrin ERD14 | 2 |
| AT5G44560 | VPS2.2 | Vacuolar protein sorting-associated protein 2 homolog 2 | 2 |
| AT2G06530 | VPS2.1 | Vacuolar protein sorting-associated protein 2 homolog 1 | 2 |
| AT4G35550 | WOX13 | WUSCHEL-related homeobox 13 | 2 |
| AT5G53800 | MGN6.19 | Nucleic acid-binding protein | 2 |
| AT3G23920 | BAM1-2 | Beta-amylase 1, chloroplastic | 2 |
| AT4G17940 | T6K21.120 | Tetratricopeptide repeat (TPR)-like superfamily protein | 2 |
| AT3G26340 | PBE2 | Proteasome subunit beta type-5-B | 2 |
| AT2G33810 | SPL3 | Squamosa promoter-binding-like protein 3 | 2 |
| AT4G15880 | ESD4 | Ubiquitin-like-specific protease ESD4 | 2 |
| AT1G19980 | T20H2.28 | Cytomatrix protein-like protein | 2 |

**Table S4. Oligonucleotides used in the quantitative real-time PCR analysis**

| Oligonucleotide Name | NCBI Gene ID | *De novo* Gene ID | Oligonucleotide Sequence (5'→ 3') |
|---|---|---|---|
| PcActin_qPCR_Fw; | 129287134 | TRINITY_GG_2011_c145_g1_i5 | 5'-GTGTCCAAGTTCTTCACCCAGT-3' |
| PcActin_qPCR_Rv; | | | 5'-ACAAACCCAATCATCCATTCCACT-3' |
| 129311911_qPCR_Fw; | 129311911 | TRINITY_GG_5409_c0_g1_i3 | 5'-GCCGTCGAATATTTTGCAGGATT-3' |
| 129311911_qPCR_Rv; | | | 5'-GCTCGTCAAACTCCGTCGAC-3' |
| 129320898_qPCR_Fw; | 129320898 | TRINITY_GG_3434_c161_g1_i1 | 5'-CCTGCACAGCTTCCAAACCC-3' |
| 129320898_qPCR_Rv; | | | 5'-TGCATGCGAGAGAGAGGGAG-3' |
| 129320117_qPCR_Fw; | 129320117 | TRINITY_GG_1469_c2999_g1_i1 | 5'-AGGAATTAATCGAGATGCACGGAA-3' |
| 129320117_qPCR_Rv; | | | 5'-CTCGTCGCTACTGTCATATGCT-3' |
| 129312011_qPCR_Fw; | 129312011 | TRINITY_GG_5409_c0_g1_i1 | 5'-TGTTGATGAGCATGGCGGAG-3' |
| 129312011_qPCR_Rv; | | | 5'-CTTCACGTCATTCCAGCTCCA-3' |
| 129293970_qPCR_Fw; | 129293970 | TRINITY_GG_5322_c186_g2_i1 | 5'-AGTTGGTGATTCTTTTCATCCCAGA-3' |
| 129293970_qPCR_Rv; | | | 5'-ACAAGTGCATTTTCCCTCAGACT-3' |

Oligonucleotide primers used in the study for real-time quantitative PCR are shown. Both the NCBI gene IDs corrsponding to the United Arab Emirates cultivar of *Prosopis cineraria,* as well as the gene IDs from the *de novo* assembly of the Indian cultivar of *P. cineraria* are given. Primers were designed from the *de novo* assembled sequences.

**Table S5** Statistics of stable hydrogen bonds between the DNA binding domain of ERF group VII and VIIIa from *Arabidopsis thaliana* and *Prosopis cineraria* and the DNA in all-atom MD simulations.

| Protein | Number of H-bonds | H-bonding residues |
|---|---|---|
| *Arabidopsis thaliana* group VII (A0A5S9XC16) | 6 | A0014ARG–C0018DG<br>A0033ARG–B0006DG<br>A0038THR–B0003DG<br>A0016ARG–C0019DG<br>A0011ARG–C0015DG<br>A0057ARG–C0014DT |
| *Prosopis cineraria* group VII (XP_054790670.1) | 11 | A0033ARG–B0006DG<br>A0057ARG–C0015DG<br>A0014ARG–C0018DG<br>A0035TRP–B0003DG<br>A0012GLY–C0016DG<br>A0038THR–B0003DG<br>A0016ARG–C0019DG<br>A0011ARG–C0015DG<br>A0049TYR–C0015DG<br>A0026ARG–C0015DG<br>A0006ARG–C0016DG |
| *Arabidopsis thaliana* group VIIIa (A0A654G7S4) | 7 | A0031ARG–B0006DG<br>A0010GLY–C0016DG<br>A0009ARG–C0015DG<br>A0047TYR–C0015DG<br>A0055ARG–C0015DG<br>A0024ARG–C0015DG<br>A0014ARG–C0019DG |
| [illegible] | | A0010GLY–C0016DG<br>A0012ARG–C0018DG<br>A0047TYR–C0015DG<br>A0007ARG–C0016DG<br>A0024ARG–C0015DG |

| | | |
|---|---|---|
| *Prosopis cineraria* group VIIIa (XP_054818156.1) | 11 | A0031ARG–C0016DG<br>A0009ARG–C0015DG<br>A0018ARG–B0002DA<br>A0033TRP–B0003DG<br>A0014ARG–B0003DG<br>A0055ARG–C0015DG |

The third column details each hydrogen bond the protein (chain ‘A’) makes with the DNA double helix comprising chains ‘B’ and ‘C.’ In each entry, information for the hydrogen bond forming residue (in 3-letter code) is given on the left, and the DNA nucleotide is given on the right (DA: Adenine; DT: Thymine, DG: Guanine; DC: Cytosine). Hydrogen bonds were calculated using the HBPLUS program.
